# Shifting the selectivity of pyrido[2,3-d]pyrimidin-7(8*H*)-one inhibitors towards the salt-inducible kinase (SIK) subfamily

**DOI:** 10.1101/2023.03.24.534094

**Authors:** Marcel Rak, Roberta Tesch, Lena M. Berger, Ekaterina Shevchenko, Monika Raab, Amelie Tjaden, Rezart Zhubi, Dimitrios-Ilias Balourdas, Andreas C. Joerger, Antti Poso, Andreas Krämer, Lewis Elson, Aleksandar Lučić, Thales Kronenberger, Thomas Hanke, Klaus Strebhardt, Mourad Sanhaji, Stefan Knapp

**Affiliations:** Institute of Pharmaceutical Chemistry, Johann Wolfgang Goethe University, Max-von-Laue-Str. 9, Frankfurt am Main 60438, Germany; Structural Genomics Consortium (SGC), Buchmann Institute for Life Sciences, Johann Wolfgang Goethe University, Max-von-Laue-Str. 15, Frankfurt am Main 60438, Germany; Department of Obstetrics and Gynaecology, School of Medicine, Johann Wolfgang Goethe University, Theodor-Stern-Kai 7, Frankfurt am Main 60590, Germany; Institute of Pharmacy, Pharmaceutical/Medicinal Chemistry and Tübingen Center for Academic Drug Discovery (TüCAD2), Eberhard Karls University Tübingen, Auf der Morgenstelle 8, Tübingen 72076, Germany; School of Pharmacy, University of Eastern Finland, Yliopistonranta 1, Kuopio 70210, Finland.; German translational cancer network (DKTK) and Frankfurt Cancer Institute (FCI), Frankfurt am Main 60438, Germany

**Keywords:** Salt-inducible kinase, SIK, kinase inhibitor, MR22, pyrido[23-d]pyrimidin-7(8H)-one, MRIA9

## Abstract

Salt-inducible kinases 1-3 (SIK1-3) are key regulators of the LKB1-AMPK pathway and play an important role in cellular homeostasis. Dysregulation of any of the three isoforms has been associated with tumorigenesis in liver, breast, and ovarian cancers. We have recently developed the dual pan-SIK/group I p21-activated kinase (PAK) chemical probe MRIA9. However, inhibition of p21-activated kinases has been associated with cardiotoxicity *in vivo*, which complicates the use of MRIA9 as a tool compound. Here, we present a structure-based approach involving the back-pocket and gatekeeper residues, for narrowing the selectivity of pyrido[2,3-d]pyrimidin-7(8*H*)-one-based inhibitors towards SIK kinases, eliminating PAK activity. Optimization was guided by high-resolution crystal structure analysis and computational methods, resulting in a pan-SIK inhibitor, MR22, which no longer exhibited activity on STE group kinases and displayed excellent selectivity in a representative kinase panel. MR22-dependent SIK inhibition led to centrosome dissociation and subsequent cell-cycle arrest in ovarian cancer cells, as observed with MRIA9, conclusively linking these phenotypic effects to SIK inhibition. Taken together, MR22 represents a valuable tool compound for studying SIK kinase function in cells.

## INTRODUCTION

The salt-inducible kinases consist of three isoforms, namely SIK1 (SIK), SIK2 (QIK), and SIK3 (QSK), and are members of the group of calcium/calmodulin-dependent kinases (CAMK) [1]. As part of the liver kinase B1 (LKB1)-AMP-activated protein kinase (AMPK) pathway, they regulate cellular energy levels and stress signaling by modulating glucose and lipid metabolism based on hormone response and nutrient availability [2,3]. In addition, kinase activity is regulated through phosphorylation by protein kinase A (PKA), which serves as a recognition site for the binding of 14-3-3 proteins [4] and induces cellular translocation of the kinases, resulting in the modulation of downstream target phosphorylation [5]. SIK-dependent adaption of cellular metabolism occurs through modulation of gene expression by phosphorylation of downstream cAMP-response element binding protein (CREB)-regulated transcription coactivators 1-3 (CRTC1, CRTC2, and CRTC3) and class IIa histone deacetylases (HDACs) [6–9]. Dysregulation of SIK kinases has been correlated with the formation of several cancer types, revealing tissue-specific oncogenic and tumor suppressive roles [10]. In hepatocellular carcinoma (HCC), low SIK1 expression levels correlate with poor patient survival [11]. SIK3 is upregulated in 55% of patient-derived breast-cancer samples and its depletion sensitizes cancer cells to treatment with antimitotic drugs [10,12]. SIK2 is an oncogenic marker in ovarian cancer [13], and its overexpression promotes the formation of metastases in the adipose-rich omentum and induces cell proliferation by upregulating free fatty acid synthesis [14]. Recently, we developed the potent and highly selective chemical probe MRIA9 (**1**) [15]. MRIA9 (**1**) acts as a pan-SIK/group I PAK inhibitor displaying excellent kinome-wide selectivity (S(35) = 0.018) (Figure 1) [15]. Potent cellular on-target activity against SIK1-3 was found with IC_50_ values of 516 ± 5 nM (SIK1), 180 ± 40 nM (SIK2), and 127 ± 23 nM (SIK3) (Figure 1) [15]. Combined treatment of ovarian cancer cells with MRIA9 (**1**) and the cytostatic drug paclitaxel resulted in a significant reduction in tumor cell growth compared with single agent treatment [16]. The anti-proliferative effect of MRIA9-derived SIK2 inhibition was further investigated, revealing that MRIA9 triggered the uncoupling of the centrosomes from the nucleus during interphase and delayed the G2/M transition in SKOV3 cells by blocking the centrosome disjunction [16]. Moreover, MRIA9-dependent inhibition of SIK2 interfered with spindle assembly and orientation during mitosis, leading to aneuploidy, which promoted apoptosis [16].

**Figure 1.**
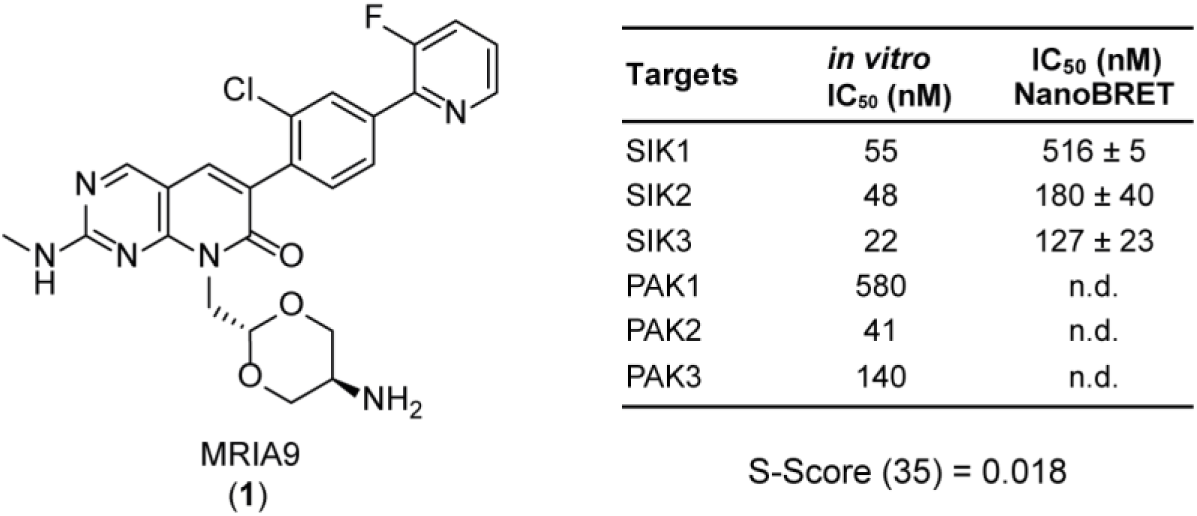
Structure and selectivity profile of dual pan-SIK/group I PAK inhibitor MRIA9 (**1**), recently published by Tesch *et al.* [15]. *In vitro* (^33^PanQinase activity assay, *Reaction Biology*) and *in cellulo* (NanoBRET) IC50 values of the main targets are shown. NanoBRET IC50 values were determined in a 10-point dose-response curve (duplicate measurements). Kinome-wide selectivity was determined on 443 kinases (^33^PanQinase assay panel; *Reaction Biology*) at a concentration of 1 µM.

The basic pyrido[2,3-d]pyrimidin-7(8*H*)-one scaffold of MRIA9 (**1**) has already been exploited for the development of multiple PAK inhibitors, including the lead structure G-5555 (**2**) [17–20]. However, inhibition of group I PAKs has been associated with *in vivo* lethality, due to cardiotoxicity [20]. Several promiscuous compounds commonly targeting group I PAK kinases have been tested in a tolerability study at a dosage of 12.5–300 mg/kg (po and/or ip), showing a direct correlation between the potency of the compounds against PAK1/PAK2 and the dosage at which lethal effects have been observed. A Langendorff heart assay using G-5555 (**2**), for example, revealed a dose-dependent correlation with the occurrence of cardiac arrhythmia and heart failure [20]. Although PAK1 function has previously been linked to cardiac diseases, including arrhythmias caused by impaired Ca^2+^ homeostasis, its knockout did not prove lethal in mice [21,22]. Therefore, the effects observed after compound administration may be induced by PAK2 inhibition as knockout of PAK2 leads to lethality in mice at an embryonic developmental stage and blocks the formation of the vasculature [22]. Importantly, the compounds used in the above-mentioned tolerability study were not tested for their kinome-wide selectivity, and further studies are needed to conclusively assess the effects of PAK inhibition, especially because no lethality was described in a pharmacokinetic (PK) study of G-5555 (**2**), administering 2 mg/kg (po) and 25 mg/kg (ip) per dosage [19].

Overall, these data highlight that there is a need for more selective inhibitors for studying SIK kinase function in cells. Recently, a pan-SIK inhibitor, SK-124, has been reported to show good *in vivo* activity stimulating bone formation and a good kinome-wide selectivity with, however, PDGFRα as the main off-target [23]. Here, we have used a structure-guided approach exploiting differences in the back pocket of CAMK and STE group kinases to shift the selectivity of the pyrido[2,3-d]pyrimidin-7(8*H*)-one inhibitors MRIA9 (**1**) and G-5555 (**2**) towards SIK kinases and to eliminate group I PAK activity.

## RESULTS AND DISCUSSION

### Structure-based strategy for modulating the selectivity of pyrido[2,3-d]pyrimidin-7(8*H*)-one derived compounds

We performed a systematic structural analysis of STE group PAK kinases (PAK1, PAK2, and PAK3) and mammalian STE20-like kinases (MST3 and MST4) to guide the optimization of pyrido[2,3-d]pyrimidin-7(8*H*)-one inhibitors towards SIK kinase selectivity. Both kinase subfamilies share a high sequence similarity between their isoforms, with more than 93% sequence identity in the kinase domain of PAK1 compared with PAK2 and PAK3, and about 90% sequence conservation between the kinase domains of MST3 and MST4. We decided to focus on the structurally best-characterized isoforms PAK1 and MST3 for further analysis. No crystal structures of SIK isoforms are yet available, and it remains challenging to obtain stable SIK2 protein for crystallization studies. We, therefore, used our established homology model of SIK2 [15], based on the reported canonical binding mode of G-5555 (**2**) in MST4, as a surrogate model for the kinase domain of all three highly conserved SIK isoforms (78% and 68% sequence conservation when comparing SIK1 with SIK2 and SIK3, respectively). A key structural feature that can be exploited for improving the selectivity of inhibitors is the gatekeeper residue, which modulates specific access to the back pocket of kinases. The most common amino acids at this position are methionine (39%), threonine (19%), leucine (18%), or phenylalanine (14%) [24]. In addition, the overall distribution of gatekeeper amino acids differs between different kinase groups, with a methionine residue found most frequently in the STE and CAMK group kinases, as well as in the group of protein kinases A, G, and C (AGC) [24]. In contrast, a threonine residue is most frequent in tyrosine (TK) and tyrosine-like kinases (TLK) [24]. Remarkably, all three SIK isoforms feature a small threonine gatekeeper residue, despite their phylogenetic classification as CAMK kinases, based on overall sequence similarity. This represents a major difference compared with the STE group PAKs and MSTs, which all feature a methionine at this site. When analyzing the structures of the kinase domains of SIK2, PAK1, and MST3, a unique subpocket was found that extends the back pocket next to the gatekeeper in SIK2. This significant structural difference led us to hypothesize that targeting the gatekeeper region would be beneficial for eliminating STE group kinase activity. We therefore systematically analyzed the co-crystal structures of kinases harboring a small threonine gatekeeper. The structure of human leucine zipper- and sterile alpha motif-containing kinase (ZAK) in complex with vemurafenib (**3**) (PDB code 5HES) [25] was particularly interesting in this context. Vemurafenib (**3**) is an FDA-approved kinase inhibitor developed to specifically target the BRAF V600E mutant in metastatic melanoma [26]. Follow-up studies, however, revealed significant off-target inhibition of ZAK, with an IC_50_ value of 4.0 nM [27]. Comparison of the structures of ZAK and SIK2 with PAK1 and MST3 showed that the kinases with a small gatekeeper residue (Thr82 and Thr96 for ZAK and SIK2, respectively) not only present a facilitated access to the back pocket but also share an overall larger pocket (Figure 2). In the case of ZAK, the back pocket is occupied by the aliphatic sulfonamide moiety of vemurafenib (**3**). The kinases with a larger methionine gatekeeper (Met344 and Met111 for PAK1 and MST3, respectively) showed no significant extension of the back pocket in that region (Figure 2), and, accordingly, vemurafenib (**3**) stabilized neither PAK1 nor MST3 in a differential scanning fluorimetry (DSF) assay.

**Figure 2.**
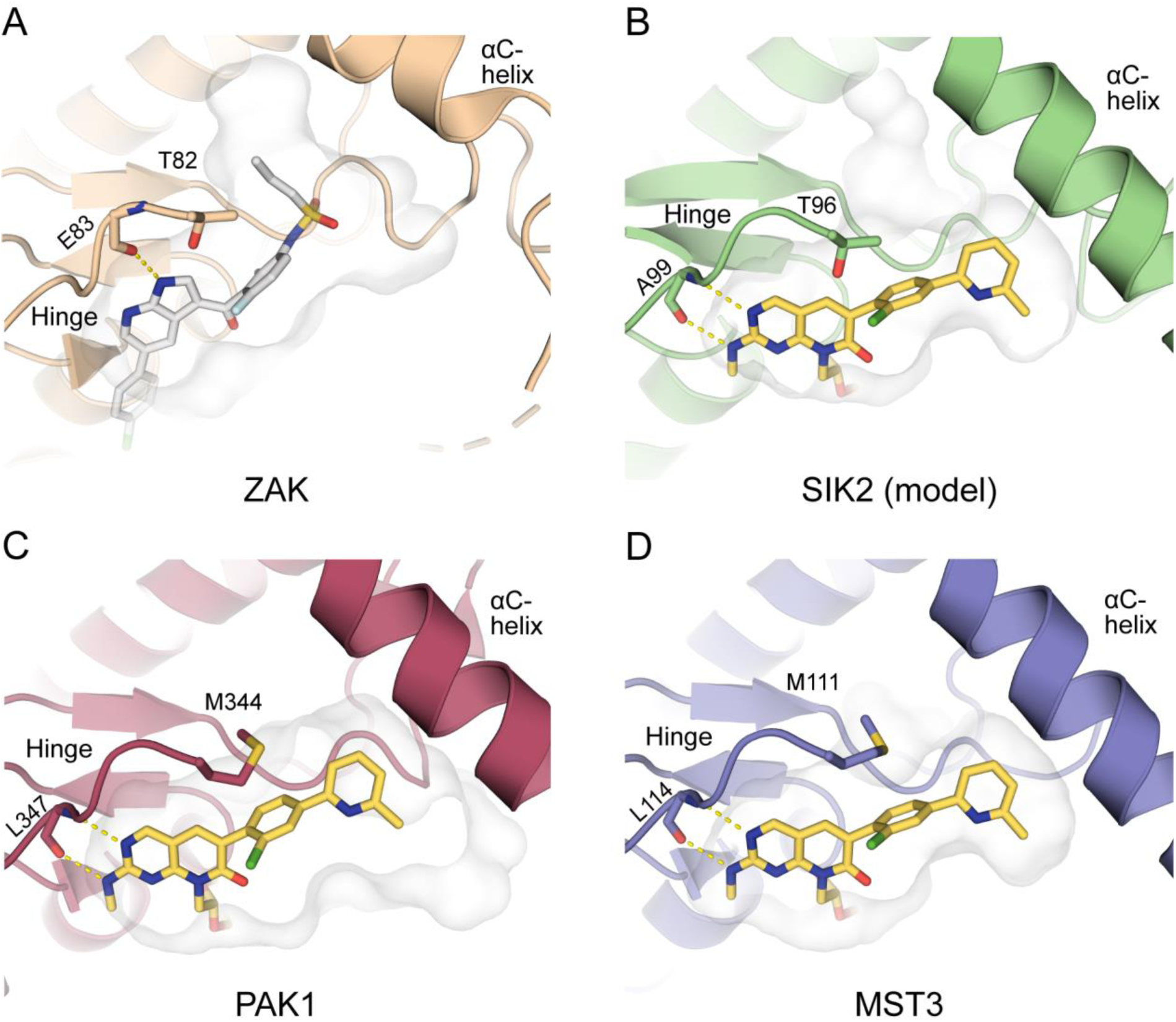
Comparison of the back-pocket region in the structure of ZAK in complex with vemurafenib (**3**) (PDB code 5HES; light brown; A) and of SIK2 (homology model; green; B), PAK1 (PDB code 5DEY; red; C), and MST3 (PDB code 7B30; blue; D) in complex with G-5555 (**2**). The surface is shown in light gray. The gatekeeper and hinge-binding amino acids are shown as stick models. Hydrogen bonds are shown as dashed yellow lines.

We further analyzed the differences in amino acids lining the back pocket next to the gatekeeper to guide our inhibitor design by also focusing on the amino acids at +4 of the αC-helix and the loop between the αC-helix and β4-strand as well as on the DFG motif and its -1 amino acid (Figure 3). Our previous studies had shown that the amino acid at the +4 position of the αC-helix limits the tolerance of substituents at position 3 of the pendant pyridine ring of G-5555 (**2**) in SIK2 [15]. In addition, the residue at this position modulates the accessibility of the back pocket next to the gatekeeper. PAK1 and SIK2 share a larger methionine residue (Met319 and Met71, respectively) at this site, whereas a smaller leucine (Leu86) is found in MST3.

**Figure 3.**
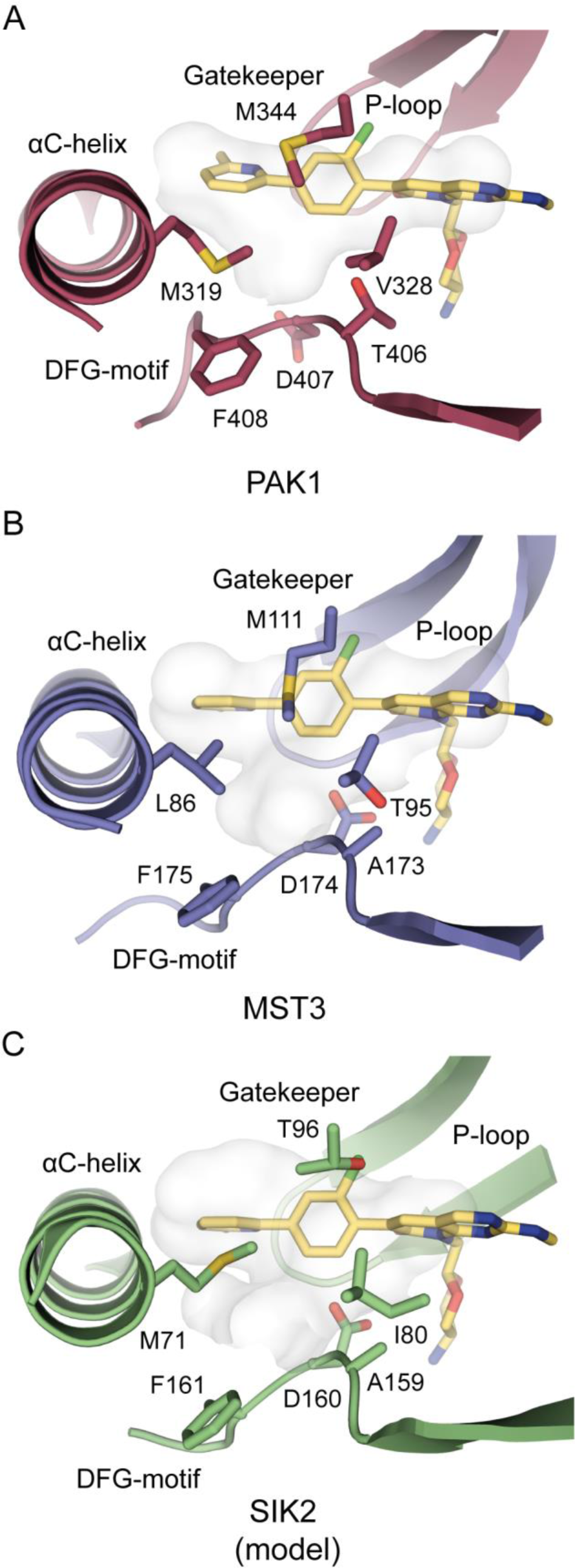
Comparison of the back pocket of PAK1 (PDB code 5DEY; red; A), MST3 (PDB code 7B30; blue; B), and SIK2 (model; green; C) in complex with G-5555 (**2**). The surface of the binding pocket in and around the area of the back pocket is shown. Important amino acids and structural motifs are highlighted.

Entrance to the back pocket is further modulated by an amino acid of the loop between the αC-helix and the β4-strand. PAK1 harbors a valine residue (Val328), MST3 a threonine (Thr95), and SIK2 an isoleucine residue (Ile80) at this position. Finally, this part of the pocket is also lined by amino acids of the DFG motif and its adjacent -1 residue. Interestingly, the loop containing the DFG motif was found to be oriented closer toward the binding pocket in PAK1 compared with MST3 and the model of SIK2. In addition, both SIK2 and MST3 share an alanine residue (Ala159 and Ala173, respectively) at position -1 of the DFG motif, whereas PAK1 has a larger threonine (Thr406). Overall, the back-pocket region differs significantly between the CAMK and STE group kinases due to key differences in the amino acids lining the entrance of the pocket, which can be exploited by modifying the substituents on the phenyl ring of MRIA9 (**1**) or G-5555 (**2**). We, therefore, decided to change the substitution pattern of the 2-chloro-4-(6-methylpyridyl)phenyl ring from position 4 to 5 to target a different region of the back pocket, as shown in Figure 4 in an overlay of G-5555 (**2**) and vemurafenib (**3**).

**Figure 4.**
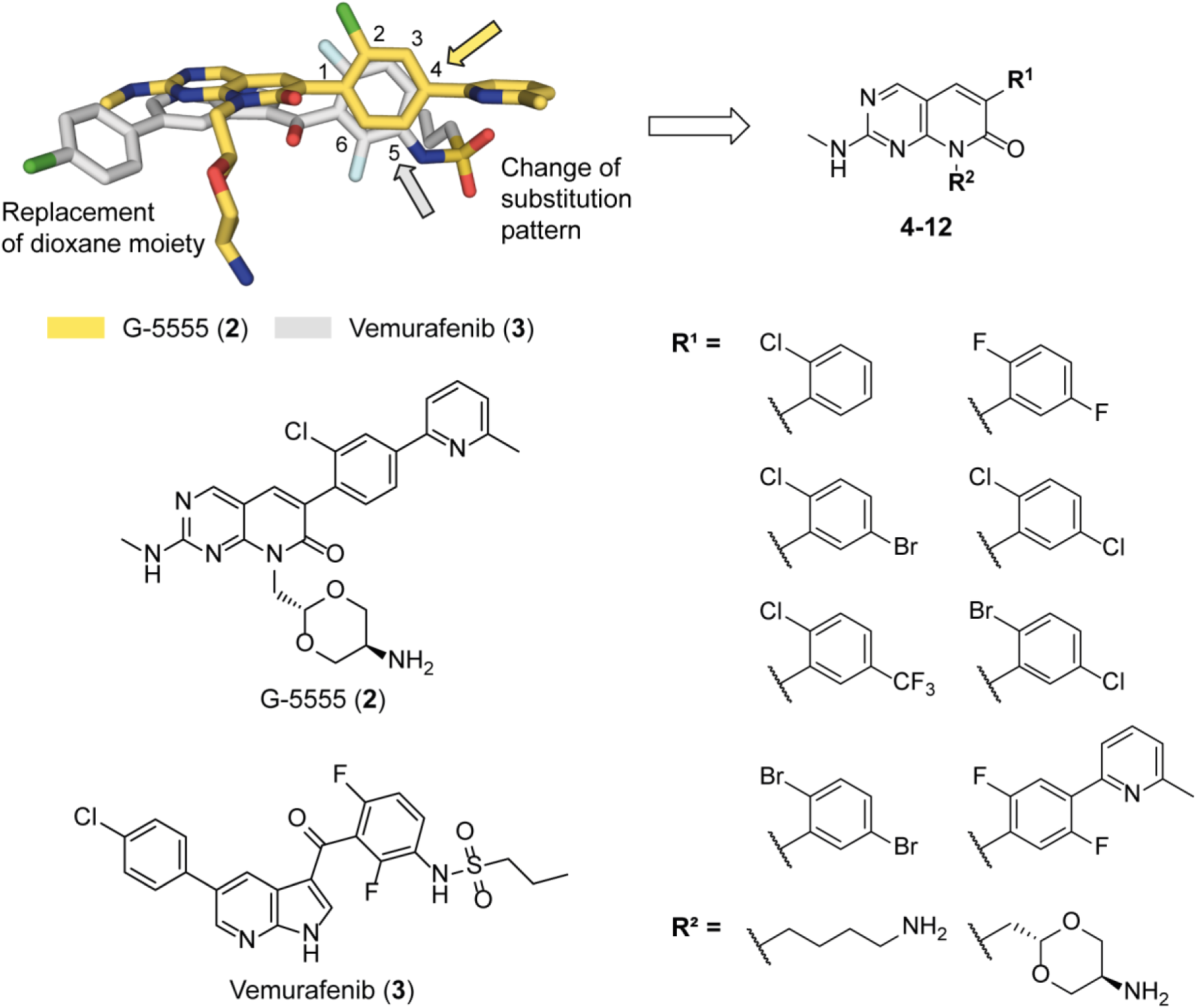
Design idea for a focused series of pyrido[2,3-d]pyrimidin-7(8*H*)-one-based inhibitors targeting unique features of the back pocket of SIK2. Modifications of the lead structure are highlighted in the comparison of the structures of G-5555 (yellow; **2**) and vemurafenib (gray; **3**). Modifications were performed at R^1^ and R^2^. Various halogenated aromatic rings (position 2 and/or 5 of the phenyl ring) were introduced at position R^1^. For R^2^, either a (2*r*,5*r*)-2-methyl-1,3-dioxan-5-amine group or a 4-aminobutyl group was chosen. For combinations of R^1^ and R^2^ see Table 1.

The terminal pyridine ring in G-5555 (**2**) was removed to investigate the effect on the selectivity and potency of the derivatives with respect to SIK2. Halogen atoms (F, Br and Cl) and a CF_3_-group were introduced in different combinations at positions 2 and 5 of the phenyl ring to exploit differences in the gatekeeper region and to specifically impair the inhibition of STE group kinases. For group R^2^, which interacts with the DFG motif and catalytic-loop amino acids, the (2*r*,5*r*)-2-methyl-1,3-dioxan-5-amine group of MRIA9 (**1**)/G-5555 (**2**) was used. In addition, a 4-aminobutyl group was introduced instead of the dioxane moiety, which was anticipated to be beneficial for modulating the overall potency of the lead structure by strengthening the interactions formed with amino acids of the catalytic loop and the DFG motif. Only one derivative was synthesized with a 6-methylpyridyl group in the back pocket. For this, the most promising phenyl ring decoration was chosen to investigate its influence on selectivity.

### Synthesis of compounds targeting differences in the back pocket

A targeted series of nine compounds was synthesized, based on the synthetic route previously published by Tesch *et al.* [15] and the route of Rudolph *et al.* [20]. For compounds **4**–**6**, the synthetic route consisted of six (Scheme 1), and for compounds **7**–**11** it consisted of five steps (Scheme 2). Compound **12** was synthesized in eight steps according to Scheme 3. The first reaction step for all compounds except for **4** was a Fischer esterification of the acetic acid starting materials **14**–**19** to obtain the corresponding ethyl acetate derivatives. In the next step, a cyclocondensation reaction of **20**–**26** with 4-amino-2-(methylthio)-pyrimidine-5-carbaldehyde was performed, leading to the formation of the pyrido[2,3-d]pyrimidin-7(8*H*)-one scaffold in **27**–**33**. Intermediates **27**–**29** were then used in a following nucleophilic substitution (S_N_) reaction with *tert*-butyl (4-bromobutyl)carbamate, leading to **34**–**36**. To implement the hinge binding moiety, the methyl sulfide group was first oxidized to a mixture of the corresponding sulfoxide and sulfone product using 3-chloroperbenzoic acid (*m*-CPBA) and further converted in a nucleophilic aromatic substitution (S_N_Ar) using methylamine (**37**–**39**). In the last step, a deprotection of the BOC-protecting group was performed, using trifluoroacetic acid (TFA) to obtain compounds **4**–**6** (Scheme 1).

**Scheme 1.**
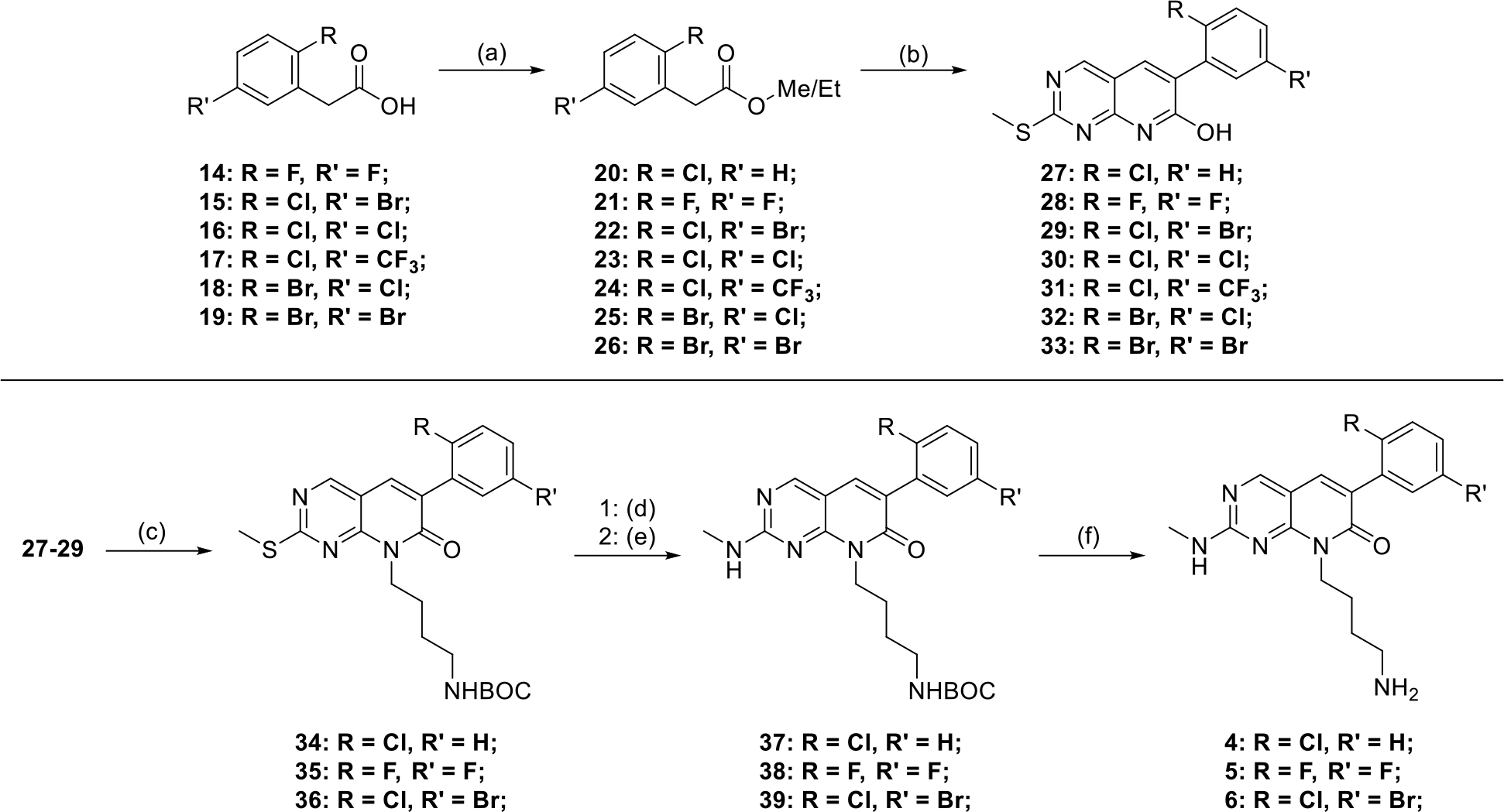
Synthesis of compounds **4**–**6**. Reagents and conditions: (a) H2SO4, ethanol, 100 °C, 15 h; (b) 4-amino-2-(methylthio)-pyrimidine-5-carbaldehyde, K2CO3, DMF, 120 °C, 16 h; (c) *tert*-butyl (4-bromobutyl)carbamate, Cs2CO3, DMF, 110 °C, 18 h; (d) 3-chloroperbenzoic acid (*m*-CPBA), DCM, RT, up to 3 h; (e) methylamine, *N,N’*-diisopropylethylamine (DIPEA), ethanol, 85 °C, 17 h; (f) TFA (20 vol%), DCM, RT, 0.5 h.

**Scheme 2.**
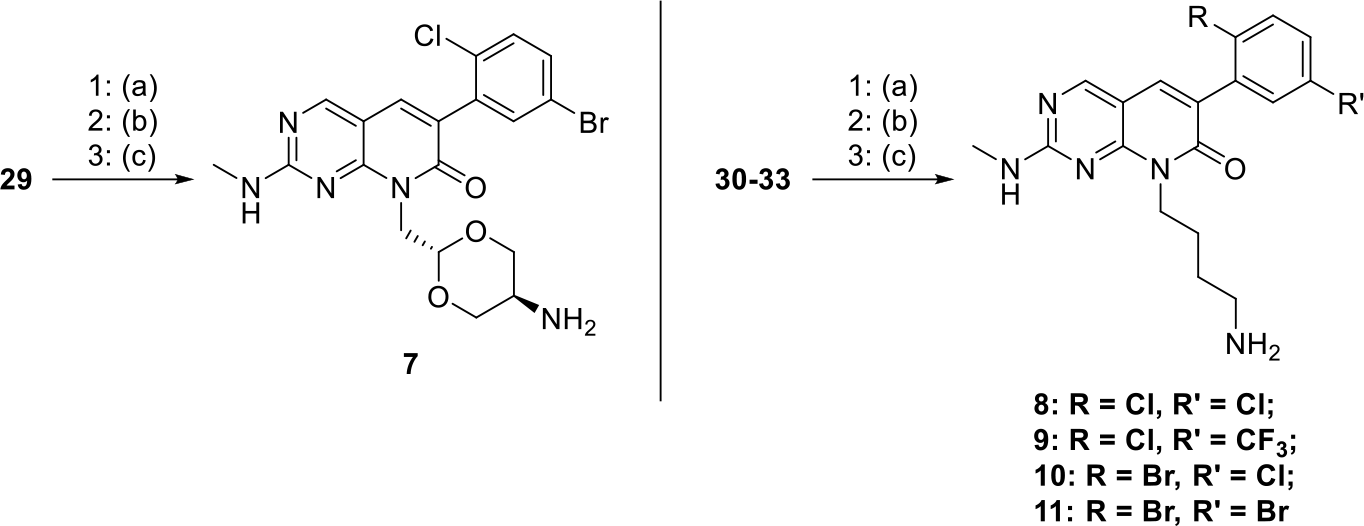
Synthesis of compounds **7**–**11**. Reagents and conditions: (a) 2-((2*r*,5*r*)-2-(bromomethyl)-1,3-dioxan-5-yl)isoindoline-1,3-dione or 2-(4-bromobutyl)isoindoline-1,3-dione, Cs2CO3, DMF, 110 °C, 18 h; (b) *m*-CPBA, DCM, RT, up to 3 h; (c) methylamine, DIPEA, ethanol, 85 °C, 17 h.

For intermediates **29**–**33**, various phthalimide-protected aliphatic amine moieties were used in the following S_N_ reaction. 2-((2*r*,5*r*)-2-(bromomethyl)-1,3-dioxan-5-yl)isoindoline-1,3-dione, used in the reaction with **29**, was synthesized as previously described [15]. In the case of **30**–**33**, 2-(4-bromobutyl)isoindoline-1,3-dione was used in this reaction. For all phthalimide-protected intermediates, the formation of a mixture of the corresponding isoindoline-1,3-dione and 2-carbamoylbenzoic acid derivative was observed, due to partial cleavage of the protecting group. This mixture was used in the following reactions without further purification. The final steps were the oxidation of the methyl sulfide to a mixture of the sulfoxide/sulfone-product using *m*-CPBA, followed by an S_N_Ar reaction with methylamine to obtain compounds **7**–**11** (Scheme 2).

For compound **12**, a Miyaura borylation reaction was carried out after the esterification reaction in the first step (**41**) to implement a boron pinacol ester at position four of the phenyl ring. A subsequent Pd-catalyzed Suzuki reaction of **42** with 2-bromo-6-methylpyridine led to intermediate **43**. The final steps of this synthetic route were similar to those already described in Scheme 1. The pyrido[2,3-d]pyrimidin-7(8*H*)-one scaffold **44** was formed in a cyclocondensation reaction of **43** with 4-amino-2-(methylthio)-pyrimidine-5-carbaldehyde, followed by the S_N_-reaction with *tert*-butyl (4-bromobutyl)carbamate, resulting in **45**. The methyl sulfide was oxidized using *m*-CPBA, followed by conversion of the sulfoxide/sulfone-product in the S_N_Ar using methylamine under basic conditions, and the acidic deprotection of the BOC-protected amine **46** to obtain compound **12** (Scheme 3).

**Scheme 3.**
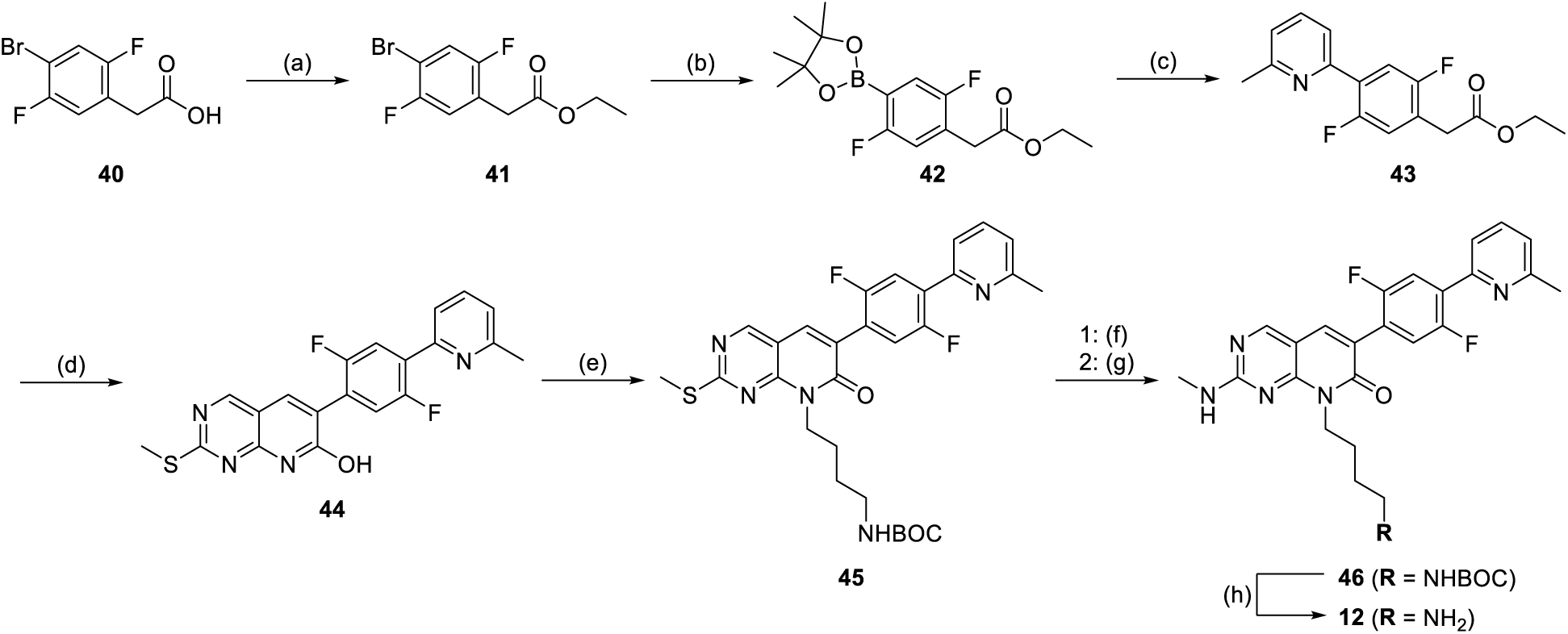
Synthesis of compound **12**. Reagents and conditions: (a) H2SO4, ethanol, 100 °C, 15 h; (b) bis(pinacolato)diboron, [1,1×-bis(diphenylphosphino)ferrocene]dichloropalladium(II) (Pd(dppf)Cl2), KOAc, dioxane, 115 °C, 18 h; (c) 2-bromo-6-methylpyridine, Pd(dppf)Cl2, KOAc, dioxane/H2O (2:1), 107 °C, 18 h; (d) 4-amino-2-(methylthio)pyrimidine-5-carbaldehyde, K2CO3, DMF, 120 °C, 16 h. (e) *tert*-butyl (4-bromobutyl)carbamate, Cs2CO3, DMF, 110 °C, 18 h; (f) *m*-CPBA, DCM, RT, up to 3 h; (g) methylamine, DIPEA, ethanol, 85 °C, 17 h; (h) TFA (20 vol%), DCM, RT, 0.5 h.

### Structure-activity relationship (SAR) of pyrido[2,3-d]pyrimidin-7(8*H*)-one-based inhibitors 4**–**12

Due to the lack of recombinant SIK1-3 protein, on-target IC_50_ values were measured using a cellular target engagement NanoBRET assay (Table 1). For SIK1, IC_50_ values were determined in lysed-mode measurements. Correlation plots of all three SIK assays performed in lysed and intact modes revealed good agreement of the data (Figure S1). Off-target activity against kinases of the STE group was assessed in a differential scanning fluorimetry (DSF) assay against MST1, MST2, MST3, MST4, and PAK1 (Table 1). The overall selectivity of the compounds synthesized in this work was tested against a panel of up to 100 kinases in a subsequent DSF assay (Table 1). Representative kinases from each branch of the phylogenetic kinome tree were included in this panel to examine the stabilization induced by the compounds and to estimate kinome-wide selectivity (Figure S2). All kinases included are listed in Table S1. In addition, a comparison of our internal DSF panel with the scanMAX assay panel against 468 kinases (*Eurofins Scientific*) revealed an overlap of 90 kinases (Figure S2). Compounds staurosporine, MRIA9 (**1**), and G-5555 (**2**) were added as controls (Table 1).

**Table 1.**
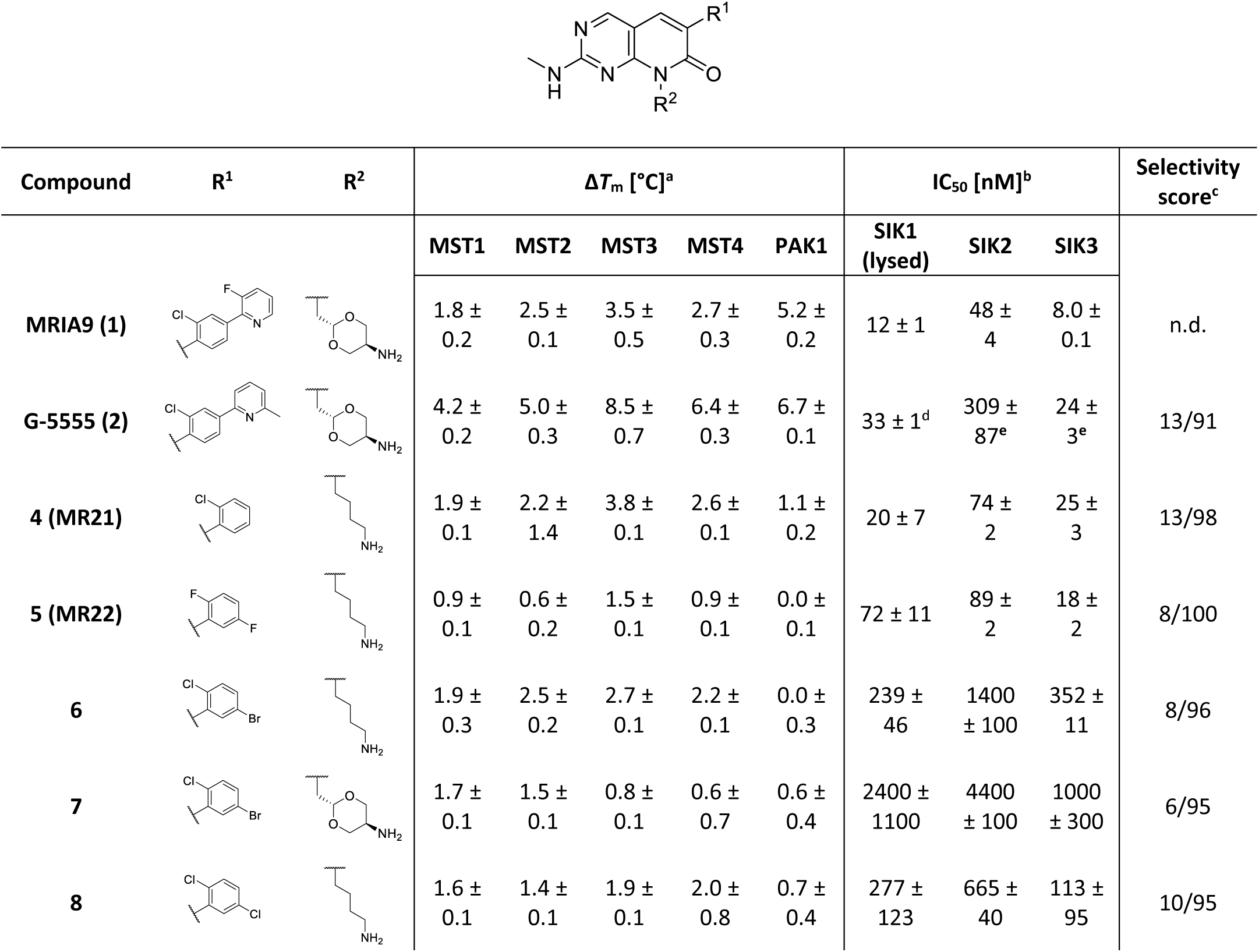
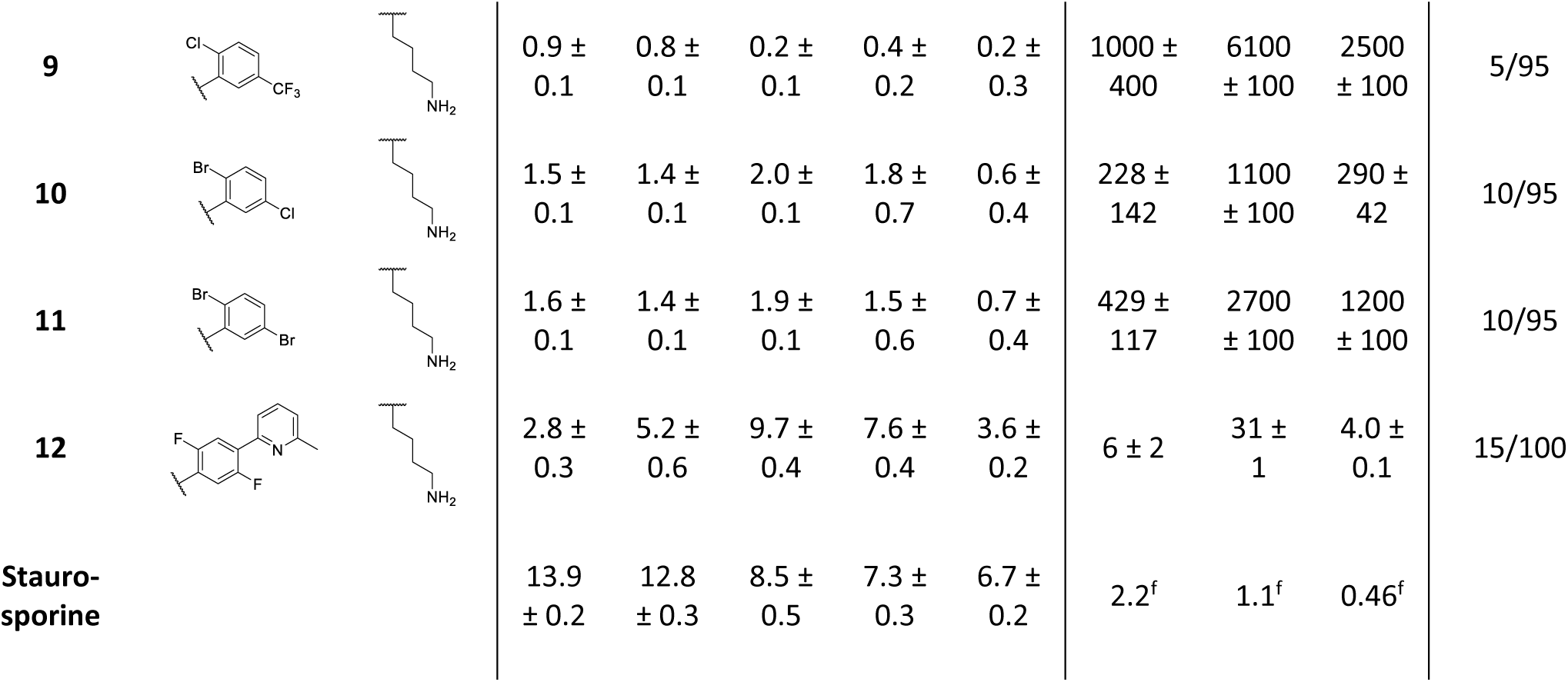
Selectivity profile of pyrido[2,3-d]pyrimidin-7(8*H*)-one-based inhibitors as pan-SIK inhibitors. ^a^Average of three measurements and the standard deviation of the mean. Negative values are shown as 0.0 °C. ^b^IC50 values with standard deviation of the mean were determined using the NanoBRET assay in a 11-point dose-response curve (duplicate measurements). SIK1 was measured in lysed and SIK2 and SIK3 in intact mode. The curves for the measurement of the lysed mode can be found in Figure S1. ^c^Selectivity score was determined measuring the Δ*T*m [°C] against a maximal number of 100 kinases. Two measurements were performed and the average was calculated. Only kinases with a *ΔT*m value higher than 50% compared with a corresponding reference compound were considered off-targets. Data, including reference compounds, can be found in Figure S7 and Tables S1 and S2. Compounds were tested at a final concentration of 10 µM. ^d^IC50 value of G-5555 (**2**) against SIK1 measured in intact mode. ^e^Published IC50 values of G-5555 (**2**) [15]. ^f^Literature Kd values for staurosporine on SIK1, SIK2, and SIK3 [28,29].

The obtained Δ*T*_m_ and IC_50_ values correlated well with published data. MRIA9 (**1**) induced a Δ*T*_m_ = 3.5 ± 0.5 °C against MST3, compared with a published value of Δ*T*_m_ = 4.0 ± 0.4 °C [15]. Surprisingly, the on-target activity of MRIA9 (**1**) was found to be slightly increased for all three SIK isoforms, with IC_50_ values of 12 ± 1 nM, 48 ± 4 nM, and 8.0 ± 0.1 nM against SIK1 (lysed), SIK2, and SIK3, respectively. The data obtained for G-5555 (**2**) was also consistent with literature values, with a Δ*T*_m_ value of 6.7 ± 0.1 °C against PAK1 *vs* 6.6 ± 0.2 °C in the literature [15]. In our internal selectivity panel against 91 kinases, 13 kinases were stabilized by G-5555 (**2**) by more than 50% compared to a corresponding reference compound. These data correlated well with the published scanMAX selectivity data against 468 kinases obtained at a concentration of 1 µM [15], highlighting the suitability of the in-house DSF panel for estimating kinome-wide selectivity (Figure S3). Removal of the 6-methylpyridine group in the back-pocket region and the substitution of the (2*r*,5*r*)-2-methyl-1,3-dioxan-5-amine group of G-5555 (**2**) with a 4-aminobutyl group in **4** (MR21) led to a dramatic decrease in stabilization of STE group kinases (MSTs and PAKs). For MST3, the Δ*T*_m_ value decreased to about half the shift of G-5555 (**2**), and only a low Δ*T*_m_ of 1.1 ± 0.2 °C was found for PAK1. Surprisingly, cellular on-target potency against SIK kinases increased compared with G-5555 (**2**), with IC_50_ values of 20 ± 7 nM (SIK1; lysed), 74 ± 2 nM (SIK2), and 25 ± 3 nM (SIK3). Moreover, the selectivity score also improved to 13/98 kinases.

Interestingly, **4** (MR21) also showed activity towards ephrin kinase A2 (EpHA2), which prompted us to determine the crystal structure of EpHA2 in complex with **4**. The superposition of this structure with the published structure of **2** (G-5555) in MST3 (Figure S4) showed that the kinase domain and the main scaffolds of both compounds superimposed well, with variations found mainly in flexible regions. Interestingly, an up- and outward movement of the αC-helix was observed in the structure of **4** (MR21) in EpHA2, which enabled the formation of the conserved salt bridge between Lys646 on the β3-strand and Glu663 on the αC-helix. In the MST3 complex, due to a different αC-helix orientation, the 6-methylpyridine group nitrogen of the back-pocket motif of **2** (G-5555) was inserted into the conserved salt bridge, forming a direct hydrogen bond with Lys65. A similar reorientation of the αC-helix of EpHA2 upon binding of G-5555 (**2**) would lead to a clash of the methyl group in position 6 of the terminal pyridine ring with Phe660 in the αC-helix, providing an explanation for the lack of potent stabilization of ephrin kinases by G-5555 (**2**) in Δ*T*_m_ assays. In MST3, a smaller residue (Ile79) is found at this site. Alteration of the hydrogen-bond network in the back pocket may also contribute to the impaired STE group activity.

For **5** (MR22), the addition of fluorine atoms at positions 2 and 5 of the phenyl ring led to a significant on-target potency, with IC_50_ values of 72 ± 11 nM, 89 ± 2 nM, and 18 ± 2 nM against SIK1 (lysed), SIK2, and SIK3, respectively. However, the stabilization of off-target kinases was further decreased, with the highest Δ*T*_m_ shift of 1.5 ± 0.1 °C against MST3 and no stabilization at all for PAK1. The selectivity determined against 100 kinases increased, with only 8 kinases stabilized by more than 50% compared to a corresponding reference compound. No significant effect on the off-target profile was observed when replacing the substituents with a chlorine at position 2 and a bromine at position 5 (compound **6**). However, the activity against SIK kinases decreased, especially against SIK2, with an IC_50_ value of 1400 ± 100 nM. Substitution of the 4-aminobutyl moiety by a (2*r*,5*r*)-2-methyl-1,3-dioxan-5-amine group in **7** blocked the inhibition of all three SIK isoforms, with the lowest IC_50_ value found against SIK3 of 1 μM. In addition, the overall selectivity increased, with only 6/95 kinases stabilized, and no potent inhibition of STE group kinases. Due to the low on-target activity of this compound, the next series of compounds were synthesized again with the 4-aminobutyl group. In **8**, the bromine substituent at position 5 of compound **6** was replaced by a chlorine, resulting in partial restoration of activity against all three SIK isoforms. IC_50_ values of 277 ± 123 nM (SIK1; lysed), 665 ± 40 nM (SIK2), and 113 ± 95 nM (SIK3) were found, indicating that the smaller chlorine halogen can be better accommodated in the pocket facing the gatekeeper residue. The selectivity score slightly increased to 10/95 kinases. A bulkier CF_3_-group at position 5 (**9**) proved to be detrimental to the inhibition of kinases of the STE and CAMK groups, as indicated by the overall low Δ*T*_m_ and the high IC_50_ values of more than 1 μM. **9** was also co-crystalized with EpHA2, and the structure of the complex was compared with that of **4** (MR21) in EpHA2 to elucidate the influence of the CF_3_-group on the binding mode (Figure S5). The superposition of the two structures revealed an identical overall orientation of the compounds in the binding pocket, with only minor shifts in the orientation of amino acid side chains in the back-pocket region, upon occupation by the CF_3_-group.

Compounds **10** and **11** harboring a bromine at position 2 and either a chlorine or a bromine atom at position 5, respectively, displayed reduced off-target activity against the MSTs and PAK1, as found for all compounds with truncated back-pocket motif. Of particular interest, in compound **12**, the 6-methylpyridine group of G-5555 (**2**) was reattached onto the 2,5-difluorophenyl ring of **5** (MR22) to evaluate the effect on the overall potency. This addition increased the activity on the MST kinases, with a Δ*T*_m_ value of 9.7 ± 0.4 °C on MST3 compared with 8.5 ± 0.7 °C for G-5555 (**2**). In the case of PAK1, the 6-methylpyridyl group did not restore stabilization to the same extent, and the Δ*T*_m_ shift was only about half of that for G-5555 (**2**) (Δ*T*_m_ = 3.6 ± 0.2 °C *vs* 6.7 ± 0.1 °C, respectively). This argues for unfavorable interaction of the compound in and around the back pocket next to the gatekeeper. For the SIK kinases, increased activity was found, with IC_50_ values of 6 ± 2 nM (SIK1; lysed), 31 ± 1 nM (SIK2), and 4.0 ± 0.1 nM (SIK3). The selectivity was slightly improved with 15/100 kinases *vs* 13/91 kinases for G-5555 (**2**). Because of the strong stabilization of MST3 protein by **12**, we determined the crystal structure of the complex and compared the binding mode of **12** with that of **2** (G-5555) (Figure S6). While the two main scaffolds of **12** and **2** formed identical hinge-binding interactions with Leu114, a different orientation of the phenyl ring could be observed. The fluorine atoms at positions 2 and 5 induced a 30° rotation of the phenyl ring, with the fluorine at position 5 pointing toward the subpocket next to the gatekeeper. Notably, PAK1, which has no pronounced subpocket in that region, showed the lowest thermostabilization upon treatment with **12**.

Taken together, we were able to eliminate the commonly observed activity of the pyrido[2,3-d]pyrimidin-7(8*H*)-one main scaffold toward STE group kinases (MSTs and PAKs) by removing the terminal pyridine group interacting with the conserved salt bridge in the back pocket. However, even with the terminal pyridine group attached, the incorporation of a substituent at position 5 of the phenyl ring (as in **12**) significantly lowered the stabilization of PAK kinases. In addition, introducing substituents at position 5 of the phenyl ring was sufficient to effectively target the subpocket next to the gatekeeper, as observed in the crystal structure of EpHA2 with bound **9**. Importantly, different combinations of the substituents at position 2 and 5 of the phenyl ring modulated the activity against SIK kinases, and the highest potency was observed for small halogen atoms, as in **4** (MR21), **8**, and **5** (MR22).

### The water network in the back pocket of SIK2 influences the binding energy of MR22 (5)

To investigate the influence of the water network on the binding of compound MR22 (**5**), we predicted the thermodynamic profile of the hydration sites within the binding pocket, using WaterMap calculations. Explicit water molecules in molecular dynamic (MD) simulations of SIK2 were clustered into hydration sites, and free energy, enthalpy, and entropy properties were calculated (Figure 5).

**Figure 5.**
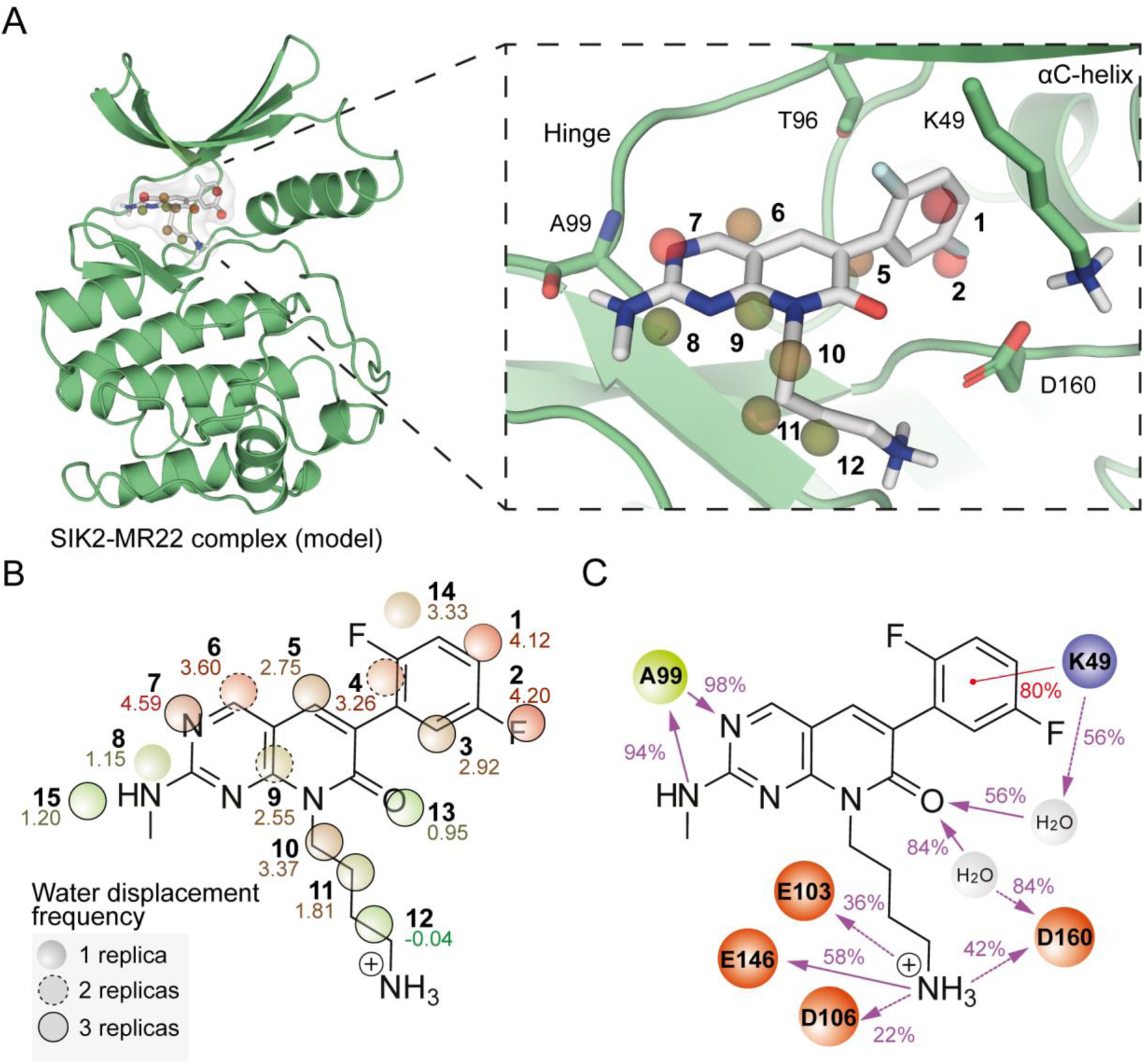
Comparison of the stabilization of **5** (MR22) in SIK2 (model) during MD simulations and evaluation of the thermodynamic water profile. (A) WaterMap simulation of MR22 (**5**; gray) in the binding pocket of SIK2 (homology model; green), highlighting favorable (green) and unfavorable (red) hydration sites during the simulation and their location relative to bound MR22 (**5**). The binding pocket is shown as a white surface. Hydration sites are presented as colored spheres according to their WaterMap-based free energy values (ΔG). Green spheres represent regions where stable water molecules could be placed, while red and orange spheres represent regions where ligand occupancy could contribute to binding affinity by enthalpic energy gain. The hydration site’s index number and color are identical to those in B. (B) Water displacement frequency among replicas with the average free energy values (ΔG). Complete thermodynamic properties of the hydration sites are presented in Table S4. (C) Interaction frequency between the inhibitor and a particular residue during the MD simulation. Charged residues are either orange or purple and the hydrophobic residue is highlighted in green. The red line represents the π-cation interaction, and purple lines represent H-bonds. A dashed line is used as an indication of side-chain interaction and a straight line is used for the backbone. The percentage on the line shows the conservation of the interaction along the simulation time.

MD simulations of the MR22-SIK2 complex (5 replicas x 500 ns = 2.5 μs; Figure S8) were carried out, and representative clusters from these MD simulations were then used as input for WaterMap analyses, revealing that seven of the fifteen hydration sites located in the SIK2 binding pocket display a poor thermodynamic profile (Figure 5A,B; Table S4). These data suggest a potential enthalpic energy gain upon MR22 (**5**) binding through the displacement of energetically unfavorable water molecules. Notably, abundant water displacement with MR22 (**5**) was facilitated by the change in the substitution pattern of the phenyl ring (Figure 4), which resulted in a displacement of two of the most energetically unfavorable water molecules (Hydration site index 1,2; Figure 5A,B; Table S4). Comparing the results of our WaterMap calculations of MR22 (**5**) in SIK2 with the published analysis of the thermodynamic water profile of G-5555 (**2**) binding to MST3 [15] allows an estimate of the contribution of the back-pocket pyridine group to the free energy.

The pyridine nitrogen replaces a favorable water molecule interacting with Lys65, and the C_5_ atom of the pyridine ring displaces an unfavorable water molecule close to the αC-helix [15]. Removal of the terminal pyridine group, as in compounds MR21 (**4**) and MR22 (**5**), results in retention of those unfavorable waters upon binding, which could negatively affect the stabilization of STE kinases. In SIK2, as described above, this can potentially be compensated for by the displacement of other unfavorable waters by the 2,5-difluorophenyl ring, thus contributing to the potent inhibition by MR22 (**5**).

Next, we identified MR22-SIK2 interactions that occurred during more than 20% of the analyzed simulation time (Figure 5C). This analysis demonstrated a nearly constant hydrogen-bond interaction with the hinge (Ala99, 98% of simulation time) and a π-cation interaction with the catalytic lysine (Lys49, 80% of simulation time), complemented by a water bridge between the lysine and the inhibitor carbonyl group (56% of simulation time). Further, the 4-aminobutyl moiety engaged in hydrogen bonding with the catalytic loop (Glu146, 58% of simulation time) and the -1 residue of the αD-helix (Glu103, 36% of simulation time). Additionally, MR22 (**5**) displayed a bidentate interaction with the DFG residue Asp160 via hydrogen bonding with the 4-aminobutyl group (42% of simulation time) and a water bridge to the carbonyl group of the pyrido[2,3-d]pyrimidin-7(8*H*)-one moiety (84% of simulation time), contributing to the robust fixation of the inhibitor within the binding site. Finally, **5** displayed an intermittent salt-bridge interaction with the αD-helix (Asp106, 22% of simulation time). In addition to the main interactions during the MD simulations, we also analyzed the torsional conformations of rotatable bonds in MR22 (**5**) (Figure S9). The 2,5-difluorophenyl group in the back pocket showed a fixed dihedral angle of 90° relative to the central scaffold over the simulation time. To facilitate strong hydrogen-bond interactions in the region of the DFG motif and the catalytic loop, the bond between the C_2_ and C_3_ atoms of the 4-aminobutyl group adopted a dihedral angle of 90° and 180°.

### Cellular off-target evaluation of second-generation SIK inhibitors in NanoBRET assays revealed improved selectivity

The most potent SIK binders with the best balance between on-target potency and selectivity, **4** (MR21), **8,** and **5** (MR22), were further evaluated in the cellular context, using the NanoBRET target engagement assay. For compounds **4** (MR21) and **8**, binding affinity was evaluated on RSK1b, ABL1, BMX, GAK, JNK1, and ephrin receptor kinases (EpHA2, EpHA4, EpHA5, EpHB1, and EpHB3) and compared with the on-target pan-SIK activity (Figure S10). Due to high sequence similarity, additional ephrin receptor kinases (EpHA1, EpHA3, EpHB2, and EpHB4) were also investigated. Both **4** and **8** showed a comparable off-target profile, with a selectivity score of 13/98 and 10/95 kinases, respectively. As described above, MR21 (**4**) potently inhibited all three SIK kinases with IC_50_ values of 20 ± 7 nM (SIK1; lysed), 74 ± 2 nM (SIK2), and 25 ± 3 nM (SIK3). *In vitro* stabilization of RSK1b, ABL1, GAK, EpHB2, and EpHB3 was not confirmed *in cellulo*, with IC_50_ values all above 1 μM. Notably, all three compounds further evaluated strongly stabilized RSK1b in the DSF assay (Figure S10; 6B). Activity of MR21 (**4**) against NEK1, FGFR1, BMPR2, MST3, MST4, and CK1e was not investigated *in cellulo* as no NanoBRET assay is currently available for these kinases. Activity was confirmed against 7 ephrin isoforms with IC_50_ values of 12 ± 2 nM (EpHA1), 210 ± 36 nM (EpHA2), 313 ± 121 nM (EpHA3), 66 ± 38 nM (EpHA4), 928 ± 6 nM (EpHA5), 78 ± 2 nM (EpHB1), and 50 ± 12 nM (EpHB4). Addition of a chlorine atom at position 5 in compound **8** decreased the on-target activity, with IC_50_ values of 277 ± 123 nM, 665 ± 40 nM, and 113 ± 95 nM, against SIK1 (lysed), SIK2, and SIK3, respectively. Compound **8** showed negligible off-target activity with IC_50_ values above 1 µM for RSK1b, ABL1, BMX, as well as EpHA2, EpHA3, EpHA5, EpHB2, and EpHB3. Only EpHA1, EpHA4, EpHB1, and EpHB4 were potently inhibited, with IC_50_ values of 82 ± 3 nM, 346 ± 1 nM, 302 ± 26 nM, and 188 ± 17 nM, respectively. Compounds **4** and **8** can therefore be considered pan-SIK/pan-ephrin kinase inhibitors with improved cellular on-target activity.

### Selectivity profile of the pan-SIK inhibitor MR22 (5)

With MR22 (**5**), the lead structures MRIA9 (**1**) and G-5555 (**2**) were improved by removing the activity against PAK and MST kinases while maintaining potent inhibition of all three SIK isoforms. This was achieved by three modifications (Figure 6A). The selectivity against STE group kinases was improved by removal of the terminal pyridine moiety. MR22 (**5**) showed no stabilization of PAK1. The implementation of a 2,5-difluorophenyl group in the back pocket further improved the selectivity over ephrin receptor kinase off-targets. Additionally, the loss of on-target activity by the modifications in the back pocket were at least partially compensated for by replacing the (2*r*,5*r*)-2-methyl-1,3-dioxan-5-amine group with a 4-aminobutyl residue, thereby enhancing the interaction with the DFG motif and the amino acids of the catalytic-loop (comparison of NanoBRET data of **2** with **7** and **8**). MR22 (**5**) showed improved selectivity in our in-house DSF panel against 100 kinases, with only 8 kinases stabilized by up to 50% compared to a corresponding reference compound (Figure 6B). This included the kinases RSK1b, ABL1, JNK1, GAK, and ephrin kinases EpHA2, EpHA4, EpHB1, and EpHB3. MR22 (**5**) showed similar on-target activity against SIK kinases in the cellular NanoBRET assay as the dual pan-SIK/group I PAK chemical probe MRIA9 (**1**), with IC_50_ values of 72 ± 11 nM (SIK1; lysed), 89 ± 2 nM (SIK2), and 18 ± 2 nM (SIK3) (Figure 6C). The two fluorine substituents, however, significantly reduced off-target activity, with cellular IC_50_ values below 1 µM found only for EpHA1 (IC_50_ = 145 ± 17 nM), EpHB1 (IC_50_ = 632 ± 10 nM), and EpHB4 (IC_50_ = 607 ± 37 nM) (Figure 6D). Unfortunately, crystallographic studies with MR22 (**5**) were not successful due to its low activity towards the surrogates MST3 and EpHA2. In Figure 6E the most prominent cluster of the SIK2-MR22 (5) MD simulation shows key interactions within the binding pocket, including the canonical hinge binding to Ala99 and interactions of the 4-aminobutyl group with Asp160 in the DFG motif and Glu146 located in the catalytic loop.

**Figure 6.**
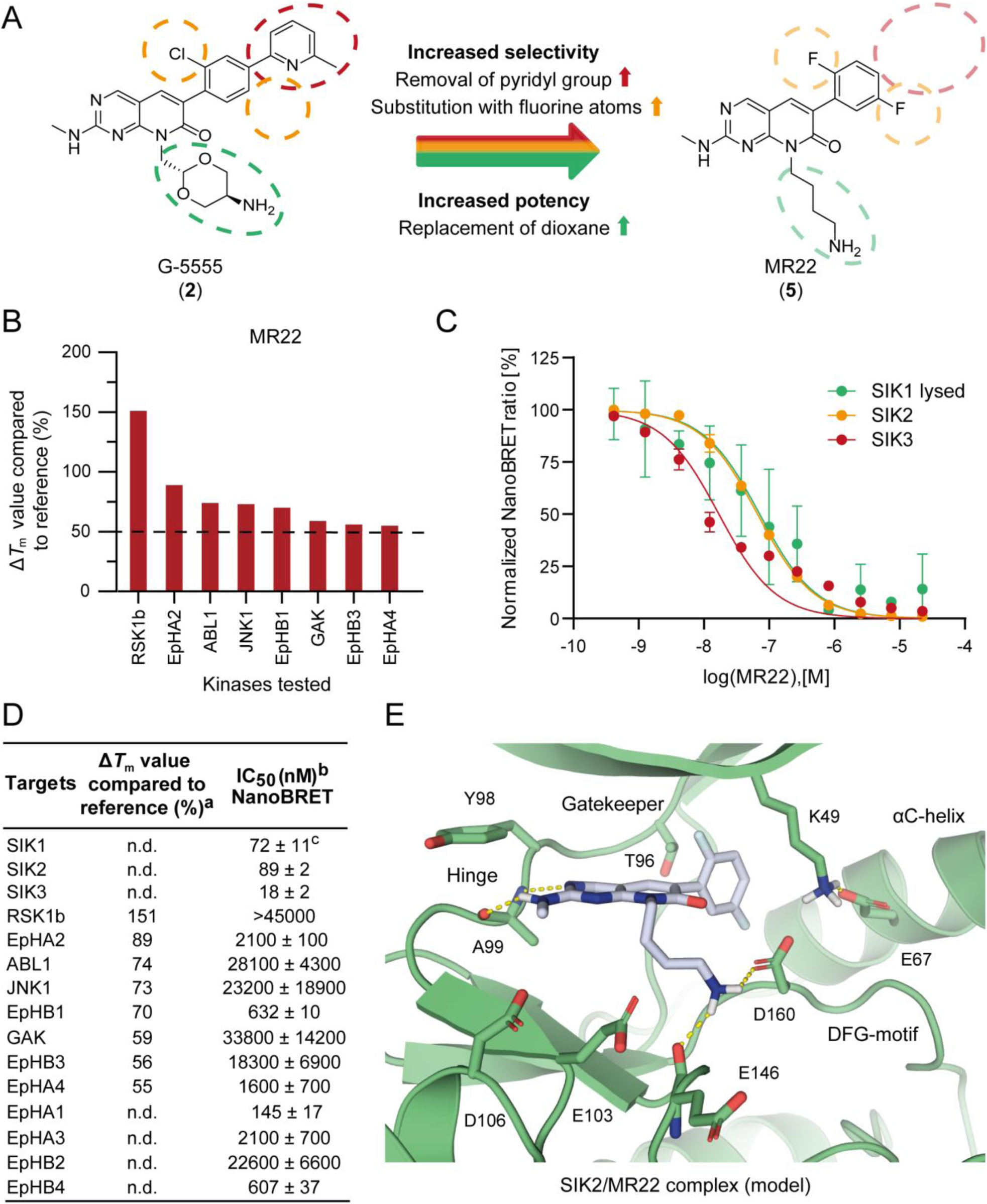
Selectivity profile of pan-SIK inhibitor MR22 (**5**). (A) Design of the potent and selective SIK subfamily inhibitor MR22 (**5**). Removal of the pyridine moiety in the back pocket diminished MST and PAK activity. Fluorine substituents on the phenyl ring modulated ephrin receptor activity. Replacement of the (2*r*,5*r*)-2-methyl-1,3-dioxan-5-amine group of G-5555 (**2**) with a 4-aminobutyl moiety increased binding affinity. (B) Selectivity profile of MR22 (**5**) against 100 kinases, tested in-house in a differential scanning fluorimetry assay. Two measurements were performed and the average was calculated. Obtained *ΔT*m values were compared to those of corresponding reference compounds and are shown in percent (%). Only kinases showing >50% stabilization compared to the reference were considered a potential off-target and are presented. Additional data is shown in Figure S7. MR22 (**5**) was tested at a final concentration of 10 µM. (C) NanoBRET curves obtained against SIK1 (lysed), SIK2, and SIK3 using compound MR22 (**5**). The normalized NanoBRET ratio was plotted against the logarithmic function of the increasing inhibitor concentration. (D) *In cellulo* on- and off-target evaluation using the NanoBRET system. In addition to SIK1-3, IC50 values were obtained against all kinases with a *ΔT*m value >50% as well as against additional ephrin kinases (EpHA1, EpHA3, EpHB2, and EpHB4). (E) Most prominent cluster of the MD simulation of **5** (MR22) bound to SIK2 (model), with the kinase shown in green and the compound in gray. Important amino acids and structural motifs are highlighted, and hydrogen-bond interactions are shown as yellow dashed lines. ^a^Included data can be found in Table S1. ^b^IC50 values with the standard deviation of the mean were determined using NanoBRET assay in a 11-point dose-response curve (duplicate measurements). ^c^SIK1 was measured in lysed mode.

### *In vitro* evaluation of the new pan-SIK inhibitors

We next investigated whether MR22 (**5**) and **4** (MR21) could reproduce phenotypic effects observed upon MRIA9-dependent SIK2 inhibition. First, an *in vitro* kinase assay was performed, monitoring glutathione S-transferase (GST)-tagged SIK2 phosphorylation levels at Ser358 upon administration of increasing concentrations (0.1 nM to 5 µM) of MRIA9 (**1**), MR21 (**4**), and MR22 (**5**) (Figure 7).

**Figure 7.**
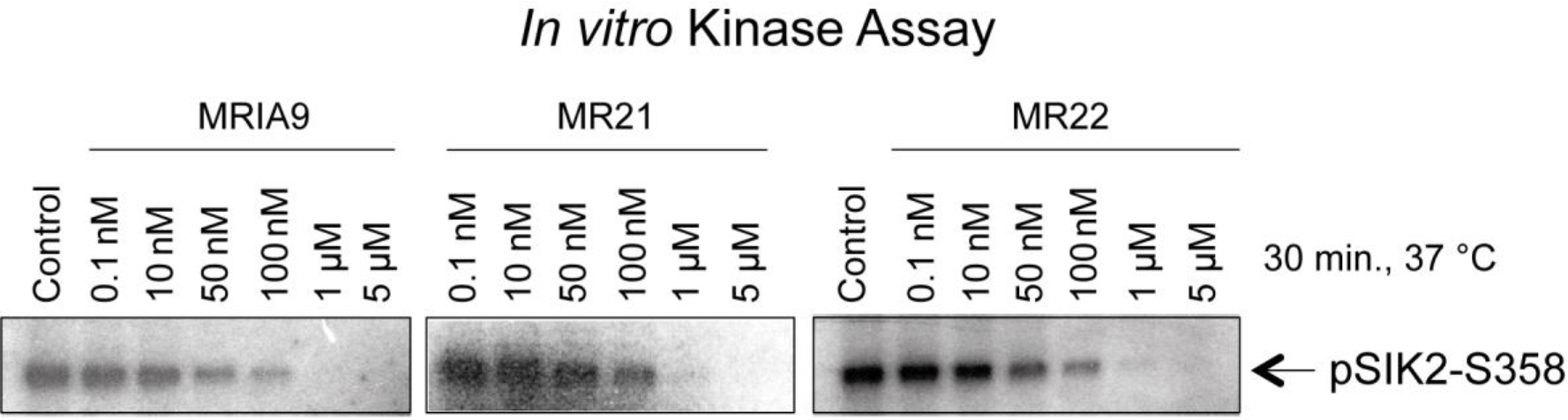
Dose-dependent *in vitro* inhibition of SIK2 by MRIA9 (**1**), MR21 (**4**), and MR22 (**5**). *In vitro* kinase assay using recombinant GST-tagged SIK2 protein incubated with increasing concentrations (0.1 nM to 5 µM) of MRIA9 (**1**), MR21 (**4**), and MR22 (**5**). Data is shown as immunoblot using an antibody monitoring autophosphorylation of SIK2 at Ser358.

Comparing the results for MRIA9 (**1**), MR21 (**4**), and MR22 (**5**), a dose-dependent decrease in SIK2 phosphorylation levels can be observed. Phosphorylation was completely blocked for all three compounds at a concentration of 5 µM. MR21 (4) and MR22 (5) completely mimicked the effects observed for MRIA9-induced inhibition. We have recently demonstrated that SIK2 is responsible for cell-cycle progression, particularly the G2/M transition, through CEP250 phosphorylation at Ser2392, which is associated with centrosome cohesion [16,30]. In addition, SIK2 mediates mitotic spindle assembly, and its expression is cell-cycle dependent [16]. Further, chemical inhibition of SIK2 by MRIA9 (**1**) blocks centrosome disjunction in the late G2 phase [16]. We, therefore, tested whether MR22 (**5**) and MR21 (**4**) can reproduce this effect in G2-arrested SKOV3 ovarian cancer cells (Figure 8). The cells were incubated with MRIA9 (**1**), MR22 (**5**), and MR21 (**4**) at a concentration of 5 μM for 48 hours.

**Figure 8.**
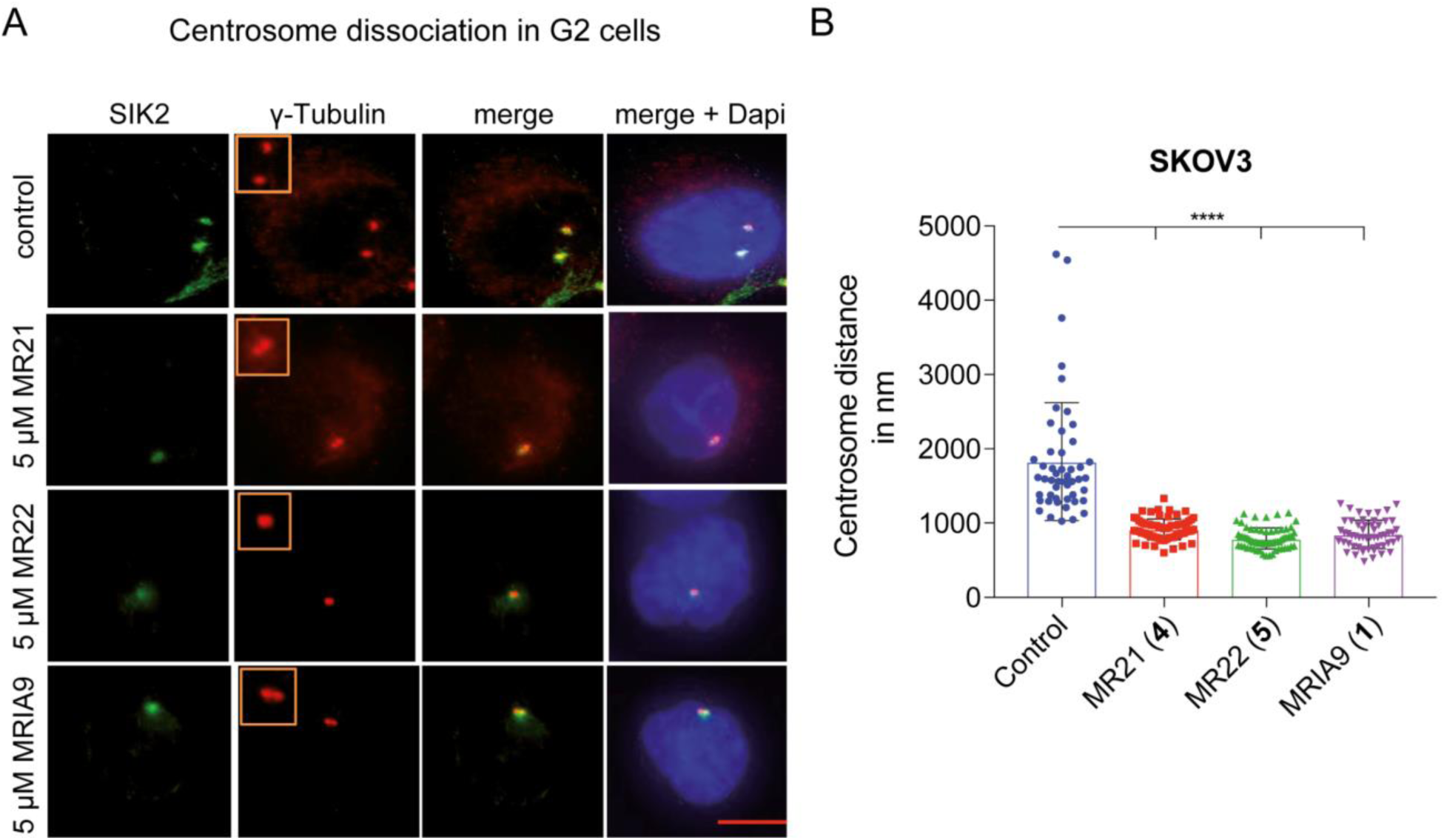
MR22 (**5**)- and MR21 (**4**)-dependent inhibition of SIK2 suppressed centrosome dissociation in G2 phase, as reported for the pan-SIK/PAK1-3 chemical probe MRIA9 (**1**). SKOV3 cells were treated with 5 µM MRIA9 (**1**), MR21 (**4**), and MR22 (**5**) for 48 hours. The cells were released in fresh medium containing the corresponding inhibitor, synchronized using 5 µM RO3306 (CDK1 inhibitor), released in early mitosis, fixed and processed for IF analysis with SIK2, γ-tubulin, and DAPI. DMSO was used in the control cells. (A) Representative images of blocked centrosome disjunction induced by chemical inhibition of SIK2 (MRIA9 (**1**), MR21 (**4**), and MR22 (**5**)), with the magnifications in the insets showing the centrosome disjunction in the different treatment groups. Scale bar = 10 µm. (B) Centrosome distance in the different treatment groups in nm, measured between the separate SIK2 and γ-tubulin signals. Data is shown as a scatter plot, calculated from 100 cells per treatment group and analyzed statistically (**** p < 0.001).

Compared with control cells, a significant reduction in centrosome dissociation was observed with all three compounds. MR21 (**4**) and MR22 (**5**) completely reproduced the effect observed for MRIA9-dependent SIK2 inhibition. The mean centrosome distance was reduced from approximately 1.8 µm in the control group to less than 1 µm after SIK inhibition by MRIA9 (**1**), MR22 (**5**), and MR21 (**4**). Furthermore, this data supported the on-target correlation of the phenotype to the inhibition of SIK kinases, the only common target kinase subfamily of the three inhibitors. Blocking centrosome separation led to subsequent cell-cycle arrest in interphase, preventing the cells from entering a new cell cycle. To investigate this further, we performed a cell-cycle analysis in SKOV3 cells after SIK2 inhibition by **1** (MRIA9), **4** (MR21), and **5** (MR22) (Figure 9).

**Figure 9.**
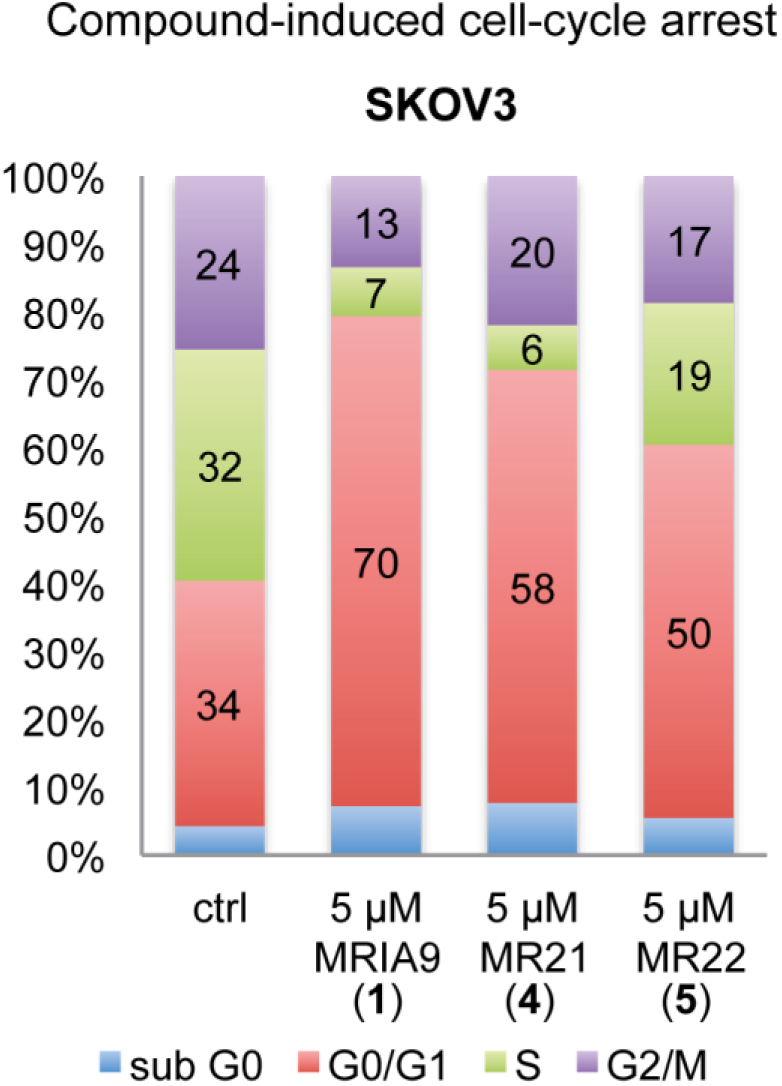
Cell-cycle analysis of SKOV3 cells incubated for 24 hours with 5 µM of MRIA9 (**1**), MR21 (**4**), and MR22 (**5**), and with DMSO as control. Cell-cycle distribution was determined by flow cytometry and is shown in percent (%).

Cell-cycle analysis was performed by incubation of the SKOV3 cells with all three compounds for 24 hours and examined by flow cytometry. We recently described the cell cycle-dependent expression of SIK2, with highest expression levels found in G1 and G2 phases [16]. Control cells were found to be almost evenly distributed between the cell-cycle phases G1, S, and G2, with 34%, 32%, and 24%, respectively. MRIA9 (**1**) as well as MR21 (**4**) and MR22 (**5**) blocked SKOV3 cells in G1 phase and inhibited cell entry into mitosis. This observation is consistent with published data on ovarian [30] and prostate [31] cancers upon SIK2 depletion or chemical inhibition with a promiscuous kinase inhibitor. The most prominent arrest was observed for the chemical probe MRIA9 (**1**), where 70% of the cells were blocked in G1 phase and the proportion of cells in G2 phase decreased to only 13%. In case of MR21 (**4**), the fraction of cells in G1 phase was still significantly increased to 58%, but slightly lower compared with MRIA9 (**1**). MR22 (**5**), in comparison, blocked 50% of the cells in G1 phase. Interestingly, the fraction of cells in S phase increased to 19% compared with 7% and 6% observed with MRIA9 (**1**) and MR21 (**4**), respectively. Overall, **5** (MR22) blocked 69% of the cells in interphase. G1 arrest observed after SIK2 inhibition has been suggested to be induced as an escape mechanism of the cells in response to attenuated mitotic progression [31].

The effect of the newly synthesized compounds on phosphorylation of the SIK2 downstream target protein kinase B (AKT) was examined by Western blot (Figure S11). The phosphoinositide 3-kinase (PI3K)/AKT pathway was activated by treating SKOV3 cells with 2 nM rapamycin for 16 hours, prior to the administration of compounds MRIA9 (**1**), MR21 (**4**), and MR22 (**5**) (5 μM concentration) for 48 hours. While all three compounds reduced the phosphorylation levels of AKT, the most profound results were found for MR22 (**5**), which completely blocked the p-AKT (Ser473) signal (Figure S11). Furthermore, MR22 (**5**) increased the expression levels of PLK1 and cyclin A compared with MRIA9 (**1**), suggesting a potential delay of the cell cycle also in G2 phase.

We further investigated the cell health of HCT116 cells when treated with increasing concentrations of MRIA9 (**1**), MR21 (**4**), and MR22 (**5**) to exclude the occurrence of unwanted side effects. Therefore, we assessed the effect of all three compounds on cell viability and clustered the cells administered with **5** (MR22) for 48 hours into different fractions, based on the morphology of their nuclei (Figure 10A,B) [32]. All compounds were tested at 1 µM and 10 µM over an administration time of 80 hours. For MRIA9 (**1**), no effect on cell viability was observed at 1 µM, which is consistent with the results found for control cells at a final DMSO concentration of 0.1%. MR21 (**4**) and MR22 (**5**) showed a slight decrease in cell count after long exposure (> 60 hours). At the higher dosage of 10 µM, all three compounds showed a significant decrease in cell viability after 24 hours. As the compounds may induce apoptosis by blocking cell-cycle progression, the observed effect on viability can be considered on-target. However, this must be considered in long-term experiments above the recommended usage concentration. For **5**, at a concentration of 1 µM, only a small fraction of cells showed pyknosed nuclei, indicating apoptosis. This number increased to about 30% at a concentration of 10 µM. Nevertheless, the majority of cells in both treatment groups represented a healthy state.

**Figure 10:**
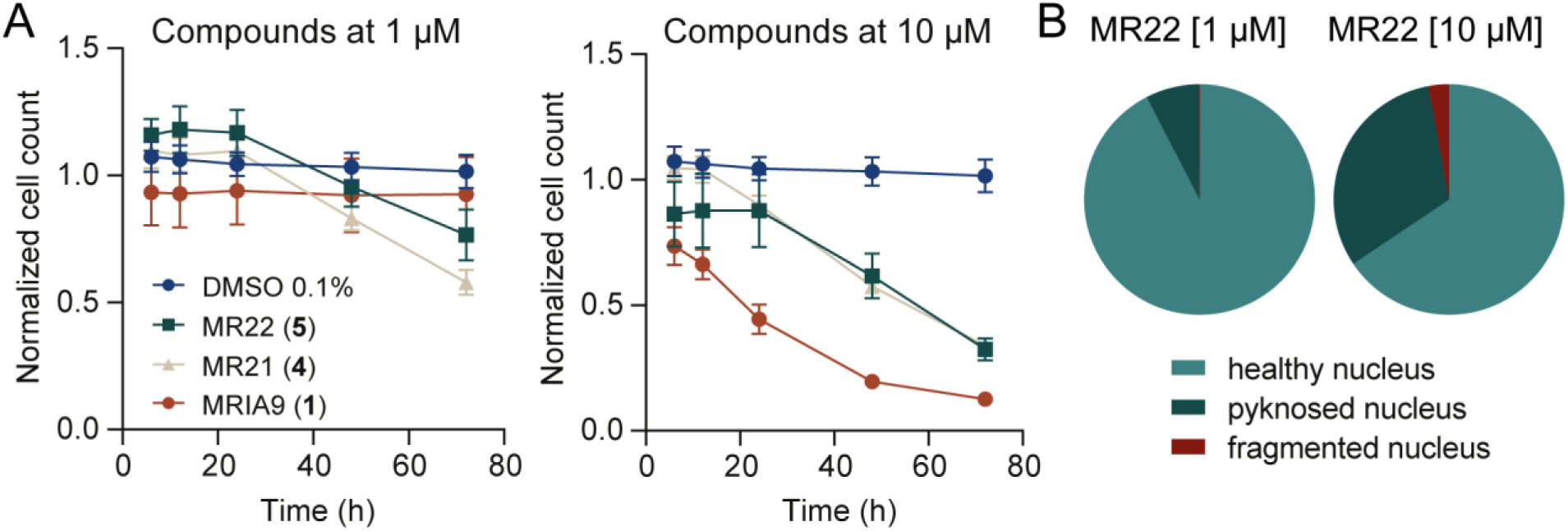
Cell viability assessment using high-content imaging in HCT116 cells. A) Normalized cell count of HCT116 cells after 6 h, 12 h, 24 h, 48 h, and 72 h of treatment with 1 µM and 10 µM of MRIA9 (**1**), MR21 (**4**), and MR22 (**5**) compared with cells exposed to 0.1% DMSO. Error bars show the SEM of two biological replicates. B) Fraction of healthy, fragmented and pyknosed nuclei after 48 h of compound exposure (MR22 (**5**)) at 1 µM and 10 µM, in HCT116 cells. Averages of two biological duplicates are shown.

### Pharmacokinetic profile prediction of MR22 (5)

To further profile compound MR22 (**5**) for potential use *in vivo*, its microsomal stability was determined on activated liver microsomes of Sprague-Dawley rats. Microsomes were incubated with MR22 (**5**) at 37 °C for 1 hour, and metabolization was monitored via high-performance liquid chromatography (HPLC) at four time points (0 min., 15 min., 30 min., and 60 min.). Unfortunately, **5** was rapidly metabolized within 1 hour, which should be considered before using this compound *in vivo*. In comparison, an excellent microsomal stability of 90% is published for MRIA9 (**1**) [15]. In addition, the pharmacokinetic properties of MR22 (**5**) were calculated (QikProp; *Schrödinger*) and compared with published values for G-5555 (**2**) and MRIA9 (**1**) (Figure S12) [15]. MR22 (**5**) showed excellent predicted cell permeability in CACO-2 (196.0 nm/sec.) and HDCK cells (476.8 nm/sec.) compared with G-5555 (**2**)/MRIA9 (**1**). MR22 (**5**) is, therefore, a suitable tool compound for investigating the role of SIK kinases in phenotypic and mechanistic studies both *in vitro* and in a cellular context.

## SUMMARY AND CONCLUSION

SIK2 is an oncogenic marker for ovarian cancer, and its overexpression promotes metastases [13,14]. We have recently developed the dual pan-SIK/PAK1-3 chemical probe MRIA9 (**1**) [15] and demonstrated that it can sensitize ovarian cancer cells to taxane treatment [16]. Excellent kinome-wide selectivity was observed, with remaining *in vitro* off-target activity only against group I PAK kinases [15]. However, chemical inhibition of group I PAKs has been reported to induce *in vivo* lethality at high doses [20].

Here, we aimed to design selective compounds that eliminate the PAK activity of pyrido[2,3-d]pyrimidin-7(8*H*)-one-based inhibitors such as MRIA9 (**1**) and G-5555 (**2**), while maintaining their potent activity against SIK kinases. A set of new inhibitors was designed by focusing on differences in the kinase domains of SIK2, PAK1, and MST3. A comprehensive analysis of published co-crystal structures revealed that vemurafenib (**3**) targets a specific region of the back pocket next to the gatekeeper residue in ZAK. A similar subpocket was identified in all three kinases investigated, although its size and accessibility differed, depending on the gatekeeper residue (a threonine in SIK2 and a larger methionine in PAK1 and MST3). The smallest subpocket was observed in PAK1, which led us to hypothesize that exploiting unique features in the back pocket would abolish activity on STE group kinases. The lead structure G-5555 (**2**) was modified at positions 2 and 5 of the phenyl ring. The SAR analysis was supported by protein crystallography, MD simulations and WaterMap calculations. This structure-guided design approach resulted in compound **5** (MR22), showing the best balance between excellent on-target activity against SIK kinases and overall selectivity. MR22 (**5**) reproduced the effect of MRIA9-dependent SIK2 inhibition in ovarian cancer cells by blocking the centrosome disjunction occurring during the G2 phase and inducing cell-cycle arrest in interphase. Furthermore, our data support the correlation of the observed phenotype induced by MRIA9 (**1**) with on-target SIK activity as MR22 (**5**) had no residual PAK activity *in vitro*. Although the *in vivo* stability of MR22 (**5**) may be an issue, we have provided a valuable tool compound for studying SIK kinase function in cells. Especially when used as a set together with MRIA9 (**1**) and the PAK1-specific chemical probe SGC-NVS-PAK1 [33], MR22 (**5**) can support target deconvolution in future phenotypic screenings on SIK kinases.

## EXPERIMENTAL SECTION

### Chemistry

The synthetic procedure of compounds **4**–**12** will be explained in the following. Further information regarding the analytical data for compounds **4**–**12** and intermediates **21**–**46** can be found in the supporting information. All commercial chemicals were purchased from common suppliers in reagent grade and used without further purification. Reactions were performed under argon atmosphere. Purification by flash chromatography was performed on a puriFlash^®^ XS 420 device with a UV-VIS multi-wave detector (200-400 nm) from *Interchim*. Prepacked normal phase PF-SIHP silica columns were used with a particle size of 30 µm (*Interchim*). All compounds synthesized were characterized by ^1^H/^13^C NMR and mass spectrometry (ESI +/-). In addition, final compounds were identified by high-resolution mass spectra (HRMS) and their purity was evaluated by HPLC. All compounds used for further testing showed >95% purity. ^1^H and ^13^C NMR spectra were measured on a DPX250, an AV400, an AV500 HD Avance III and an AV600 spectrometer from *Bruker Corporation*. Chemical shifts (*δ*) are reported in parts per million (ppm) and coupling constants (*J*) in hertz. DMSO-d_6_ was used as solvent, and the spectra were calibrated to the solvent signal: 2.50 ppm (^1^H NMR) or 39.52 ppm (^13^C NMR). Mass spectra (ESI +/-) were measured on a Surveyor MSQ device from *Thermo Fisher Scientific*. HRMS were obtained on a MALDI LTQ Orbitrap XL (*Thermo Fischer Scientific)*. Preparative purification by high-performance liquid chromatography (HPLC) was carried out on an *Agilent* 1260 Infinity II device using an Eclipse XDB-C18 (*Agilent*, 21.2 x 250mm, 7µm) reversed-phase column. A suitable gradient (flow rate 21 ml/min.) was used, with water (A; 0.1% TFA) and acetonitrile (B; 0.1% TFA) as a mobile phase. All compounds purified by preparative HPLC chromatography were obtained as TFA salts. Determination of the compound purity by HPLC was carried out, on the same *Agilent* 1260 Infinity II device, together with HPLC-MS measurements using a LC/MSD device (G6125B, ESI pos. 100-1000). The compounds were analyzed on a Poroshell 120 EC-C18 (*Agilent*, 3 x 150 mm, 2.7 µm) reversed phase column, with water (A; 0.1% formic acid) and acetonitrile (B; 0.1% formic acid) as a mobile phase, using the following gradient: 0 min. 5% B - 2 min. 5% B - 8 min. 98% B - 10 min. 98% B (flow rate of 0.5 mL/min.). UV-detection was performed at 280 and 310 nm.

### General Procedure I for esterification reaction

The corresponding acetic acid (1.0 eq) was dissolved in ethanol (80 mL) and cooled to 0 °C. Sulfuric acid (1.0 eq) was added, and the reaction solution was allowed to warm to room temperature. The reaction was stirred at 100 °C for 15 hours. Afterwards, the solvent was evaporated under reduced pressure and the remaining residue was dissolved in ethyl acetate (30 mL). The organic layer was washed two times with saturated NaHCO_3_-solution (20 mL) and brine (20 mL). The aqueous layer was extracted with ethyl acetate (20 mL) and the combined organic layers were dried over MgSO_4_. The solvent was evaporated under reduced pressure to obtain the corresponding ethyl acetate derivative.

### General Procedure II for cyclocondensation reaction

The corresponding methyl or ethyl acetate derivative (1.0 eq), 4-amino-2-(methylthio)-pyrimidine-5-carbaldehyde (1.0 eq) and potassium carbonate (3.0 eq) were dissolved in anhydrous DMF (70 mL) and stirred at 120 °C for 16 hours. Water (300 mL) was added to the reaction solution leading to the precipitation of the corresponding pyrido[2,3-d]pyrimidin-7(8*H*)-ol product. After filtration, the aqueous layer was extracted three times with ethyl acetate (30 mL) and the combined organic layers were dried over MgSO_4_. The solvent was evaporated under reduced pressure, and the remaining residue was recrystallized from acetone.

### General procedure III for nucleophilic substitution reaction

The corresponding 2-(methylthio)pyrido[2,3-d]pyrimidin-7(8*H*)-ol derivative (1 eq) and primary alkyl bromide (1.0 or 1.5 eq), together with cesium carbonate (3.0 eq) were dissolved in anhydrous DMF (20 mL) and stirred at 110 °C for 18 hours. The solvent was evaporated under reduced pressure, and the remaining residue was purified by flash chromatography on silica gel using DCM/MeOH as an eluent. In case of 2-((2*r*,5*r*)-2-(bromomethyl)-1,3-dioxan-5-yl)isoindoline-1,3-dione or 2-(4-bromobutyl)isoindoline-1,3-dione as a primary alkyl bromide, a mixture of the corresponding isoindoline-1,3-dione and 2-carbamoylbenzoic acid derivative was obtained and used in the next step without further separation.

### General procedure IV for oxidation of the methyl sulfide to a sulfoxide/sulfone

**The** corresponding methyl sulfide derivative (1 eq.) was dissolved in anhydrous DCM (10 mL). *M*-chloroperoxybenzoic acid (*m*-CPBA) (≤ 77%, 2.1 eq) was added in one portion, and the reaction solution was stirred at room temperature for up to three hours. The reaction was monitored by HPLC analysis and stopped after complete transformation by adding saturated aq. NaHCO_3_-solution (10 mL). The aqueous layer was extracted three times with a mixture of DCM/MeOH (4:1; 15 mL) and DCM (15 mL). The combined organic layers were dried over Na_2_SO_4_, and the solvent was evaporated under reduced pressure. The remaining residue, consisting a mixture of the corresponding sulfoxide and sulfone product, was used in the next step without further purification.

### General procedure V for nucleophilic aromatic substitution reaction

The remaining residue of the corresponding product obtained from general procedure IV (1.0 eq) was dissolved in ethanol (10 mL), and a solution of methylamine in THF (2M, 4.0 eq) and *N*,*N*-diisopropylethylamine (5.0 eq) were added. The reaction solution was stirred at 85 °C for 17 hours. Afterwards the solvent was evaporated under reduced pressure. The remaining residue was purified either by flash chromatography on silica gel using DCM/MeOH as an eluent or by preparative HPLC chromatography on silica gel (H_2_O/ACN; 0.1% TFA).

### General procedure VI for deprotection of BOC-protected amines

The corresponding BOC-protected amine was dissolved in anhydrous DCM (5 mL), and TFA (20 vol%) was added. The reaction solution was stirred at room temperature for 30 min. Afterwards the reaction solution was evaporated under reduced pressure. If necessary, the remaining residue was purified either by flash chromatography on silica gel using DCM/MeOH as an eluent or by preparative HPLC chromatography on silica gel (H_2_O/ACN; 0.1% TFA). Compounds obtained from this reaction were isolated as TFA salt.

*Ethyl 2-(2,5-difluorophenyl)acetate (**21**).*Compound **21** was prepared according to general procedure I, using **14** (1.00 g, 5.81 mmol) as starting material. **21** was obtained as a white solid with low melting temperature, in a yield of 99% (1.15 g). ^1^H NMR (500 MHz, DMSO-d_6_, 300K): *δ* = 7.27 - 7.21 (m, 2H), 7.19 - 7.12 (m, 1H), 4.10 (q, *J* = 7.1 Hz, 2H), 3.73 (d, *J* = 1.4 Hz, 2H), 1.18 (t, *J* = 7.1 Hz, 3H) ppm. ^13^C NMR (126 MHz, DMSO-d_6_, 300K): *δ* = 169.7, 158.3 (dd, *J* = 118.3, 2.2 Hz), 156.4 (dd, *J* = 119.4, 2.2 Hz), 123.6 (dd, *J* = 18.9, 8.8 Hz), 118.3 (dd, *J* = 24.6, 4.7 Hz), 116.4 (dd, *J* = 24.8, 9.0 Hz), 115.4 (dd, *J* = 24.0, 8.7 Hz), 60.6, 33.8, 14.0 ppm. MS (ESI+): m/z = 342.1 [2M_fr._]^+^; calculated: 342.1.

*Ethyl 2-(2-chloro-5-bromophenyl)acetate (**22**).* Compound **22** was prepared according to general procedure I, using **15** (4.00 g, 16.0 mmol) as starting material. **22** was obtained as a pale yellow solid with low melting temperature, in a yield of 92% (4.10 g). ^1^H NMR (500 MHz, DMSO-d_6_, 300K): *δ* = 7.68 (d, *J* = 2.4 Hz, 1H), 7.52 (dd, *J* = 8.5, 2.4 Hz, 1H), 7.42 (d, *J* = 8.5 Hz, 1H), 4.10 (q, *J* = 7.1 Hz, 2H), 3.82 (s, 2H), 1.18 (t, *J* = 7.1 Hz, 3H) ppm. ^13^C NMR (126 MHz, DMSO-d_6_, 300K): *δ* = 169.7, 135.3, 134.7, 133.2, 131.8, 131.0, 119.8, 60.7, 38.1, 14.1 ppm. MS (ESI+): m/z = 301.0 [M + Na]^+^; calculated: 300.9.

*Ethyl 2-(2,5-dichlorophenyl)acetate (**23**).* Compound **23** was prepared according to general procedure I, using **16** (5.00 g, 24.4 mmol) as starting material. **23** was obtained as a white solid with low melting temperature, in a yield of 99% (5.66 g). ^1^H NMR (500 MHz, DMSO-d_6_, 300K): *δ* = 7.55 (d, *J* = 2.6 Hz, 1H), 7.49 (d, *J* = 8.6 Hz, 1H), 7.40 (dd, *J* = 8.6, 2.6 Hz, 1H), 4.10 (q, *J* = 7.1 Hz, 2H), 3.82 (s, 2H), 1.18 (t, *J* = 7.1 Hz, 3H) ppm. ^13^C NMR (126 MHz, DMSO-d_6_, 300K): *δ* = 169.5, 134.9, 132.5, 131.8, 131.5, 130.6, 128.8, 60.6, 38.1, 14.0 ppm. MS (ESI+): m/z = 255.1 [M + Na]^+^; calculated: 255.0.

*Ethyl 2-(2-chloro-5-(trifluoromethyl)phenyl)acetate (**24**).* Compound **24** was prepared according to general procedure I, using **17** (1.00 g, 4.19 mmol) as starting material. **24** was obtained as a white solid with low melting temperature, in a yield of 99% (1.11 g). ^1^H NMR (500 MHz, DMSO-d_6_, 300K): *δ* = 7.91 - 7.84 (m, 1H), 7.75 - 7.66 (m, 2H), 4.11 (q, *J* = 7.1 Hz, 2H), 3.94 (s, 2H), 1.19 (t, *J* = 7.1 Hz, 3H) ppm. ^13^C NMR (126 MHz, DMSO-d_6_, 300K): *δ* = 169.5, 138.2, 134.4, 130.1, 128.9 (q, *J* = 3.7 Hz), 127.8 (q, *J* = 32.3 Hz), 125.8 (q, *J* = 3.7 Hz), 123.8 (q, *J* = 272.3 Hz), 60.6, 38.2, 14.0 ppm. MS (ESI+): m/z = 289.1 [M + Na]^+^; calculated: 289.0.

*Ethyl 2-(2-bromo-5-chlorophenyl)acetate (**25**).* Compound **25** was prepared according to general procedure I, using **18** (1.00 g, 4.0 mmol) as starting material. **25** was obtained as a white solid with low melting temperature, in a yield of 94% (1.04 g). ^1^H NMR (500 MHz, DMSO-d_6_, 300K): *δ* = 7.64 (d, *J* = 8.5 Hz, 1H), 7.55 (d, *J* = 2.6 Hz, 1H), 7.31 (dd, *J* = 8.5, 2.6 Hz, 1H), 4.11 (q, *J* = 7.1 Hz, 2H), 3.82 (s, 2H), 1.19 (t, *J* = 7.1 Hz, 3H) ppm. ^13^C NMR (126 MHz, DMSO-d_6_, 300K): *δ* = 169.5, 136.8, 133.9, 132.2, 131.9, 129.0, 123.0, 60.6, 40.6, 14.0 ppm. MS (ESI+): m/z = 261.0 [M_fr._]^+^; calculated: 260.9.

*Ethyl 2-(2,5-dibromophenyl)acetate (**26**).* Compound **26** was prepared according to general procedure I, using **19** (1.00 g, 3.4 mmol) as starting material. **26** was obtained as a white solid with low melting temperature, in a yield of 99% (1.09 g). ^1^H NMR (500 MHz, DMSO-d_6_, 300K): *δ* = 7.67 (d, *J* = 2.5 Hz, 1H), 7.57 (d, *J* = 8.5 Hz, 1H), 7.43 (dd, *J* = 8.5, 2.5 Hz, 1H), 4.10 (q, *J* = 7.1 Hz, 2H), 3.82 (s, 2H), 1.19 (t, *J* = 7.1 Hz, 3H) ppm. ^13^C NMR (126 MHz, DMSO-d_6_, 300K): *δ* = 169.5, 137.1, 134.7, 134.1, 131.9, 123.7, 120.5, 60.6, 40.5, 14.0 ppm. MS (ESI+): m/z = 345.0 [M + Na]^+^; calculated: 344.9.

*6-(2-Chlorophenyl)-2-(methylthio)pyrido[2,3-d]pyrimidin-7-ol (**27**).* Compound **27** was prepared according to general procedure II, using **20** (1.00 g, 5.42 mmol) as starting material. **27** was obtained as a yellow solid in a combined yield of 65% (1.07 g). ^1^H NMR (500 MHz, DMSO-d_6_, 300K): *δ* = 12.63 (s, 1H), 8.89 (s, 1H), 7.95 (s, 1H), 7.55 (d, *J* = 6.9 Hz, 1H), 7.46 - 7.41 (m, 3H), 2.59 (s, 3H) ppm. ^13^C NMR (126 MHz, DMSO-d_6_, 300K): *δ* = 172.1, 161.4, 156.9, 154.2, 136.4, 134.8, 132.8, 131.8, 131.7, 130.0, 129.2, 127.1, 108.8, 13.7 ppm. MS (ESI+): m/z = 304.0 [M + H]^+^; calculated: 304.0.

*6-(2,5-Difluorophenyl)-2-(methylthio)pyrido[2,3-d]pyrimidin-7-ol (**28**).* Compound **28** was prepared according to general procedure II, using **21** (1.16 g, 5.79 mmol) as starting material. **28** was obtained as a pale-yellow solid in a combined yield of 96% (1.70 g). ^1^H NMR (500 MHz, DMSO-d_6_, 300K): *δ* = 12.70 (s, 1H), 8.91 (s, 1H), 8.09 (s, 1H), 7.42 - 7.31 (m, 3H), 2.58 (s, 3H) ppm. ^13^C NMR (126 MHz, DMSO-d_6_, 300K): *δ* = 172.4, 161.3, 157.1, 156.7 (td, *J* = 237.9, 3.9 Hz, 2C), 154.1, 137.4, 126.7, 124.7 (dd, *J* = 17.5, 9.1 Hz), 118.0 (dd, *J* = 24.7, 3.8 Hz), 117.1 (dd, *J* = 25.1, 9.2 Hz), 116.7 (dd, *J* = 23.9, 8.9 Hz), 108.8, 13.7 ppm. MS (ESI+): m/z = 306.0 [M + Na]^+^; calculated: 306.1.

*6-(5-Bromo-2-chlorophenyl)-2-(methylthio)pyrido[2,3-d]pyrimidin-7-ol (**29**).* Compound **29** was prepared according to general procedure II, using **22** (2.00 g, 7.21 mmol) as starting material. **29** was obtained as a pale-yellow solid in a combined yield of 80% (2.22 g). ^1^H NMR (500 MHz, DMSO-d_6_, 300K): *δ* = 12.70 (s, 1H), 8.89 (s, 1H), 8.01 (s, 1H), 7.67 - 7.62 (m, 2H), 7.52 (d, *J* = 8.4 Hz, 1H), 2.58 (s, 3H) ppm. ^13^C NMR (126 MHz, DMSO-d_6_, 300K): *δ* = 172.4, 161.2, 157.1, 154.3, 137.0, 136.9, 134.1, 132.7, 132.3, 131.2, 130.4, 119.7, 108.7, 13.7 ppm. MS (ESI+): m/z = 405.9 [M + Na]^+^; calculated: 405.9.

*6-(2,5-Dichlorophenyl)-2-(methylthio)pyrido[2,3-d]pyrimidin-7-ol (**30**).* Compound **30** was prepared according to general procedure II, using **23** (2.00 g, 8.58 mmol) as starting material. **30** was obtained as a yellow solid in a combined yield of 93% (2.69 g). ^1^H NMR (500 MHz, DMSO-d_6_, 300K): *δ* = 12.67 (s, 1H), 8.87 (s, 1H), 7.99 (s, 1H), 7.59 (d, *J* = 8.5 Hz, 1H), 7.55 - 7.50 (m, 2H), 2.58 (s, 3H) ppm. ^13^C NMR (126 MHz, DMSO-d_6_, 300K): *δ* = 172.2, 161.5, 156.9, 154.5, 136.9, 136.7, 131.7, 131.4, 131.3, 130.9, 130.4, 129.7, 108.7, 13.7 ppm. MS (ESI+): m/z = 338.0 [M + H]^+^; calculated: 338.0.

*6-(2-Chloro-5-(trifluoromethyl)phenyl)-2-(methylthio)-pyrido*[2,3-d]*pyrimidin-7-ol (**31**).* Compound **31** was prepared according to general procedure II, using **24** (1.20 g, 4.5 mmol) as starting material. **31** was obtained as a pale-yellow solid in a combined yield of 47% (0.79 g). ^1^H NMR (500 MHz, DMSO-d_6_, 300K): *δ* = 12.73 (s, 1H), 8.90 (s, 1H), 8.07 (s, 1H), 7.85 - 7.81 (m, 3H), 2.59 (s, 3H) ppm. ^13^C NMR (126 MHz, DMSO-d_6_, 300K): *δ* = 172.5, 161.2, 157.1, 154.3, 137.5, 137.3, 135.9, 130.4, 130.3, 128.6 (q, *J* = 3.5 Hz), 127.8 (q, *J* = 32.5 Hz), 126.7 (q, *J* = 3.5 Hz), 123.7 (q, *J* = 272.5 Hz), 108.7, 13.7 ppm. MS (ESI+): m/z = 394.1 [M + Na]^+^; calculated: 394.0.

*6-(2-Bromo-5-chlorophenyl)-2-(methylthio)pyrido*[2,3-d]*pyrimidin-7-ol (**32**).* Compound **32** was prepared according to general procedure II, using **25** (1.00 g, 3.6 mmol) as starting material. **32** was obtained as a pale-yellow solid in a combined yield of 84% (1.16 g). ^1^H NMR (500 MHz, DMSO-d_6_, 300K): *δ* = 12.70 (s, 1H), 8.89 (s, 1H), 7.97 (s, 1H), 7.74 (d, *J* = 8.6 Hz, 1H), 7.52 (d, *J* = 2.6 Hz, 1H), 7.44 (dd, *J* = 8.6, 2.6 Hz, 1H), 2.59 (s, 3H) ppm. ^13^C NMR (126 MHz, DMSO-d_6_, 300K): *δ* = 172.3, 161.2, 157.0, 154.4, 138.7, 136.6, 134.0, 132.2, 132.2, 131.3, 129.9, 121.8, 108.7, 13.7 ppm. MS (ESI+): m/z = 381.9 [M + Na]^+^; calculated: 381.9.

*6-(2,5-Dibromophenyl)-2-(methylthio)pyrido[2,3-d]pyrimidin-7-ol (**33**).* Compound **33** was prepared according to general procedure II, using **26** (1.20 g, 3.73 mmol) as starting material. **33** was obtained as a yellow solid in a combined yield of 67% (1.06 g). ^1^H NMR (500 MHz, DMSO-d_6_, 300K): *δ* = 12.69 (s, 1H), 8.88 (s, 1H), 7.97 (s, 1H), 7.67 (d, *J* = 8.5 Hz, 1H), 7.63 (d, *J* = 2.5 Hz, 1H), 7.55 (dd, *J* = 8.5, 2.5 Hz, 1H), 2.58 (s, 3H) ppm. ^13^C NMR (126 MHz, DMSO-d_6_, 300K): *δ* = 172.4, 161.1, 157.0, 154.2, 139.0, 136.7, 134.2, 134.1, 132.8, 132.1, 122.5, 120.4, 108.7, 13.7 ppm. MS (ESI+): m/z = 427.8 [M + H]^+^; calculated: 427.9.

*tert-Butyl (4-(6-(2-chlorophenyl)-2-(methylthio)-7-oxopyrido[2,3-d]pyrimidin-8(7H)-yl)butyl)carbamate (**34**).* Compound **34** was prepared according to general procedure III, using **27** (300 mg, 0.99 mmol) and *tert*-butyl (4-bromobutyl)carbamate (249 mg, 0.99 mmol) as starting materials. **34** was obtained as a white solid in a yield of 88% (414 mg). ^1^H NMR (500 MHz, DMSO-d_6_, 300K): *δ* = 8.91 (s, 1H), 7.99 (s, 1H), 7.56 - 7.54 (m, 1H), 7.45 - 7.42 (m, 3H), 6.78 (t, *J* = 5.4 Hz, 1H), 4.35 (t, *J* = 7.2 Hz, 2H), 2.95 - 2.91 (m, 2H), 2.62 (s, 3H), 1.69 - 1.65 (m, 2H), 1.45 - 1.41 (m, 2H), 1.35 (s, 9H) ppm. ^13^C NMR (126 MHz, DMSO-d_6_, 300K): *δ* = 172.1, 160.4, 157.4, 155.6, 153.4, 135.4, 135.1, 132.9, 131.7, 130.7, 130.0, 129.2, 127.1, 109.1, 77.3, 40.6, 40.1, 28.2 (3C), 27.1, 24.9, 13.9 ppm. MS (ESI+): m/z = 375.1 [M-BOC + H]^+^; calculated: 375.1.

*tert-Butyl (4-(6-(2,5-difluorophenyl)-2-(methylthio)-7-oxopyrido[2,3-d]pyrimidin-8(7H)-yl)butyl)carbamate (**35**).* Compound **35** was prepared according to general procedure III, using **28** (200 mg, 0.66 mmol) and *tert*-butyl (4-bromobutyl)carbamate (165 mg, 0.66 mmol) as starting materials. **35** was obtained as a white solid in a yield of 86% (267 mg). ^1^H NMR (500 MHz, DMSO-d_6_, 300K): *δ* = 8.93 (s, 1H), 8.13 (s, 1H), 7.40 - 7.32 (m, 3H), 6.78 (t, *J* = 5.3 Hz, 1H), 4.35 (t, *J* = 7.2 Hz, 2H), 2.95 - 2.92 (m, 2H), 2.62 (s, 3H), 1.69 - 1.64 (m, 2H), 1.46 - 1.42 (m, 2H), 1.35 (s, 9H) ppm. ^13^C NMR (126 MHz, DMSO-d_6_, 300K): *δ* = 172.4, 160.2, 157.6, 156.8 (td, *J* = 237.5, 4.1 Hz, 2C), 155.55, 153.35, 136.4, 125.6, 125.0 (dd, *J* = 14.1, 12.3 Hz), 118.0 (dd, *J* = 22.3, 6.3 Hz), 117.2 - 116.5 (m, 2C), 109.1, 77.3, 40.8, 40.1, 28.2 (3C), 27.1, 24.8, 13.9 ppm. MS (ESI+): m/z = 377.0 [M-BOC + H]^+^; calculated: 377.1.

*tert-Butyl (4-(6-(5-bromo-2-chlorophenyl)-2-(methylthio)-7-oxopyrido[2,3-d]pyrimidin-8(7H)-yl)butyl)carbamate (**36**).* Compound **36** was prepared according to general procedure III, using **29** (200 mg, 0.52 mmol) and *tert*-butyl (4-bromobutyl)carbamate (132 mg, 0.52 mmol) as starting materials. **36** was obtained as a yellow solid in a yield of 86% (249 mg). ^1^H NMR (500 MHz, DMSO-d_6_, 300K): *δ* = 8.91 (s, 1H), 8.05 (s, 1H), 7.67 - 7.64 (m, 2H), 7.52 (d, *J* = 8.8 Hz, 1H), 6.77 (t, *J* = 5.3 Hz, 1H), 4.34 (t, *J* = 7.1 Hz, 2H), 2.95 - 2.90 (m, 2H), 2.62 (s, 3H), 1.70 - 1.63 (m, 2H), 1.45 - 1.40 (m, 2H), 1.34 (s, 9H) ppm. ^13^C NMR (126 MHz, DMSO-d_6_, 300K): *δ* = 172.4, 160.2, 157.6, 155.6, 153.5, 137.2, 136.1, 134.1, 132.7, 132.4, 131.2, 129.3, 119.7, 109.0, 77.4, 40.7, 40.1, 28.3 (3C), 27.1, 24.9, 14.0 ppm. MS (ESI+): m/z = 453.0 [M-BOC + H]^+^; calculated: 453.0.

*tert-Butyl (5-(6-(2-chlorophenyl)-2-(methylamino)-7-oxopyrido[2,3-d]pyrimidin-8(7H)-yl)butyl)carbamate (**37**).* Compound **37** was prepared according to general procedures IV and V, using **34** (390 mg, 0.82 mmol) as starting material. The crude product was purified by flash chromatography on silica gel using DCM/MeOH as an eluent. **37** was obtained as a white solid in a yield of 53% (198 mg) over two steps. ^1^H NMR (500 MHz, DMSO-d_6_, 300K): *δ* = 8.70 - 8.60 (m, 1H), 7.89 - 7.81 (m, 1H), 7.75 (s, 1H), 7.52 - 7.50 (m, 1H), 7.41 - 7.37 (m, 3H), 6.79 (s, 1H), 4.32 - 4.19 (m, 2H), 2.96 - 2.90 (m, 5H), 1.70 - 1.58 (m, 2H), 1.46 - 1.40 (m, 2H), 1.35 (s, 9H) ppm. ^13^C NMR (126 MHz, DMSO-d_6_, 300K): *δ* = 162.0, 160.9, 159.3, 155.6, 155.0, 136.2, 136.0, 133.2, 132.0, 130.7, 129.5, 129.1, 127.0, 103.9, 77.3, 40.1 (2C), 28.3 (3C), 27.9, 27.3, 24.9 ppm. MS (ESI+): m/z = 358.1 [M-BOC + H]^+^; calculated: 358.1.

*tert-Butyl (4-(6-(2,5-difluorophenyl)-2-(methylamino)-7-oxopyrido[2,3-d]pyrimidin-8(7H)-yl)butyl)carbamate (**38**).* Compound **38** was prepared according to general procedures IV and V, using **35** (100 mg, 0.21 mmol) as starting material. The crude product was purified by flash chromatography on silica gel using DCM/MeOH as an eluent. **38** was obtained as a white solid in a yield of 46% (44 mg) over two steps. ^1^H NMR (500 MHz, DMSO-d_6_, 300K): *δ* = 8.75 - 8.60 (m, 1H), 7.92 - 7.88 (m, 2H), 7.37 - 7.22 (m, 3H), 6.85 - 6.73 (m, 1H), 4.34 - 4.19 (m, 2H), 2.96 - 2.89 (m, 5H), 1.70 - 1.56 (m, 2H), 1.49 - 1.40 (m, 2H), 1.35 (s, 9H) ppm. ^13^C NMR (126 MHz, DMSO-d_6_, 300K): *δ* = 162.0, 160.7, 159.6, 157.8 (dd, *J* = 224.1, 3.6 Hz), 155.8 (dd, *J* = 224.3, 3.8 Hz), 155.6, 155.0, 137.1, 125.9 (t, *J* = 13.1 Hz), 119.5, 118.0 (dd, *J* = 22.4, 6.0 Hz), 116.8 (dd, *J* = 19.4, 15.3 Hz), 115.9 (dd, *J* = 21.9, 10.6 Hz), 104.0, 77.3, 40.2, 40.1, 28.2 (3C), 27.8, 27.2, 24.8 ppm. MS (ESI+): m/z = 360.1 [M-BOC + H]^+^; calculated: 360.2.

*tert-Butyl (4-(6-(5-bromo-2-chlorophenyl)-2-(methylamino)-7-oxopyrido[2,3-d]pyrimidin-8(7H)-yl)butyl)carbamate (**39**).* Compound **39** was prepared according to general procedures IV and V, using **36** (226 mg, 0.41 mmol) as starting material. The crude product was purified by flash chromatography on silica gel using DCM/MeOH as an eluent. **39** was obtained as a pale yellow solid in a yield of 45% (98 mg) over two steps. ^1^H NMR (500 MHz, DMSO-d_6_, 300K): *δ* = 8.72 -8.59 (m, 1H), 7.89 (d, *J* = 4.8 Hz, 1H), 7.82 (s, 1H), 7.63 - 7.58 (m, 2H), 7.49 (d, *J* = 8.6 Hz, 1H), 6.78 (s, 1H), 4.32 - 4.18 (m, 2H), 2.96 - 2.90 (m, 5H), 1.70 - 1.55 (m, 2H), 1.46 - 1.39 (m, 2H), 1.35 (s, 9H) ppm. ^13^C NMR (126 MHz, DMSO-d_6_, 300K): *δ* = 162.0, 160.7, 159.5, 155.5, 155.0, 138.0, 136.7, 134.3, 132.6, 132.1, 131.0, 123.3, 119.5, 103.8, 77.3, 40.1, 39.9, 28.2 (3C), 27.8, 27.2, 24.8 ppm. MS (ESI+): m/z = 436.1 [M-BOC + H]^+^; calculated: 436.1.

*8-(4-Aminobutyl)-6-(2-chlorophenyl)-2-(methylamino)pyrido[2,3-d]pyrimidin-7(8H)-one* Compound **4** was prepared according to general procedure VI, using **37** (138 mg, 0.30 mmol) as starting material. The crude product was purified by flash chromatography on silica gel, using DCM/MeOH as an eluent, to obtain **4** as a yellow solid in a yield of 41% (73 mg). ^1^H NMR (500 MHz, DMSO-d_6_, 300K): *δ* = 8.73 - 8.60 (m, 1H), 7.89 (d, *J* = 4.3 Hz, 1H), 7.80 - 7.69 (m, 4H), 7.55 - 7.49 (m, 1H), 7.43 - 7.36 (m, 3H), 4.40 - 4.24 (m, 2H), 2.92 (d, *J* = 4.7 Hz, 3H), 2.84 (s, 2H), 1.84 - 1.67 (m, 2H), 1.67 - 1.52 (m, 2H) ppm. ^13^C NMR (126 MHz, DMSO-d_6_, 300K): *δ* = 162.0, 161.0, 159.4, 155.0, 136.3, 135.8, 133.2, 131.9, 129.5, 129.1, 126.9, 124.8, 103.9, 40.1, 38.7, 27.9, 24.7, 24.4 ppm. HRMS (FTMS +p MALDI): m/z = 358.1430 [M + H]^+^; calculated for C_18_H_21_ClN_5_O: 358.1429. HPLC: t_R_ = 7.02 min.; purity ≥ 95% (UV: 280/310 nm). LCMS (ESI+): m/z = 358.2 [M + H]^+^; calculated: 358.1.

*8-(4-Aminobutyl)-6-(2,5-difluorophenyl)-2-(methylamino)pyrido[2,3-d]pyrimidin-7(8H)-one* Compound **5** was prepared according to general procedure VI, using **38** (20 mg, 0.04 mmol) as starting material. The crude product was purified by flash chromatography on silica gel, using DCM/MeOH as an eluent, to obtain **5** as a yellow solid in a yield of 43% (11 mg). ^1^H NMR (500 MHz, DMSO-d_6_, 300K): *δ* = 8.77 - 8.62 (m, 1H), 7.99 - 7.90 (m, 2H), 7.71 (s, 3H), 7.37 - 7.25 (m, 3H), 4.38 - 4.22 (m, 2H), 2.92 (d, *J* = 4.7 Hz, 3H), 2.87 - 2.81 (m, 2H), 1.83 - 1.70 (m, 2H), 1.64 - 1.56 (m, 2H) ppm. ^13^C NMR (126 MHz, DMSO-d_6_, 300K): *δ* = 162.5, 161.3, 160.1, 158.2 (dd, *J* = 225.2, 3.7 Hz), 156.3 (dd, *J* = 228.2, 3.6 Hz), 155.5, 137.8, 126.3 (dd, *J* = 16.9, 9.6 Hz), 119.9, 118.5 (dd, *J* = 20.9, 7.5 Hz), 117.3 (dd, *J* = 24.5, 10.2 Hz), 116.5 (dd, *J* = 21.9, 10.8 Hz), 104.4, 40.6, 39.2, 28.4, 25.2, 24.9 ppm. HRMS (FTMS +p MALDI): m/z = 360.1627 [M + H]^+^; calculated for C_18_H_20_F_2_N_5_O: 360.1630. HPLC: t_R_ = 7.05 min.; purity ≥ 95% (UV: 280/310 nm). LCMS (ESI+): m/z = 360.2 [M + H]^+^; calculated: 360.2.

*8-(4-Aminobutyl)-6-(5-bromo-2-chlorophenyl)-2-(methylamino)pyrido[2,3-d]pyrimidin-7(8H)-one (**6**).* Compound **6** was prepared according to general procedure VI, using **39** (84 mg, 0.16 mmol) as starting material. The crude product was purified by flash chromatography on silica gel, using DCM/MeOH as an eluent, to obtain **6** as a yellow solid in a yield of 75% (78 mg). ^1^H NMR (500 MHz, DMSO-d_6_, 300K): *δ* = 8.73 - 8.61 (m, 1H), 7.93 (d, *J* = 4.6 Hz, 1H), 7.88 - 7.78 (m, 4H), 7.63 - 7.58 (m, 2H), 7.49 (dd, *J* = 7.7, 1.2 Hz, 1H), 4.38 - 4.21 (m, 2H), 2.91 (d, *J* = 4.8 Hz, 3H), 2.84 (s, 2H), 1.82 - 1.67 (m, 2H), 1.63 - 1.54 (m, 2H) ppm. ^13^C NMR (126 MHz, DMSO-d_6_, 300K): *δ* = 162.0, 160.8, 159.6, 155.1, 137.9, 137.0, 134.3, 132.7, 132.2, 131.1, 123.3, 119.5, 103.9, 40.1, 38.7, 27.9, 24.7, 24.4 ppm. HRMS (FTMS +p MALDI): m/z = 436.0535 [M + H]^+^; calculated for C_18_H_20_BrClN_5_O: 436.0534. HPLC: t_R_ = 7.37 min.; purity ≥ 95% (UV: 280/310 nm). LCMS (ESI+): m/z = 436.1 [M + H]^+^; calculated: 436.1.

*8-(((2r,5r)-5-Amino-1,3-dioxan-2-yl)methyl)-6-(5-bromo-2-chlorophenyl)-2-(methylamino)pyrido[2,3-d]pyrimidin-7(8H)-one (**7**).* Compound **7** was prepared according to general procedures III-V. **29** (300 mg, 0.78 mmol) was used as starting material. The crude product was purified by HPLC chromatography on silica gel (H_2_O/ACN; 0.1% TFA), to obtain **7** as a yellow solid in a yield of 16% (89 mg) over three steps. ^1^H NMR (500 MHz, DMSO-d_6_, 300K): *δ* = 8.72 - 8.60 (m, 1H), 8.15 (s, 3H), 8.01 - 7.94 (m, 1H), 7.86 (s, 1H), 7.65 - 7.58 (m, 2H), 7.52 - 7.47 (m, 1H), 5.15 - 4.97 (m, 1H), 4.57 - 4.43 (m, 2H), 4.20 (dd, *J* = 10.7, 3.9 Hz, 2H), 3.57 (s, 2H), 3.40 - 3.34 (m, 1H), 2.91 (d, *J* = 4.8 Hz, 3H) ppm. ^13^C NMR (126 MHz, DMSO-d_6_, 300K): *δ* = 161.9, 160.8, 159.6, 155.3, 137.7, 137.4, 134.3, 132.6, 132.2, 131.1, 123.1, 119.6, 103.8, 97.4, 66.3 (2C), 42.3, 42.1, 27.9 ppm. HRMS (FTMS +p MALDI): m/z = 480.0434 [M + H]^+^; calculated for C_19_H_20_BrClN_5_O_3_: 480.0433. HPLC: t_R_ = 7.42 min.; purity ≥ 95% (UV: 280/310 nm). LCMS (ESI+): m/z = 482.1 [M + H]^+^; calculated: 482.0.

*8-(4-Aminobutyl)-6-(2,5-dichlorophenyl)-2-(methylamino)pyrido[2,3-d]pyrimidin-7(8H)-one (**8**).* Compound **8** was prepared according to general procedures III-V. **30** (300 mg, 0.89 mmol) was used as starting material. The crude product was purified by HPLC chromatography on silica gel (H_2_O/ACN; 0.1% TFA), to obtain **8** as a yellow solid in a yield of 11% (60 mg) over three steps. ^1^H NMR (500 MHz, DMSO-d_6_, 300K): *δ* = 8.78 - 8.60 (m, 1H), 8.01 - 7.93 (m, 1H), 7.85 (s, 1H), 7.72 (s, 3H), 7.57 (dd, *J* = 7.8, 1.1 Hz, 1H), 7.51 - 7.46 (m, 2H), 4.40 - 4.20 (m, 2H), 2.92 (d, *J* = 3.9 Hz, 3H), 2.87 - 2.81 (m, 2H), 1.79 - 1.71 (m, 2H), 1.62 - 1.54 (m, 2H) ppm. ^13^C NMR (126 MHz, DMSO-d_6_, 300K): *δ* = 161.9, 160.8, 159.4, 155.1, 137.6, 136.9, 132.0, 131.5, 131.3, 130.8, 129.2, 123.4, 103.9, 40.1, 38.7, 27.9, 24.7, 24.4 ppm. HRMS (FTMS +p MALDI): m/z = 392.1034 [M + H]^+^; calculated for C_18_H_20_Cl_2_N_5_O: 392.1039. HPLC: t_R_ = 7.21 min.; purity ≥ 95% (UV: 280/310 nm). LCMS (ESI+): m/z = 392.1 [M + H]^+^; calculated: 392.1.

*8-(4-Aminobutyl)-6-(2-chloro-5-(trifluoromethyl)phenyl)-2-(methylamino)pyrido[2,3-d]pyrimidin-7(8H)-one (**9**).* Compound **9** was prepared according to general procedures III-V. **31** (200 mg, 0.54 mmol) was used as starting material. The crude product was purified by HPLC chromatography on silica gel (H_2_O/ACN; 0.1% TFA), to obtain **9** as a yellow solid in a yield of 16% (57 mg) over three steps. ^1^H NMR (500 MHz, DMSO-d_6_, 300K): *δ* = 8.76 - 8.62 (m, 1H), 8.00 - 7.94 (m, 1H), 7.92 (s, 1H), 7.80 - 7.78 (m, 2H), 7.78 - 7.77 (m, 1H), 7.66 (s, 3H), 4.37 - 4.27 (m, 2H), 2.93 (d, *J* = 4.5 Hz, 3H), 2.87 - 2.81 (m, 2H), 1.81 - 1.72 (m, 2H), 1.63 - 1.55 (m, 2H) ppm. ^13^C NMR (126 MHz, DMSO-d_6_, 300K): *δ* = 162.0, 160.8, 159.6, 155.1, 137.8 - 137.7 (m), 137.2, 136.9, 130.4, 128.9 - 128.6 (m), 127.7 (q, *J* = 32.6 Hz), 126.4 - 126.0 (m), 123.8 (q, *J* = 272.6 Hz), 123.1, 103.9, 40.1, 38.7, 27.9, 24.7, 24.4 ppm. HRMS (FTMS +p MALDI): m/z = 426.1304 [M + H]^+^; calculated for C_19_H_20_ClF_3_N_5_O: 426.1303. HPLC: t_R_ = 7.41 min.; purity ≥ 95% (UV: 280/310 nm). LCMS (ESI+): m/z = 426.2 [M + H]^+^; calculated: 426.1.

*8-(4-Aminobutyl)-6-(2-bromo-5-chlorophenyl)-2-(methylamino)pyrido[2,3-d]pyrimidin-7(8H)-one (**10**).* Compound **10** was prepared according to general procedures III-V. **32** (250 mg, 0.65 mmol) was used as starting material. The crude product was purified by HPLC chromatography on silica gel (H_2_O/ACN; 0.1% TFA), to obtain **10** as a yellow solid in a yield of 13% (55 mg) over three steps. ^1^H NMR (500 MHz, DMSO-d_6_, 300K): *δ* = 8.74 - 8.62 (m, 1H), 7.96 (s, 1H), 7.83 - 7.80 (m, 1H), 7.72 (d, *J* = 8.5 Hz, 4H), 7.46 (d, *J* = 2.6 Hz, 1H), 7.40 (dd, *J* = 8.6, 2.6 Hz, 1H), 4.41 - 4.23 (m, 2H), 2.92 (d, *J* = 2.9 Hz, 3H), 2.89 - 2.80 (m, 2H), 1.83 - 1.68 (m, 2H), 1.62 - 1.56 (m, 2H) ppm. ^13^C NMR (126 MHz, DMSO-d_6_, 300K): *δ* = 161.9, 160.7, 159.4, 155.1, 139.7, 136.7, 133.9, 132.1, 131.5, 129.4, 125.3, 122.4, 103.9, 40.1, 38.7, 27.9, 24.7, 24.4 ppm. HRMS (FTMS +p MALDI): m/z = 438.0510 [M + H]^+^; calculated for C_18_H_20_BrClN_5_O: 438.0514. HPLC: t_R_ = 7.36 min.; purity ≥ 95% (UV: 280/310 nm). LCMS (ESI+): m/z = 438.1 [M + H]^+^; calculated: 438.1.

*8-(4-Aminobutyl)-6-(2,5-dibromophenyl)-2-(methylamino)pyrido[2,3-d]pyrimidin-7(8H)- one (**11**).* Compound **11** was prepared according to general procedures III-V. **33** (250 mg, 0.59 mmol) was used as starting material. The crude product was purified by HPLC chromatography on silica gel (H_2_O/ACN; 0.1% TFA), to obtain **11** as a yellow solid in a yield of 11% (47 mg) over three steps. ^1^H NMR (500 MHz, DMSO-d_6_, 300K): *δ* = 8.74 - 8.61 (m, 1H), 7.97 - 7.92 (m, 1H), 7.82 (s, 1H), 7.71 (s, 3H), 7.65 (d, *J* = 8.5 Hz, 1H), 7.58 (d, *J* = 2.4 Hz, 1H), 7.52 (dd, *J* = 8.5, 2.5 Hz, 1H), 4.36 - 4.29 (m, 2H), 2.92 (d, *J* = 4.5 Hz, 3H), 2.89 - 2.79 (m, 2H), 1.81 - 1.70 (m, 2H), 1.64 - 1.52 (m, 2H) ppm. ^13^C NMR (126 MHz, DMSO-d_6_, 300K): *δ* = 162.0, 160.7, 159.5, 155.1, 140.0, 136.7, 134.2, 134.2, 132.3, 125.2, 123.0, 120.3, 103.8, 40.4, 38.7, 27.9, 24.7, 24.4 ppm. HRMS (FTMS +p MALDI): m/z = 482.0001 [M + H]^+^; calculated for C_18_H_20_Br_2_N_5_O: 482.0009. HPLC: t_R_ = 7.44 min.; purity ≥ 95% (UV: 280/310 nm). LCMS (ESI+): m/z = 482.0 [M + H]^+^; calculated: 482.0.

*Ethyl 2-(4-bromo-2,5-difluorophenyl)acetate (**41**).* Compound **41** was prepared according to general procedure I, using **40** (1.00 g, 3.98 mmol) as starting material. **41** was obtained as a white solid with low melting temperature, in a yield of 99% (1.10 g). ^1^H NMR (500 MHz, DMSO- d_6_, 300K): *δ* = 7.70 (dd, *J* = 8.9, 5.8 Hz, 1H), 7.45 (dd, *J* = 9.0, 6.4 Hz, 1H), 4.10 (q, *J* = 7.1 Hz, 2H), 3.73 (d, *J* = 0.9 Hz, 2H), 1.18 (t, *J* = 7.1 Hz, 3H) ppm. ^13^C NMR (126 MHz, DMSO-d_6_, 300K): *δ* = 169.4, 156.6 (dd, *J* = 245.7, 2.6 Hz), 154.5 (dd, *J* = 240.9, 2.7 Hz), 123.5 (dd, *J* = 18.7, 7.8 Hz), 119.8 (d, *J* = 27.5 Hz), 119.1 (dd, *J* = 25.2, 5.1 Hz), 106.8 (dd, *J* = 23.5, 10.5 Hz), 60.7, 33.6, 14.0 ppm. MS (ESI+): m/z = 279.0 [M + H]^+^; calculated: 279.0.

*Ethyl 2-(2,5-difluoro-4-(4,4,5,5-tetramethyl-1,3,2-dioxaborolan-2-yl)phenyl)acetate (**42**).* **41** (500 mg, 1.79 mmol), bis(pinacolato)diboron (591 mg, 2.33 mmol), potassium acetate (527 mg, 5.37 mmol) and [1,1’-bis(diphenylphosphino)ferrocene]dichloropalladium(II) (92 mg, 0.13 mmol) were dissolved in anhydrous dioxane (10 mL) and stirred at 115 °C for 18 hours. The solvent was evaporated under reduced pressure, and the remaining residue was purified by flash chromatography on silica gel using cyclohexane/ethyl acetate as an eluent. **42** was obtained as a colorless oil in a yield of 39% (230 mg). ^1^H NMR (500 MHz, DMSO-d_6_, 300K): *δ* = 7.29 (dd, *J* = 9.3, 4.6 Hz, 1H), 7.22 (dd, *J* = 9.0, 5.6 Hz, 1H), 4.10 (q, *J* = 7.1 Hz, 2H), 3.76 (s, 2H), 1.30 (s, 12H), 1.18 (t, *J* = 7.1 Hz, 3H) ppm. ^13^C NMR (126 MHz, DMSO-d_6_, 300K): *δ* = 169.5, 161.8 (d, *J* = 246.0 Hz), 156.5 (d, *J* = 244.4 Hz), 127.6 (dd, *J* = 19.0, 9.5 Hz), 121.2 (dd, *J* = 22.9, 9.3 Hz), 118.8 (dd, *J* = 27.3, 4.2 Hz), 84.1 (2C), 60.6, 34.0, 24.5 (4C), 14.0 ppm. MS (ESI+): m/z = 327.1 [M + H]^+^; calculated: 327.2.

*Ethyl 2-(2,5-difluoro-4-(6-methylpyridin-2-yl)phenyl)acetate (**43**).* **42** (230 mg, 0.71 mmol), potassium acetate (138 mg, 1.41 mmol), [1,1′-bis(diphenylphosphino)- ferrocene]dichloropalladium(II) (36 mg, 0.05 mmol) and 2-bromo-6-methylpyridine (121 mg, 0.71 mmol) were dissolved in a mixture of dioxane/water (2:1; 12 mL) and stirred at 107 °C for 18 hours. The solvent was evaporated under reduced pressure and the remaining residue was purified by flash chromatography on silica gel using cyclohexane/ethyl acetate as an eluent. **43** was obtained as a pale green oil in a yield of 44% (91 mg). ^1^H NMR (500 MHz, DMSO-d_6_, 300K): *δ* = 7.81 (t, *J* = 7.8 Hz, 1H), 7.72 (dd, *J* = 10.4, 6.5 Hz, 1H), 7.63 (dd, *J* = 7.5, 1.5 Hz, 1H), 7.39 (dd, *J* = 11.2, 6.2 Hz, 1H), 7.30 (d, *J* = 7.6 Hz, 1H), 4.12 (q, *J* = 7.1 Hz, 2H), 3.80 (s, 2H), 2.55 (s, 3H), 1.20 (t, *J* = 7.1 Hz, 3H) ppm. ^13^C NMR (126 MHz, DMSO-d_6_, 300K): *δ* = 169.7, 158.2, 157.1 (dd, *J* = 177.5, 4.6 Hz), 155.2 (dd, *J* = 182.8, 4.7 Hz), 150.2, 137.3, 127.2 (dd, *J* = 13.9, 7.9 Hz), 124.1 (dd, *J* = 19.0, 9.5 Hz), 122.8, 121.1 (d, *J* = 9.7 Hz), 119.4 (dd, *J* = 26.2, 5.2 Hz), 116.1 (dd, *J* = 25.3, 4.4 Hz), 60.7, 33.7, 24.2, 14.0 ppm. MS (ESI+): m/z = 292.1 [M + H]^+^; calculated: 292.1.

*6-(2,5-Difluoro-4-(6-methylpyridin-2-yl)phenyl)-2-(methylthio)pyrido[2,3-d]pyrimidin-7-ol (**44**).* Compound **44** was prepared according to general procedure II, using **43** (85 mg, 0.29 mmol) as starting material. **44** was obtained as a pale yellow solid in a yield of 96% (111 mg). ^1^H NMR (500 MHz, DMSO-d_6_, 300K): *δ* = 12.73 (s, 1H), 8.93 (s, 1H), 8.16 (s, 1H), 7.89 - 7.80 (m, 2H), 7.69 (d, *J* = 7.5 Hz, 1H), 7.57 (dd, *J* = 11.3, 5.8 Hz, 1H), 7.32 (d, *J* = 7.6 Hz, 1H), 2.59 (s, 3H), 2.57 (s, 3H) ppm. ^13^C NMR (126 MHz, DMSO-d_6_, 300K): *δ* = 172.5, 161.2, 158.3, 157.2, 156.5 (dd, *J* = 55.9, 1.5 Hz), 154.6 (dd, *J* = 55.7, 1.4 Hz), 154.1, 150.0, 137.5 (d, *J* = 2.1 Hz), 137.4, 128.2 (dd, *J* = 14.2, 7.9 Hz), 126.1, 124.8 (dd, *J* = 17.3, 9.2 Hz), 123.0, 121.3 (d, *J* = 10.1 Hz), 119.2 (dd, *J* = 27.2, 3.2 Hz), 116.6 (dd, *J* = 26.7, 3.7 Hz), 108.8, 24.2, 13.7 ppm. MS (ESI+): m/z = 397.1 [M + H]^+^; calculated: 397.1.

tert-Butyl (4-(6-(2,5-difluoro-4-(6-methylpyridin-2-yl)phenyl)-2-(methylthio)-7- oxopyrido[2,3-d]pyrimidin-8(7H)-yl)butyl)carbamate (**45**). Compound **45** was prepared according to general procedure III, using **44** (105 mg, 0.26 mmol) and tert-butyl (4-bromobutyl)carbamate (67 mg, 0.26 mmol) as starting materials. The crude product was purified by flash chromatography on silica gel using DCM/MeOH as an eluent. **45** was obtained as a white solid in a yield of 71% (107 mg). ^1^H NMR (500 MHz, DMSO-d_6_, 300K): δ = 8.97 (s, 1H), 8.21 (s, 1H), 7.87 - 7.83 (m, 2H), 7.70 (d, J = 7.0 Hz, 1H), 7.57 (dd, J = 11.4, 5.9 Hz, 1H), 7.33 (d, J = 7.7 Hz, 1H), 6.79 (t, J = 5.6 Hz, 1H), 4.37 (t, J = 7.2 Hz, 2H), 2.97 - 2.93 (m, J = 12.8, 6.6 Hz, 2H), 2.63 (s, 3H), 2.57 (s, 3H), 1.72 - 1.66 (m, 2H), 1.48 - 1.43 (m, 2H), 1.35 (s, 9H) ppm. ^13^C NMR (126 MHz, DMSO-d_6_, 300K): δ = 172.5, 160.2, 158.4, 157.7, 156.5 (dd, J = 52.9, 1.6 Hz), 155.6, 154.6 (dd, J = 54.3, 1.8 Hz), 153.4, 150.0, 137.4, 136.6 (d, J = 2.1 Hz), 128.3 (dd, J = 14.1, 8.1 Hz), 125.2 (dd, J = 19.0, 7.8 Hz), 125.1, 123.0, 121.3 (d, J = 10.1 Hz), 119.2 (dd, J = 27.0, 3.5 Hz), 116.6 (dd, J = 26.6, 3.6 Hz), 109.1, 77.4, 40.8, 40.1, 28.2 (3C), 27.1, 24.8, 24.3, 13.9 ppm. MS (ESI+): m/z = 468.2 [M-BOC + H]^+^; calculated: 468.2.

tert-Butyl (4-(6-(2,5-difluoro-4-(6-methylpyridin-2-yl)phenyl)-2-(methylamino)-7- oxopyrido[2,3-d]pyrimidin-8(7H)-yl)butyl)carbamate (**46**). Compound **46** was prepared according to general procedures IV and V, using **45** (95 mg, 0.17 mmol) as starting material. The crude product was purified by flash chromatography on silica gel using DCM/MeOH as an eluent. **46** was obtained as a pale yellow solid in a yield of 37% (34 mg) over two steps. ^1^H NMR (500 MHz, DMSO-d_6_, 300K): δ = 8.75 - 8.63 (m, 1H), 7.97 (s, 1H), 7.95 - 7.90 (m, 1H), 7.84 - 7.78 (m, 2H), 7.68 (d, J = 7.3 Hz, 1H), 7.54 - 7.50 (m, 1H), 7.31 (d, J = 7.7 Hz, 1H), 6.82 - 6.75 (m, 1H), 4.39 - 4.21 (m, 2H), 2.98 - 2.93 (m, 2H), 2.91 (d, J = 4.8 Hz, 3H), 2.57 (s, 3H), 1.71 - 1.58 (m, 2H), 1.49 - 1.41 (m, 2H), 1.35 (s, 9H) ppm. ^13^C NMR (126 MHz, DMSO-d_6_, 300K): δ = 162.0, 160.7, 159.7, 158.3, 156.6 (dd, J = 62.2, 1.5 Hz), 155.6, 155.0, 154.6 (dd, J = 64.0, 1.5 Hz), 150.1, 137.4, 137.3 (d, J = 1.6 Hz), 128.8, 127.4 (dd, J = 14.0, 7.6 Hz), 126.1 (dd, J = 17.1, 9.8 Hz), 122.8, 121.2 (d, J = 10.3 Hz), 119.1 (dd, J = 27.0, 3.5 Hz), 116.4 (dd, J = 26.9, 3.6 Hz), 104.0, 77.3, 40.2, 40.1, 28.2 (3C), 27.8, 27.2, 24.8, 24.3 ppm. MS (ESI+): m/z = 551.3 [M + H]^+^; calculated: 551.3.

*8-(4-Aminobutyl)-6-(2,5-difluoro-4-(6-methylpyridin-2-yl)phenyl)-2- (methylamino)pyrido[2,3-d]pyrimidin-7(8H)-one (**12**).* Compound **12** was prepared according to general procedure VI, using **46** (21 mg, 0.04 mmol) as starting material. **12** was obtained as a yellow solid in a yield of 81% (21 mg). ^1^H NMR (500 MHz, DMSO-d_6_, 300K): *δ* = 8.79 - 8.67 (m, 1H), 8.10 (s, 1H), 8.05 - 8.00 (m, 2H), 7.85 - 7.79 (m, 2H), 7.73 (s, 3H), 7.57 (dd, *J* = 11.4, 6.0 Hz, 1H), 7.49 (d, *J* = 7.9 Hz, 1H), 4.42 - 4.24 (m, 2H), 2.94 (s, 3H), 2.88 - 2.82 (m, 2H), 2.63 (s, 3H), 1.81 - 1.72 (m, 2H), 1.65 - 1.57 (m, 2H) ppm. ^13^C NMR (126 MHz, DMSO-d_6_, 300K): *δ* = 161.4, 160.9, 159.0, 157.6, 156.5 (dd, *J* = 71.1, 1.6 Hz), 155.3, 154.6 (dd, *J* = 73.1, 1.7 Hz), 148.7, 139.8, 137.6, 126.9 (dd, *J* = 17.2, 9.5 Hz), 125.5 (dd, *J* = 14.1, 8.5 Hz), 124.2, 122.5 (d, *J* = 8.6 Hz), 119.3 (dd, *J* = 26.3, 3.6 Hz), 119.2, 117.1 (dd, *J* = 27.1, 3.1 Hz), 104.2, 40.1, 38.9, 28.0, 24.9, 24.5, 23.0 ppm. HRMS (FTMS +p MALDI): m/z = 451.2048 [M + H]^+^; calculated for C_24_H_25_F_2_N_6_O: 451.2052. HPLC: t_R_ = 6.83 min.; purity ≥ 95% (UV: 280/310 nm). LCMS (ESI+): m/z = 451.2 [M + H]^+^; calculated: 451.2.

### Statistical analyses and figures

Structural images were generated using PyMol 2.4.3, (*Schrödinger LCC*) and graphs were plotted using GraphPad Prism version 8.4, (*GraphPad Software*, San Diego, California, USA, www.graphpad.com).

### NanoBRET assay

The assay was performed as described previously [34,35]. In brief: full-length kinases were obtained as plasmids cloned in frame with a terminal NanoLuc fusion (*Promega*) as specified in Table S3. Plasmids were transfected into HEK293T cells using FuGENE HD (*Promega*, E2312), and proteins were allowed to express for 20 h. Serially diluted inhibitor and the corresponding NanoBRET Kinase Tracer (*Promega*) at a concentration determined previously as the Tracer IC_50_ (Table S3) were pipetted into white 384-well plates (*Greiner* 781207) using an Echo acoustic dispenser (*Labcyte*). The corresponding protein-transfected cells were added and reseeded at a density of 2 x 10^5^ cells/mL after trypsinization and resuspending in Opti-MEM without phenol red (*Life Technologies*). The system was allowed to equilibrate for 2 hours at 37°C/5% CO_2_ prior to BRET measurements. To measure BRET, NanoBRET NanoGlo Substrate + Extracellular NanoLuc Inhibitor (*Promega*, N2540) was added as per the manufacturer’s protocol, and filtered luminescence was measured on a PHERAstar plate reader (*BMG Labtech*) equipped with a luminescence filter pair (450 nm BP filter (donor) and 610 nm LP filter (acceptor)). Competitive displacement data were then graphed using GraphPad Prism 9 software using a normalized 3-parameter curve fit with the following equation: Y=100/(1+10^(X-LogIC50)).

### Differential scanning fluorimetry (DSF) assay

The assay was performed as previously described [36,37]. Briefly, recombinant protein kinase domains at a concentration of 2 μM were mixed with 10 μM compound in a buffer containing 20 mM HEPES, pH 7.5, and 500 mM NaCl. SYPRO Orange (5000×, Invitrogen) was added as a fluorescence probe (1 µl per mL). Subsequently, temperature-dependent protein unfolding profiles were measured using the QuantStudio 5 realtime PCR machine (Thermo Fisher). Excitation and emission filters were set to 465 nm and 590 nm, respectively. The temperature was raised at a heating of 3 °C per minute. Data points were analyzed with the internal software (Thermal Shift Software Version 1.4, Thermo Fisher) using the Boltzmann equation to determine the inflection point of the transition curve. Differences in melting temperature are given as *ΔT*_m_ values in °C.

### Protein expression and purification

MST3 (STK24) and EPHA2 were expressed and purified as previously described by Tesch *et al.* [15] and Gande *et al*. [38], respectively. MST3 (R4-D301) was subcloned in pNIC-CH vector (non-cleavable C-terminal His_6_-tag), while EPHA2 (D596-G900) was in pET28a vector (N-terminal His_6_-tag, followed by a TEV cleavage site). The expression plasmids were transformed in *Escherichia coli* Rosetta BL21(D3)-R3-pRARE2 and BL21(D3)-R3-λPP competent cells respectively. Initially, cells were cultured in Terrific Broth (TB) media at 37°C to an optical density (OD) 2.8 prior to induction with 0.5 mM IPTG at 18 °C overnight. Cells were harvested and resuspended in a buffer containing (i) for MST3, 50 mM HEPES pH 7.5, 500 mM NaCl, 30 mM imidazole, 5% glycerol, and 0.5 mM TCEP; (ii) for EPHA2, 50 mM Tris, pH 7.5, 500 mM NaCl, 5 mM imidazole, 5% glycerol, and 1 mM TCEP and lysed by sonication. The recombinant proteins were initially purified by Ni^2+^-affinity chromatography. The Ni-NTA fraction containing MST3 was concentrated using a 30 kDa cutoff ultrafiltration device and further purified by size exclusion chromatography (SEC; HiLoad 16/600 Superdex 200) pre-equilibrated with SEC buffer (25 mM HEPES pH 7.5, 200 mM NaCl, 0.5 mM TCEP, 5% glycerol). In the case of EPHA2, the histidine tag was removed by TEV protease treatment and the cleaved protein was separated by reverse Ni^2+^-affinity purification. The protein was further purified by size exclusion chromatography using a HiLoad 16/600 Superdex 200 with a buffer containing 30 mM Tris, pH 7.5, 150 mM NaCl, 0.5 mM TCEP and 5% glycerol. Quality control was performed by SDS-gel electrophoresis and ESI-MS (MST3: expected 34,507.7 Da, observed 34,506.9 Da, EPHA2: expected 34,418.8 Da, observed 34,419.2 Da).

### Crystallization and structure determination

Co-crystallization trials were performed using the sitting-drop vapor-diffusion method at 293 K with a mosquito crystallization robot (*TTP Labtech*, Royston UK). MST3 (11 mg/mL in 25 mM HEPES pH 7.5, 200 mM NaCl, 0.5 mM TCEP, 5% glycerol) and EPHA2 (12 mg/mL in 30 mM Tris pH 7.5, 150 mM NaCl, 0.5 mM TCEP) proteins were incubated with inhibitor at a final concentration of 1 mM prior to setting up crystallization trials. Crystals grew under the following conditions: (i) MST3 with **12**, 20% PEG 3350, 0.1M bis-tris pH 6.5, 1:1 volume ratio; and (ii) EPHA2 with **4** or **9**, 20% PEG 3350, 10% ethylene glycol, 0.2 M potassium thiocyanate, 2:1 volume ratio. Crystals were cryo-protected with mother liquor supplemented with 20% ethylene glycol and flash-frozen in liquid nitrogen. X-ray diffraction data sets were collected at 100 K at beamline X06SA of the Swiss Light Source, Villigen, Switzerland. Diffraction data were integrated with the program XDS [39] and scaled with AIMLESS [40], which is part of the CCP4 package [41]. The ephrin receptor kinase structures were solved by molecular replacement using PHASER [42] (with PDB entry 5NKA as a starting model), and the MST3 structures were solved by difference Fourier analysis using PHENIX [43] with PDB entry 7B32 as a starting model. Structure refinement was performed using iterative cycles of manual model building in COOT [44] and refinement in PHENIX. Dictionary files for the compounds were generated using the Grade Web Server (http://grade.globalphasing.org). X-ray data collection and refinement statistics are shown in Table S5.

### Homology modeling and molecular docking

System preparation and docking calculations were performed using the *Schrödinger* Drug Discovery suite for molecular modeling (version 2021.2). Protein−ligand complexes were prepared with the Protein Preparation Wizard [45] to fix protonation states of amino acids, add hydrogens, and fix missing side-chain atoms. The template structures for homology modeling were identified using BLAST [46], and microtubule affinity regulating kinase 2 (MARK2; PDB code 5EAK) and MST4 (PDB code 7B36) were selected as the main templates. The model was built as a multi-template composite/chimera type with Prime [47], followed by hydrogen bond assignment and energy minimization with Protein Preparation Wizard. All ligands for docking were drawn using Maestro and prepared using LigPrep [48] to generate the 3D conformation, adjust the protonation state to physiological pH (7.4), and calculate the partial atomic charges with the OPLS4 [49] force field. Docking studies with the prepared ligands were performed using Glide (Glide V7.7) [50,51] with the flexible modality of induced-fit docking with extra precision (XP), followed by a side-chain minimization step using Prime. Ligands were docked within a grid around 12 Å from the centroid of the co-crystallized ligand, generating ten poses per ligand.

### Molecular dynamics simulation

MD simulations were carried out using Desmond [52] with the OPLS4 force-field [49]. The simulated system encompassed the protein-ligand complexes, a predefined water model (TIP3P [53]) as a solvent and counter ions (5 Cl^-^ adjusted to neutralize the overall system charge). The system was treated in a cubic box with periodic boundary conditions specifying the box’s shape and size as 10 Å distance from the box edges to any atom of the protein. In all simulations, we used a time step of 1 fs, the short-range coulombic interactions were treated using a cut-off value of 9.0 Å using the short-range method, while the Smooth Particle Mesh Ewald method (PME) handled long-range coulombic interactions [54]. Initially, the system’s relaxation was performed using Steepest Descent and the limited-memory Broyden-Fletcher-Goldfarb-Shanno algorithms in a hybrid manner, according to the established protocol available in the Desmond standard settings. During the equilibration step, the simulation was performed under the NPT ensemble for 5 ns implementing the Berendsen thermostat and barostat methods [55]. A constant temperature of 310 K was kept throughout the simulation using the Nosé-Hoover thermostat algorithm [56] and Martyna-Tobias-Klein Barostat [57] algorithm to maintain 1 atm of pressure, respectively. After minimization and relaxation of the system, we continued with the production step of at least 500 ns, with frames being recorded/saved every 1,000 ps. Five independent replicas were produced for MR22 (**5**), resulting in a total of 2.5 µs simulation. Trajectories and interaction data are available on the Zenodo repository (DOI: 10.5281/zenodo.7638649). The representative structures were selected by inspecting changes in the Root-mean-square deviation (RMSD). Figure S8 represents the variation of the RMSD values along with the simulation for both template crystal structures and simulations with docking pose. Moreover, the torsional analysis of rotatable bonds of MR22 (**5**) during the MD simulations is shown in Figure S9. Finally, protein-ligand interactions were determined using the Simulation Event Analysis pipeline implemented in Maestro (Maestro v2021.2).

### WaterMap calculation

Relevant cluster structures from the MD simulations were selected for WaterMap calculations. WaterMap [58,59] simulations where non-solvent heavy atoms are restrained were run using default settings with 5.0 ns simulation time with TIP4P water model. Waters within 10.0 Å from the ligand were included in the analysis. All hydration-site information was solely based on the WaterMap simulation as all previous water molecules from the MDs were removed prior to the calculation. Enthalpy values of the hydration sites were obtained by averaging over the non-bonded interaction for each water molecule in the cluster. Entropy and enthalpy values for each hydration site were calculated using inhomogeneous solvation theory.

### PK-Prediction

QikProp (implemented in the *Schrödinger* Drug Discovery suite for molecular modeling, version 2021.2) was used to predict pharmacokinetic properties, and acceptable limits were derived from the references in the manual.

### *In vitro* kinase assay

To measure kinase activity of SIK2, *in vitro* kinase assays were performed as described by Matthess *et al.* [60]. Samples were resolved by sodium dodecyl sulfate-polyacrylamide gel electrophoresis and subjected to autoradiography. Active recombinant human SIK2 was purchased from ProQinase Reaction Biology, product no. 1371-0000-1.

### Cell-cycle analysis

For cell cycle analysis, cells were harvested, washed, fixed with 70% EtOH, and stained as previously described [61]. Cell cycle quantification was performed using FACS Calibur and Cellquest Pro software (both *BD Biosciences*, Franklin Lakes, NJ, USA).

### Immunofluorescence assays

Cells were grown on a glass coverslip, fixed and permeabilized with methanol (−20 °C), and washed with PBS before adding appropriate primary and secondary antibodies. Cells were stained with the following primary antibodies: mouse anti-γ-tubulin (*Sigma*, Germany), rabbit anti-SIK2 (*Cell signaling*). The secondary antibodies: FITC-conjugated goat anti-rabbit, or Cy3 conjugated goat anti-mouse (*Jackson Immunoresearch*, West Grove, PA, USA). DNA was stained using DAPI (4′,6-diamidino-2-phenylindol-dihydrochloride) (*Roche*, Mannheim, Germany). Slides were examined using an Axio Imager 7.1 microscope (*Zeiss*, Göttingen, Germany), and images were taken using an Axio Cam MRm camera (*Zeiss*, Göttingen, Germany).

### Western blot analysis and antibodies

Western blot analysis was performed as previously described [61], using the following antibodies: rabbit polyclonal anti-SIK2 (*Cell Signaling*, Danvers, MA, USA), rabbit polyclonal anti-SIK2-pS385 (*Kinexus*, Sydney, Australia), rabbit polyclonal anti-AKT (*Cell Signaling*), rabbit polyclonal anti-AKT-pS473 (*Cell Signaling*), mouse monoclonal anti-β-actin (*Sigma-Aldrich*), mouse monoclonal anti-cyclin A (*Santa Cruz Biotechnology*, Dallas, TX, USA), mouse monoclonal anti-PLK1 (*Santa Cruz Biotechnology*).

### Viability assessment

To assess cell viability, a live-cell assay based on nuclear morphology was performed as previously described by Tjaden *et al.* [32]. In brief, HCT116 cells were stained with 60 nM Hoechst33342 (*Thermo Scientific*) and seeded at a density of 1200 cells per well (cell culture microplate, PS, f-bottom, µClear®, 781091, *Greiner*) in culture medium (50 µL per well) in a 384-well plate. After 6 h, 12 h, 24 h, 48 h, and 72 h of compound treatment, fluorescence and cellular shape was measured, using the CQ1 high-content confocal microscope (Yokogawa). Compounds were added directly to the cells at three different concentrations (1 µM, 5 µM, and 10 µM). The following parameters were used for image acquisition: Ex 405 nm/Em 447/60 nm, 500 ms, 50%; Ex 561 nm/Em 617/73 nm, 100 ms, 40%; Ex 488/Em 525/50 nm, 50 ms, 40%; Ex 640 nm/Em 685/40, 50 ms, 20%; bright field, 300 ms, 100% transmission, one centered field per well, 7 z stacks per well with 55 µm spacing. Analysis of images was performed using the CellPathfinder software (*Yokogawa*) as described previously [62]. The cell count was normalized against the cell count of cells treated with 0.1% DMSO. Cells showing Hoechst High Intensity Objects were detected. All normal gated cells were further classified in cells containing healthy, fragmented or pyknosed nuclei. Error bars show SEM of biological duplicates.

### Microsomal stability

The solubilized test compound (5 μL, final concentration 10 µM) was preincubated at 37 °C in 432 μL of phosphate buffer (0.1 M, pH 7.4) together with 50 μL NADPH regenerating system (30 mM glucose-6-phosphate, 4 U/mL glucose-6-phosphate dehydrogenase, 10 mM NADP, 30 mM MgCl_2_). After 5 min, the reaction was started by the addition of 13 μL of microsome mix from the liver of Sprague−Dawley rats (Invitrogen; 20 mg protein/mL in 0.1 M phosphate buffer) in a shaking water bath at 37 °C. The reaction was stopped by adding 500 μL of ice-cold methanol at 0, 15, 30, and 60 min. The samples were centrifuged at 5000 g for 5 min at 4 °C, and the supernatants were analyzed and test compound was quantified by HPLC: The composition of the mobile phase is adapted to the test compound in a range of MeOH 40-90% and water (0.1% formic acid) 10-60%; flow-rate: 1 mL/min; stationary phase: Purospher® STAR, RP18, 5 μm, 125×4, precolumn: Purospher® STAR, RP18, 5 μm, 4×4; detection wavelength: 254 and 280 nm; injection volume: 50 μL. Control samples were performed to check the test compound’s stability in the reaction mixture: first control was without NADPH, which is needed for the enzymatic activity of the microsomes, second control was with inactivated microsomes (incubated for 20 min at 90 °C), and third control was without test compound (to determine the baseline). The amounts of the test compound were quantified by an external calibration curve. Data are expressed as the mean ± SEM remaining compound from three independent experiments.

## ASSOCIATED CONTENT

### Supporting Information

The Supporting Information is available free of charge.

Validation of NanoBRET results on SIK1-3; Comparison of scanMAX kinase assay panel and in-house DSF panel; Comparison of published kinome-wide selectivity of G-5555 (**2**) and in-house DSF-panel results; Crystal structure comparison of G-5555 (**2**)/MST3 and MR21 (**4**)/EpHA2 complex; Comparison of binding mode of **9** and **4** in EpHA2; Analysis of the binding mode of **12** in MST3 compared to that of G-5555 (**2**); Processed selectivity data of compounds **4**–**12** obtained with the in-house DSF panel; RMSD values of SIK2-MR22 system; MR22 (**5**) torsional conformations during the MD simulations; NanoBRET off-target evaluation of **4** and **8**; Western blot analysis of downstream AKT activity after chemical inhibition of SIK2; Metabolism of MR22 (**5**); In-house DSF selectivity profiles of compounds **2** and **4**–**12**; NanoBRET assay information; Thermodynamic parameters of hydration sites in WaterMap calculation. X-ray data collection and refinement statistics; Analytical data of all compounds presented. (PDF)

### Author Contributions

M.Rak, R.T., and S.K. designed the project; M. Rak, A.L., and T.H. synthesized the compounds; R.T., D.-I.B. and R.Z. expressed, purified, and co-crystallized MST3 or EpHA2; D.-I.B., R.T., R.Z. A.K. and A.C.J. refined the structures of MST3 and EpHA2 inhibitor complexes; E.S. conceptualized, executed and analyzed the molecular modeling experiments; T.K. helped to conceptualize and write the modeling part; A.P. provided resources for modeling; A.K., L.E. and M.Rak performed DSF assays; L.B. performed NanoBRET assays; A.T. performed cell viability assay; M. Raab, M.S. and K.S. performed auto-phosphorylation and centrosome dissociation assays as well as cell-cycle analysis and Western blotting; and S.K. supervised the research. The manuscript was written by M. Rak, A.C.J., and S.K. with contributions from all coauthors.

### Notes

The authors declare no competing interest.

### Accession Codes

Coordinates and structure factors of the MST3- and EpHA2-inhibitor complexes are available in the Protein Data Bank (PDB) under accession codes 8BZI (MST3-**12** complex), 8BIN (EpHA2-MR21 (**4**) complex) and 8BIO (EpHA2-**9** complex).

The computational datasets generated and/or analysed during the current study are available in the Zenodo repository (DOI: 10.5281/zenodo.7638649).

## ACKNOWLEDGMENT

M. Rak is grateful for the support by the Dr. Hilmer-Stiftung. The authors are grateful to the Structural Genomics Consortium (SGC), a registered charity (No:1097737) that received funds from Bayer AG, Boehringer Ingelheim, Bristol Myers Squibb, Genentech, Genome Canada through Ontario Genomics Institute, Janssen, Merck KGaA, Pfizer, and Takeda. This project received funding from the Innovative Medicines Initiative 2 Joint Undertaking (JU) under grant agreement No. 875510. The JU receives support from the European Union’s Horizon 2020 research and innovation program, EFPIA, Ontario Institute for Cancer Research, Royal Institution for the Advancement of Learning McGill University, Kungliga Tekniska Hoegskolan, and Diamond Light Source Limited. Disclaimer: This communication reflects the views of the authors, and JU is not liable for any use that may be made of the information contained herein. R.T. was supported by the German Research Foundation (DFG) grant 397659447. A.C.J. is supported by German Research Foundation (DFG) grant JO 1473/1-3. The data collection at SLS was supported by funding from the European Union’s Horizon 2020 research and innovation program grant agreement number 730872, project CALIPSOplus. We thank the staff at beamline X06SA of the Swiss Light Source for assistance during data collection. T.K., A.P. and E.S. would like to thank the TϋCAD2, which is funded by the Federal Ministry of Education and Research (BMBF) and the Baden-Württemberg Ministry of Science as part of the Excellence Strategy of the German Federal and State Governments EXC 2180–390900677 T.K. is supported by the Fortüne initiative (No. 2613-0-0) and by the iFIT, which are both initiatives from the Excellence Strategy of the German Federal and State Governments. A.T. is supported by the SFB 1177 ‘Molecular and Functional Characterization of Selective Autophagy. The CQ1 microscope was funded by FUGG (INST 161/920-1 FUGG). We thank the CSC-Finland for the generous computational resources provided. We thank Astrid Kaiser for performing the microsomal stability assay.

## ABBREVIATIONS

ABL1: Abelson murine leukemia viral oncogene homologue 1
AGC: protein kinases A, G and C
AKT: protein kinase B
AMPK: AMP-activated protein kinase
BMPR2: bone morphogenetic protein receptor type-2
BMX: bone marrow tyrosine kinase gene in chromosome X protein
BRAF: serine/threonine protein kinase B-raf
CAMK: calmodulin/calcium-dependent kinases
CDK2: cyclin-dependent kinase 2
CEP250: centrosome-associated protein 250
CK1: casein kinase 1
CREB: cAMP-response element binding protein
CRTC: CREB-regulated transcription coactivators
DSF: differential scanning fluorimetry
EpHA: ephrin type-A receptor
EpHB: ephrin type-B receptor
FDA: food and drug administration
FGFR1: fibroblast growth factor receptor 1
GAK: cyclin-G-associated kinase
GST: glutathione S-transferase
HCC: hepatocellular carcinoma
HDAC: histone deacetylase
HPLC: high-performance liquid chromatography
JNK1: c-Jun N-terminal kinase 1
LKB1: liver kinase B1
MST: mammalian STE20-like protein kinase
NEK1: NimA-related protein kinase 1
PAK: p21-activated kinase
PI3K: phosphoinositide 3-kinase
PKA: protein kinase A
PLK1: polo-like kinase 1
QIK: Qin-induced kinase
RSK1: ribosomal S6 kinase 1
SAR: structure-activity relationship
SIK: salt-inducible kinase
STE: homologous kinases of the yeast proteins STE20, STE11, and STE7
TK: tyrosine kinases
TLK: tyrosine-like kinases
ZAK: human leucine zipper-and sterile alpha motif-containing kinase

