## Supplemental material for "Shifting the selectivity of pyrido[2,3-d]pyrimidin-7(8*H*)-one inhibitors towards the salt-inducible kinase (SIK) subfamily"

### Table of Content

Supplementary Figures S1–S12 .......................................................................................... S3-S16

Table S1: Selectivity screening results of compounds **2** and **4–7** ................................... S17-S19

Table S2: Selectivity screening results of compounds **8–12** ........................................... S20-S23

Table S3: NanoBRET assay information...................................................................... S24

Table S4: Thermodynamic properties of hydration sites in Watermap calculation................... S25

Table S5: X-ray data collection and refinement statistics......................................................... S26

Analytical data for compounds **4–12** and intermediates **21–46**........................................ S27-S90

References.................................................................................................................S91

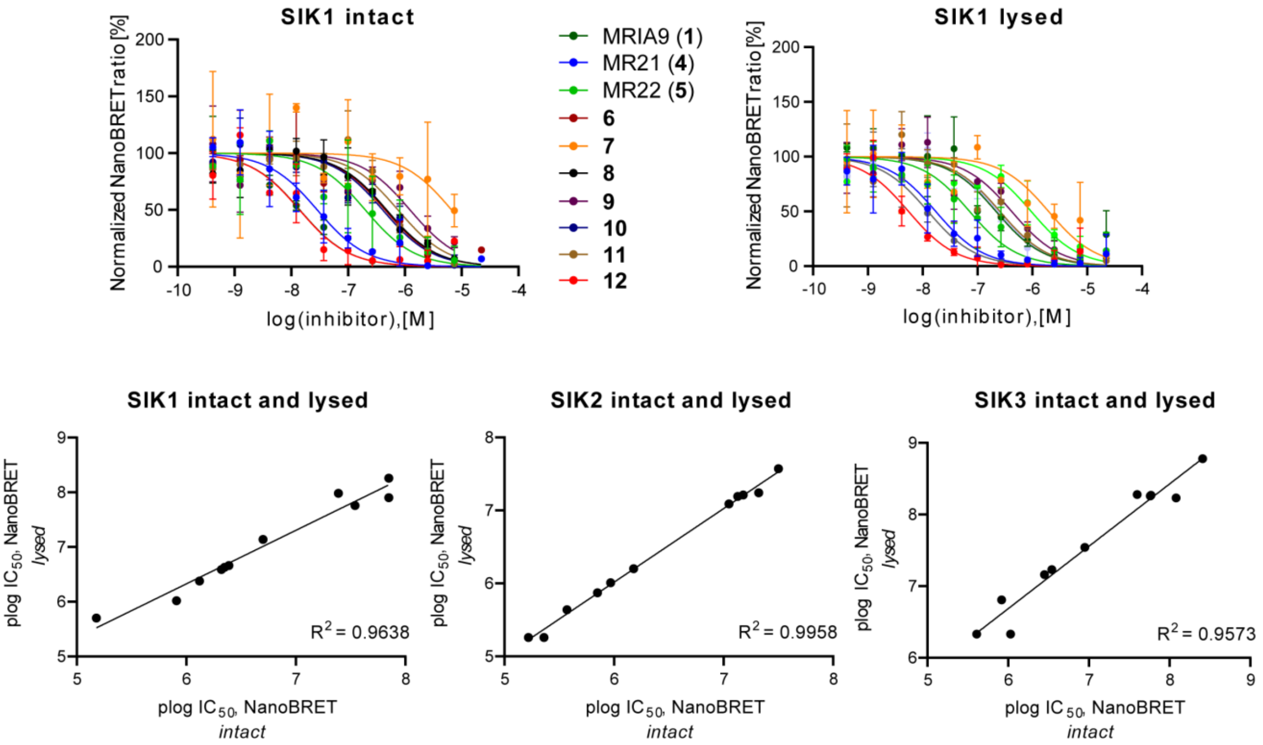


**Figure S1.** Comparison of NanoBRET $IC_{50}$ values for intact and lysed mode measurements of compounds **1** and **4–12** on SIK1-3.

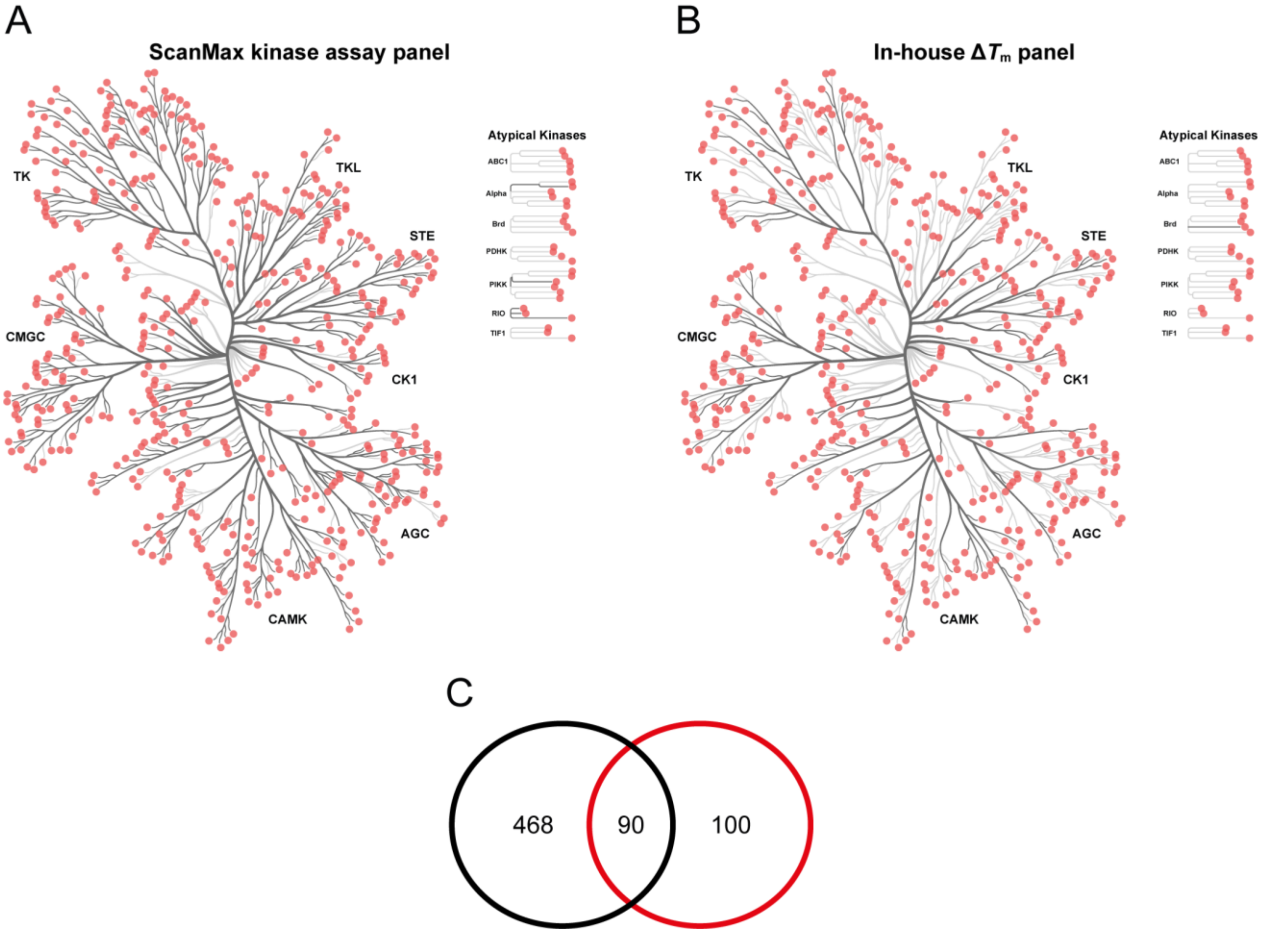


**Figure S2.** Comparison of the kinases included in the scanMAX kinase assay panel (*Eurofins Scientific*) and the in-house DSF assay panel. Both assays are shown as phylogenetic trees using the web-based tool CORAL [1]. Kinases are shown as red circles, and kinases included in the corresponding panel are highlighted as dark gray branches. (A) Phylogenetic tree of the scanMAX panel consisting of 468 kinases, including kinase mutants. (B) Phylogenetic tree of the in-house DSF assay panel consisting of 100 kinases. (C) Number of kinases of the individual panels are shown in the circles, with the scanMAX panel in black and the in-house $\Delta T_m$ panel in red. The overlap of both kinase panels is shown as intersection between the two circles.

Comparison of selectivity data of G-5555 (**2**)

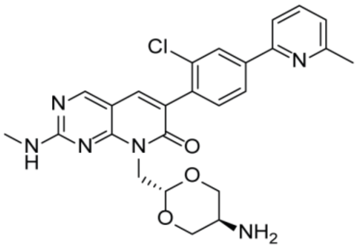

G-5555
(**2**)

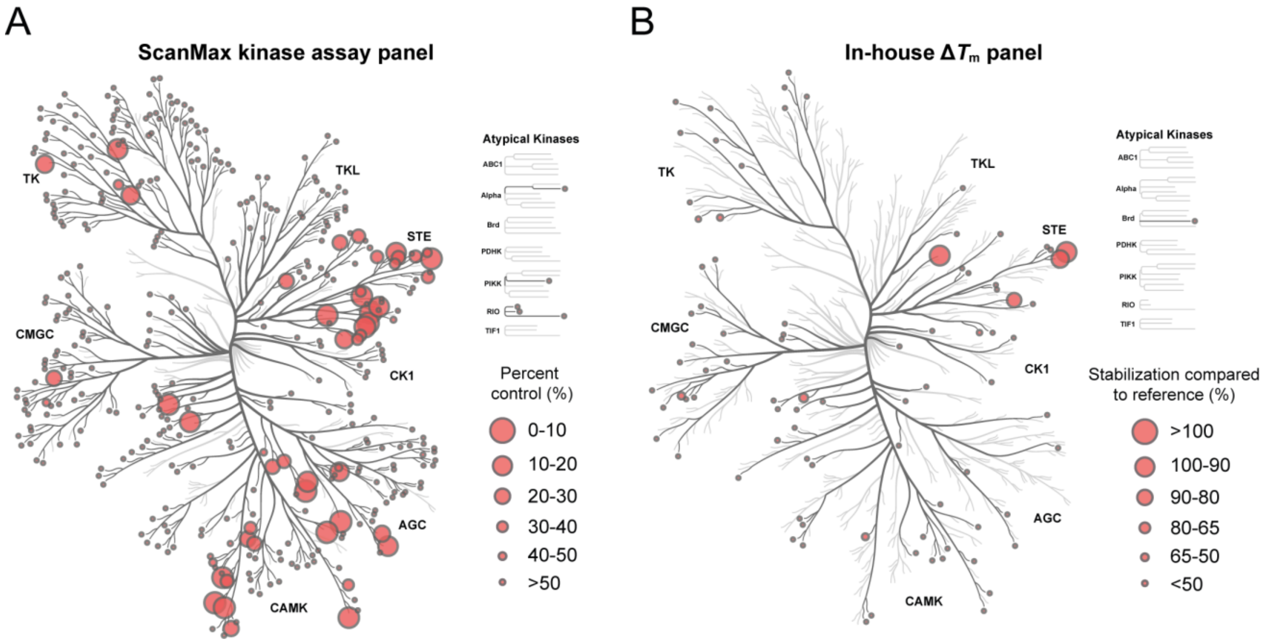


**Figure S3.** Comparison of the published [2] and in-house selectivity data of G-5555 (**2**). Data is illustrated as phylogenetic trees using the web-based tool Coral [1]. Kinases tested are shown as dark gray branches and kinases inhibited by red cycles. (A) Selectivity profile of **2** tested in the scanMAX kinase panel on 468 kinases at a concentration of 1 µM. The circle radius represents the remaining activity in percent compared with the kinase activity in the absence of the inhibitor. (B) Phylogenetic tree of the selectivity profile of **2** investigated in the in-house $\Delta T_m$ panel against 91 kinases. Obtained $\Delta T_m$ values were compared to the value of a corresponding reference compound and are shown in percent (%), represented by the circle radius. G-5555 (**2**) was tested at a final concentration of 10 µM. Data shown in this figure can be found in Table S1.

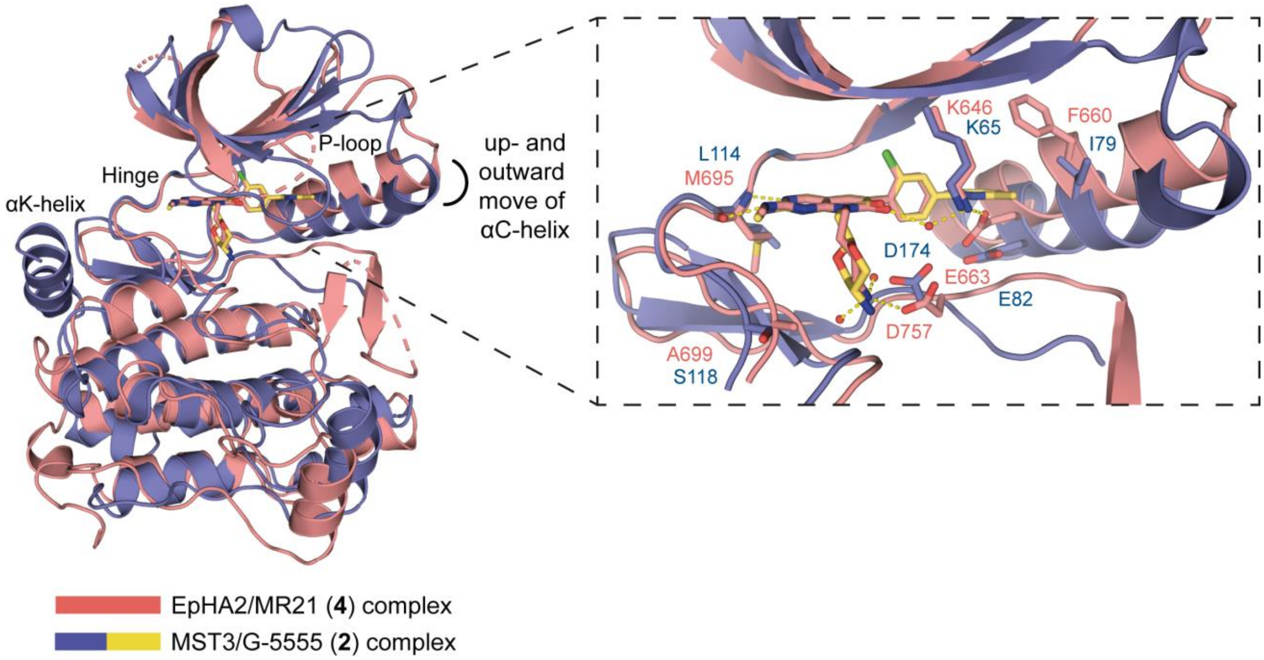


**Figure S4.** Comparison of the co-crystal structures of G-5555 (**2**) with MST3 (PDB code: 7B30) and MR21 (**4**) with EpHA2 (PDB code: 8BIN). The overall structure is shown together with an inset of the binding pocket. MST3/G-5555 (**2**) is colored in blue/yellow and EpHA2/MR21 (**4**) in pink. A different conformation of the αC-helix can be found. Key amino acids forming the binding pocket are highlighted, and hydrogen bonds in the complex of EpHA2 with MR21 (**4**) are shown as yellow dashed lines.

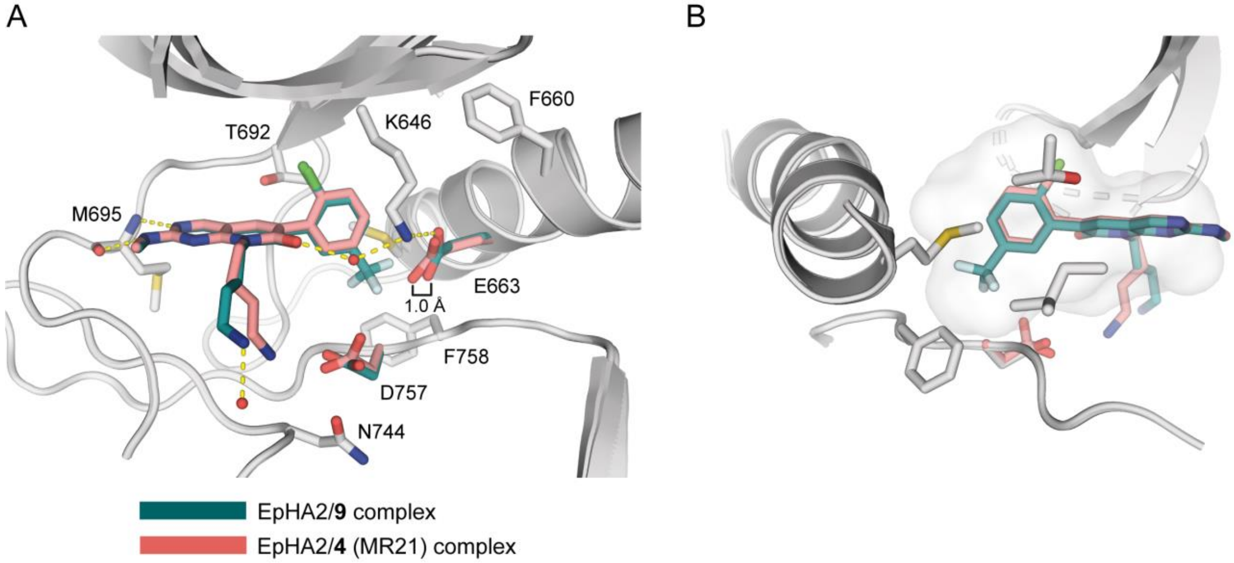


**Figure S5.** Comparison of the co-crystal structures of **9** and **4** (MR21) with EpHA2 (PDB codes 8BIO and 8BIN, respectively). (A) The binding mode of the compounds in the kinase structures is shown as an overlay. Important amino acids in the binding pocket are shown as stick models. (B) Different orientation of the overlay showing the surface of the subpocket that extends next to the gatekeeper. The kinase structures are shown in grey, and differences in the orientation of the amino acids are highlighted in the colors of the corresponding inhibitors, with **9** in dark-turquoise and **4** in pink. Hydrogen bonds between **9** and EpHA2 are shown as yellow dashed lines.

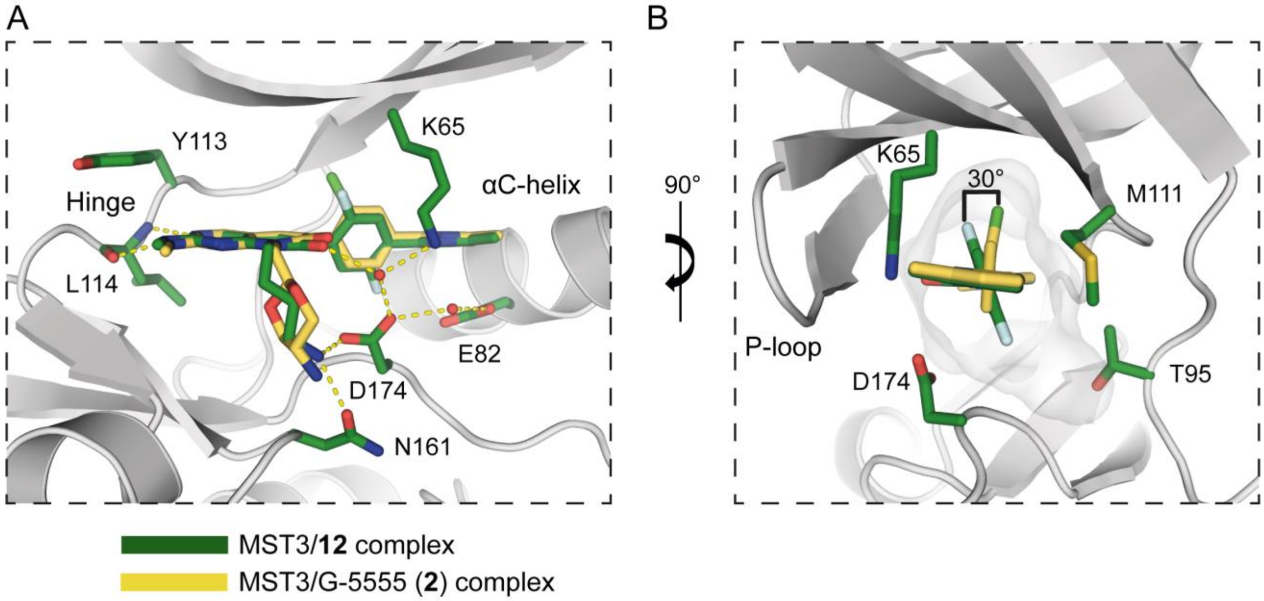


**Figure S6.** Overlay of the crystal structures of MST3 (grey) in complex with **12** (green) and G-5555 (yellow; **2**) (PDB codes 8BZI and 7B30, respectively). (A) The binding mode of the compounds in the ATP binding pocket is shown as an overlay. Amino acids are displayed based on the structure of **12**/MST3 and numbered accordingly. (B) Different orientation of the phenyl rings of **2** and **12** in the back-pocket region next to the gatekeeper. The surface inside the binding pocket is shown in light gray. The 2,5-difluorophenyl group of **12** is rotated by 30° with respect to the 2-chlorophenyl ring of G-5555 (**2**).

### In-house selectivity profile of compounds 4-12

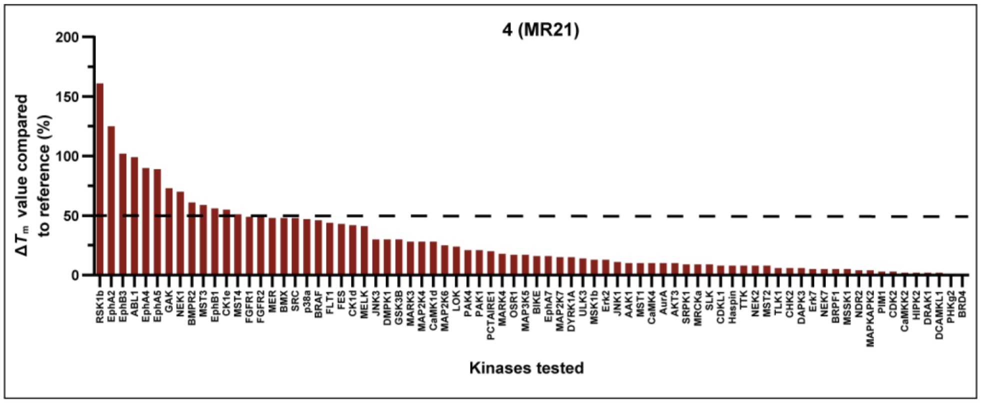


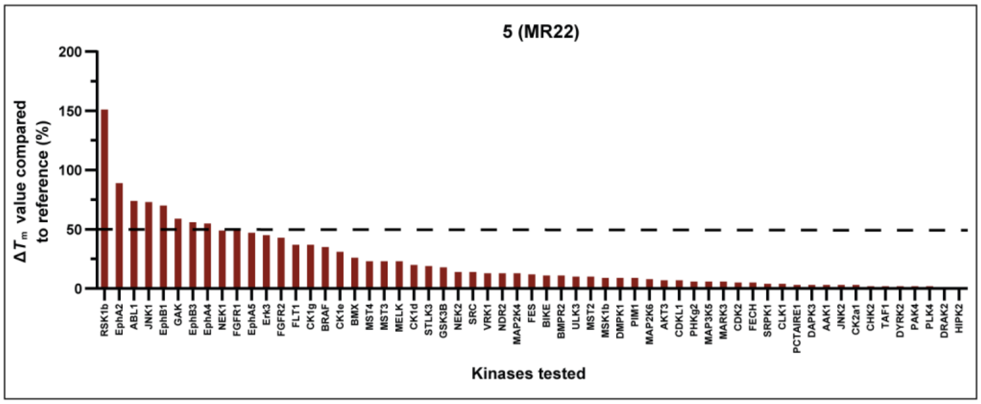


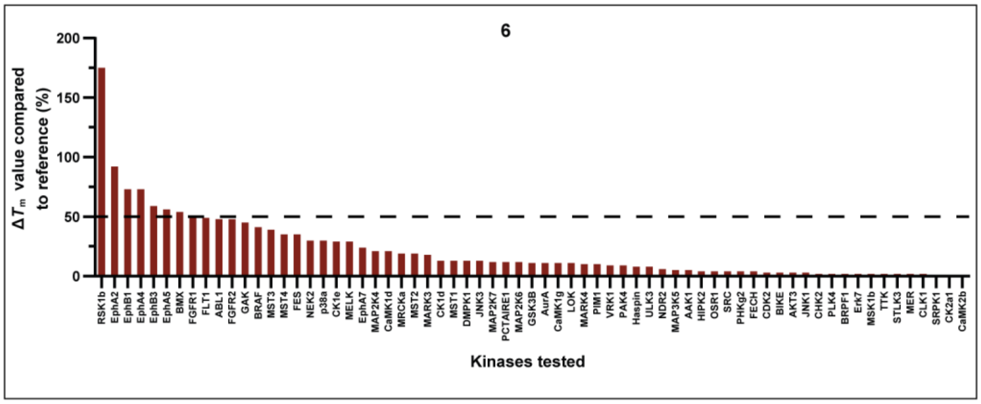


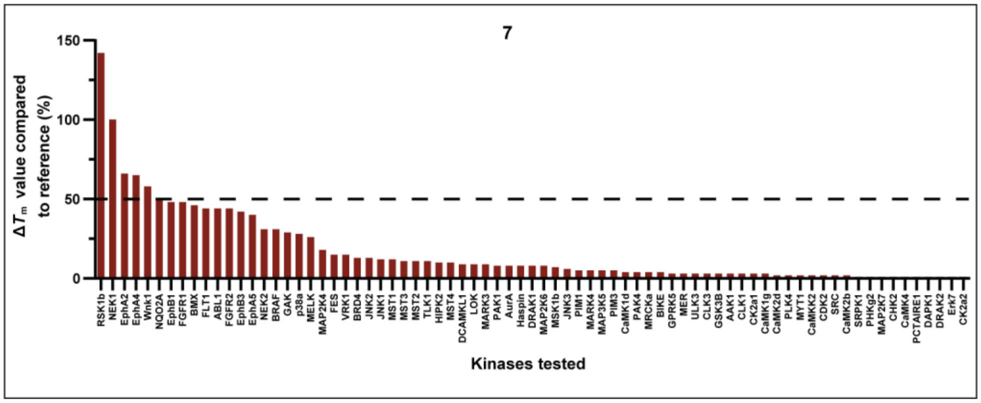

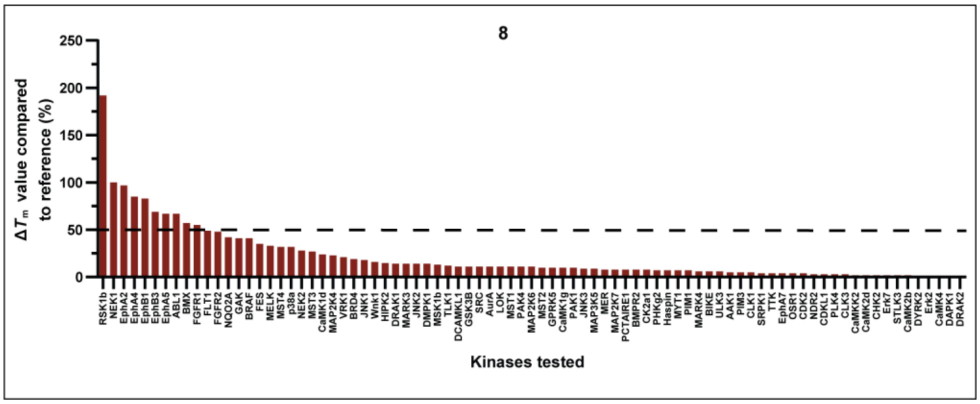
8
ΔTm value compared to reference (%)
250
200
150
100
50
0
Kinases tested
RSK1b NEK1 EphA2 EphA4 EphB1 EphB3 EphA5 ABL1 BMX FGFR1 FLT1 FGFR2 NQO2A GAK BRAF FES MELK MST4 p38a NEK2 MST3 CaMK1d MAP2K4 VRK1 BRD4 JNK1 Wnk1 HIPK2 DRAK1 MARK3 JNK2 DMPK1 MSK1b TLK1 DCAMKL1 GSK3B SRC AurA LOK MST1 PAK4 MAP2K6 MST2 GPRK5 CaMK1g PAK1 JNK3 MAP3K5 MER MAP2K7 PCTAIRE1 BMPR2 CK2a1 PHKg2 Haspin MYT1 PIM1 MARK4 BIKE ULK3 AAK1 PIM3 CLK1 SRPK1 TTK EphA7 OSR1 CDK2 NDR2 CDKL1 PLK4 CLK3 CaMKK2 CaMK2d CHK2 Erk7 STLK3 CaMK2b DYRK2 Erk2 CaMK4 DAPK1 DRAK2

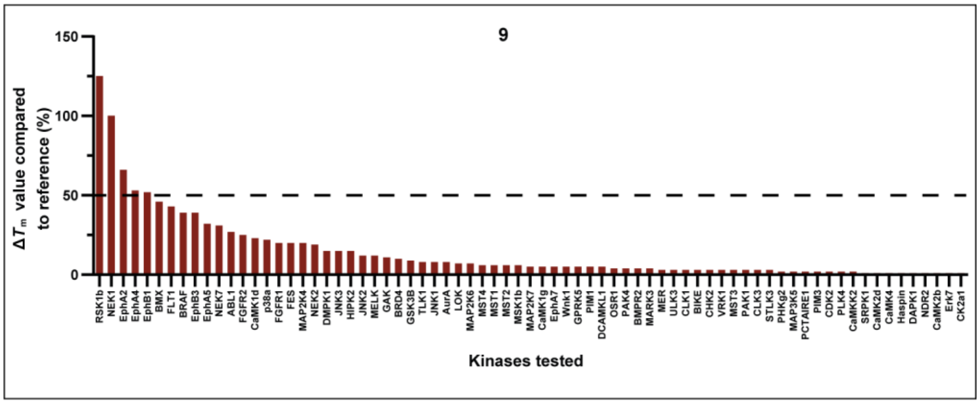
9
ΔTm value compared to reference (%)
150
100
50
0
Kinases tested
RSK1b NEK1 EphA2 EphA4 EphB1 BMX FLT1 BRAF EphB3 EphA5 NEK7 ABL1 FGFR2 CaMK1d p38a FGFR1 FES MAP2K4 NEK2 DMPK1 JNK3 HIPK2 JNK2 MELK GAK BRD4 GSK3B TLK1 JNK1 AurA LOK MAP2K6 MST4 MST1 MST2 MSK1b CaMK1g EphA7 Wnk1 GPRK5 PIM1 DCAMKL1 OSR1 PAK4 BMPR2 MARK3 MER ULK3 CLK1 BIKE CHK2 VRK1 MST3 PAK1 CLK3 STLK3 PHKg2 MAP3K5 PCTAIRE1 PIM3 CDK2 PLK4 CaMKK2 SRPK1 CaMK2d CaMK4 Haspin DAPK1 NDR2 CaMK2b Erk7 CK2a1

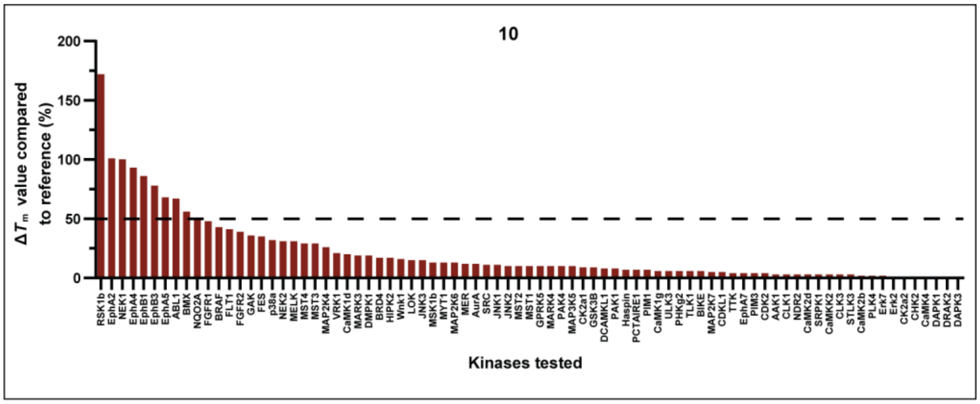
10
ΔTm value compared to reference (%)
200
150
100
50
0
Kinases tested
RSK1b EphA2 NEK1 EphA4 EphB1 EphB3 EphA5 ABL1 BMX NQO2A FGFR1 BRAF FLT1 FGFR2 GAK FES p38a NEK2 MELK MST4 MST3 MAP2K4 VRK1 CaMK1d MARK3 DMPK1 BRD4 HIPK2 Wnk1 LOK JNK3 MSK1b MYT1 MAP2K6 MER AurA SRC JNK1 JNK2 MST2 MST1 GPRK5 MARK4 PAK4 MAP3K5 CK2a1 GSK3B DCAMKL1 PAK1 Haspin PCTAIRE1 PIM1 CaMK1g ULK3 PHKg2 TLK1 BIKE MAP2K7 CDKL1 TTK EphA7 PIM3 CDK2 AAK1 CLK1 NDR2 CaMK2d SRPK1 CaMKK2 CLK3 STLK3 CaMK2b PLK4 Erk7 Erk2 CK2a2 CHK2 CaMK4 DAPK1 DRAK2 DAPK3

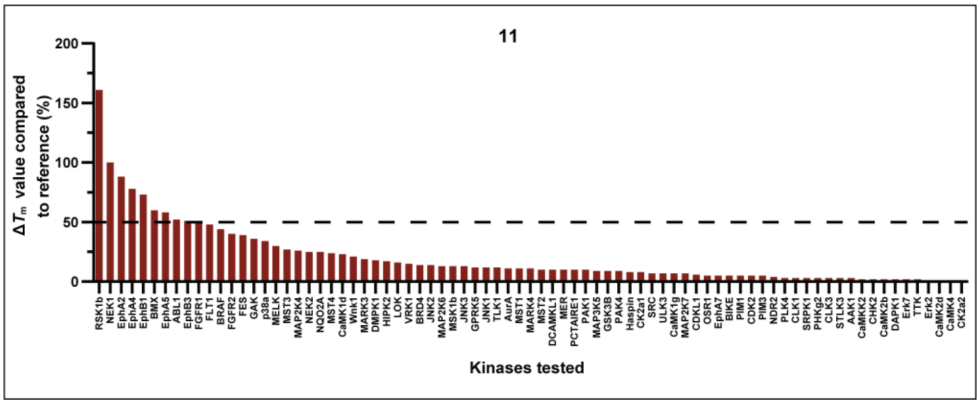
11
ΔTm value compared to reference (%)
200
150
100
50
0
Kinases tested
RSK1b NEK1 EphA2 EphA4 EphB1 BMX EphA5 ABL1 EphB3 FGFR1 FLT1 BRAF FGFR2 FES GAK p38a MELK MST3 MAP2K4 NEK2 NQO2A MST4 CaMK1d Wnk1 MARK3 DMPK1 HIPK2 LOK VRK1 BRD4 JNK2 MAP2K6 MSK1b JNK3 GPRK5 JNK1 TLK1 AurA MST1 MARK4 MST2 DCAMKL1 MER PCTAIRE1 PAK1 MAP3K5 GSK3B PAK4 Haspin CK2a1 SRC ULK3 CaMK1g MAP2K7 CDKL1 OSR1 EphA7 BIKE PIM1 CDK2 PIM3 NDR2 PLK4 CLK1 SRPK1 PHKg2 CLK3 STLK3 AAK1 CaMKK2 CHK2 CaMK2b DAPK1 Erk7 TTK Erk2 CaMK2d CaMK4 CK2a2

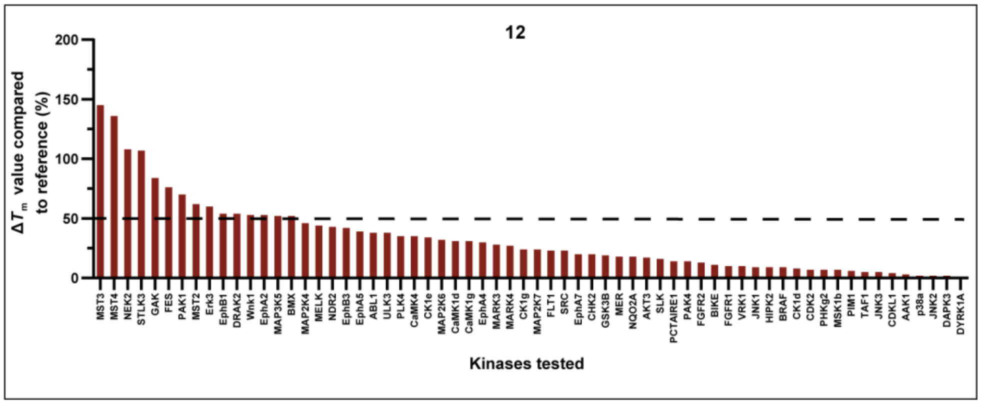


**Figure S7.** In-house DSF panel results for compounds **4–12** on up to 100 kinases, shown as bar graphs. Obtained $\Delta T_m$ values were compared with the values of corresponding reference compounds and are shown in percent (%). Only kinases that were stabilized are shown. Data included in this figure can be found in Tables S1 and S2. The average of two measurements is shown. Compounds **4–12** were tested at a final concentration of 10 µM. The 50% stabilization threshold used as off-target criterion is indicated as a dashed line.

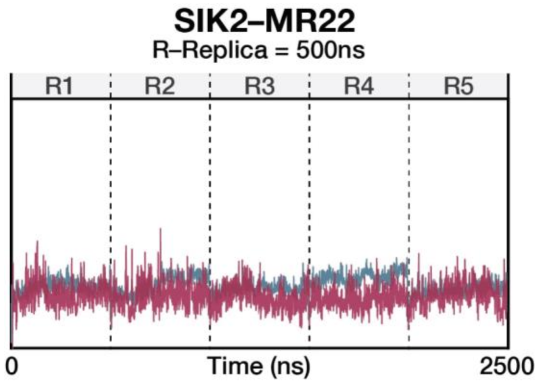


**Figure S8.** RMSD values of the SIK2-MR22 system in MD simulations. RMSDs were calculated for the Cα-atoms. The dashed lines indicate the replicas (1 replica = 500 ns)

**SIK2-MR22 (5)**

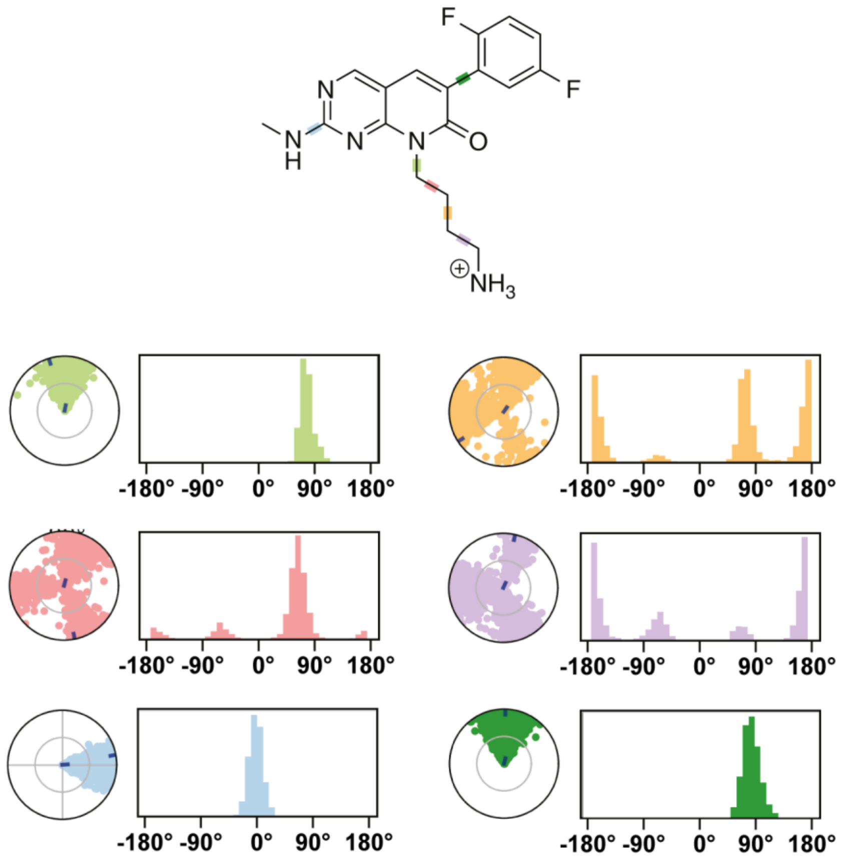


**Figure S9**. MR22 (**5**) torsional conformations of rotatable bonds during the MD simulations (5 replicas x 500 ns = 2500 ns). The polar plots show the conformation of MR22 (**5**) as a function of time, where the radial coordinate is the simulation time, and the angular coordinate the torsional angle. The bar charts show the probability of the torsions as a function of angle. They represent the angle over the simulation time. The color of the plot matches the color-coding of the rotatable bond on the MR22 (**5**) structure.

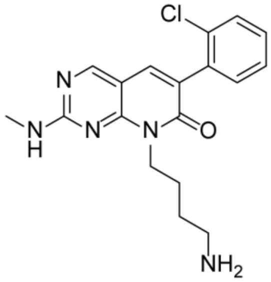

MR21
(**4**)

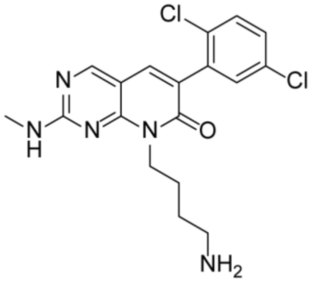

**8**

| Targets | $\Delta T_m$ value compared to reference (%)[a] | $IC_{50}$ (nM)[b] NanoBRET |
|---|---|---|
| SIK1 | n.d. | 20 ± 7[c] |
| SIK2 | n.d. | 74 ± 2 |
| SIK3 | n.d. | 25 ± 3 |
| RSK1b | 161 | >45000 |
| EpHA2 | 125 | 210 ± 36 |
| EpHB3 | 102 | 1500 ± 100 |
| ABL1 | 99 | 4600 ± 400 |
| EpHA4 | 90 | 66 ± 38 |
| EpHA5 | 89 | 928 ± 6 |
| GAK | 73 | 23500 ± 2500 |
| NEK1 | 70 | n.d. |
| BMPR2 | 61 | n.d. |
| MST3 | 59 | n.d. |
| EpHB1 | 56 | 78 ± 2 |
| CK1e | 55 | n.d. |
| MST4 | 51 | n.d. |
| EpHA1 | n.d. | 12 ± 2 |
| EpHA3 | n.d. | 313 ± 121 |
| EpHB2 | n.d. | 1300 ± 200 |
| EpHB4 | n.d. | 50 ± 12 |

| Targets | $\Delta T_m$ value compared to reference (%)[a] | $IC_{50}$ (nM)[b] NanoBRET |
|---|---|---|
| SIK1 | n.d. | 277 ± 123[c] |
| SIK2 | n.d. | 665 ± 40 |
| SIK3 | n.d. | 113 ± 95 |
| RSK1b | 192 | >45000 |
| NEK1 | 100 | n.d. |
| EpHA2 | 97 | 1400 ± 200 |
| EpHA4 | 85 | 346 ± 1 |
| EpHB1 | 83 | 302 ± 26 |
| EpHB3 | 69 | 6500 ± 600 |
| EpHA5 | 67 | 3000 ± 600 |
| ABL1 | 67 | >45000 |
| BMX | 57 | 2300 ± 500 |
| FGFR1 | 55 | n.d. |
| EpHA1 | n.d. | 82 ± 3 |
| EpHA3 | n.d. | 1600 ± 400 |
| EpHB2 | n.d. | 6200 ± 1400 |
| EpHB4 | n.d. | 188 ± 17 |

**Figure S10.** Cellular off-target evaluation of **4** and **8** in the NanoBRET assay. Selectivity data of compounds **4** (MR21) and **8** in a DSF against a panel of up to 100 kinases. The obtained $\Delta T_m$ values of each kinase are shown in % compared to a corresponding reference compound. Kinases with a $\Delta T_m$ value > 50% as well as SIK1-3 and EpHA1, EpHA3, EpHB2, and EpHB4 were further analysed, and $IC_{50}$ values were determined in a cellular NanoBRET assay. [a]Average of two measurements. Compounds MR21 (**4**) and **8** were tested at a final concentration of 10 µM. Raw data can be found in Table S1. [b]$IC_{50}$ values were determined in a 11-point dose-response curve (duplicate measurements) and are presented as standard deviation of the mean. [c]$IC_{50}$ values against SIK1 were measured in lysed mode.

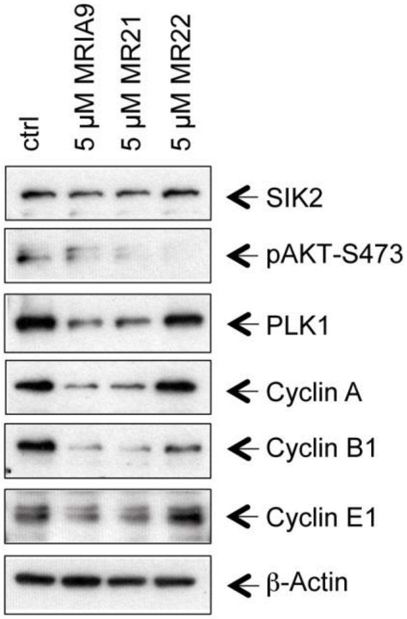


**Figure S11.** Effects of MRIA9 (**1**), MR21 (**4**), and MR22 (**5**) on SIK2 downstream AKT phosphorylation levels. SKOV-3 cells were treated with 2 nM rapamycin for 16 h and subsequently incubated with the corresponding compound for 48 hours at a concentration of 5 µM. Cells were lysed and immunoblotted with specific antibodies against SIK2, pAKT-S473, PLK1, and Cyclin A, B1 and E1, as well as β-Actin.

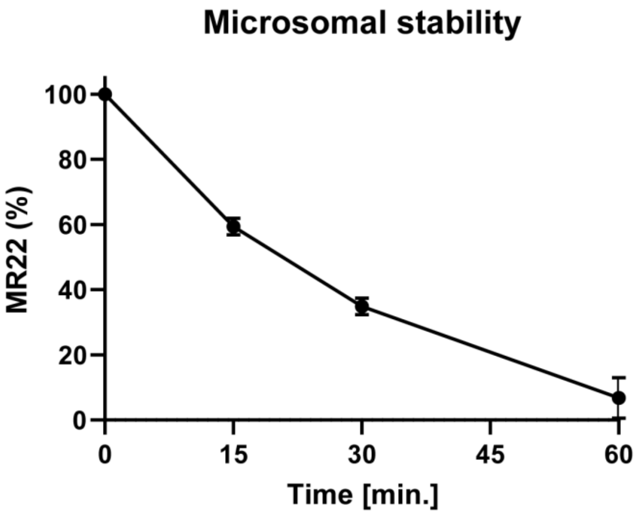


| Descriptor | Interpretation | Limit Values | G-5555[a] | MRIA9[a] | MR22[+] |
|---|---|---|---|---|---|
| $P_{Caco-2}$ | Predicted apparent Caco-2 cell permeability in nm/sec[b]. Caco-2 cells are a model for the gut-blood barrier | <25 poor, >500 great | 140.1 | 92.2 | 196.0 |
| logB/B | Predicted brain/blood partition coefficient[c] | -3.0 – 1.2 | -0.8 | -0.8 | -0.7 |
| $P_{MDCK}$ | Predicted apparent MDCK cell permeability in nm/sec[b]. MDCK cells are considered to be a good mimic for the blood-brain barrier | <25 poor, >500 great | 125.4 | 121.6 | 476.8 |
| $\log K_p$ | Predicted skin permeability | -8.0 – -1.0 | -4.4 | -4.7 | -3.4 |
| $\log K_{hsa}$ | Prediction of binding to human serum albumin | -1.5 – 1.5 | 0.1 | 0.0 | 1.0 |
| Oral Abs. | Predicted human oral absorption on 0 to 100% scale. The prediction is based on a quantitative multiple linear regression model | >80% is high <25% is poor | 81 | 76 | 75 |

**Figure S12.** Microsomal stability and prediction of pharmacokinetic properties of MR22 (**5**). Microsomal stability was measured by incubation of **5** with activated liver microsomes, monitoring its stability by HPLC over a time of one hour. Prediction of ADME properties was performed using the software QikProp (*Schrödinger, LLC, New York, NY, 2021*). [a]Published predictions for G-5555 (**2**) and MRIA9 (**1**) [2]. [b]Predictions are for non-active transport. [c]Predictions are for orally delivered drugs so, for example, dopamine and serotonin are CNS negative because they are too polar to cross the blood-brain barrier.

**Table S1**. Selectivity screening of compounds **2** and **4–7** in an in-house $\Delta T_m$ panel against up to 100 kinases. Common abbreviations are used for the kinases. The corresponding reference compound is shown with its $\Delta T_m$ shift. Average of two measurements. Compounds **2** and **4–7** were tested at a final concentration of 10 µM. (N.D. = not determined)

| | | | **$\Delta T_m$ [°C]** | | | | |
|---|---|---|---|---|---|---|---|
| **Kinase** | **Reference compound** | **$\Delta T_m$ shift reference compound** | **G-5555 (2)** | **MR21 (4)** | **MR22 (5)** | **6** | **7** |
| **AAK1** | Staurosporine | 15.3 | 0.6 | 1.6 | 0.4 | 0.7 | 0.5 |
| **ABL1** | Staurosporine | 9.3 | 5.1 | 9.2 | 6.9 | 4.5 | 4.1 |
| **AKT3** | Staurosporine | 7.0 | 1.4 | 0.7 | 0.5 | 0.2 | 0.0 |
| **AurA** | Staurosporine | 17.2 | 8.5 | 1.7 | -3.0 | 1.9 | 1.4 |
| **BIKE** | Staurosporine | 19.0 | 2.5 | 3.1 | 2.1 | 0.6 | 0.7 |
| **BMPR2** | Lestaurtinib | 2.6 | -0.4 | 1.6 | 0.3 | -0.3 | 0.0 |
| **BMX** | Staurosporine | 7.0 | 1.3 | 3.3 | 1.9 | 3.8 | 3.2 |
| **BRAF** | Dabrafenib | 26.6 | 3.9 | 12.2 | 9.2 | 10.8 | 8.3 |
| **BRD4** | JQ1 | 7.0 | -1.1 | 0.1 | -2.8 | -0.1 | 0.9 |
| **BRPF1** | GSK6853 | 14.0 | 0.8 | 0.7 | -1.0 | 0.3 | -0.7 |
| **CaMK1d** | Staurosporine | 9.9 | 4.5 | 2.7 | -1.2 | 2.1 | 0.4 |
| **CaMK1g** | Staurosporine | 11.0 | 2.3 | -0.1 | -0.4 | 1.2 | 0.3 |
| **CaMK2b** | Staurosporine | 13.2 | -1.7 | -1.1 | -0.6 | 0.1 | 0.2 |
| **CaMK2d** | Staurosporine | 16.5 | -0.1 | 0.0 | -0.6 | 0.0 | 0.4 |
| **CaMK4** | Staurosporine | 8.4 | 0.7 | 0.8 | -0.6 | -0.1 | 0.1 |
| **CaMKK2** | Staurosporine | 24.5 | 1.1 | 0.6 | -0.1 | -0.2 | 0.5 |
| **CASK** | Staurosporine | 5.0 | -0.8 | 0.0 | -0.4 | -0.3 | -0.1 |
| **CDK2** | Staurosporine | 15.0 | 0.5 | 0.4 | 0.7 | 0.5 | 0.3 |
| **CDKL1** | CEP-32496 | 6.6 | -0.1 | 0.5 | 0.5 | 0.0 | -0.1 |
| **CHK2** | Staurosporine | 16.7 | 0.9 | 0.9 | 0.4 | 0.4 | 0.2 |
| **CK1d** | SR-3029 | 6.8 | 0.4 | 2.9 | 1.3 | 0.9 | -0.1 |
| **CK1e** | PF-670462 | 9.0 | N.D. | 4.9 | 2.8 | 2.6 | N.D. |
| **CK1g** | Staurosporine | 2.0 | N.D. | N.D. | 0.7 | N.D. | N.D. |
| **CK2a1** | Similtasertib | 14.6 | 0.1 | -0.4 | 0.4 | 0.2 | 0.4 |
| **CK2a2** | Similtasertib | 15.7 | -0.6 | -0.3 | -1.0 | -0.1 | 0.1 |
| **CLK1** | Staurosporine | 12.3 | 0.1 | -1.2 | 0.4 | 0.2 | 0.4 |
| **CLK3** | T3 | 15.0 | 0.1 | 0.0 | -0.4 | -0.3 | 0.5 |

| | | | | | | | |
|---|---|---|---|---|---|---|---|
| **DAPK1** | Staurosporine | 9.4 | -1.0 | -0.2 | -0.7 | -0.2 | 0.1 |
| **DAPK3** | Staurosporine | 16.2 | 0.5 | 0.9 | 0.5 | -0.2 | -0.4 |
| **DCAMKL1** | Staurosporine | 10.7 | 0.3 | 0.2 | -1.7 | -0.5 | 1.0 |
| **DMPK1** | Staurosporine | 9.4 | -0.3 | 2.8 | 0.9 | 1.2 | -0.1 |
| **DRAK1** | Staurosporine | 7.7 | -0.7 | 0.1 | -0.7 | -0.1 | 0.6 |
| **DRAK2** | Staurosporine | 10.6 | -0.3 | -0.5 | 0.1 | -0.2 | 0.1 |
| **DYRK1A** | Staurosporine | 9.6 | -1.0 | 1.4 | 0.0 | 0.0 | -0.6 |
| **DYRK2** | Staurosporine | 7.3 | -0.1 | 0.0 | 0.2 | -0.1 | 0.0 |
| **EphA2** | Staurosporine | 7.3 | 2.2 | 9.1 | 6.5 | 6.7 | 4.8 |
| **EphA4** | Staurosporine | 5.5 | N.D. | 4.9 | 3.0 | 4.0 | 3.6 |
| **EphA5** | Staurosporine | 8.1 | 2.7 | 7.2 | 3.8 | 4.5 | 3.2 |
| **EphA7** | Staurosporine | 11.3 | 4.1 | 1.8 | -1.6 | 2.7 | -0.5 |
| **EphB1** | Staurosporine | 6.4 | N.D. | 3.6 | 4.5 | 4.7 | 3.1 |
| **EphB3** | Staurosporine | 5.9 | 1.7 | 6.0 | 3.3 | 3.5 | 2.5 |
| **Erk2** | GDC-0994 | 7.5 | 1.4 | 1.0 | -3.7 | 0.0 | 0.0 |
| **Erk3** | Staurosporine | 1.2 | 0.3 | N.D. | 0.5 | N.D. | N.D. |
| **Erk7** | Staurosporine | 14.2 | 0.3 | 0.8 | -1.5 | 0.3 | 0.1 |
| **FECH** | Vemurafenib | 5.5 | N.D. | -1.1 | 0.3 | 0.2 | -1.1 |
| **FES** | Staurosporine | 7.1 | 4.1 | 3.0 | 0.9 | 2.5 | 1.1 |
| **FGFR1** | Staurosporine | 6.0 | -0.1 | 3.3 | 3.4 | 3.0 | 2.9 |
| **FGFR2** | Staurosporine | 8.5 | 0.7 | 4.7 | 3.7 | 4.1 | 3.7 |
| **FLT1** | Staurosporine | 12.2 | 1.7 | 5.4 | 4.6 | 6.0 | 5.4 |
| **GAK** | Staurosporine | 9.1 | 6.1 | 6.6 | 5.4 | 4.1 | 2.6 |
| **GPRK5** | Staurosporine | 5.9 | -0.2 | -1.2 | -0.6 | 0.0 | 0.2 |
| **GSK3B** | Staurosporine | 9.0 | 0.1 | 2.7 | 1.6 | 1.0 | 0.3 |
| **Haspin** | Staurosporine | 8.9 | 0.2 | 0.7 | -0.3 | 0.7 | 0.7 |
| **HIPK2** | Staurosporine | 4.8 | N.D. | 0.1 | 0.1 | 0.2 | 0.5 |
| **JNK1** | Staurosporine | 7.6 | 4.7 | 0.9 | 5.5 | 0.2 | 0.9 |
| **JNK2** | SBI-0069279 | 8.6 | 0.6 | -0.1 | 0.2 | -0.4 | 1.1 |
| **JNK3** | CEP-32496 | 8.8 | 1.8 | 2.7 | -0.2 | 1.1 | 0.5 |
| **LOK** | Staurosporine | 22.8 | 10.2 | 5.4 | -0.4 | 2.4 | 2.1 |
| **MAP2K4** | Staurosporine | 11.7 | 6.4 | 3.3 | 1.5 | 2.5 | 2.1 |
| **MAP2K6** | Staurosporine | 12.0 | 4.4 | 3.1 | 1.0 | 1.4 | 0.9 |
| **MAP2K7** | Staurosporine | 7.3 | N.D. | 1.1 | -0.9 | 0.9 | 0.1 |
| **MAP3K5** | Staurosporine | 17.5 | 8.9 | 2.9 | 1.0 | 0.9 | 0.8 |
| **MAPKAPK 2** | Staurosporine | 3.4 | -0.3 | 0.1 | -0.1 | -0.7 | 0.0 |

| | | | | | | | |
|---|---|---|---|---|---|---|---|
| **MARK3** | Staurosporine | 19.1 | 8.8 | 5.3 | 1.1 | 3.5 | 1.7 |
| **MARK4** | Staurosporine | 16.8 | 7.7 | 3.0 | 0.0 | 1.7 | 0.8 |
| **MELK** | Staurosporine | 13.9 | 8.4 | 5.7 | 3.2 | 4.0 | 3.6 |
| **MER** | Staurosporine | 5.9 | 0.8 | 2.8 | -0.8 | 0.1 | 0.2 |
| **MRCKa** | Staurosporine | 2.6 | 0.9 | 0.2 | -0.5 | 0.5 | 0.1 |
| **MSK1b** | Staurosporine | 15.6 | 2.0 | 2.1 | 1.5 | 0.3 | 1.1 |
| **MSSK1** | Staurosporine | 3.7 | N.D. | 0.2 | -0.1 | N.D. | N.D. |
| **MST1** | Staurosporine | 14.7 | 4.5 | 1.5 | -0.3 | 1.9 | 1.7 |
| **MST2** | Staurosporine | 13.4 | 6.9 | 1.0 | 1.4 | 2.5 | 1.5 |
| **MST3** | Staurosporine | 7.0 | 7.5 | 4.1 | 1.6 | 2.7 | 0.8 |
| **MST4** | Staurosporine | 6.2 | 6.1 | 3.2 | 1.4 | 2.2 | 0.6 |
| **MYT1** | Dasatinib | 4.5 | -2.1 | -2.3 | -2.1 | -1.4 | 0.1 |
| **NDR2** | Staurosporine | 12.3 | 3.5 | 0.5 | 1.6 | 0.7 | -0.1 |
| **NEK1** | Staurosporine | 0.1 | -2.2 | 0.1 | 0.0 | -0.3 | 0.1 |
| **NEK2** | Staurosporine | 6.7 | 3.5 | 0.5 | 1.0 | 2.0 | 2.1 |
| **NEK7** | Staurosporine | 1.3 | 0.1 | 0.1 | -1.3 | -1.0 | -0.2 |
| **NQO2A** | Staurosporine | 1.2 | N.D. | -1.0 | -0.2 | 0.0 | 0.6 |
| **OSR1** | Staurosporine | 7.4 | 0.2 | 1.3 | -0.8 | 0.3 | 0.0 |
| **p38a** | Doramapimod | 18.9 | 1.4 | 8.8 | -0.6 | 5.6 | 5.3 |
| **PAK1** | Staurosporine | 7.3 | 6.4 | 1.6 | -0.5 | 0.0 | 0.6 |
| **PAK4** | Staurosporine | 12.9 | 3.5 | 2.8 | 0.2 | 1.1 | 0.5 |
| **PCTAIRE1** | Staurosporine | 9.0 | 1.3 | 1.8 | 0.3 | 1.1 | 0.1 |
| **PHKG2** | Staurosporine | 21.8 | 0.6 | 0.3 | 1.4 | 0.8 | 0.3 |
| **PIM1** | Staurosporine | 12.2 | -1.6 | 0.4 | 1.1 | 1.2 | 0.6 |
| **PIM3** | Staurosporine | 19.8 | -1.8 | -0.8 | -0.6 | -0.4 | 0.9 |
| **PLK4** | Staurosporine | 17.9 | 3.4 | 0.0 | 0.3 | 0.4 | 0.4 |
| **RSK1b** | Staurosporine | 3.6 | 0.0 | 5.8 | 5.4 | 6.3 | 5.1 |
| **SLK** | Staurosporine | 18.4 | 7.2 | 1.6 | -0.6 | N.D. | N.D. |
| **SRC** | Staurosporine | 5.4 | -0.2 | 2.6 | 0.7 | 0.2 | 0.1 |
| **SRPK1** | Staurosporine | 6.7 | -0.4 | 0.6 | 0.3 | 0.1 | 0.1 |
| **STLK3** | Staurosporine | 11.4 | 0.6 | -1.1 | 2.1 | 0.2 | 0.0 |
| **TAF1** | Bromosporine | 7.4 | 0.5 | -0.1 | 0.2 | -0.6 | -0.8 |
| **TLK1** | Staurosporine | 8.5 | -0.3 | 0.5 | -1.3 | 0.0 | 0.9 |
| **TTK** | Staurosporine | 11.2 | 1.5 | 0.9 | -2.6 | 0.2 | -0.2 |
| **ULK3** | Staurosporine | 17.8 | 8.0 | 2.4 | 1.9 | 1.4 | 0.6 |
| **VRK1** | Staurosporine | 3.4 | -0.1 | -0.4 | 0.4 | 0.3 | 0.5 |
| **Wnk1** | Staurosporine | 1.9 | 2.4 | -0.4 | -0.1 | 0.0 | 1.1 |

**Table S2**. Selectivity screening of compounds **8–12** in an in-house $\Delta T_m$ panel against up to 100 kinases. Common abbreviations are used for the kinases. The corresponding reference compound is shown with its $\Delta T_m$ shift. Average of two measurements. Compounds **8–12** were tested at a final concentration of 10 µM. (N.D. = not determined)

| | | | **$\Delta T_m$ [°C]** | | | | |
|---|---|---|---|---|---|---|---|
| **Kinase** | **Reference compound** | **$\Delta T_m$ shift reference compound** | **8** | **9** | **10** | **11** | **12** |
| **AAK1** | Staurosporine | 15.3 | 0.8 | 0.0 | 0.5 | 0.4 | 0.5 |
| **ABL1** | Staurosporine | 9.3 | 6.2 | 2.5 | 6.2 | 4.8 | 3.5 |
| **AKT3** | Staurosporine | 7.0 | 0.0 | 0.0 | -0.1 | -0.1 | 1.2 |
| **AurA** | Staurosporine | 17.2 | 1.9 | 1.3 | 2.0 | 1.9 | -2.4 |
| **BIKE** | Staurosporine | 19.0 | 1.1 | 0.6 | 1.1 | 1.0 | 2.2 |
| **BMPR2** | Lestaurtinib | 2.6 | 0.2 | 0.1 | 0.0 | 0.0 | -0.2 |
| **BMX** | Staurosporine | 7.0 | 4.0 | 3.2 | 3.9 | 4.2 | 3.6 |
| **BRAF** | Dabrafenib | 26.6 | 10.8 | 10.4 | 11.4 | 11.6 | 2.3 |
| **BRD4** | JQ1 | 7.0 | 1.3 | 0.7 | 1.2 | 1.0 | -2.0 |
| **BRPF1** | GSK6853 | 14.0 | -0.5 | -0.4 | -0.3 | -0.7 | -1.5 |
| **CaMK1d** | Staurosporine | 9.9 | 2.4 | 2.3 | 2.0 | 2.3 | 3.1 |
| **CaMK1g** | Staurosporine | 11.0 | 1.1 | 0.6 | 0.7 | 0.8 | 3.4 |
| **CaMK2b** | Staurosporine | 13.2 | 0.2 | 0.1 | 0.3 | 0.3 | -0.4 |
| **CaMK2d** | Staurosporine | 16.5 | 0.4 | 0.2 | 0.5 | 0.2 | -0.2 |
| **CaMK4** | Staurosporine | 8.4 | 0.1 | 0.1 | 0.1 | 0.1 | 2.9 |
| **CaMKK2** | Staurosporine | 24.5 | 0.6 | 0.4 | 0.7 | 0.6 | 0.0 |
| **CASK** | Staurosporine | 5.0 | -0.2 | -0.1 | 0.0 | -0.3 | -0.2 |
| **CDK2** | Staurosporine | 15.0 | 0.6 | 0.3 | 0.6 | 0.7 | 1.1 |
| **CDKL1** | CEP-32496 | 6.6 | 0.2 | 0.0 | 0.3 | 0.4 | 0.2 |
| **CHK2** | Staurosporine | 16.7 | 0.4 | 0.5 | 0.2 | 0.4 | 3.3 |
| **CK1d** | SR-3029 | 6.8 | 0.0 | 0.0 | 0.0 | 0.0 | 0.6 |
| **CK1e** | PF-670462 | 9.0 | N.D. | N.D. | N.D. | N.D. | 3.1 |
| **CK1g** | Staurosporine | 2.0 | N.D. | N.D. | N.D. | N.D. | 0.5 |
| **CK2a1** | Silmitasertib | 14.6 | 1.1 | 0.1 | 1.3 | 1.1 | -0.2 |
| **CK2a2** | Silmitasertib | 15.7 | 0.0 | 0.0 | 0.2 | 0.1 | -1.0 |

| | | | | | | | |
|---|---|---|---|---|---|---|---|
| **CLK1** | Staurosporine | 12.3 | 0.6 | 0.4 | 0.4 | 0.4 | -0.1 |
| **CLK3** | T3 | 15.0 | 0.4 | 0.4 | 0.4 | 0.4 | -0.3 |
| **DAPK1** | Staurosporine | 9.4 | 0.1 | 0.1 | 0.1 | 0.2 | -0.9 |
| **DAPK3** | Staurosporine | 16.2 | -0.1 | -0.1 | 0.1 | -0.2 | 0.3 |
| **DCAMKL1** | Staurosporine | 10.7 | 1.2 | 0.5 | 0.9 | 1.1 | -1.6 |
| **DMPK1** | Staurosporine | 9.4 | 1.3 | 1.4 | 1.8 | 1.7 | -0.1 |
| **DRAK1** | Staurosporine | 7.7 | 1.1 | -1.1 | -0.8 | -0.8 | 0.0 |
| **DRAK2** | Staurosporine | 10.6 | 0.1 | 0.0 | 0.1 | 0.0 | 5.7 |
| **DYRK1A** | Staurosporine | 9.6 | -0.5 | -0.5 | -0.6 | -0.3 | 0.1 |
| **DYRK2** | Staurosporine | 7.3 | 0.1 | 0.0 | 0.0 | 0.0 | -0.2 |
| **EphA2** | Staurosporine | 7.3 | 7.1 | 4.8 | 7.4 | 6.4 | 3.9 |
| **EphA4** | Staurosporine | 5.5 | 4.7 | 2.9 | 5.1 | 4.3 | 1.6 |
| **EphA5** | Staurosporine | 8.1 | 5.4 | 2.6 | 5.5 | 4.7 | 3.2 |
| **EphA7** | Staurosporine | 11.3 | 0.5 | 0.6 | 0.5 | 0.6 | 2.3 |
| **EphB1** | Staurosporine | 6.4 | 5.3 | 3.3 | 5.5 | 4.7 | 3.5 |
| **EphB3** | Staurosporine | 5.9 | 4.1 | 2.3 | 4.6 | 3.0 | 2.5 |
| **Erk2** | GDC-0994 | 7.5 | 0.1 | 0.0 | 0.1 | 0.1 | -1.8 |
| **Erk3** | Staurosporine | 1.2 | N.D. | N.D. | N.D. | N.D. | 0.7 |
| **Erk7** | Staurosporine | 14.2 | 0.3 | 0.1 | 0.3 | 0.3 | -1.3 |
| **FECH** | Vemurafenib | 5.5 | -1.7 | -1.7 | -1.7 | -1.6 | -0.6 |
| **FES** | Staurosporine | 7.1 | 2.5 | 1.4 | 2.5 | 2.8 | 5.4 |
| **FGFR1** | Staurosporine | 6.0 | 3.3 | 1.2 | 2.9 | 3.0 | 0.6 |
| **FGFR2** | Staurosporine | 8.5 | 4.1 | 2.1 | 3.3 | 3.4 | 1.1 |
| **FLT1** | Staurosporine | 12.2 | 6.0 | 5.3 | 5.0 | 5.8 | 2.8 |
| **GAK** | Staurosporine | 9.1 | 3.7 | 1.0 | 3.3 | 3.3 | 7.6 |
| **GPRK5** | Staurosporine | 5.9 | 0.6 | 0.3 | 0.6 | 0.7 | -0.4 |
| **GSK3B** | Staurosporine | 9.0 | 1.0 | 0.8 | 0.8 | 0.8 | 1.7 |
| **Haspin** | Staurosporine | 8.9 | 0.6 | 0.1 | 0.6 | 0.7 | -0.4 |
| **HIPK2** | Staurosporine | 4.8 | 0.7 | 0.7 | 0.8 | 0.8 | 0.4 |
| **JNK1** | Staurosporine | 7.6 | 1.4 | 0.6 | 0.8 | 0.9 | 0.7 |
| **JNK2** | SBI-0069279 | 8.6 | 1.2 | 1.0 | 0.9 | 1.2 | 0.2 |
| **JNK3** | CEP-32496 | 8.8 | 0.8 | 1.3 | 1.3 | 1.1 | 0.4 |
| **LOK** | Staurosporine | 22.8 | 2.5 | 1.6 | 3.5 | 3.6 | -0.4 |
| **MAP2K4** | Staurosporine | 11.7 | 2.7 | 2.3 | 3.0 | 3.0 | 5.4 |

| | | | | | | | |
|---|---|---|---|---|---|---|---|
| **MAP2K6** | Staurosporine | 12.0 | 1.3 | 0.8 | 1.5 | 1.6 | 3.8 |
| **MAP2K7** | Staurosporine | 7.3 | 0.6 | 0.4 | 0.4 | 0.5 | 1.7 |
| **MAP3K5** | Staurosporine | 17.5 | 1.5 | 0.4 | 1.7 | 1.6 | 9.1 |
| **MAPKAPK2** | Staurosporine | 3.4 | 0.0 | -0.1 | 0.0 | 0.0 | -0.3 |
| **MARK3** | Staurosporine | 19.1 | 2.7 | 0.7 | 3.7 | 3.6 | 5.3 |
| **MARK4** | Staurosporine | 16.8 | 1.0 | 0.0 | 1.7 | 1.8 | 4.5 |
| **MELK** | Staurosporine | 13.9 | 4.6 | 1.6 | 4.3 | 4.2 | 6.1 |
| **MER** | Staurosporine | 5.9 | 0.5 | 0.2 | 0.7 | 0.6 | 1.1 |
| **MRCKa** | Staurosporine | 2.6 | 0.0 | -0.5 | -0.2 | -0.2 | -0.8 |
| **MSK1b** | Staurosporine | 15.6 | 2.1 | 0.9 | 2.1 | 2.0 | 1.0 |
| **MSSK1** | Staurosporine | 3.7 | N.D. | N.D. | N.D. | N.D. | -0.2 |
| **MST1** | Staurosporine | 14.7 | 1.6 | 0.9 | 1.5 | 1.6 | -0.7 |
| **MST2** | Staurosporine | 13.4 | 1.4 | 0.8 | 1.4 | 1.4 | 8.3 |
| **MST3** | Staurosporine | 7.0 | 1.9 | 0.2 | 2.0 | 1.9 | 10.2 |
| **MST4** | Staurosporine | 6.2 | 2.0 | 0.4 | 1.8 | 1.5 | 8.4 |
| **MYT1** | Dasatinib | 4.5 | 0.3 | -0.2 | 0.6 | -0.4 | -1.8 |
| **NDR2** | Staurosporine | 12.3 | 0.4 | 0.1 | 0.4 | 0.5 | 5.2 |
| **NEK1** | Staurosporine | 0.1 | 0.1 | 0.1 | 0.1 | 0.1 | -0.4 |
| **NEK2** | Staurosporine | 6.7 | 1.9 | 1.3 | 2.1 | 1.7 | 7.2 |
| **NEK7** | Staurosporine | 1.3 | -0.2 | 0.4 | -0.5 | -0.2 | -2.1 |
| **NQO2A** | Staurosporine | 1.2 | 0.5 | -0.3 | 0.6 | 0.3 | 0.2 |
| **OSR1** | Staurosporine | 7.4 | 0.3 | 0.3 | -0.2 | 0.4 | -0.2 |
| **p38a** | Doramapimod | 18.9 | 6.0 | 4.2 | 6.1 | 6.4 | 0.4 |
| **PAK1** | Staurosporine | 7.3 | 0.7 | 0.2 | 0.6 | 0.7 | 5.1 |
| **PAK4** | Staurosporine | 12.9 | 1.4 | 0.5 | 1.3 | 1.1 | 1.8 |
| **PCTAIRE1** | Staurosporine | 9.0 | 0.7 | 0.2 | 0.6 | 0.9 | 1.3 |
| **PHKG2** | Staurosporine | 21.8 | 1.5 | 0.5 | 1.3 | 0.6 | 1.5 |
| **PIM1** | Staurosporine | 12.2 | 0.8 | 0.6 | 0.8 | 0.6 | 0.8 |
| **PIM3** | Staurosporine | 19.8 | 1.0 | 0.4 | 0.8 | 0.9 | -1.2 |
| **PLK4** | Staurosporine | 17.9 | 0.5 | 0.3 | 0.4 | 0.6 | 6.3 |
| **RSK1b** | Staurosporine | 3.6 | 6.9 | 4.5 | 6.2 | 5.8 | -0.2 |
| **SLK** | Staurosporine | 18.4 | N.D. | N.D. | N.D. | N.D. | 2.9 |
| **SRC** | Staurosporine | 5.4 | 0.6 | 0.0 | 0.6 | 0.4 | 1.2 |
| **SRPK1** | Staurosporine | 6.7 | 0.3 | 0.1 | 0.2 | 0.2 | 0.0 |

| | | | | | | | |
|---|---|---|---|---|---|---|---|
| **STLK3** | Staurosporine | 11.4 | 0.2 | 0.3 | 0.3 | 0.3 | 12.1 |
| **TAF1** | Bromosporine | 7.4 | -0.7 | -0.7 | -0.8 | -0.8 | 0.4 |
| **TLK1** | Staurosporine | 8.5 | 1.0 | 0.7 | 0.5 | 1.0 | -1.5 |
| **TTK** | Staurosporine | 11.2 | 0.5 | -0.5 | 0.5 | 0.2 | -0.7 |
| **ULK3** | Staurosporine | 17.8 | 1.0 | 0.6 | 1.1 | 1.3 | 6.7 |
| **VRK1** | Staurosporine | 3.4 | 0.7 | 0.1 | 0.7 | 0.5 | 0.3 |
| **Wnk1** | Staurosporine | 1.9 | 0.3 | 0.1 | 0.3 | 0.4 | 1.0 |

**Table S3**. NanoBRET assay information. Tracer K4, K5, K10, and K11 have the following ordering numbers at Promega: N2482, N2492, N2642, and N2652, respectively.

| Protein Kinase | Alias | Catalog #/ CAS # | NanoLuc orientation | Tracer, used | Tracer $K_{D,app}$ [nM] | Tracer, used [nM] |
|---|---|---|---|---|---|---|
| SIK1 | | NV2031 | N | K10 | 66 | 100 |
| SNF1LK2 | SIK2 | NV2061 | N | K10 | 24 | 100 |
| SIK3 | | NV2041 | N | K10 | 224 | 200 |
| ABL1 | | NV1011 | N | K4 | 247 | 250 |
| BMX | | NV1101 | C | K10 | 352 | 350 |
| GAK | | NV1421 | N | K5 | 125 | 150 |
| MAPK8 | JNK1 | NV1701 | N | K5 | 279 | 300 |
| RPS6KA1 | RSK1b | NV1981 | N | K10 | 42 | 50 |
| EPHA1 | | NV1221 | C | K10 | 477 | 500 |
| EPHA2 | | NV1231 | C | K4 | 94 | 100 |
| EPHA3 | | NV3061 | C | K11 | 188 | 200 |
| EPHA4 | | NV1241 | C | K10 | 851 | 900 |
| EPHA5 | | NV1251 | C | K4 | 46 | 100 |
| EPHB1 | | NV3071 | C | K10 | 699 | 700 |
| EPHB2 | | NV1291 | C | K4 | 34 | 100 |
| EPHB3 | | NV1301 | C | K4 | 387 | 400 |
| EPHB4 | | NV1311 | C | K10 | 552 | 500 |

**Table S4.** Thermodynamic parameters for the solvation of the 15 hydration sites within the MR22 (**5**) binding pocket in SIK2. Occupancy is calculated from the number of water-oxygen atoms found occupying a given hydration site during the five ns of molecular dynamics simulation. Enthalpic energy (ΔH), entropic energy (-TΔS) and free energy value (ΔG) are given in kcal/mol. Sites with positive ΔH values correspond to waters that are enthalpically less stable in the protein binding site than in bulk water. Water molecules in the protein binding site generally have lower entropies than in bulk water and thus have positive -TΔS values. Greater values signify water molecules with more substantial entropic penalties, which are less likely to remain stable in the binding site.

| **Hydration Site index** | **Thermodynamic properties** | | |
|---|---|---|---|
| | **ΔG** | **ΔH** | **-TΔS** |
| **1** | 4.12 | 2.78 | 1.34 |
| **2** | 4.20 | 2.23 | 1.97 |
| **3** | 2.92 | 1.86 | 1.07 |
| **4** | 3.26 | 2.43 | 0.83 |
| **5** | 2.75 | 1.62 | 1.12 |
| **6** | 3.60 | 1.23 | 2.38 |
| **7** | 4.59 | 2.43 | 2.16 |
| **8** | 1.20 | -0.34 | 1.54 |
| **9** | 2.55 | 1.73 | 0.83 |
| **10** | 3.37 | 2.61 | 0.76 |
| **11** | 1.00 | 0.08 | 0.92 |
| **12** | -0.04 | -1.28 | 1.23 |
| **13** | 0.95 | -0.04 | 1.00 |
| **14** | 3.33 | 2.45 | 0.88 |
| **15** | 1.15 | -0.27 | 1.42 |

**Table S5.** X-ray data collection and refinement statistics.

| | EPHA2-**9** | EPHA2-MR21 (**4**) | MST3-**12** |
|---|---|---|---|
| **Data collection** | | | |
| Space group | $P2_1$ | $P2_1$ | C2 |
| Molecules in asymmetric unit | 1 | 1 | 1 |
| *a, b, c* (Å) | 32.84, 107.41, 40.65 | 32.80, 107.68, 40.66 | 98.79, 58.75, 61.47 |
| α, β, γ (°) | 90.00, 109.04, 90.00 | 90.00, 108.94, 90.00 | 90.00, 93.05, 90.00 |
| Resolution (Å)[a] | 38.4-1.60 (1.63-1.60) | 38.5-1.50 (1.53-1.50) | 39.5-1.72 (1.75-1.72) |
| Unique reflections[a] | 22,7442 (6,314) | 24,7667 (2,624) | 36,820 (1,919) |
| Completeness (%)[a] | 96.5 (74.0) | 84.2 (35.3) | 98.4 (96.9) |
| Multiplicity[a] | 6.7 (4.8) | 6.9 (3.6) | 6.9 (6.6) |
| Mean $I/\sigma(I)$[a] | 15.0 (1.9) | 14.8 (2.1) | 18.4 (2.2) |
| $R_{pim}$ | 0.027 (0.366) | 0.037 (0.318) | 0.020 (0.335) |
| CC1/2[a] | 0.999 (0.720) | 0.998 (0.825) | 0.999 (0.811) |
| **Refinement** | | | |
| $R_{work}$, (%)[b] | 16.4 | 13.2 | 15.7 |
| $R_{free}$, (%)[b] | 20.1 | 17.4 | 19.9 |
| No. of atoms[c] | 2424 | 2289 | 2345 |
| Overall B-factor (Å$^2$) | 21.3 | 17.9 | 36.5 |
| RMSD bond lengths (Å) | 0.007 | 0.009 | 0.007 |
| RMSD bond angles (°) | 1.32 | 1.49 | 0.90 |
| Ramachandran favored (%)[d] | 96.1 | 97.2 | 97.0 |
| Ramachandran outliers (%)[d] | 0.0 | 0.0 | 0.0 |
| **Protein Data Bank entry** | 8BIO | 8BIN | 8BZI |

[a]Values in parentheses are for the highest-resolution shell.
[b]$R_{work}$ and $R_{free} = \sum||F_{obs}| - |F_{calc}||/\sum|F_{obs}|$, where $R_{free}$ was calculated with 5% of the reflections chosen at random and not used in the refinement.
[c]Number includes alternative conformations.
[d]MolProbity statistics [3].

### Analytical data for compounds 4–12 and intermediates 21–46

$^1$H NMR compound **21**:

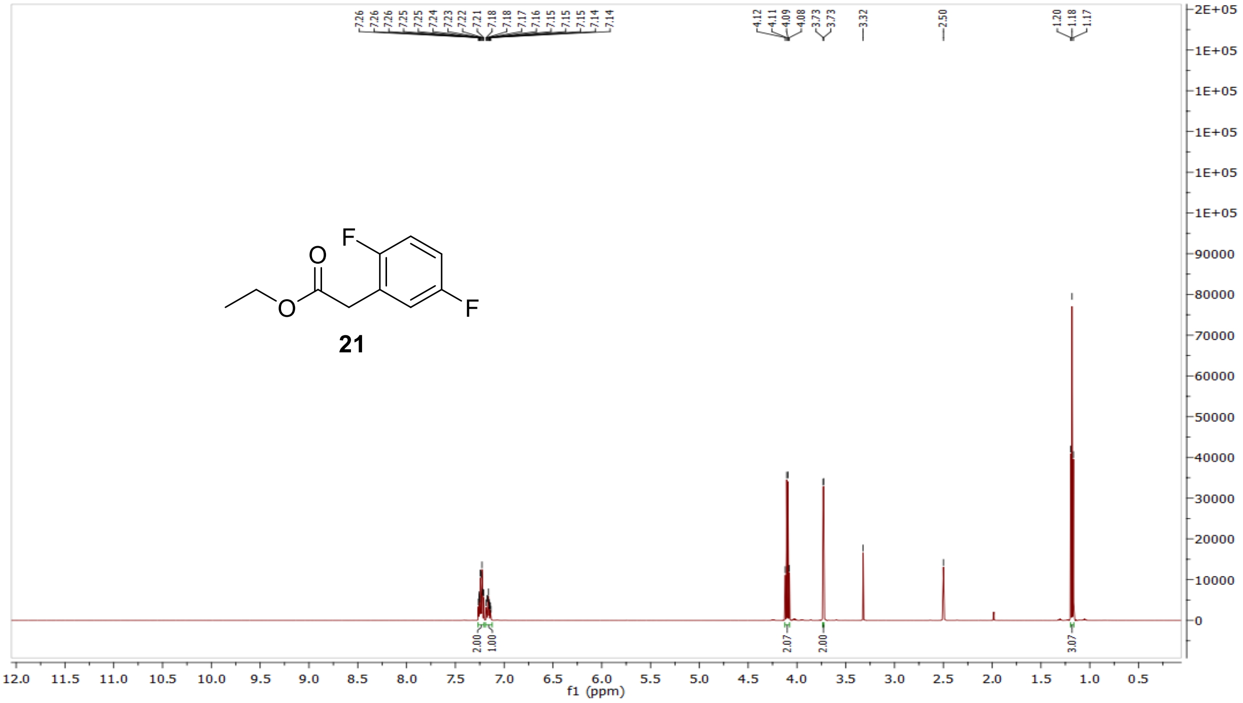


$^{13}$C NMR compound **21:**

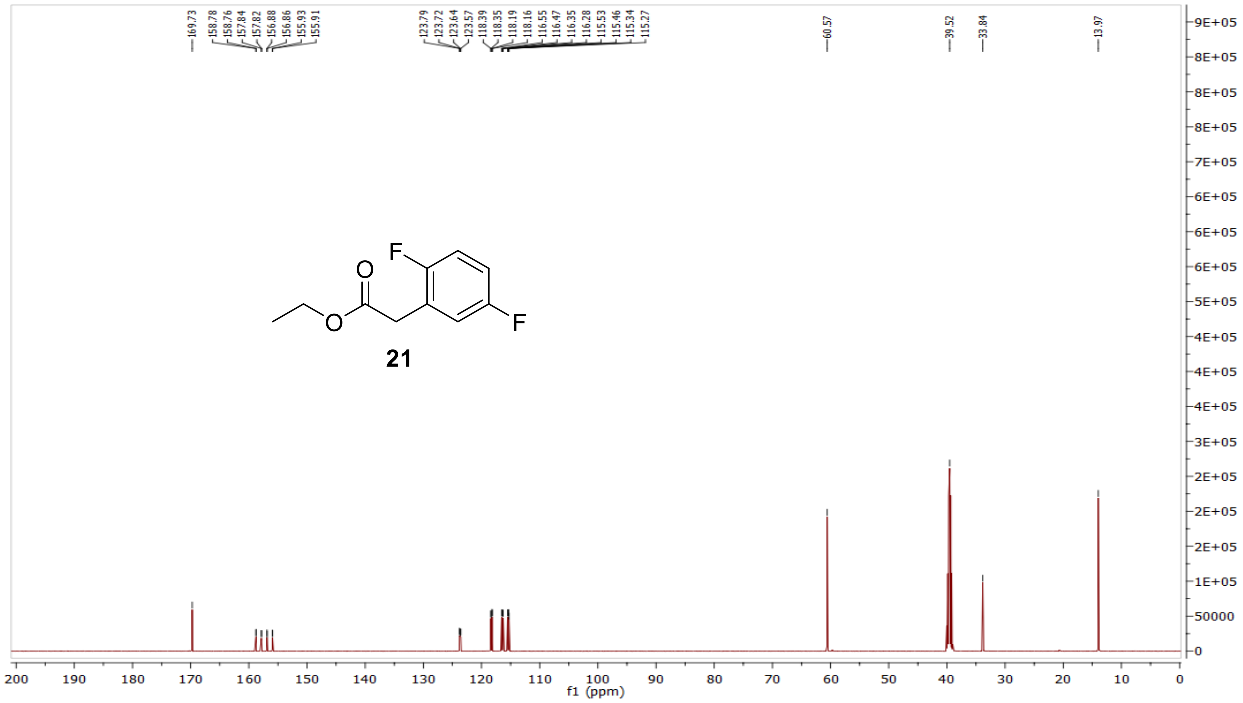

MS (ESI+) compound **21**:

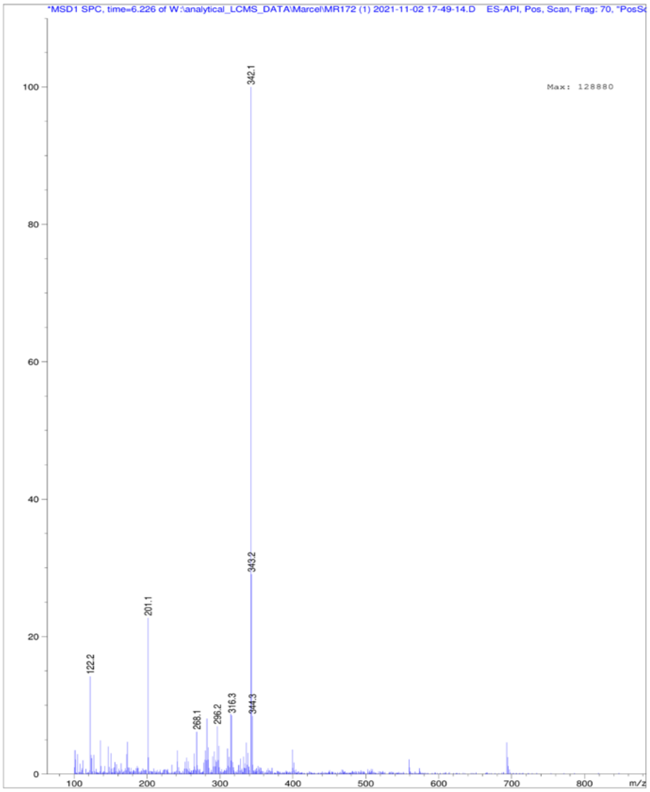


[1]H NMR compound **22**:

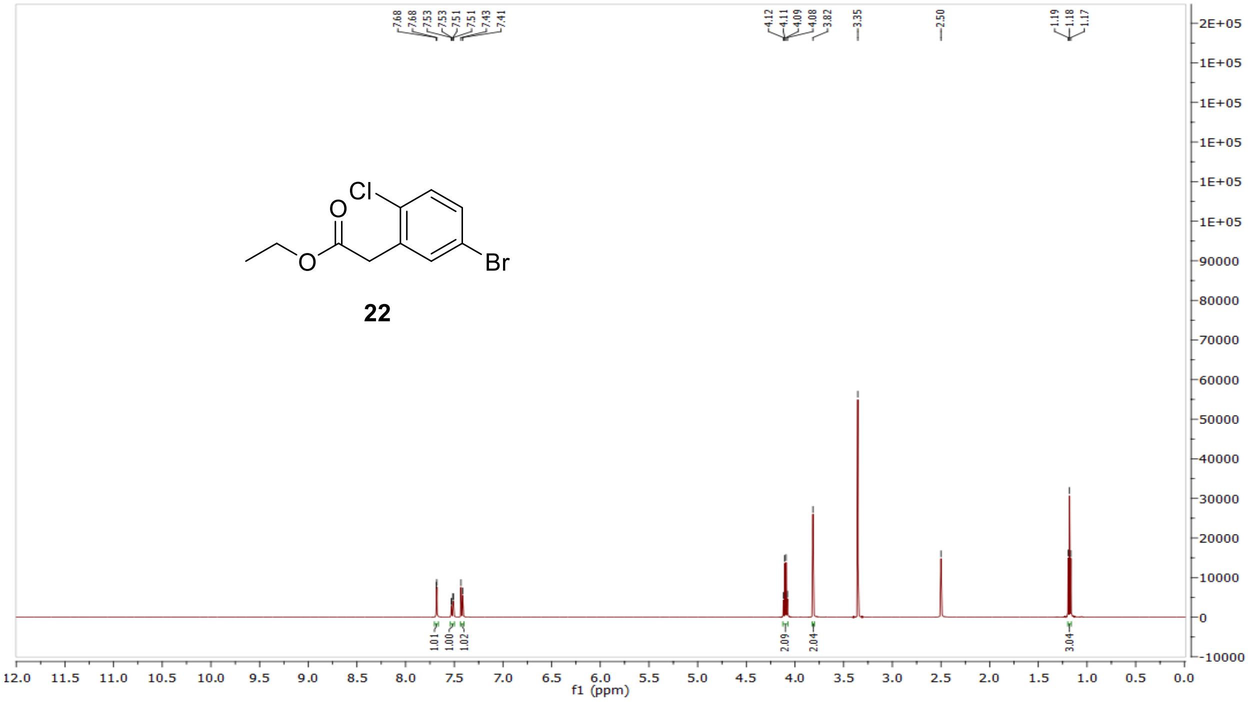

$^{13}$C NMR compound **22:**

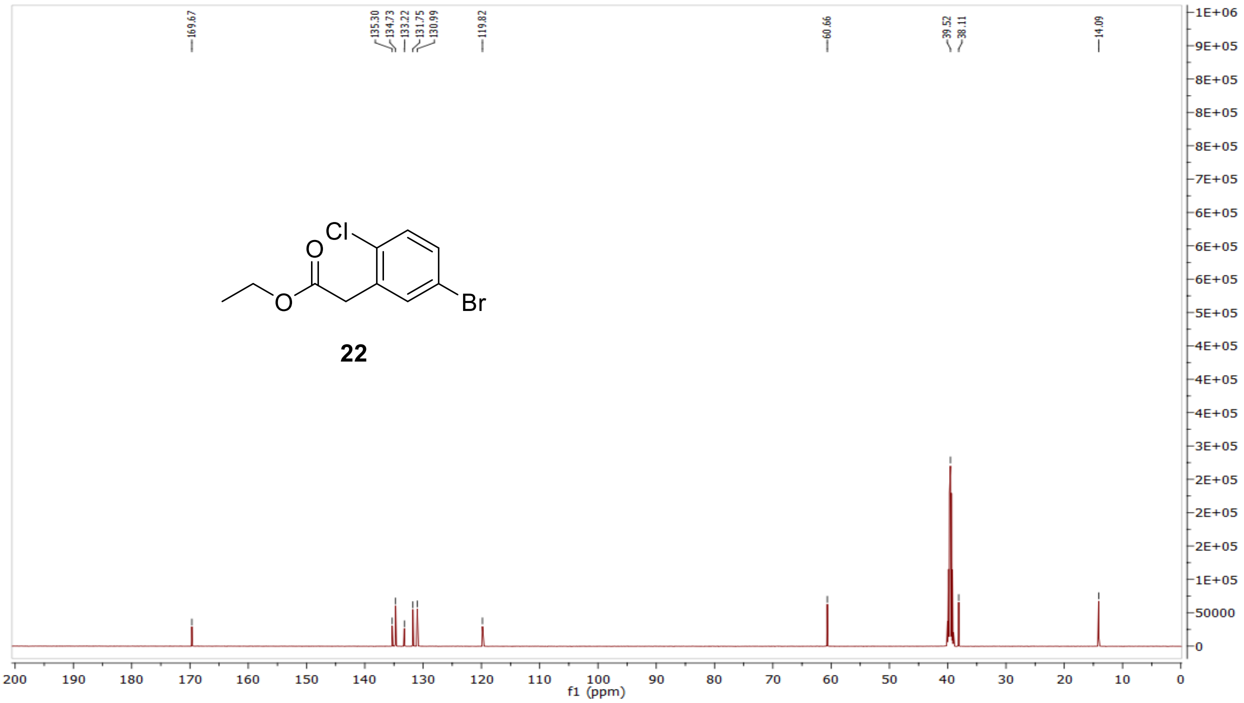


MS (ESI+) compound **22**:

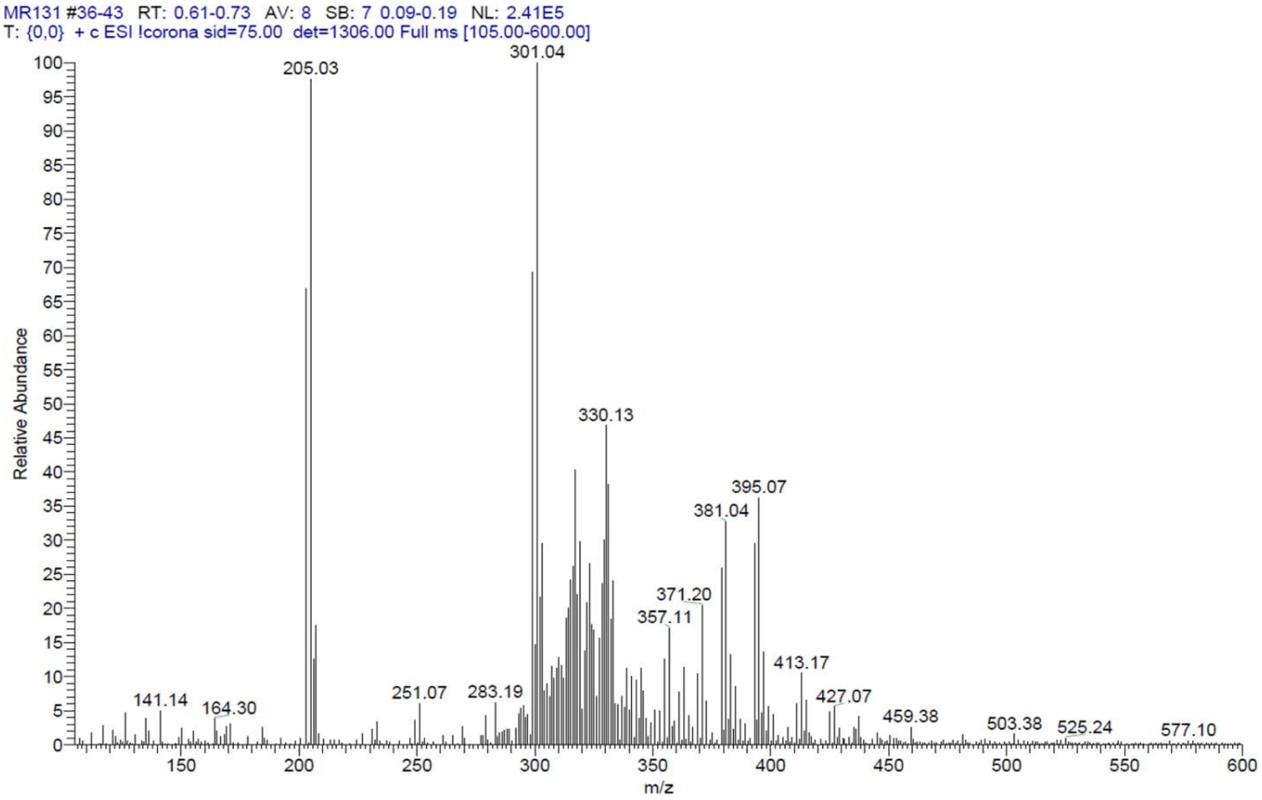

$^{1}H$ NMR compound **23**:

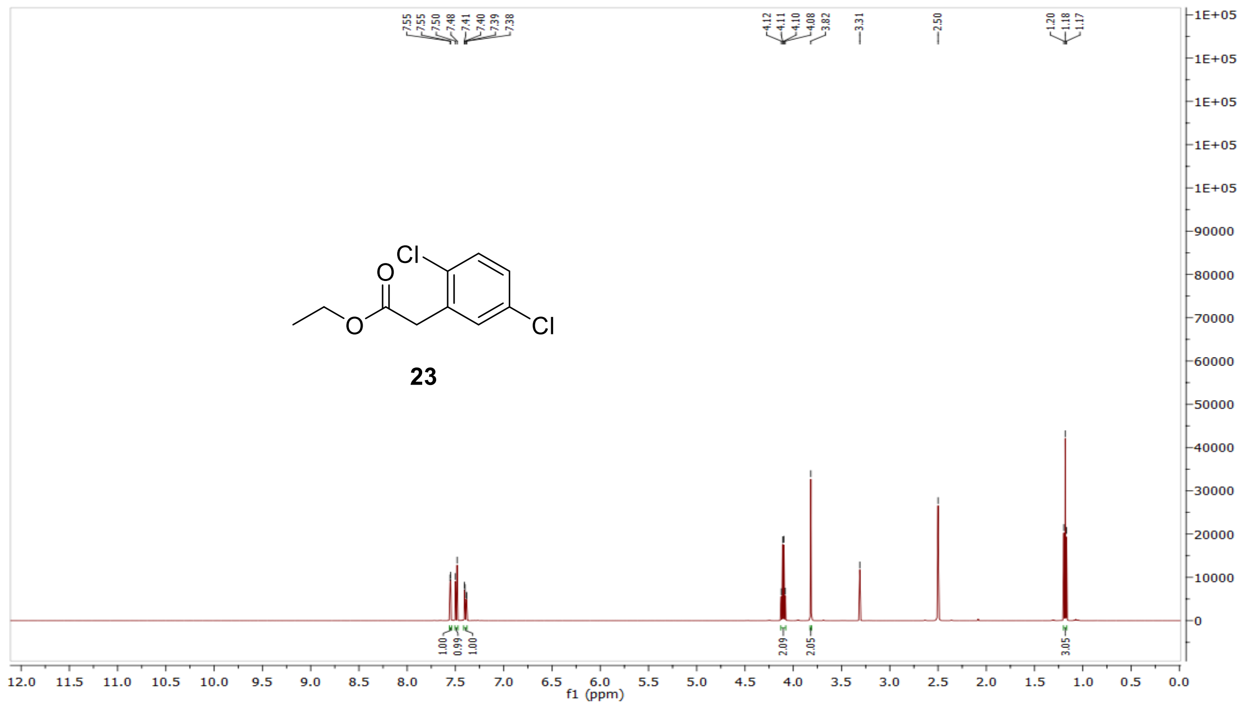


$^{13}C$ NMR compound **23:**

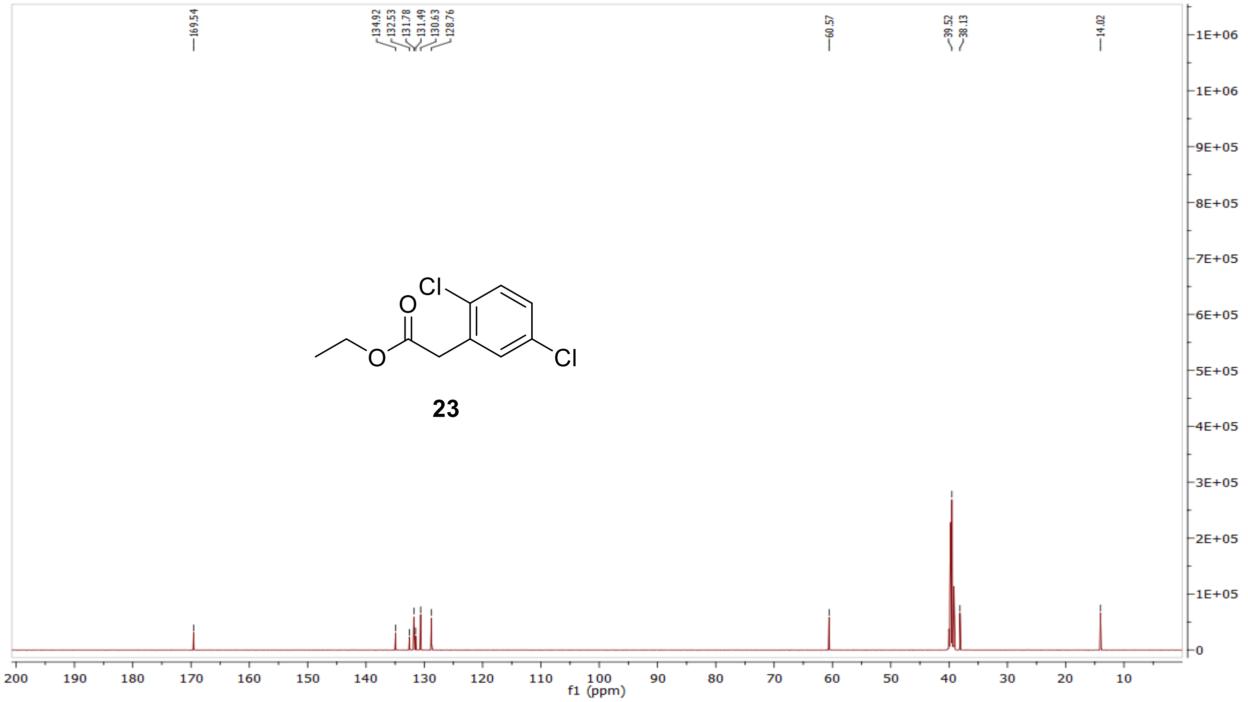

MS (ESI+) compound **23**:

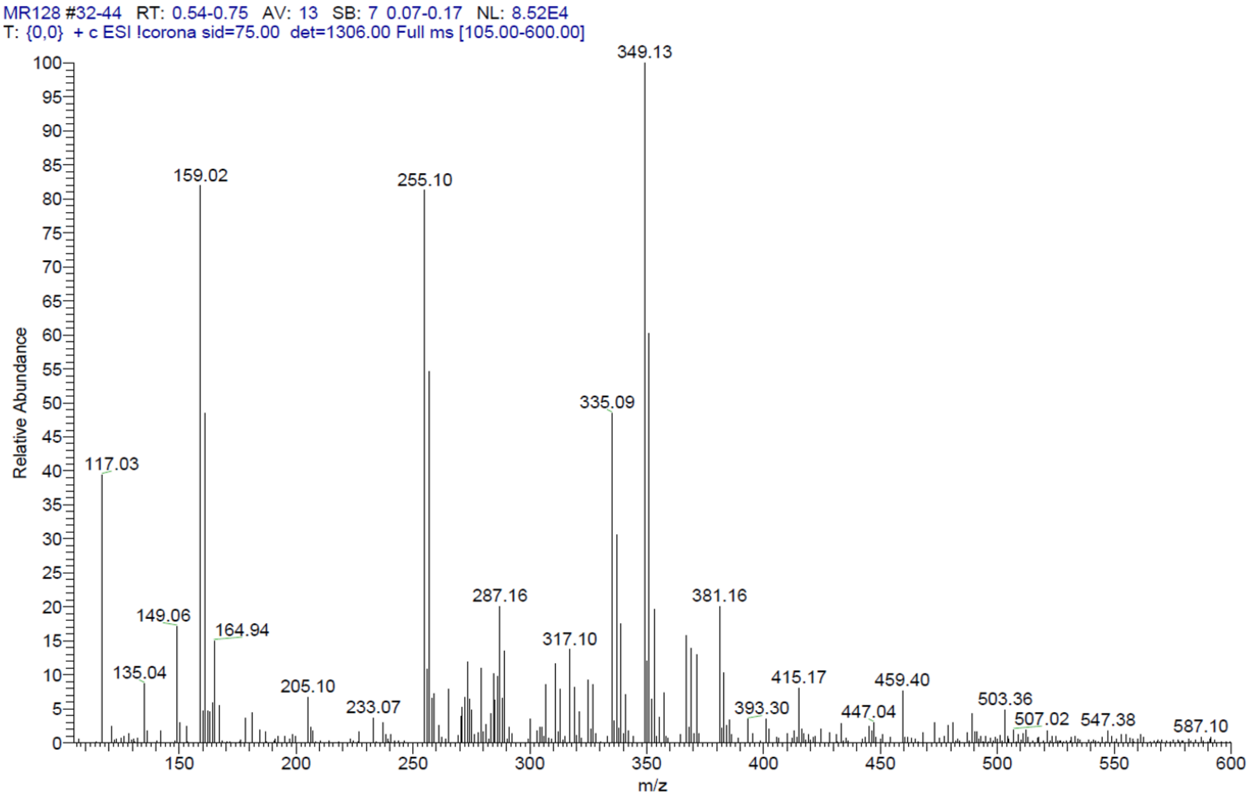


[1]H NMR compound **24**:

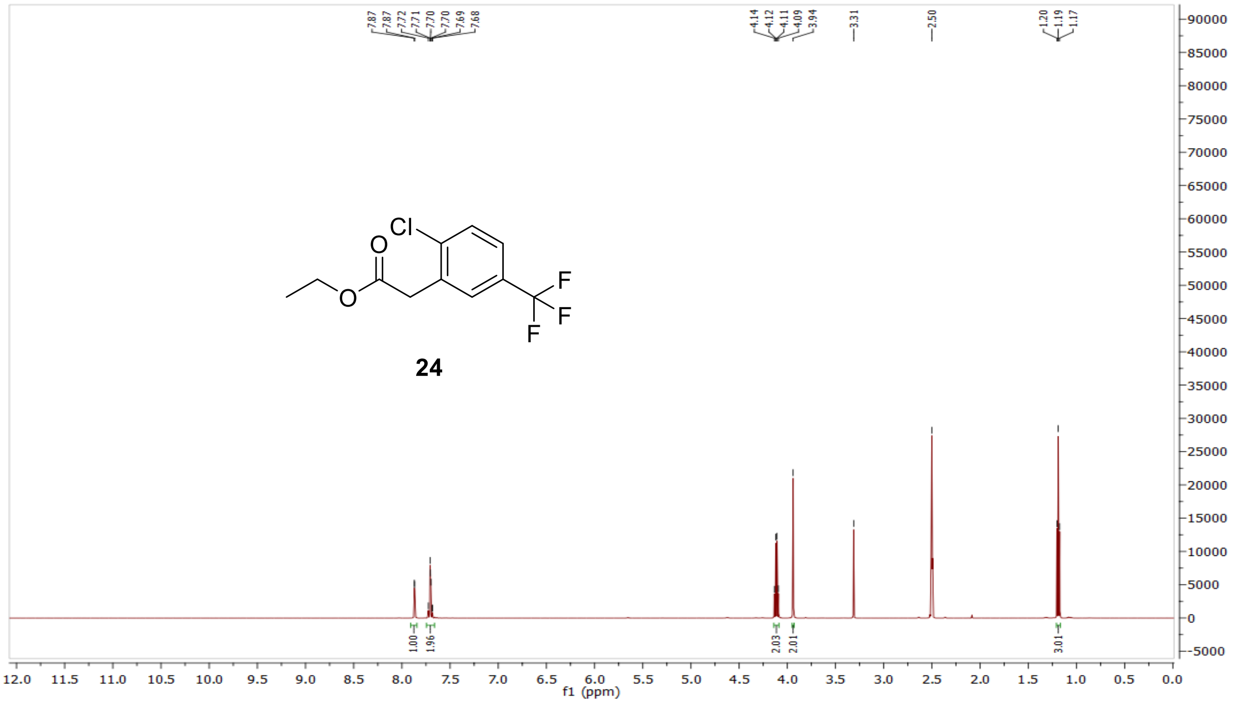

$^{13}$C NMR compound **24:**

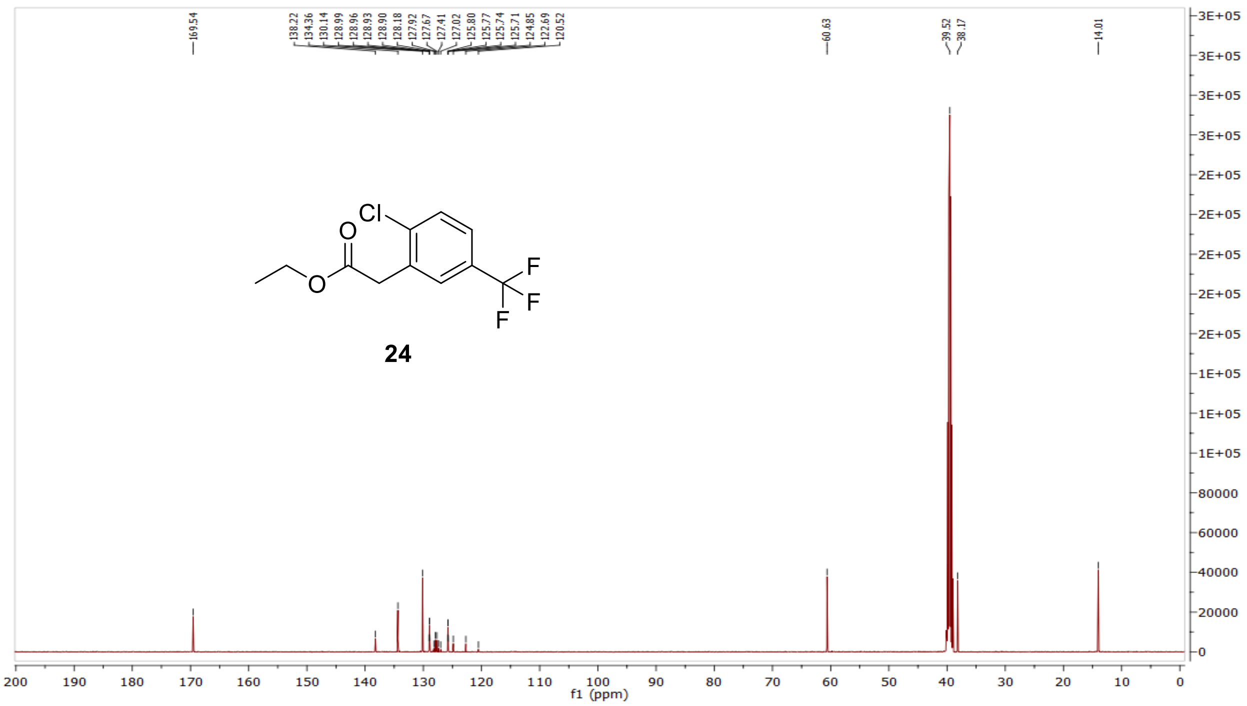

MS (ESI+) compound **24**:

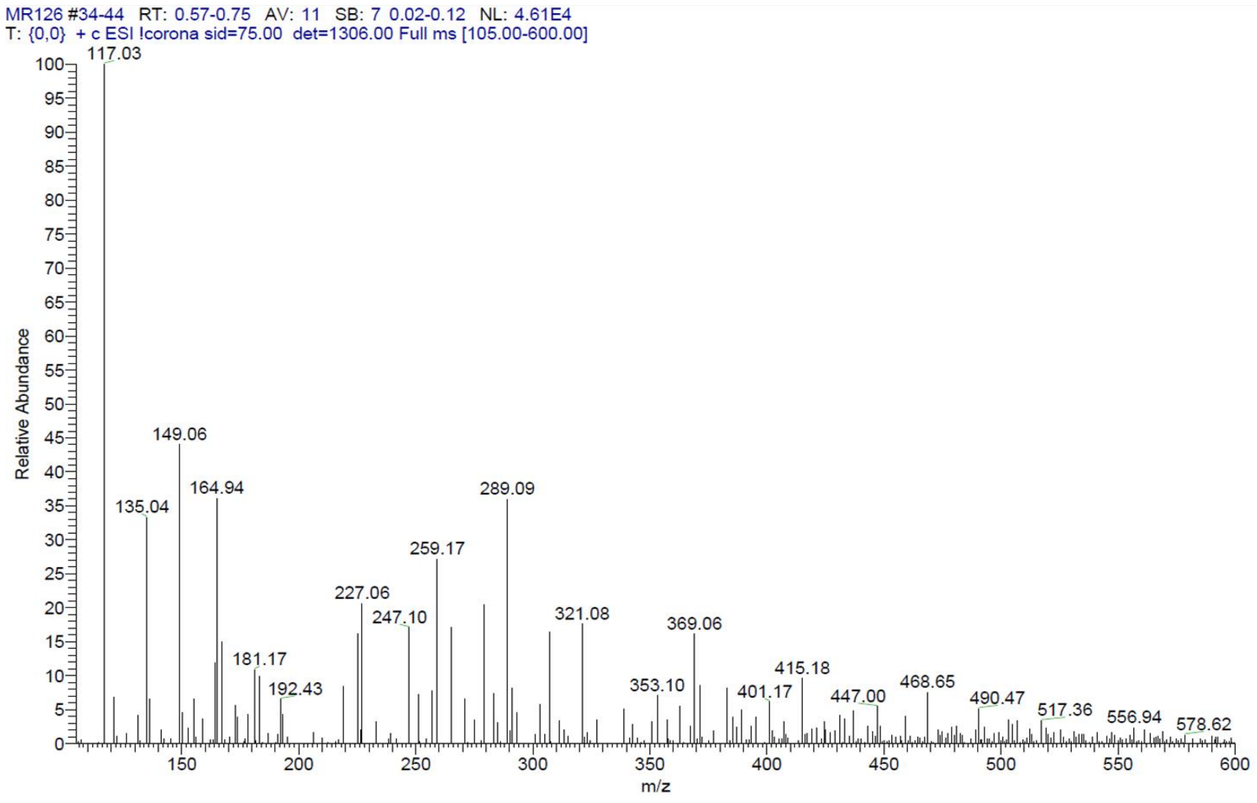

<sup></sup>[1]H NMR compound **25**:

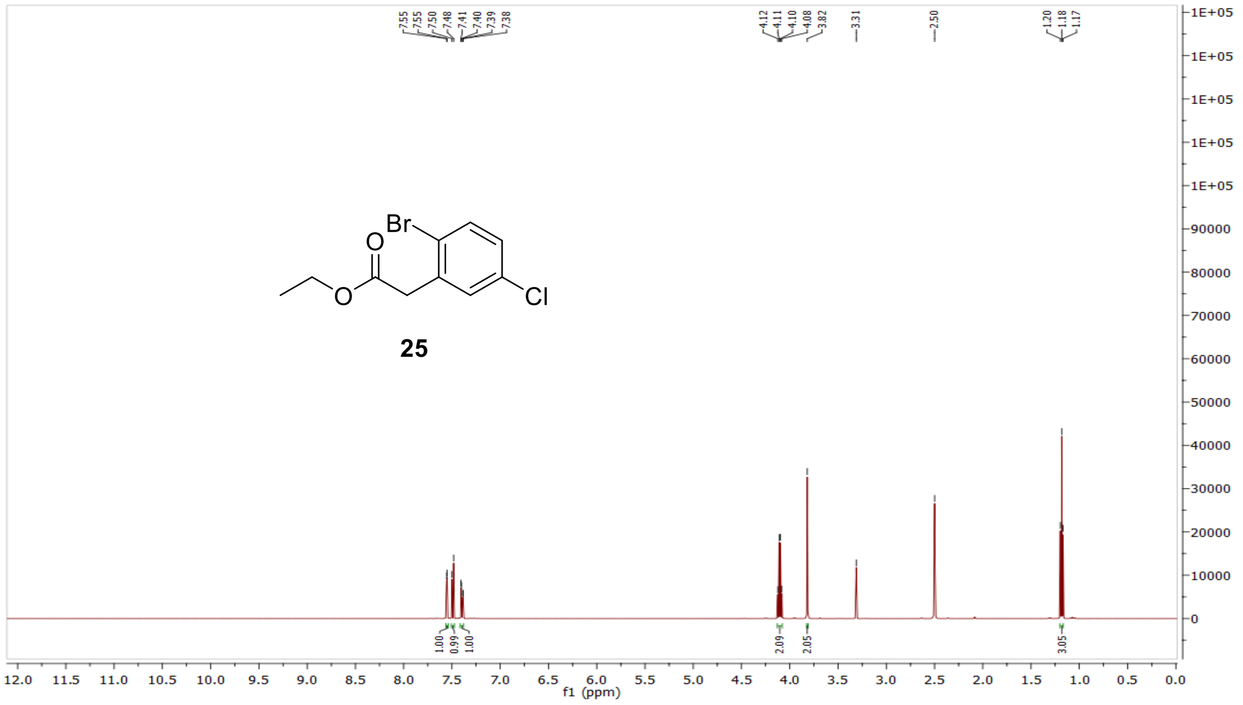

[13]C NMR compound **25:**

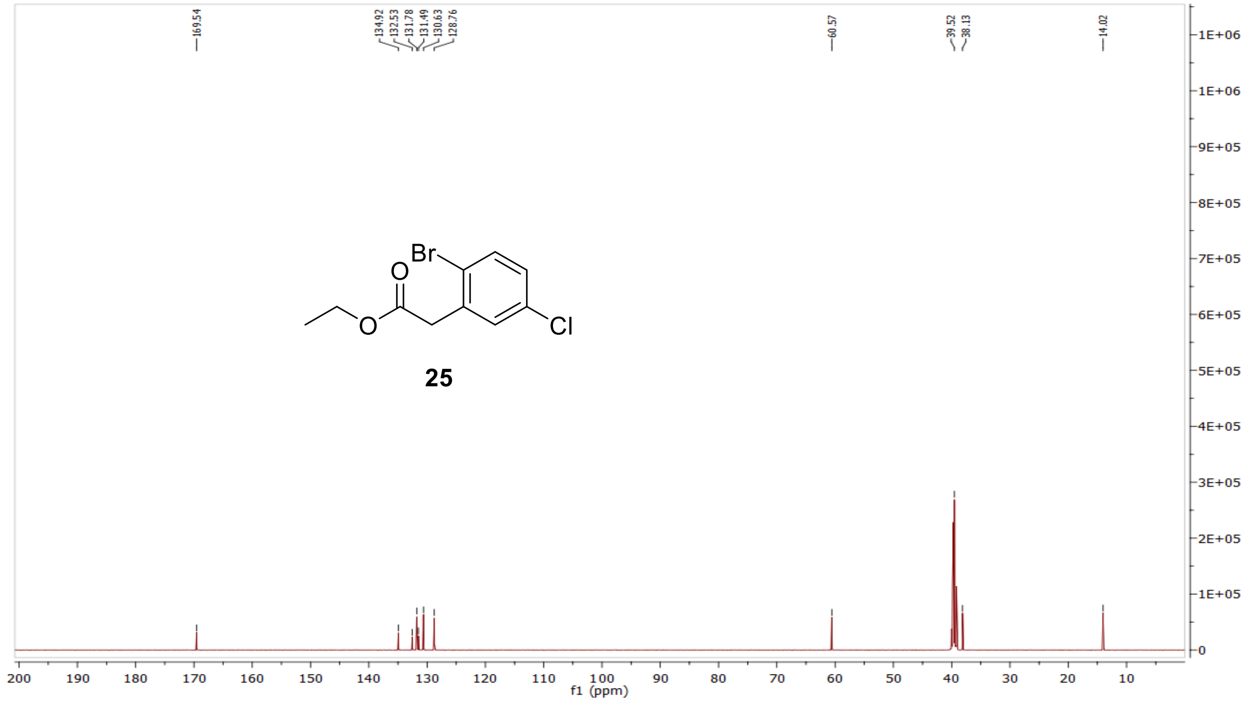

MS (ESI+) compound **25**:

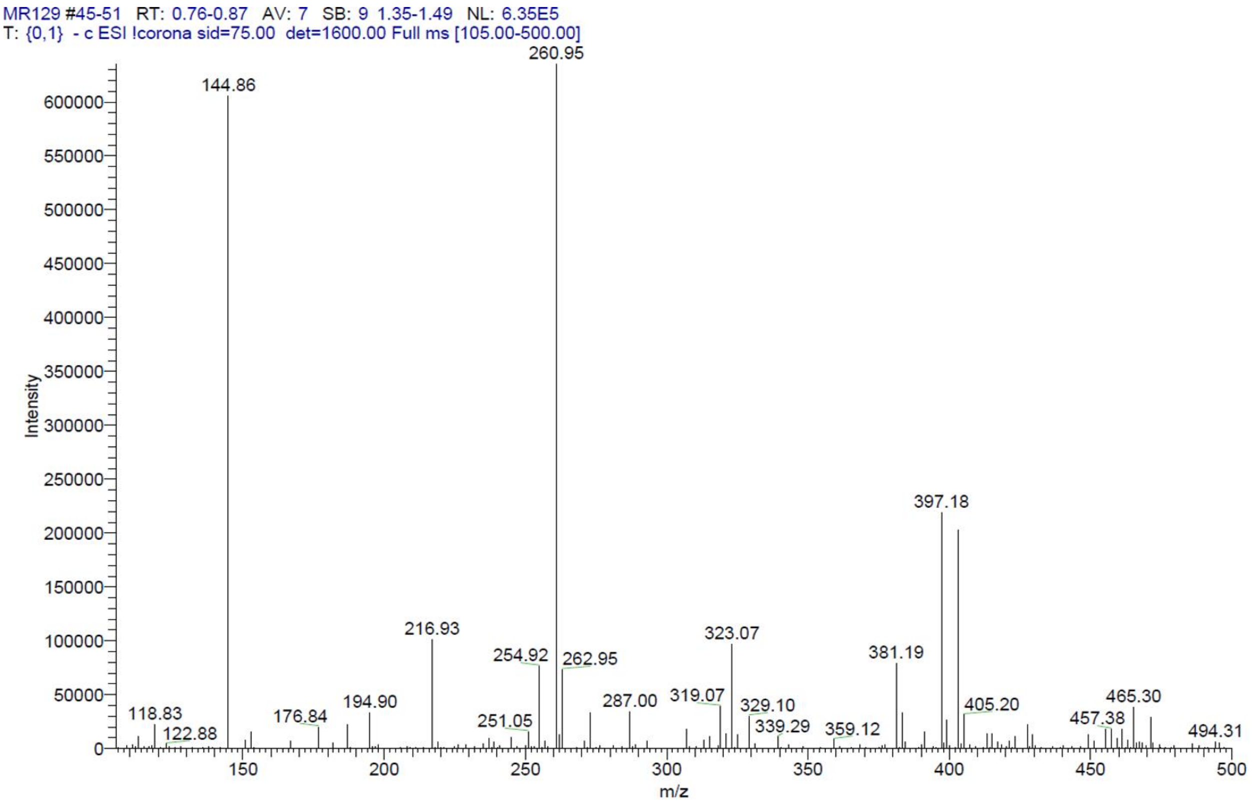


[1]H NMR compound **26**:

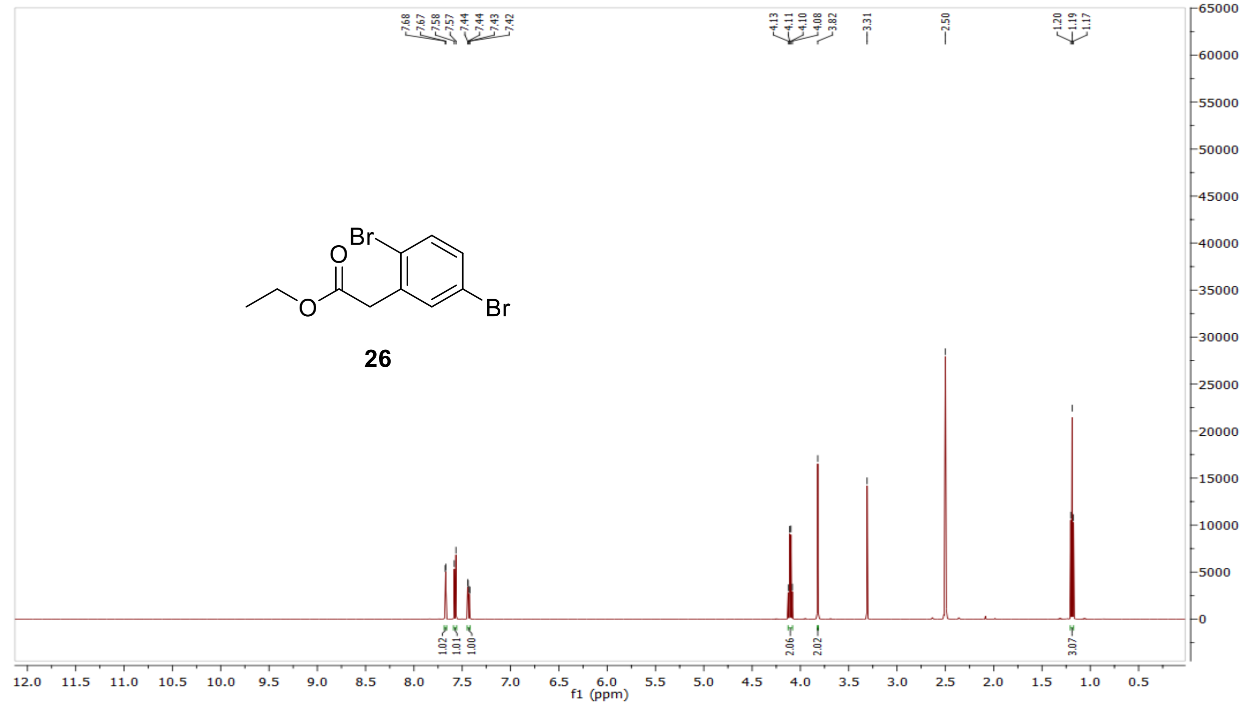

[13]C NMR compound **26:**

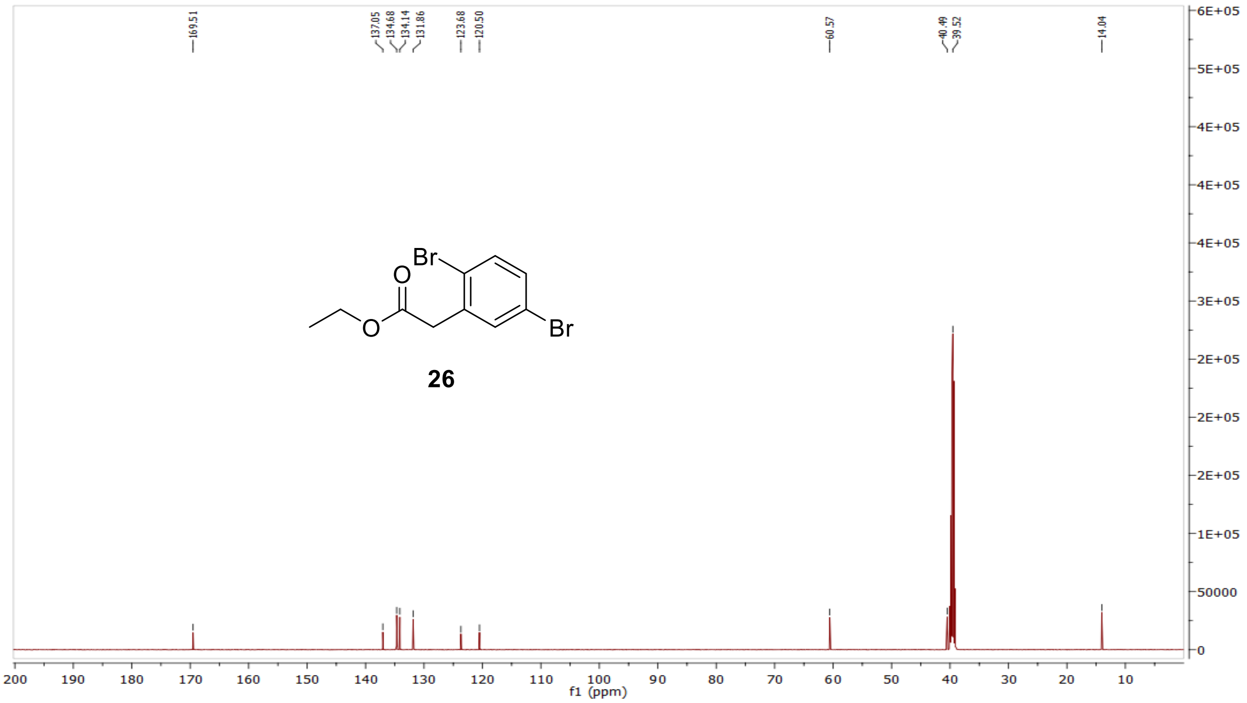


MS (ESI+) compound **26**:

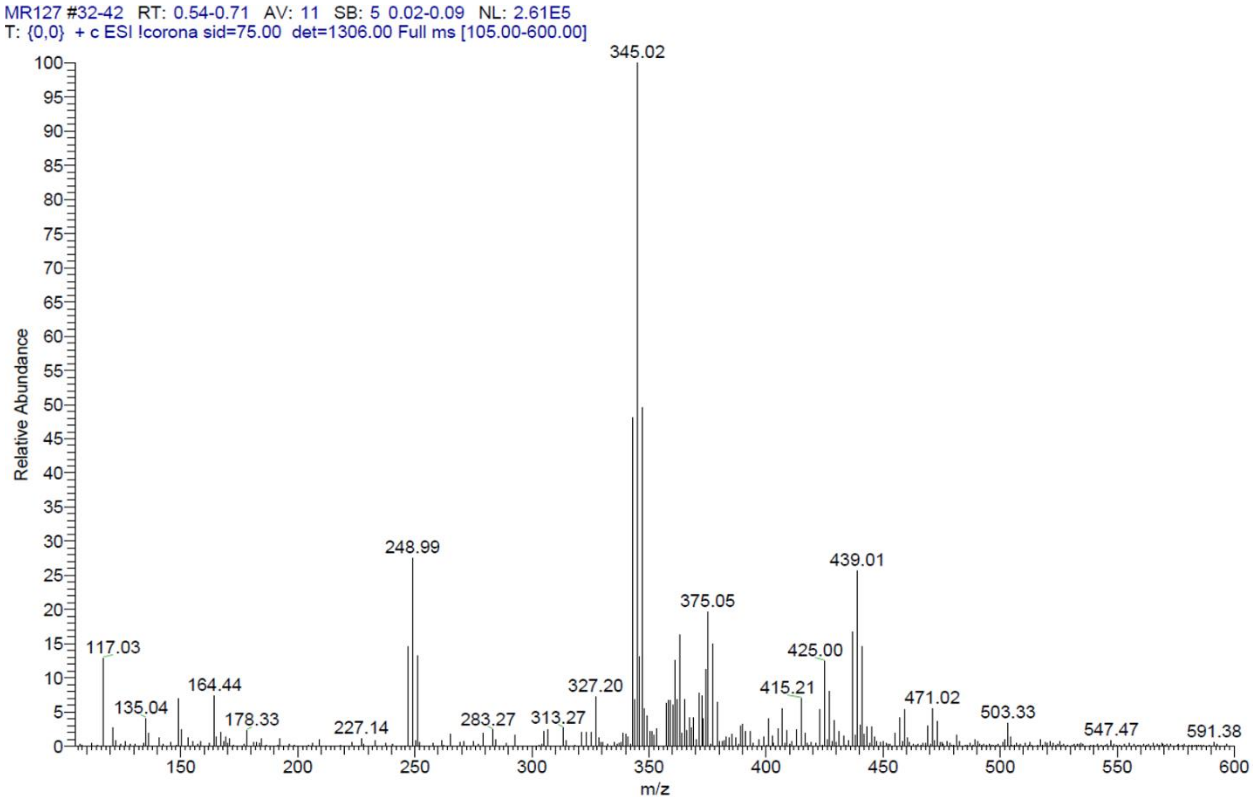

$^1$H NMR compound **27**:

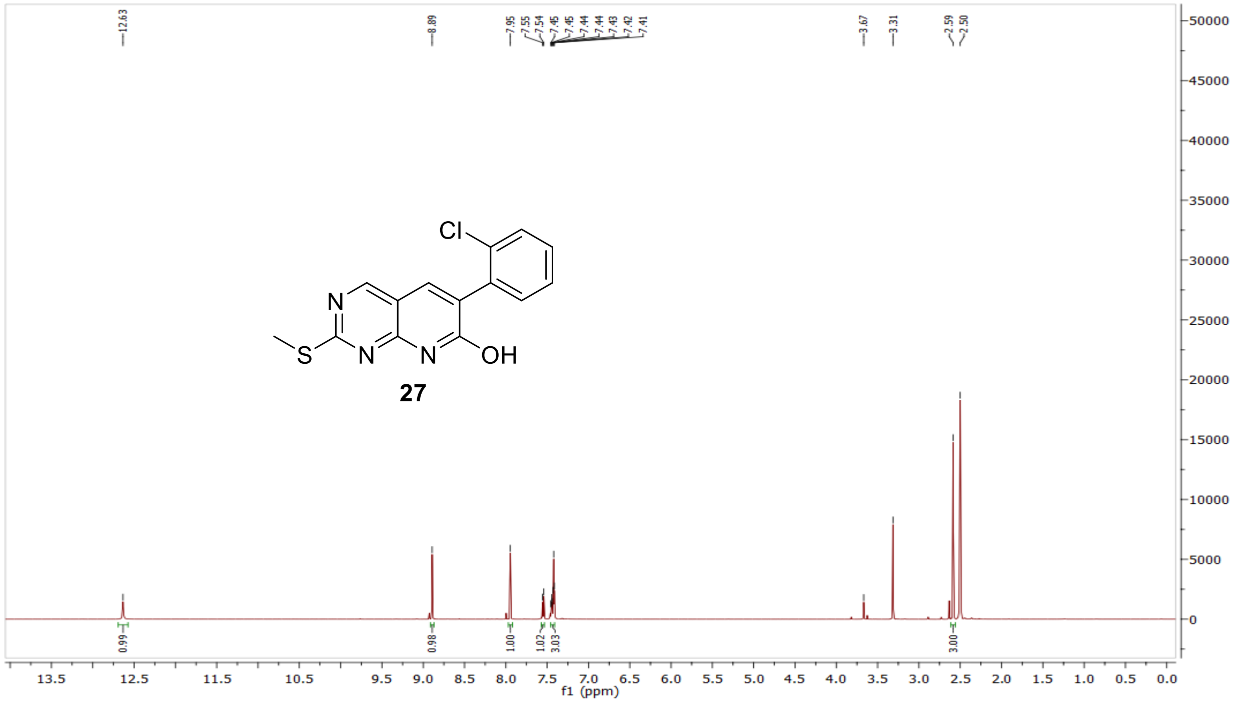


$^{13}$C NMR compound **27:**

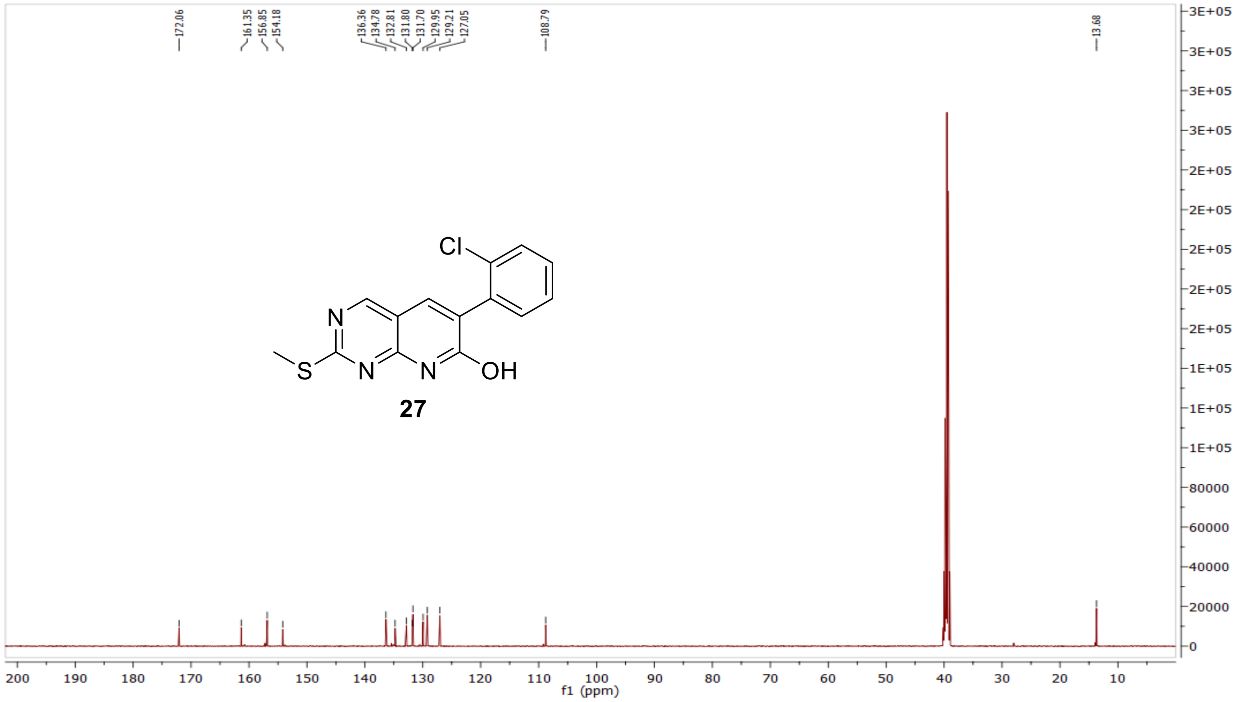

MS (ESI+) compound **27**:

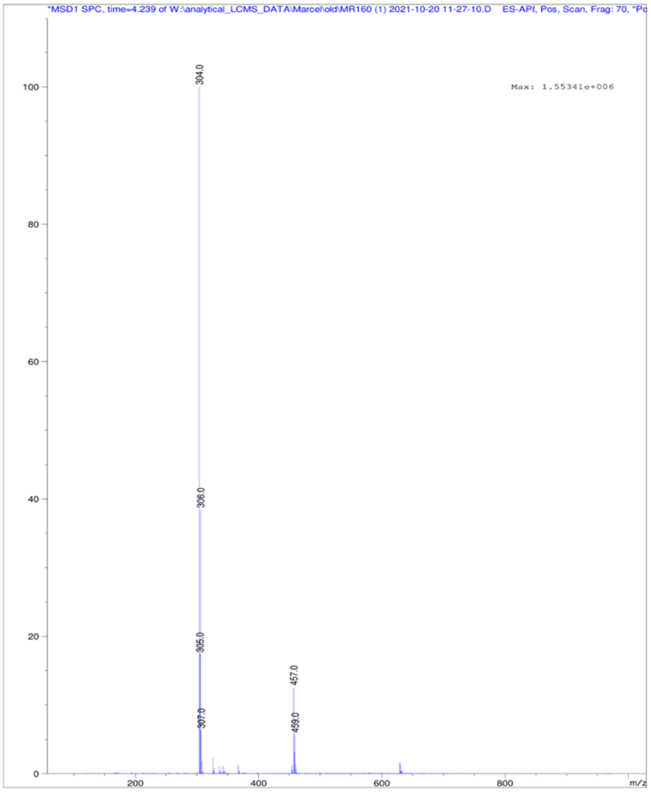


$^1$H NMR compound **28**:

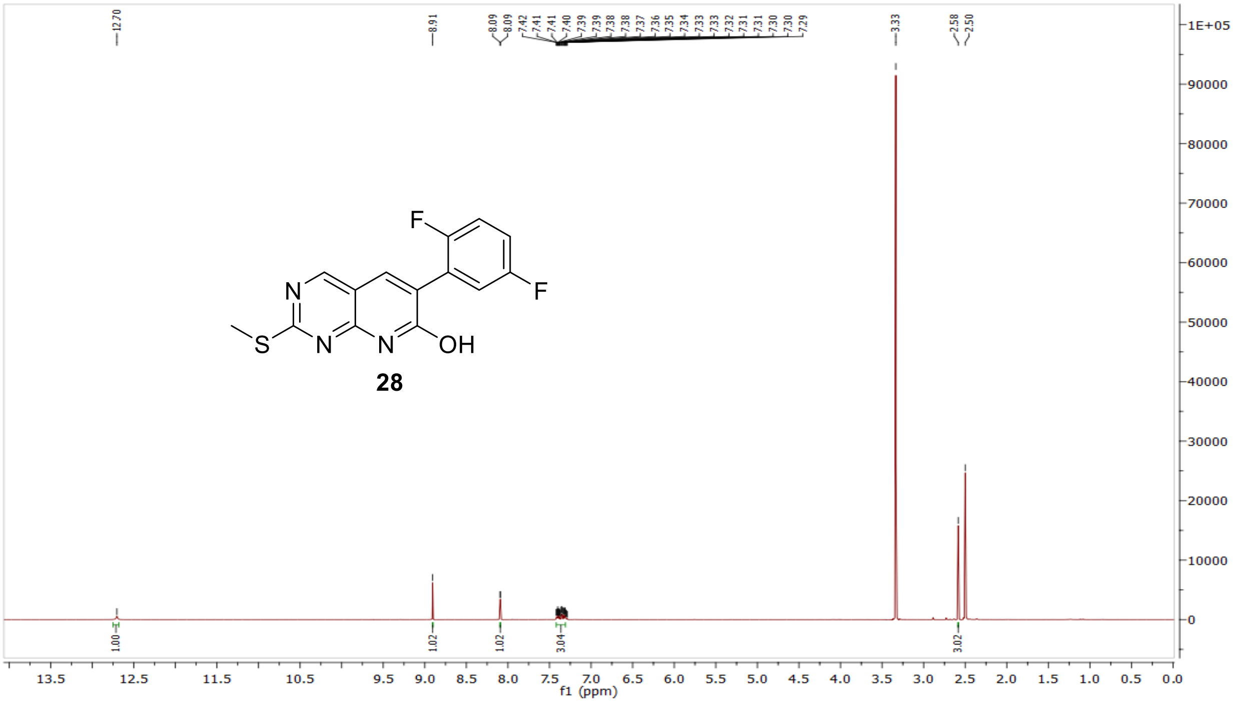

$^{13}$C NMR compound **28:**

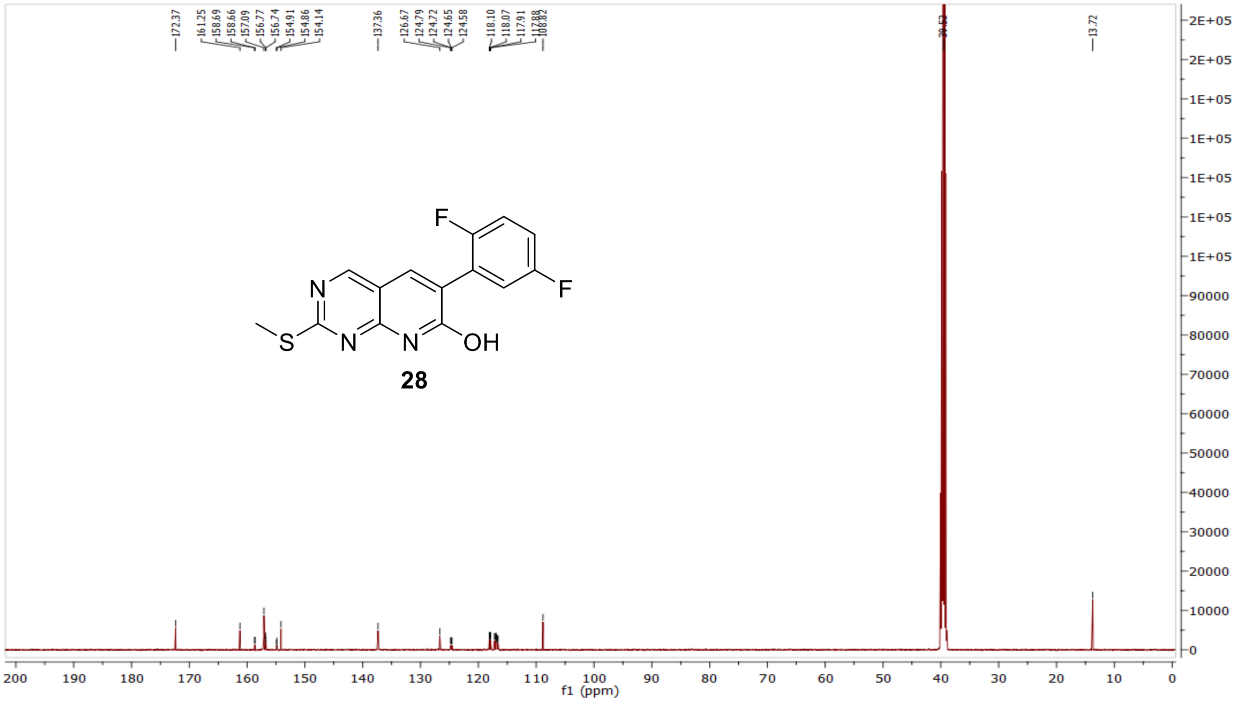


MS (ESI+) compound **28**:

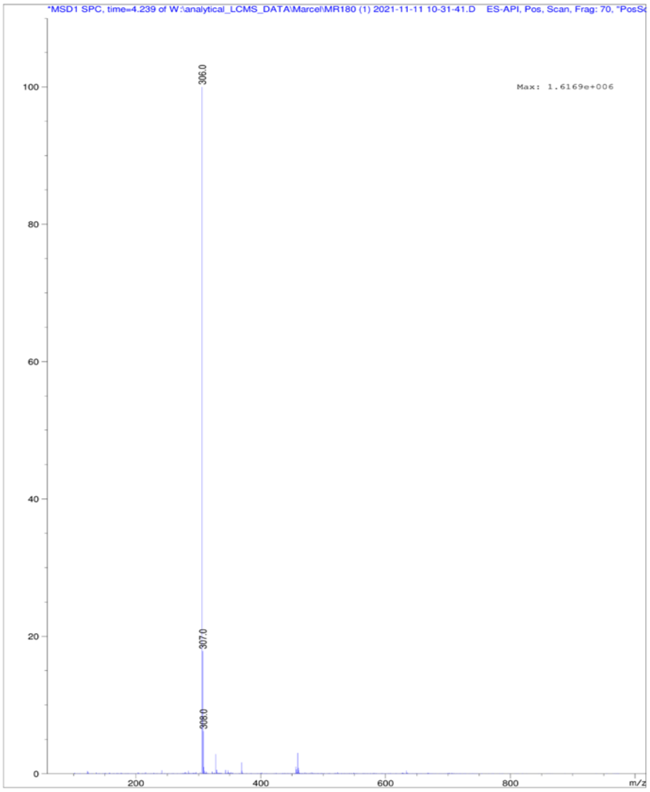

[1]H NMR compound **29**:

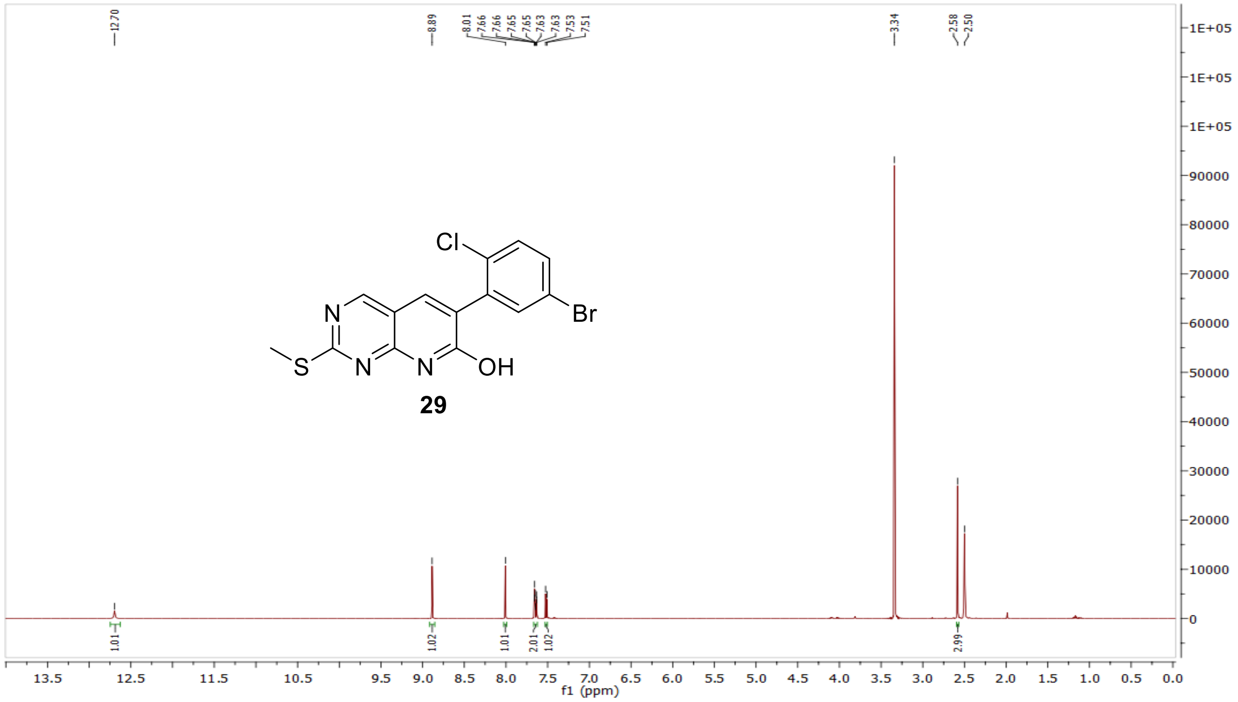

[13]C NMR compound **29:**

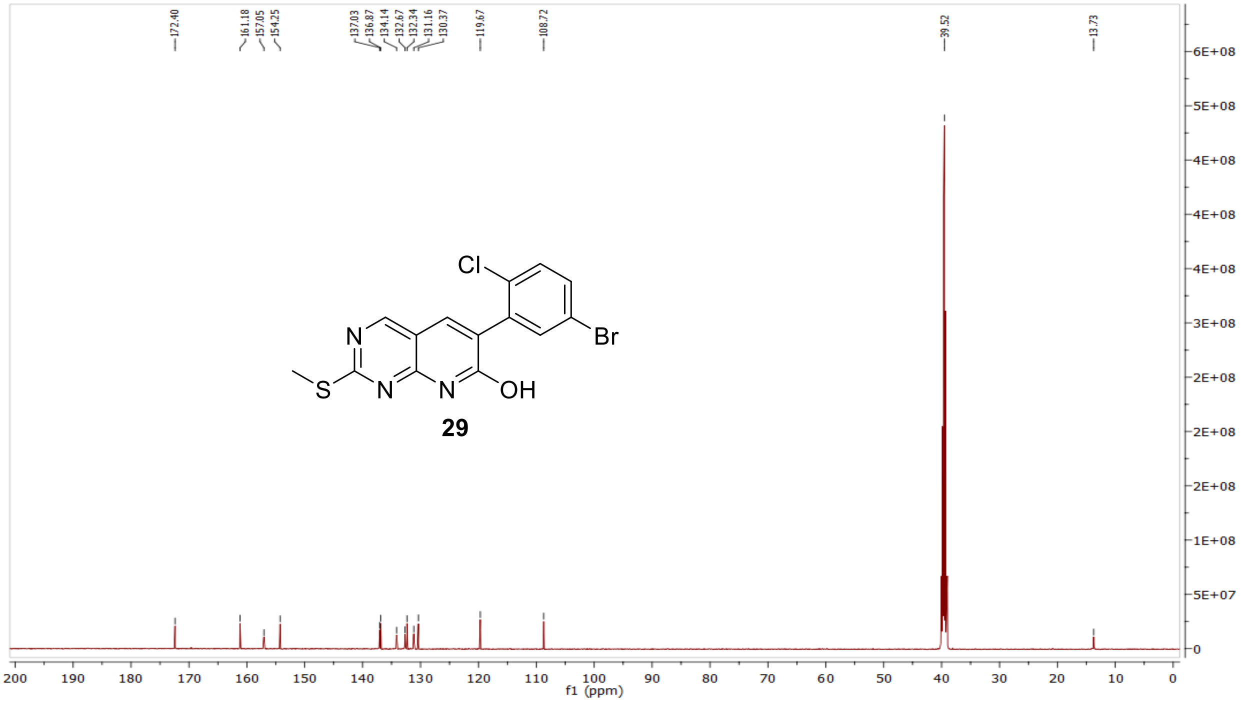

MS (ESI+) compound **29**:

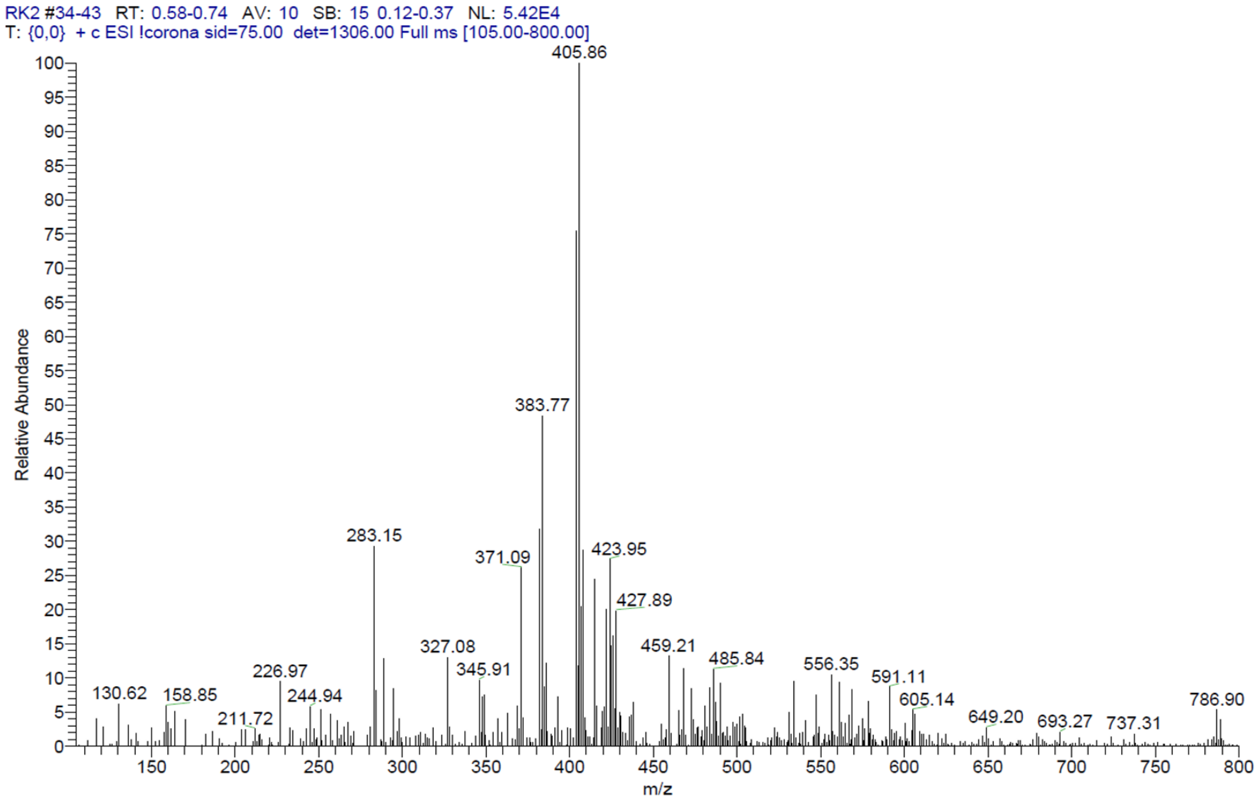


$^1$H NMR compound **30**:

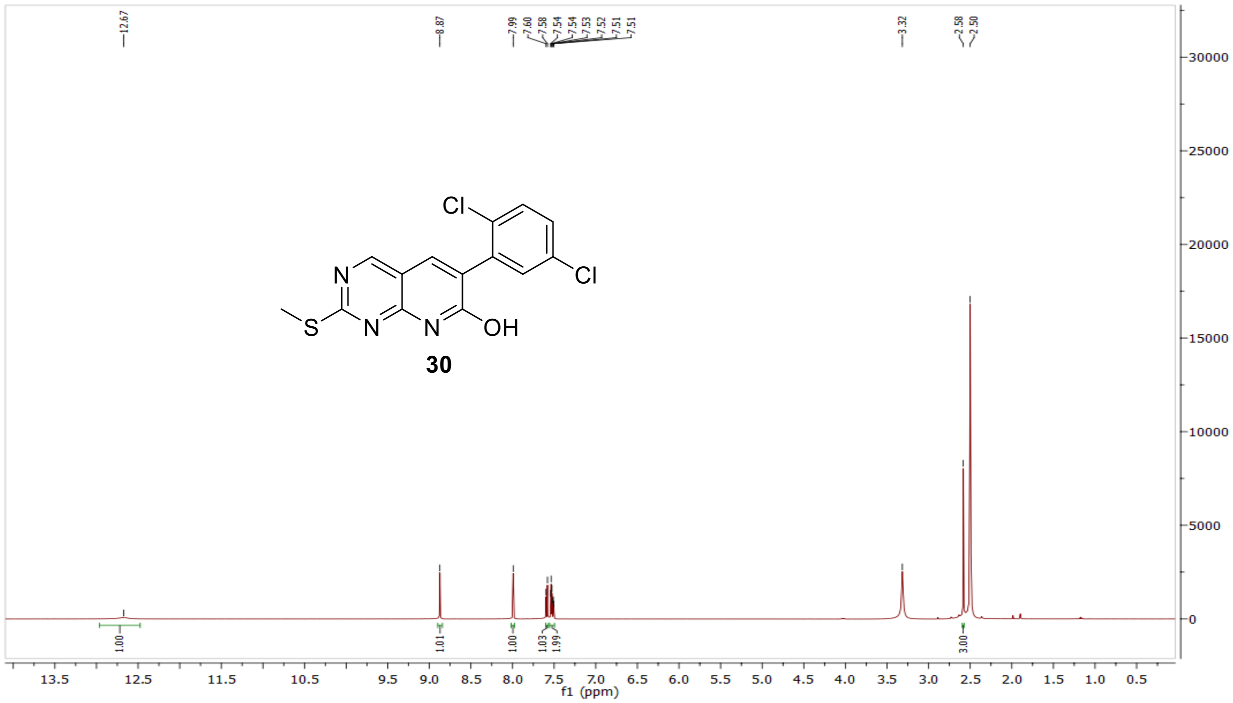

$^{13}$C NMR compound **30:**

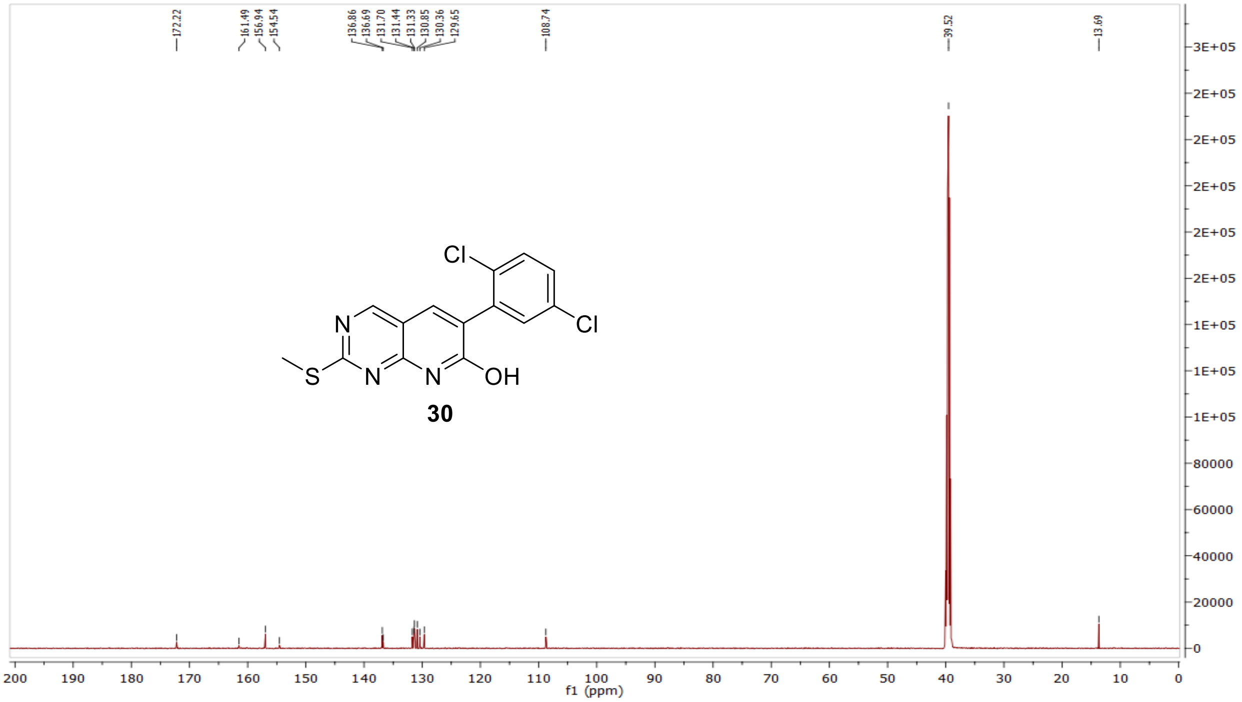


MS (ESI+) compound **30**:

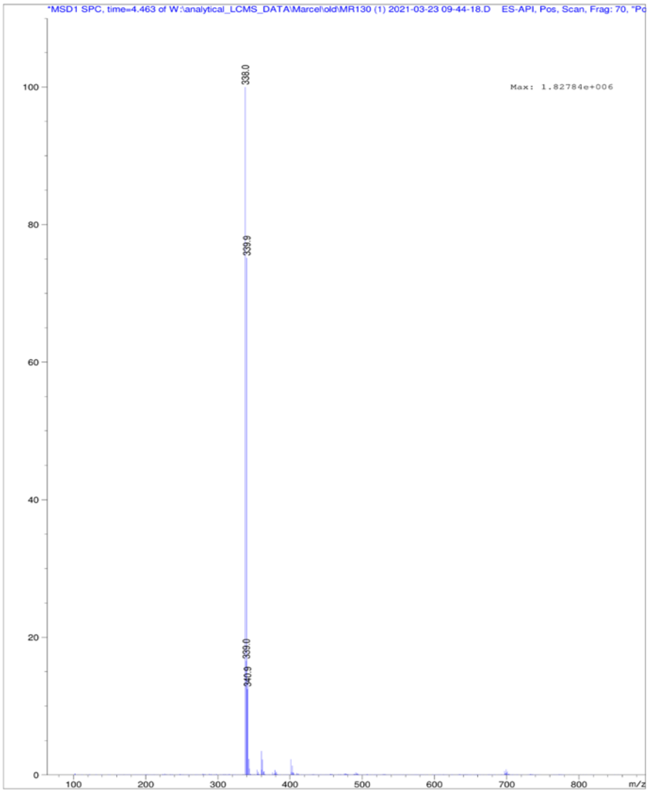

$^{1}$H NMR compound **31**:

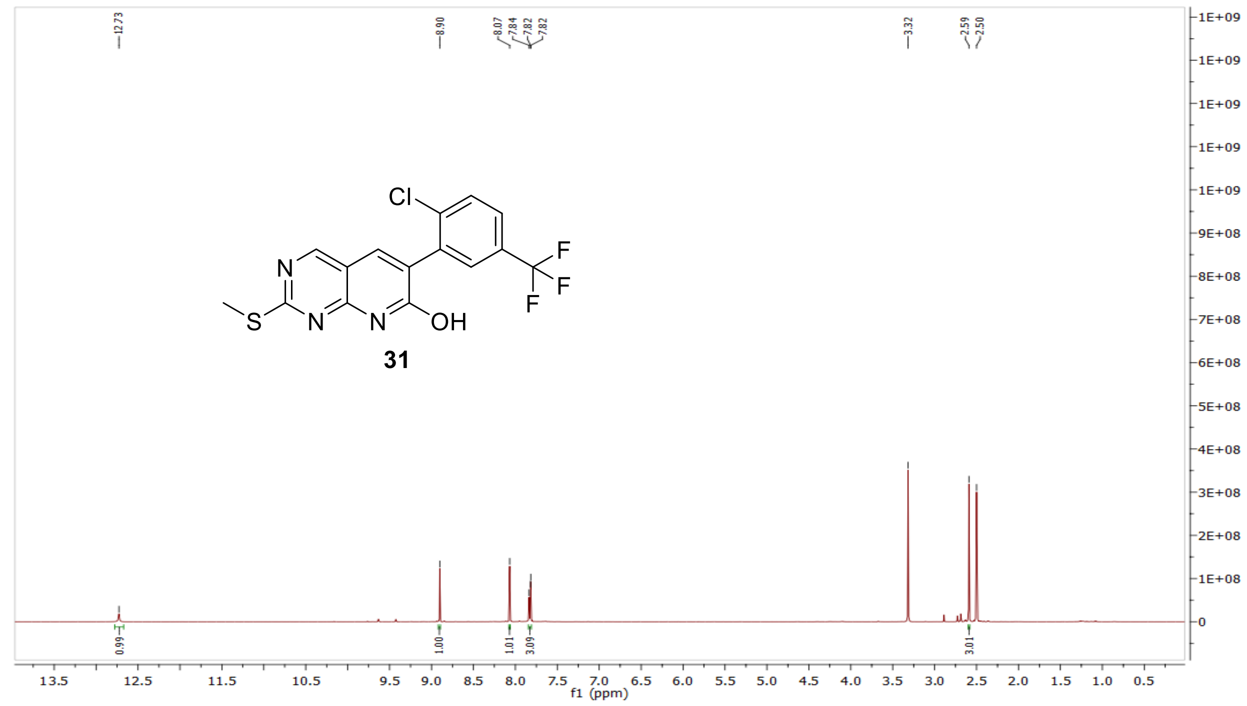

$^{13}$C NMR compound **31:**

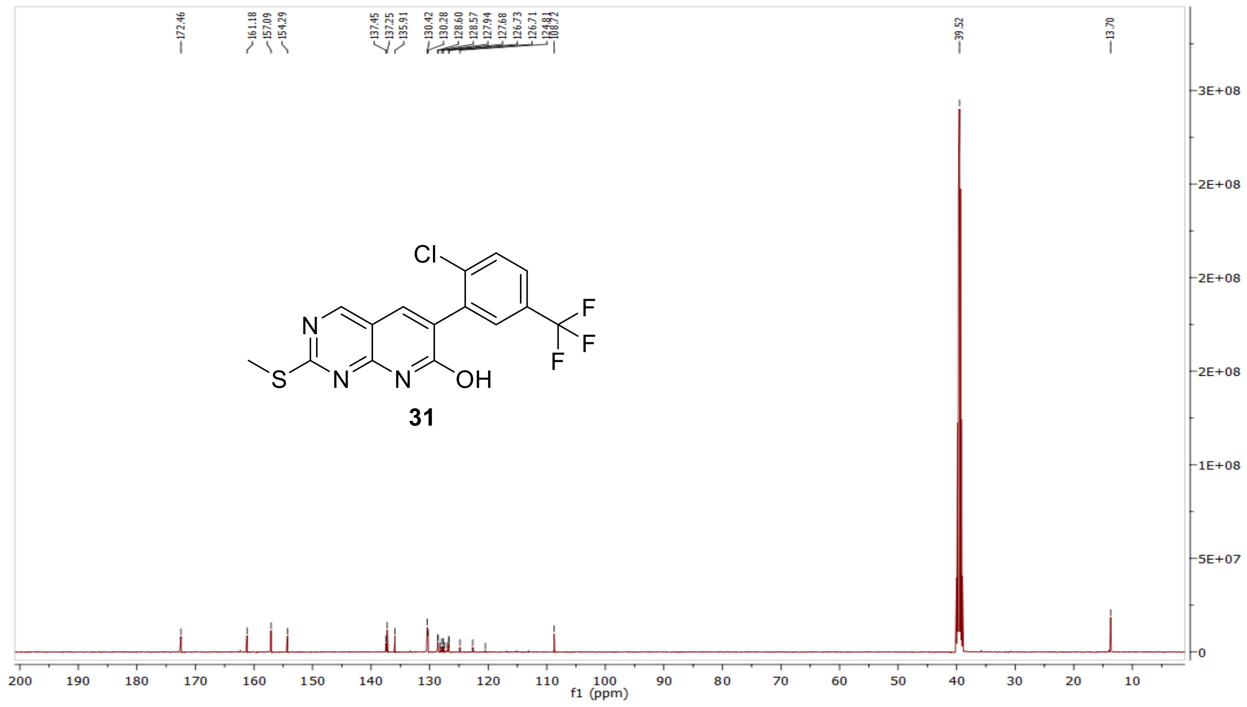

MS (ESI+) compound **31**:

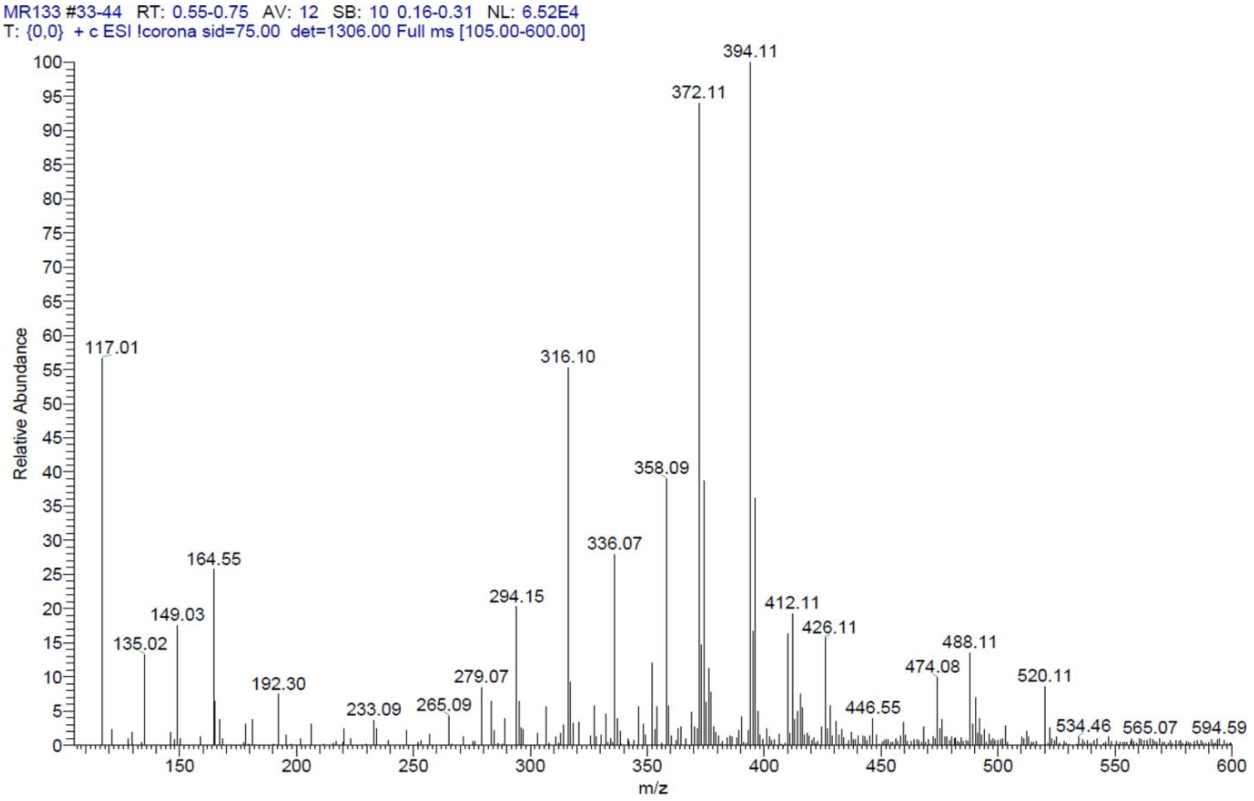


[1]H NMR compound **32**:

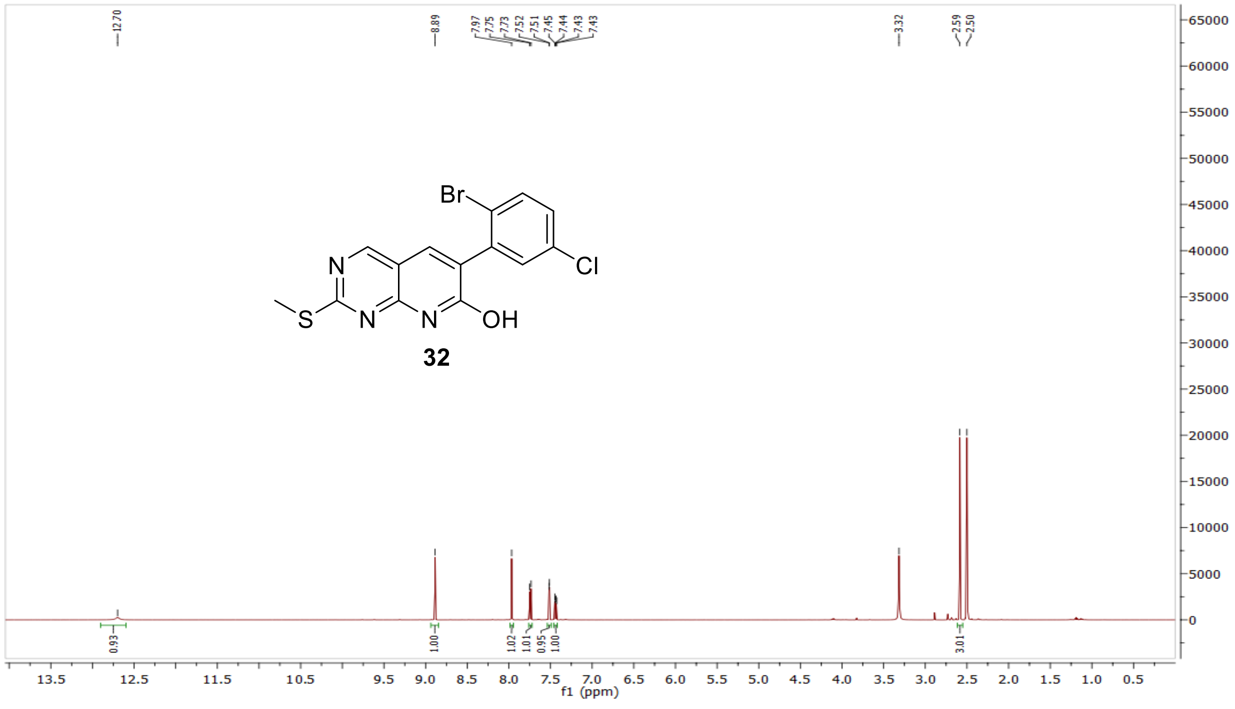

$^{13}$C NMR compound **32:**

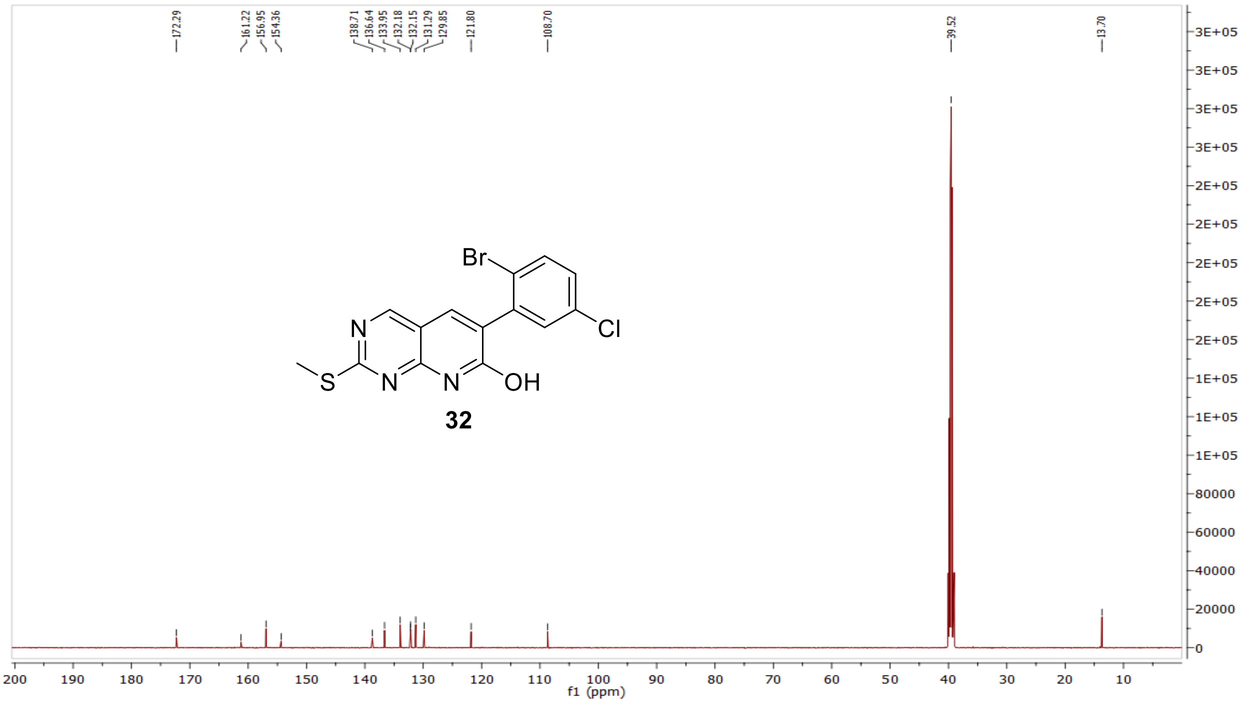


MS (ESI+) compound **32**:

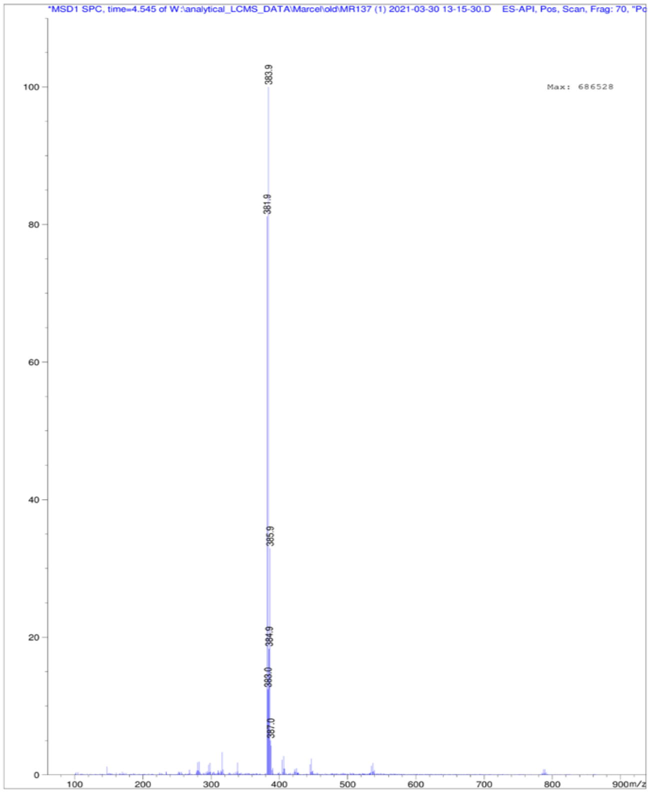

[1]H NMR compound **33**:

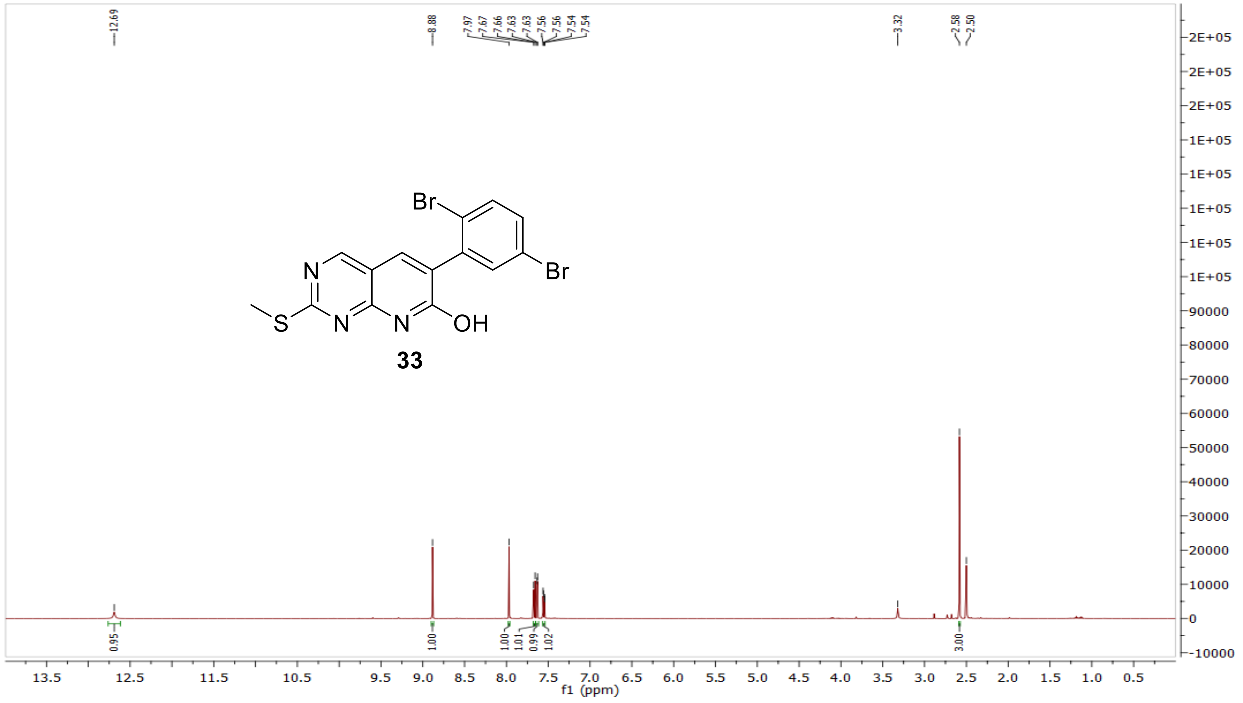


[13]C NMR compound **33:**

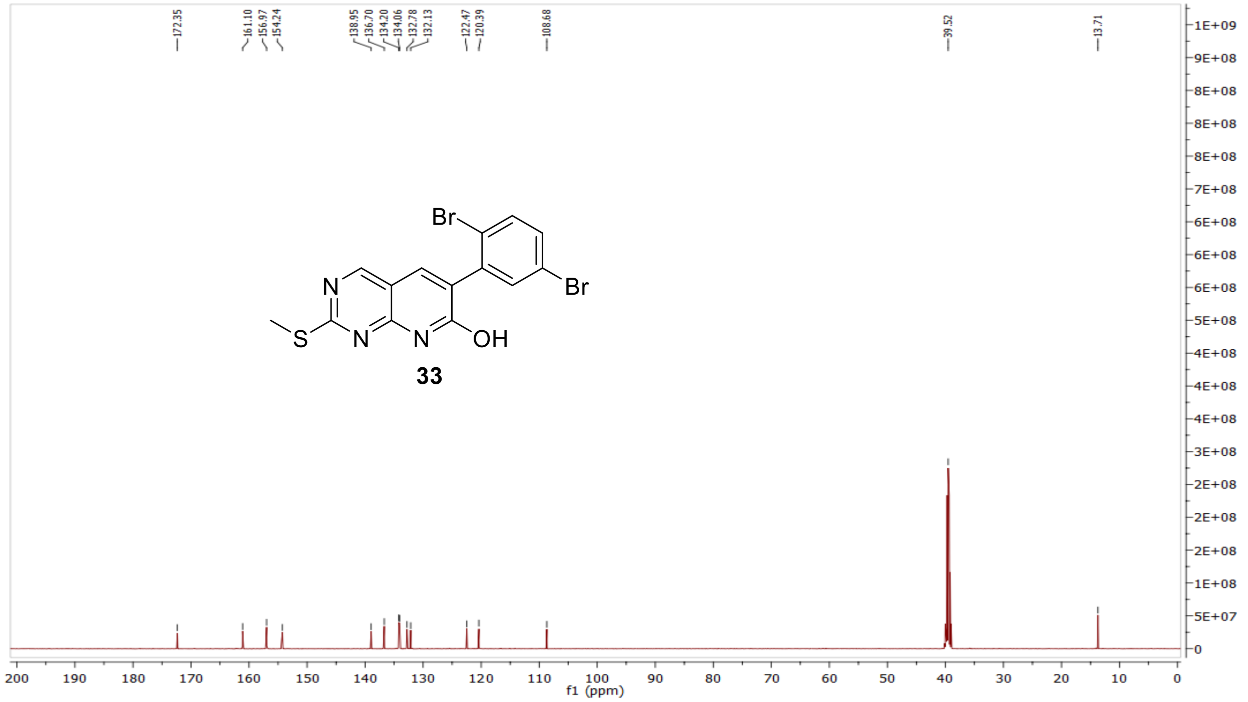

MS (ESI+) compound **33**:

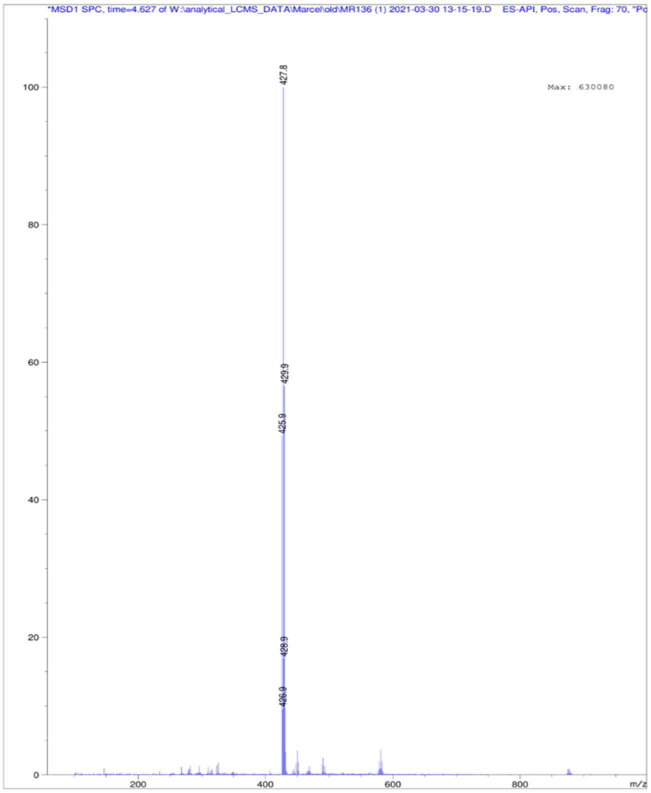


[1]H NMR compound **34**:

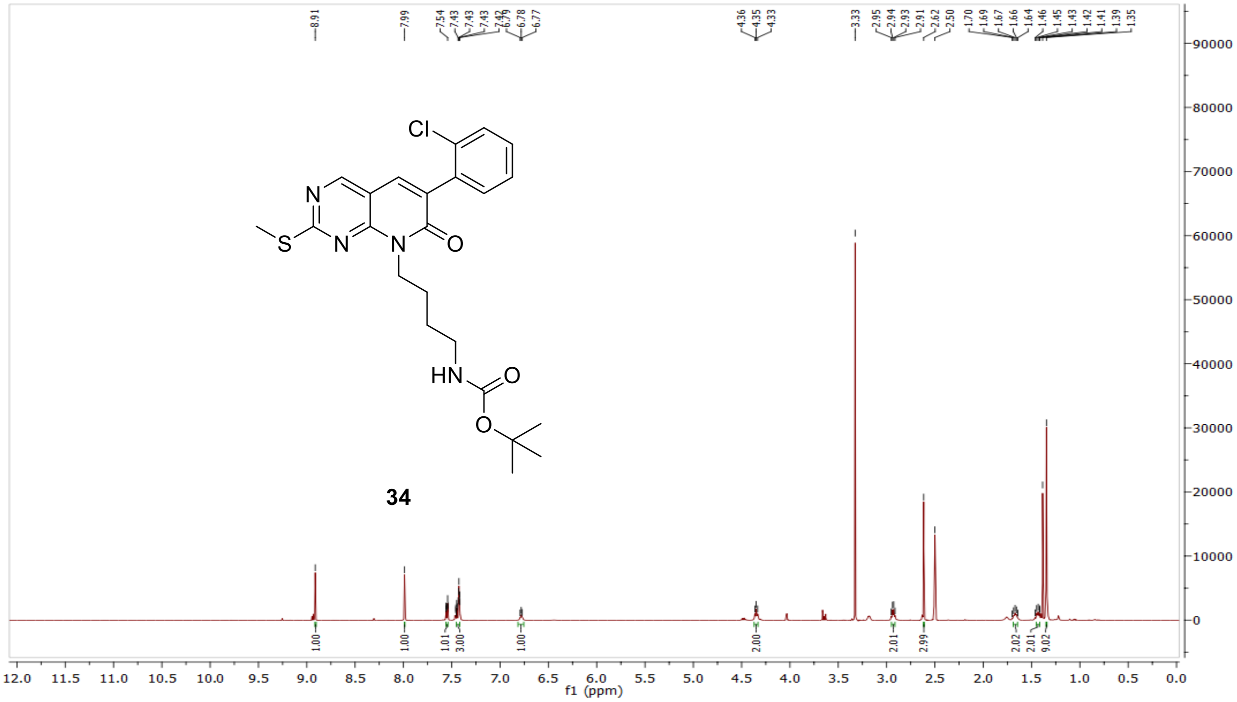

$^{13}C$ NMR compound **34:**

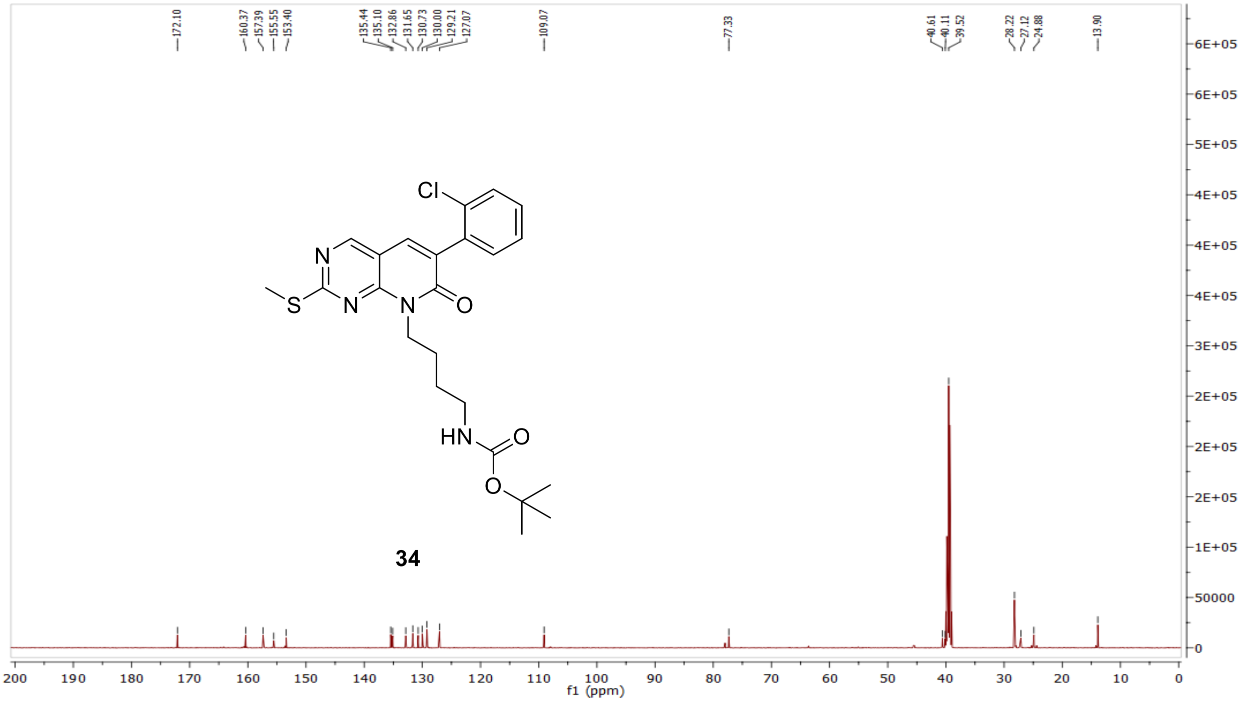


MS (ESI+) compound **34**:

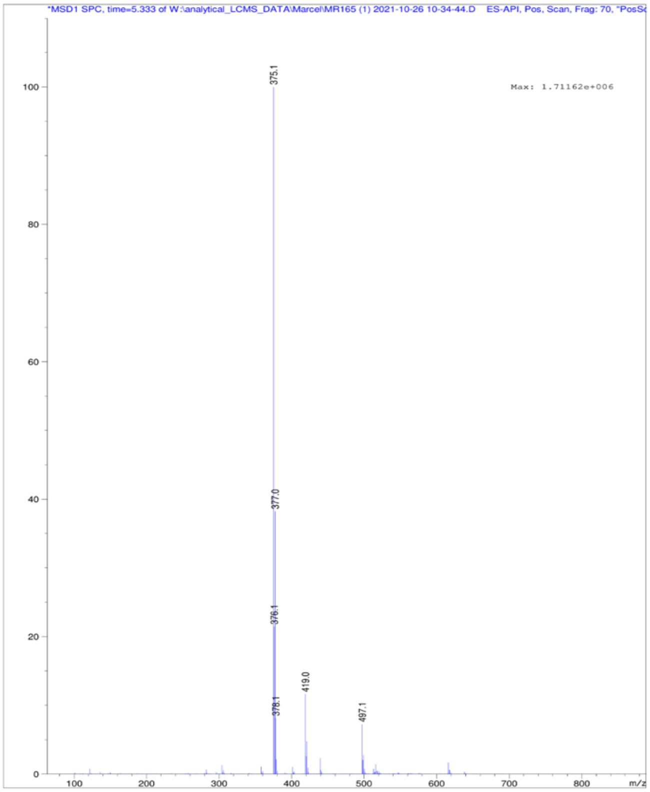

[1]H NMR compound **35**:

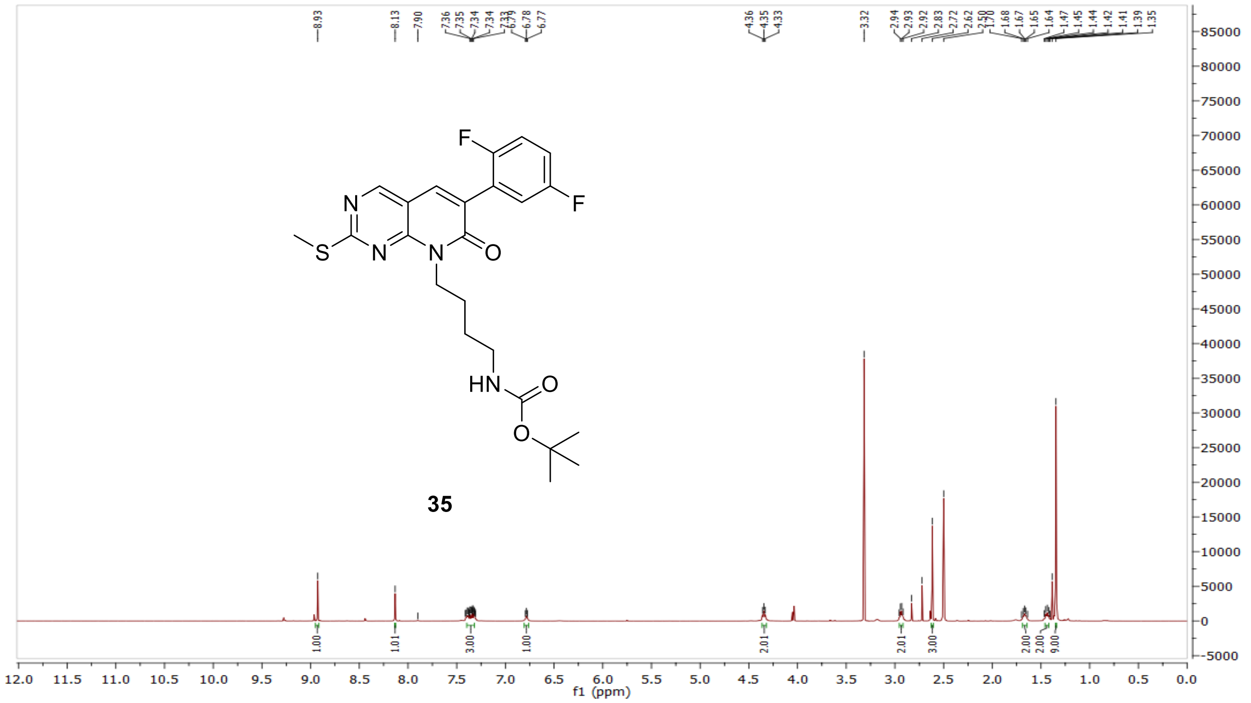


[13]C NMR compound **35:**

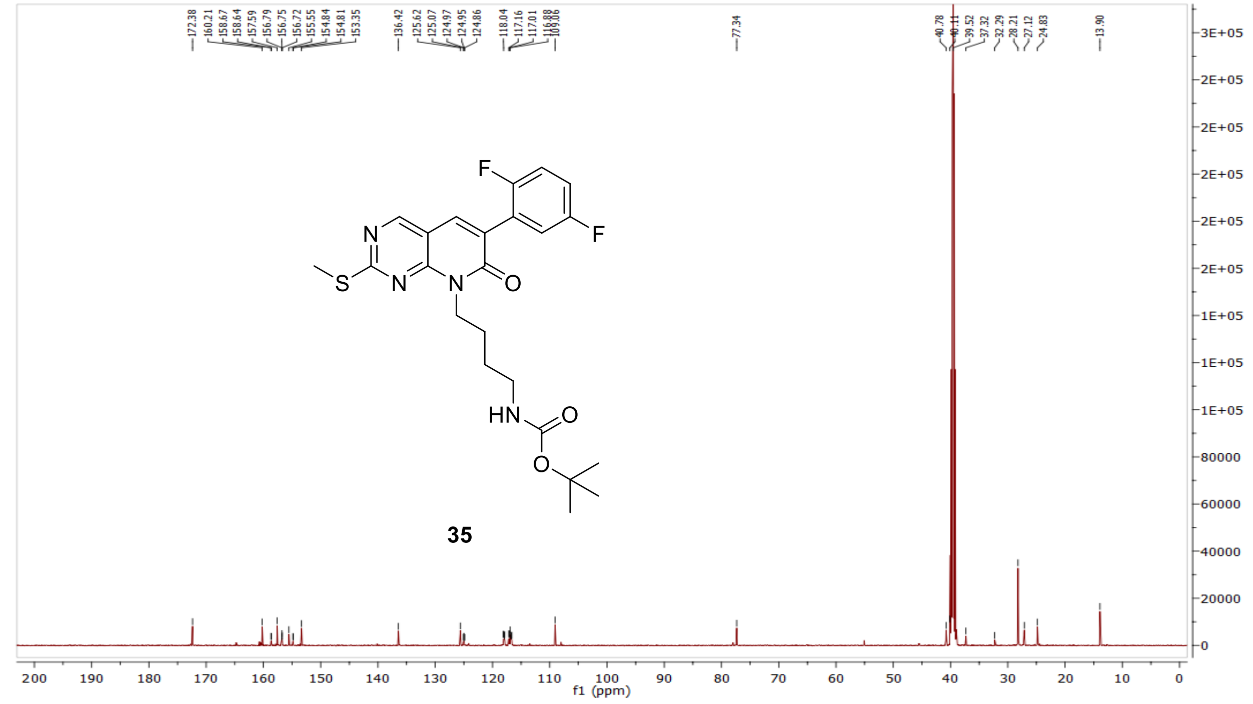

MS (ESI+) compound **35**:

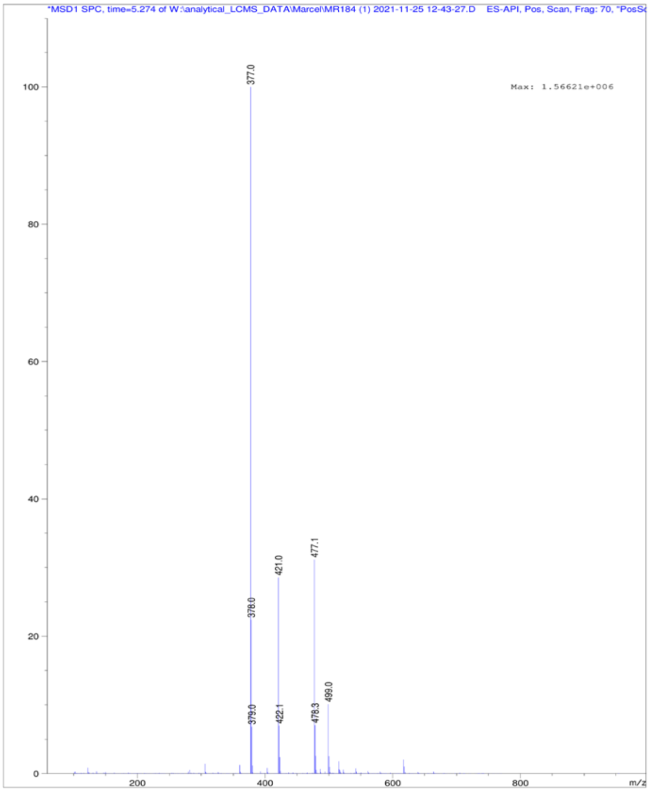

$^1$H NMR compound **36**:

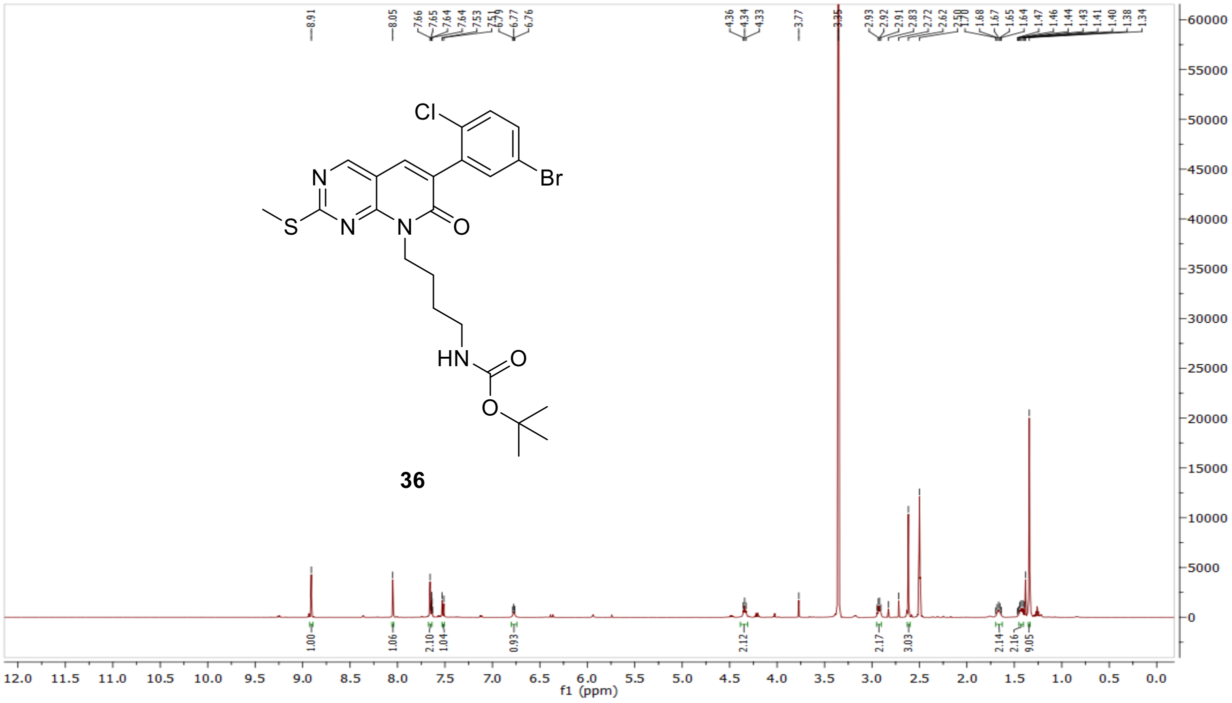

$^{13}$C NMR compound **36:**

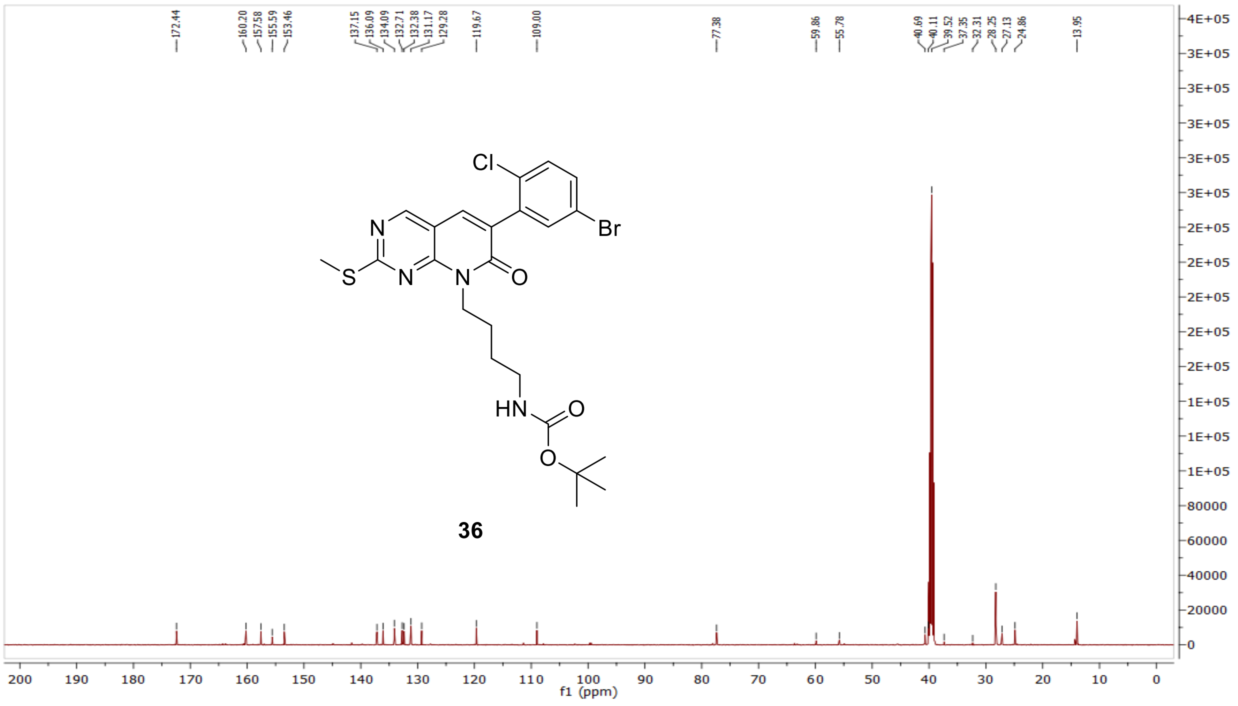


MS (ESI+) compound **36**:

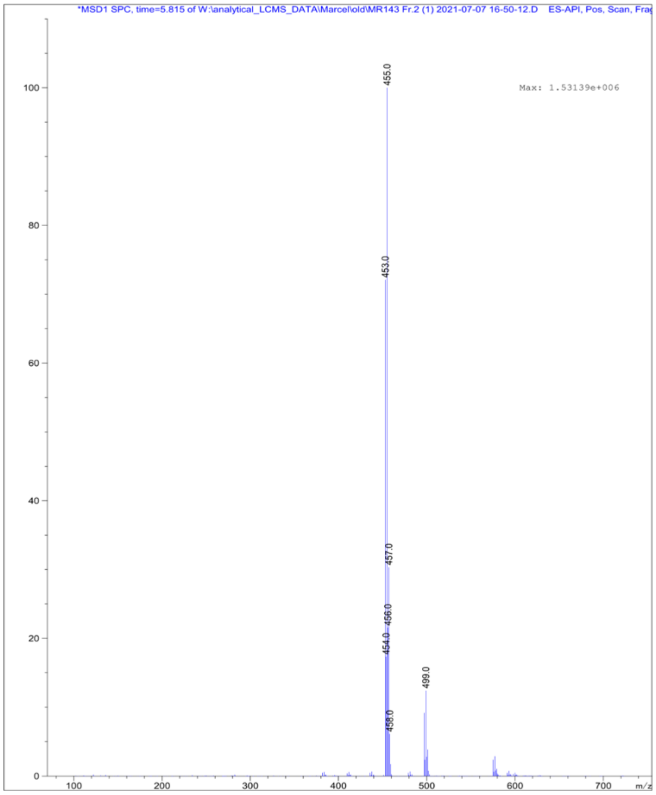

[1]H NMR compound **37**:

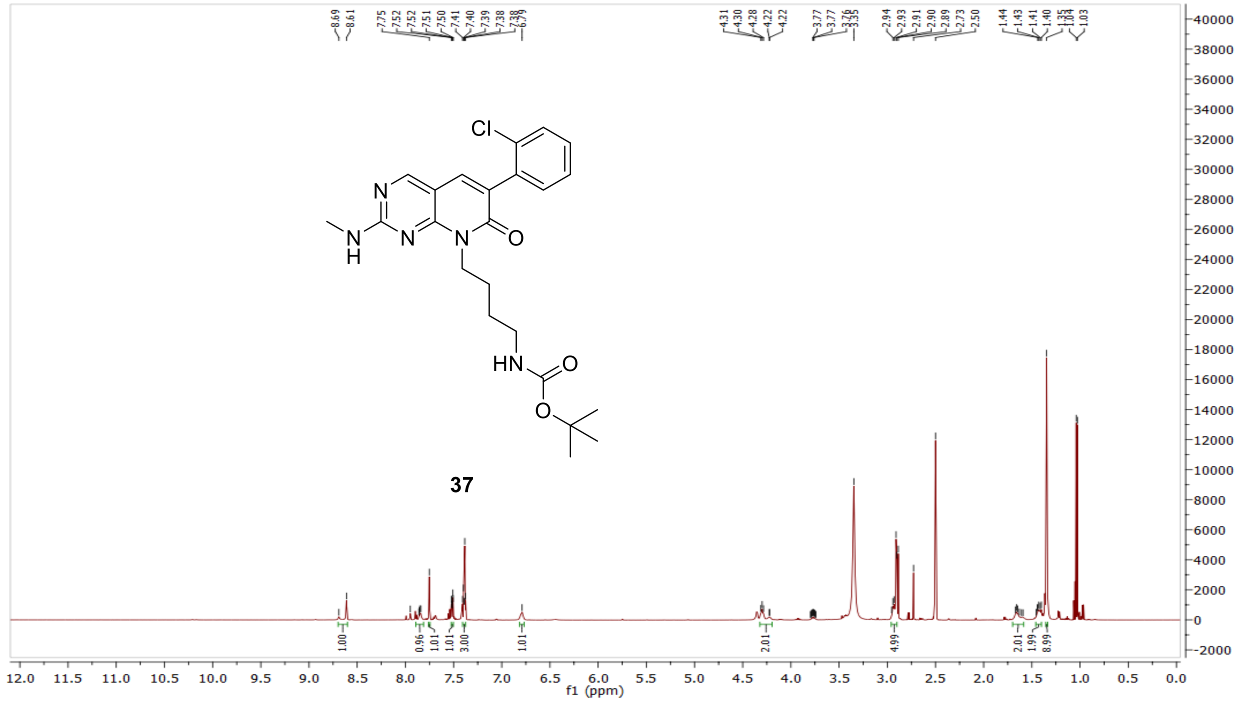

[13]C NMR compound **37:**

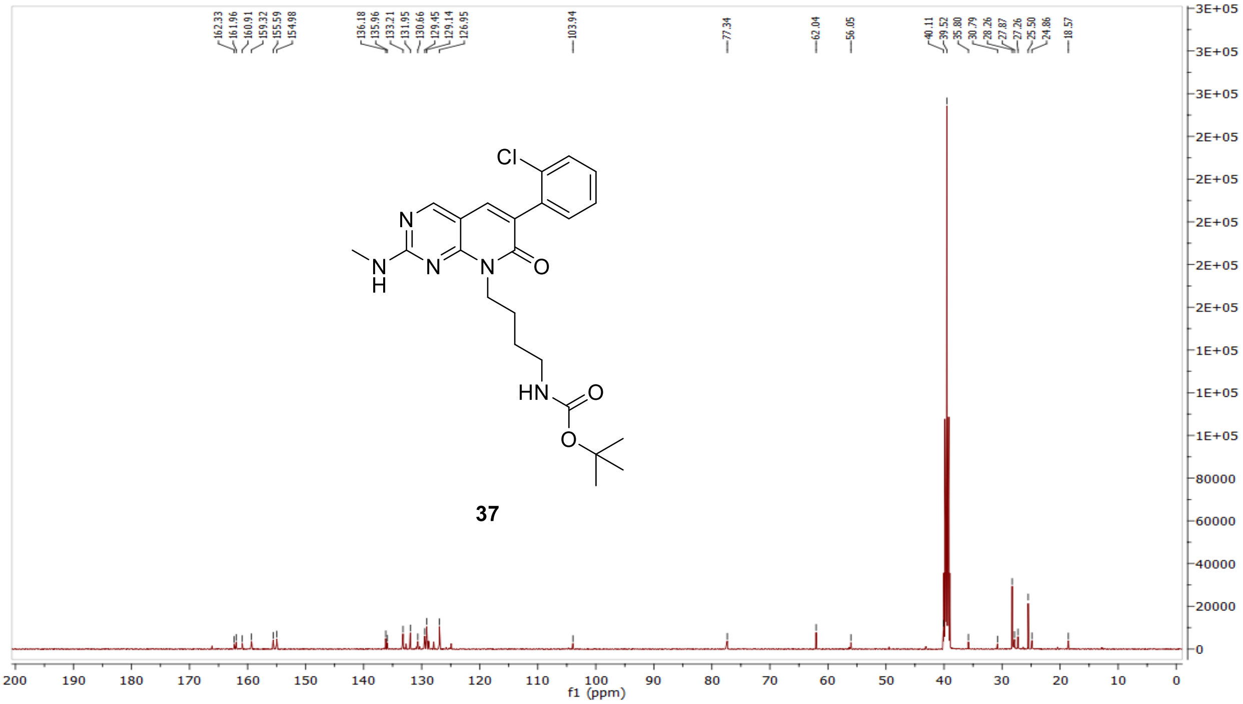

MS (ESI+) compound **37**:

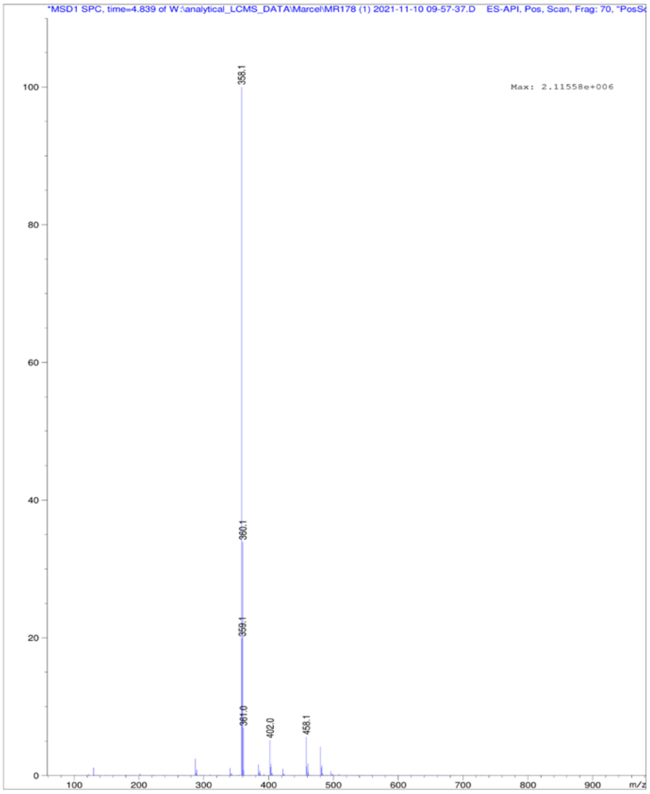


$^1$H NMR compound **38**:

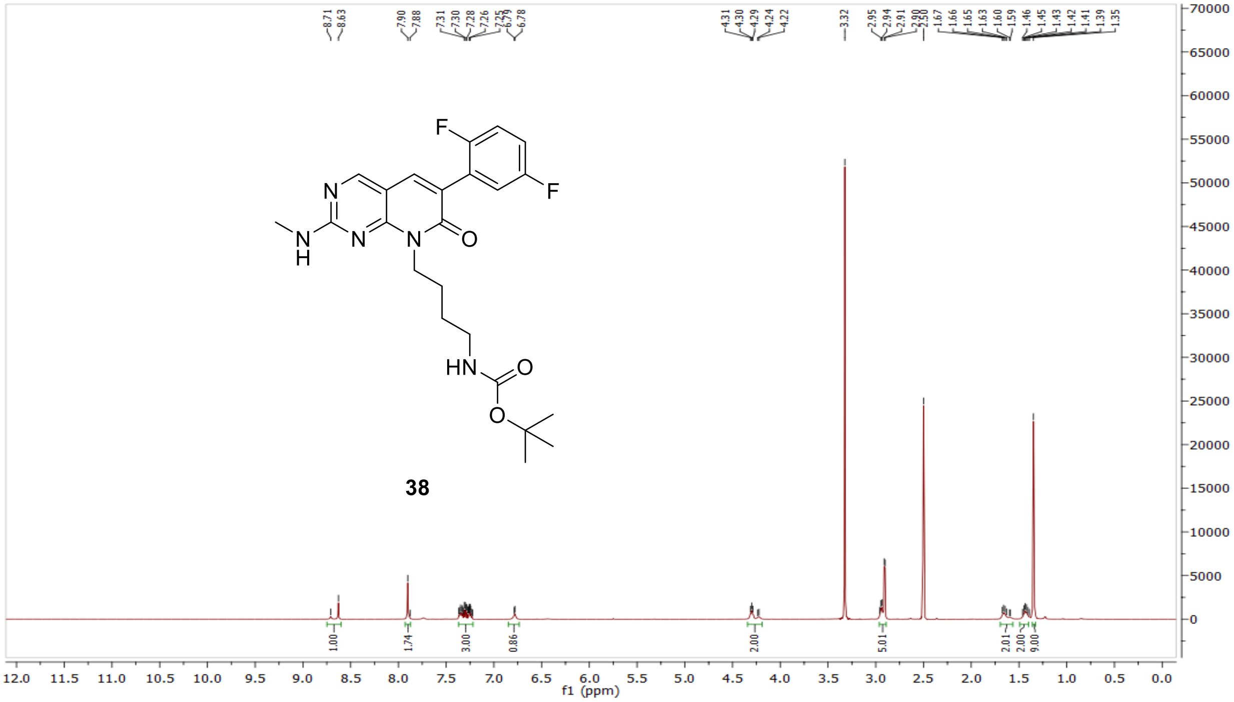

$^{13}$C NMR compound **38:**

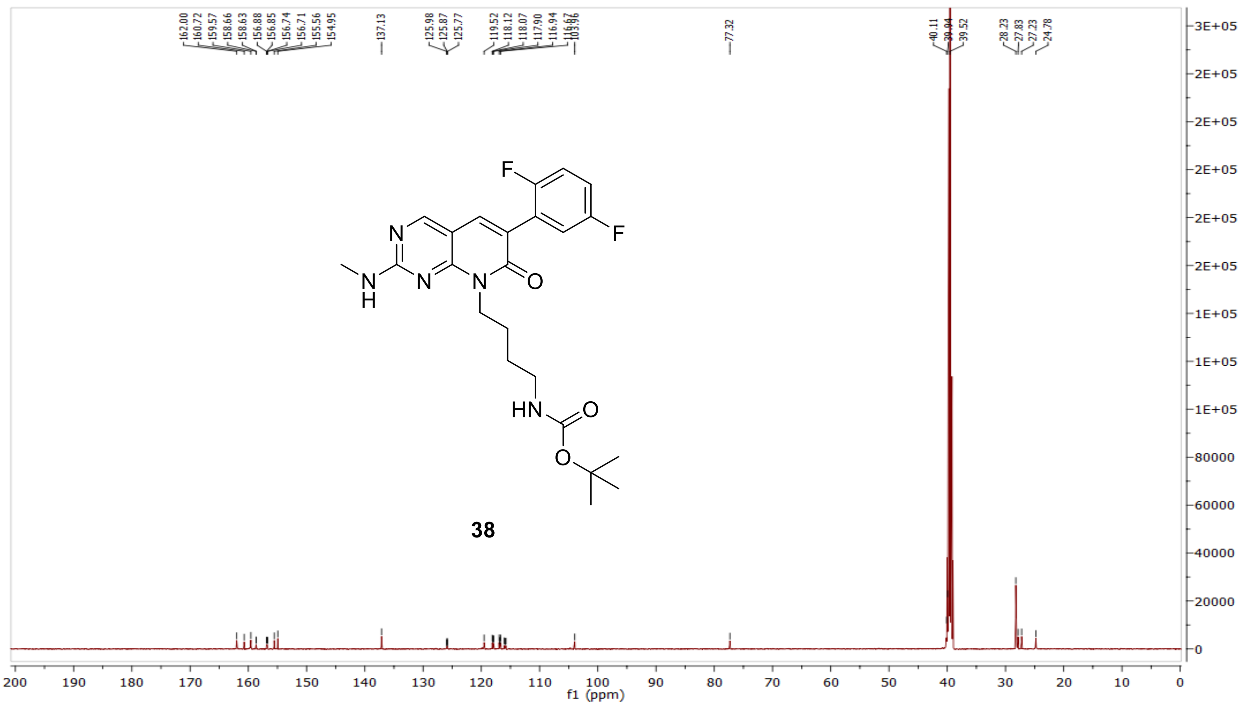


MS (ESI+) compound **38**:

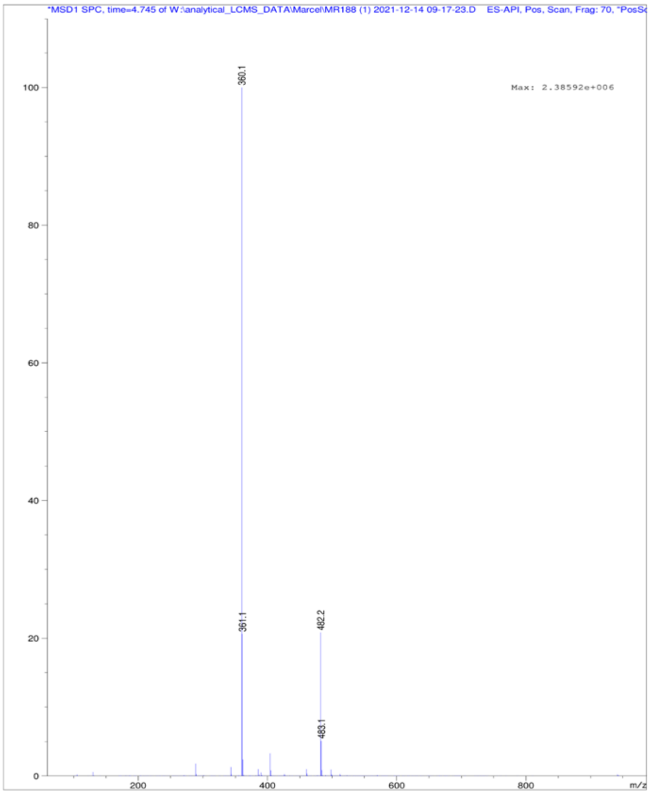

[1]H NMR compound **39**:

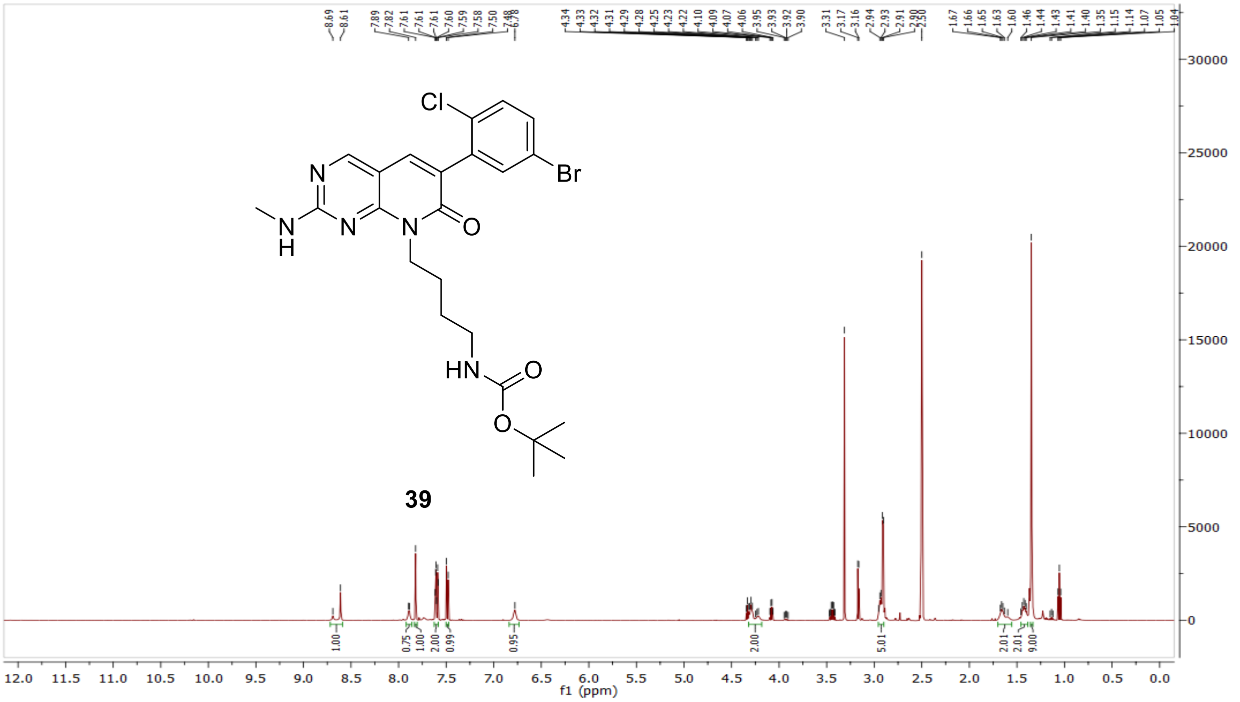


[13]C NMR compound **39:**

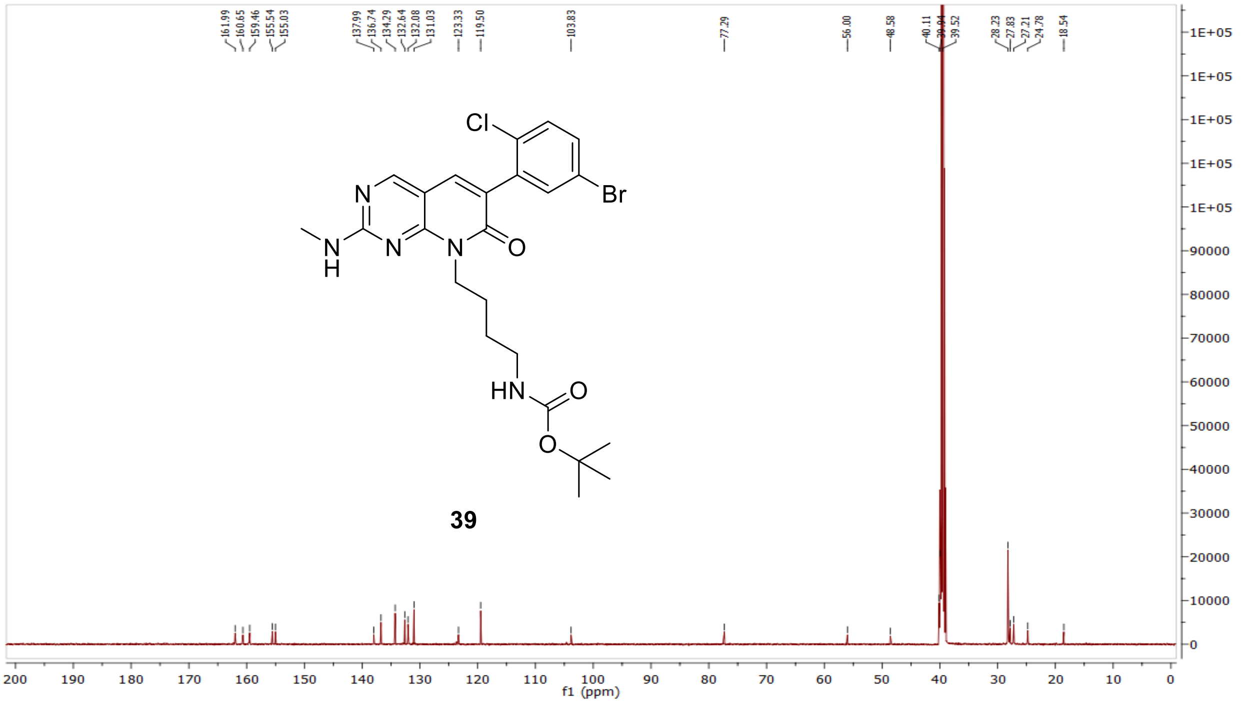

MS (ESI+) compound **39**:

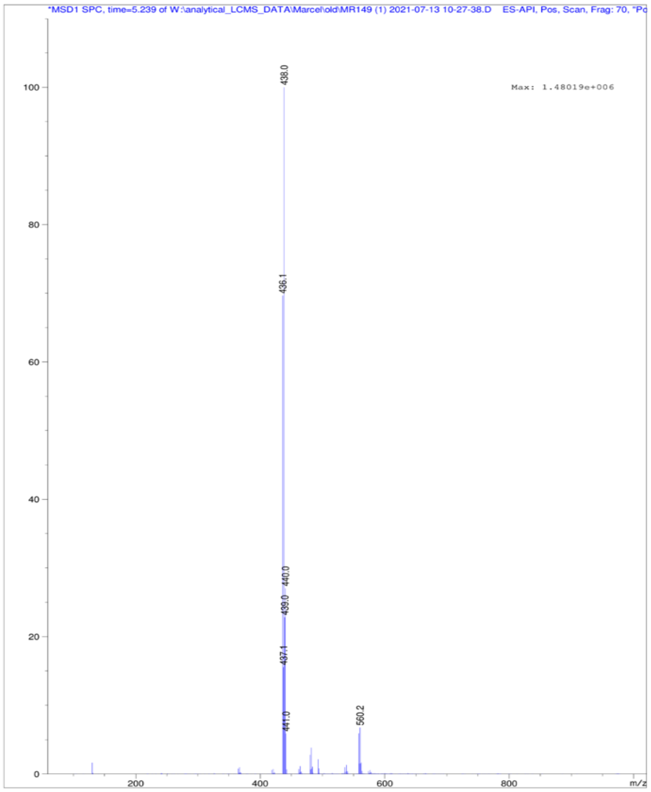

[1]H NMR compound **4**:

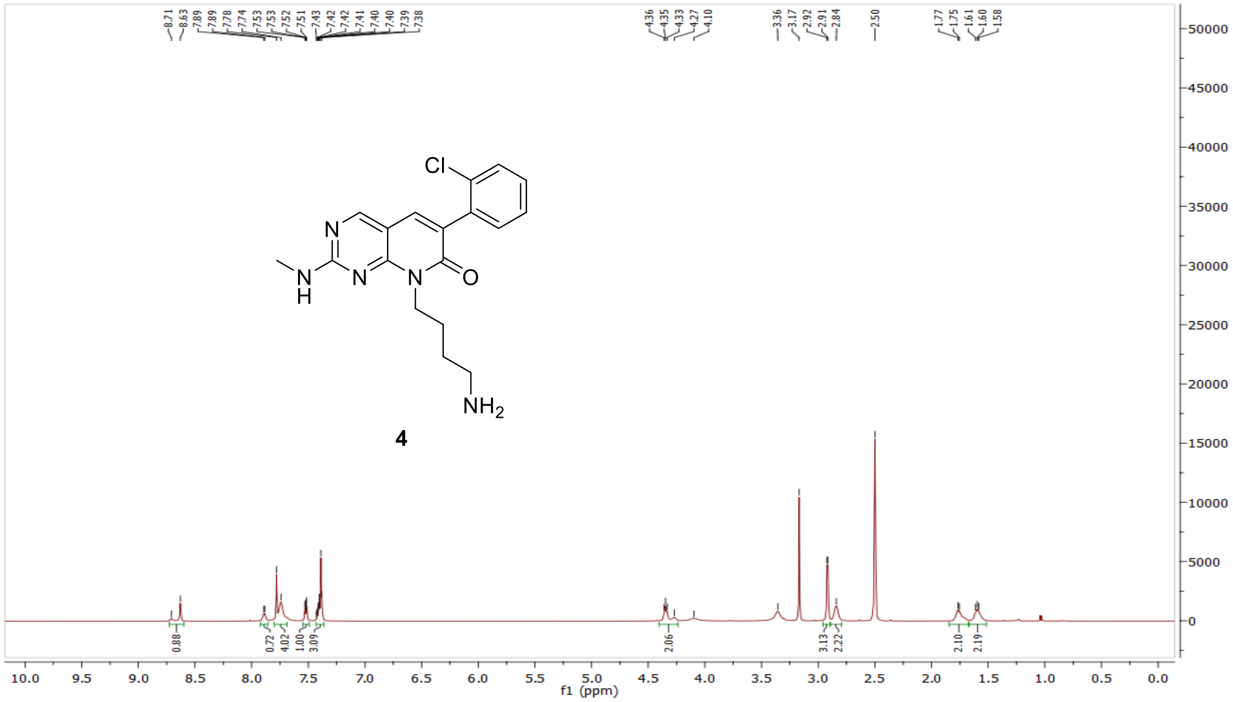

$^{13}$C NMR compound **4:**

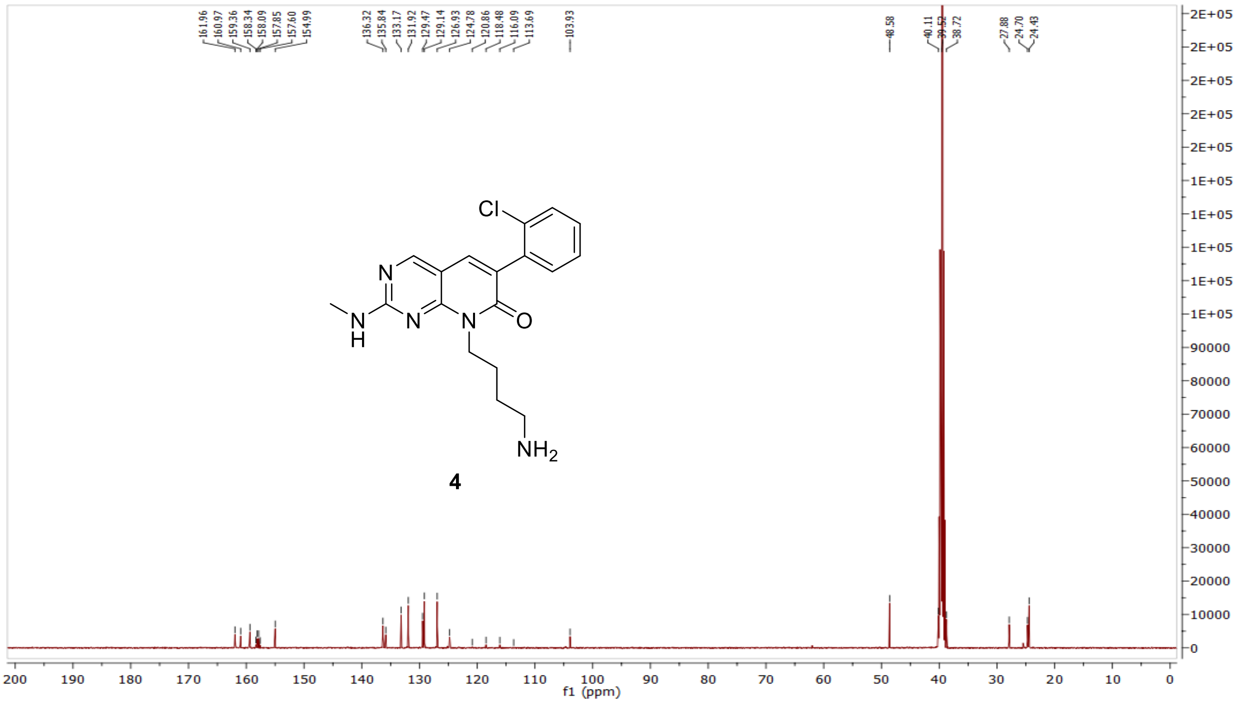


HRMS (FTMS +p MALDI) compound **4**:

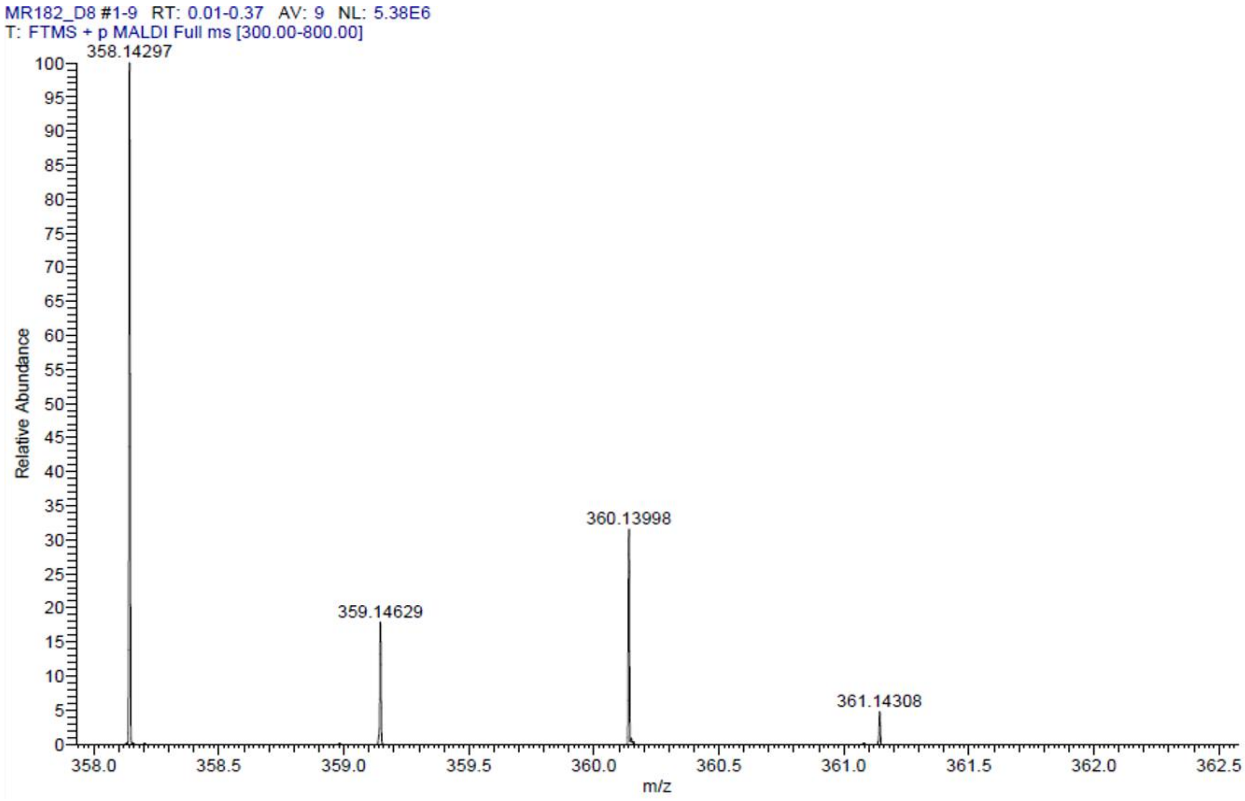

HPLC-chromatograms compound **4**:

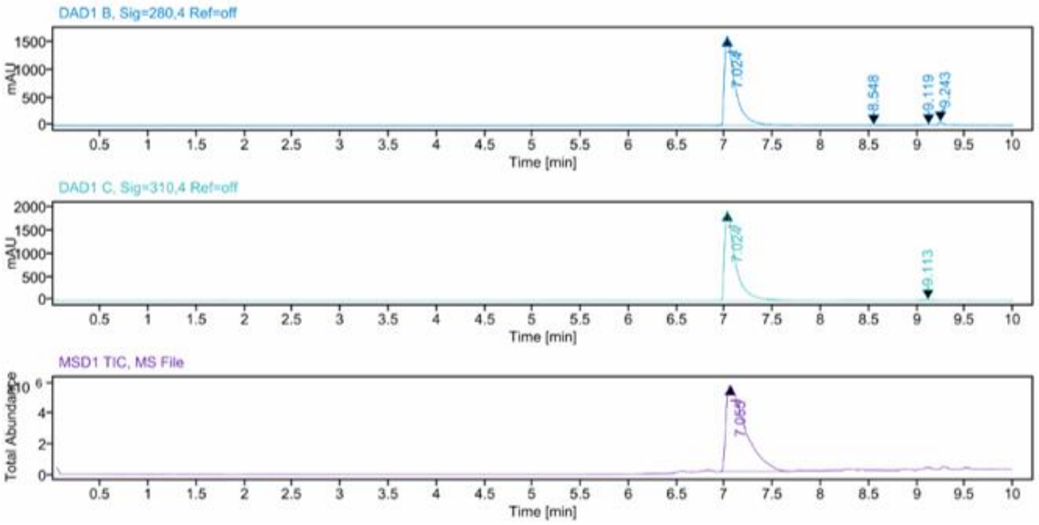


**Sample Purity**

Signal Description DAD1 B, Sig=280,4 Ref=off

| Sample Name | Name | RT | Width | Area | Area% | Height |
|---|---|---|---|---|---|---|
| MR182 | | 7.024 | 0.106 | 12452.0234 | 97.93 | 1616.9824 |
| MR182 | | 8.548 | 0.044 | 13.3111 | 0.10 | 4.3513 |
| MR182 | | 9.119 | 0.056 | 49.9194 | 0.39 | 9.8359 |
| MR182 | | 9.243 | 0.048 | 199.4718 | 1.57 | 60.3556 |

Max Area% 97.934

UV Signal Purity>95% Pass

Signal Description DAD1 C, Sig=310,4 Ref=off

| Sample Name | Name | RT | Width | Area | Area% | Height |
|---|---|---|---|---|---|---|
| MR182 | | 7.024 | 0.107 | 15087.4609 | 99.84 | 1939.1698 |
| MR182 | | 9.113 | 0.128 | 24.9326 | 0.16 | 3.6185 |

Max Area% 99.835

UV Signal Purity>95% Pass

**MSD Apex Spectra**

**RT:**

7.055

**Sample Name:**

MR182

**Signal Name:**

MSD1TIC

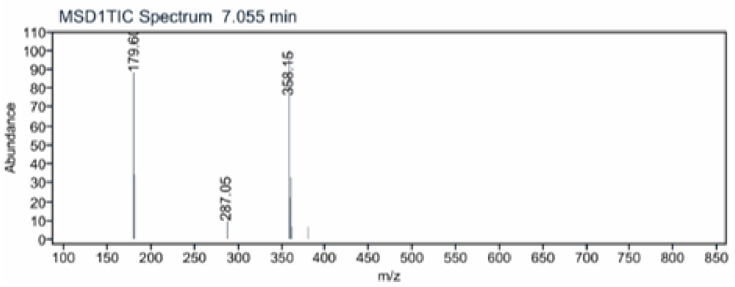

$^{1}H$ NMR compound **5**:

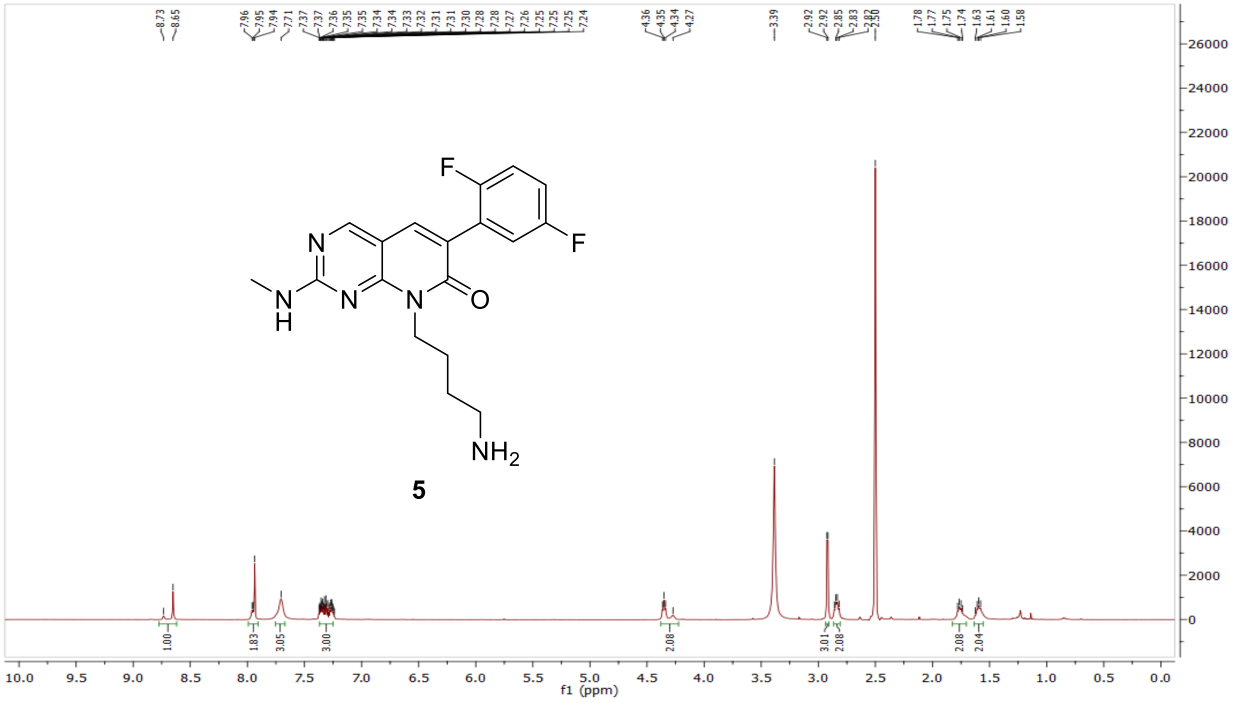

$^{13}C$ NMR compound **5:**

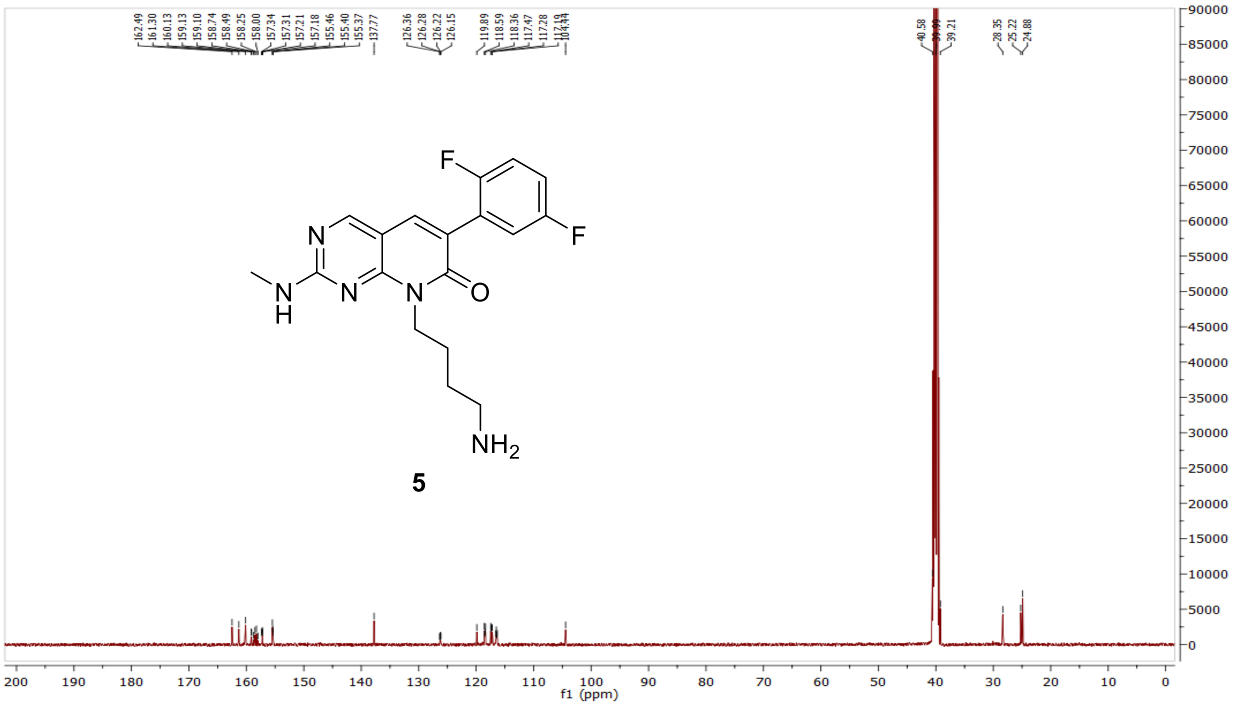

HRMS (FTMS +p MALDI) compound **5**:

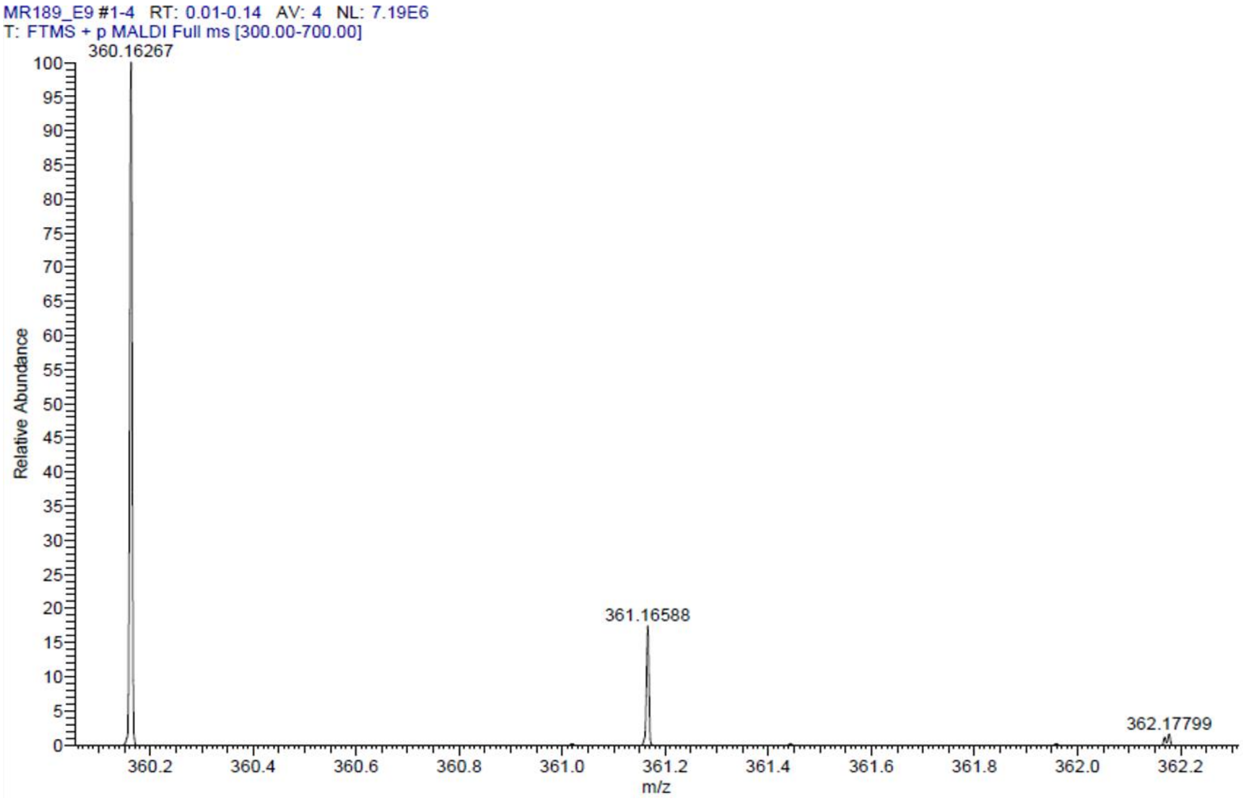

HPLC-chromatograms compound **5**

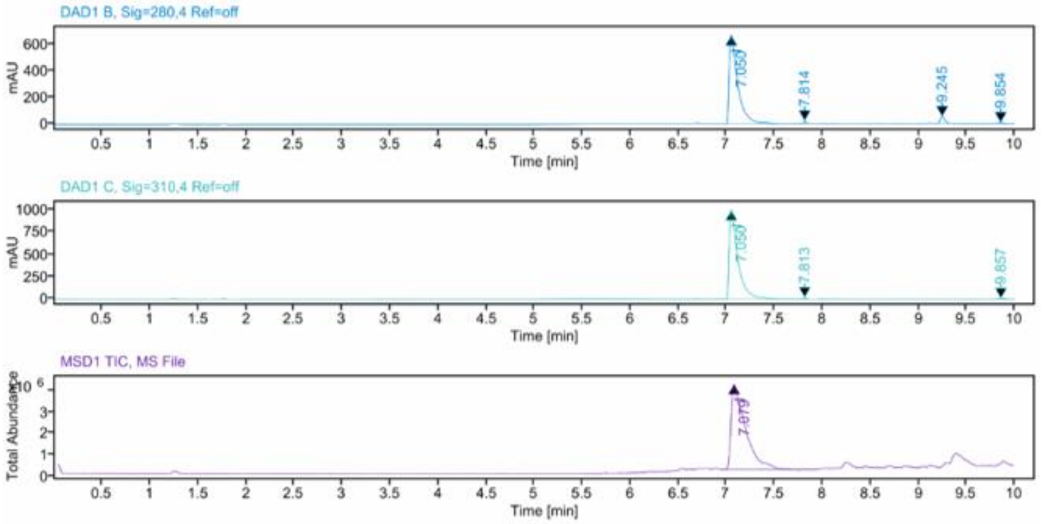


**Sample Purity**

Signal Description DAD1 B, Sig=280,4 Ref=off

| Sample Name | Name | RT | Width | Area | Area% | Height |
|---|---|---|---|---|---|---|
| MR189.12 | | 7.050 | 0.086 | 4139.7251 | 95.03 | 664.5875 |
| MR189.12 | | 7.814 | 0.012 | 27.6205 | 0.63 | 22.9001 |
| MR189.12 | | 9.245 | 0.049 | 157.3543 | 3.61 | 51.0237 |
| MR189.12 | | 9.854 | 0.052 | 31.5950 | 0.73 | 9.8429 |

Max Area% 95.029

UV Signal Purity>95% Pass

Signal Description DAD1 C, Sig=310,4 Ref=off

| Sample Name | Name | RT | Width | Area | Area% | Height |
|---|---|---|---|---|---|---|
| MR189.12 | | 7.050 | 0.086 | 6232.4932 | 99.09 | 992.8013 |
| MR189.12 | | 7.813 | 0.012 | 25.8458 | 0.41 | 20.2140 |
| MR189.12 | | 9.857 | 0.051 | 31.0812 | 0.49 | 8.7417 |

Max Area% 99.095

UV Signal Purity>95% Pass

**MSD Apex Spectra**

**RT:**

7.079

**Sample Name:**

MR189.12

**Signal Name:**

MSD1TIC

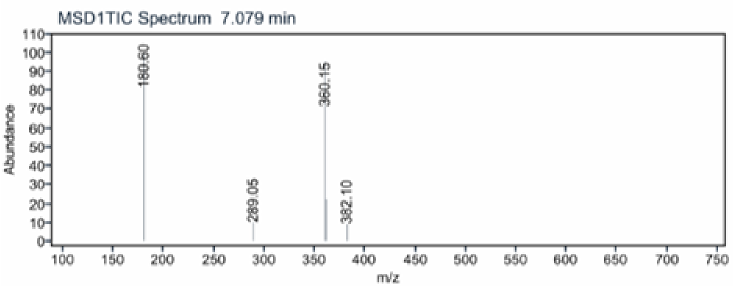

$^{1}H$ NMR compound **6**:

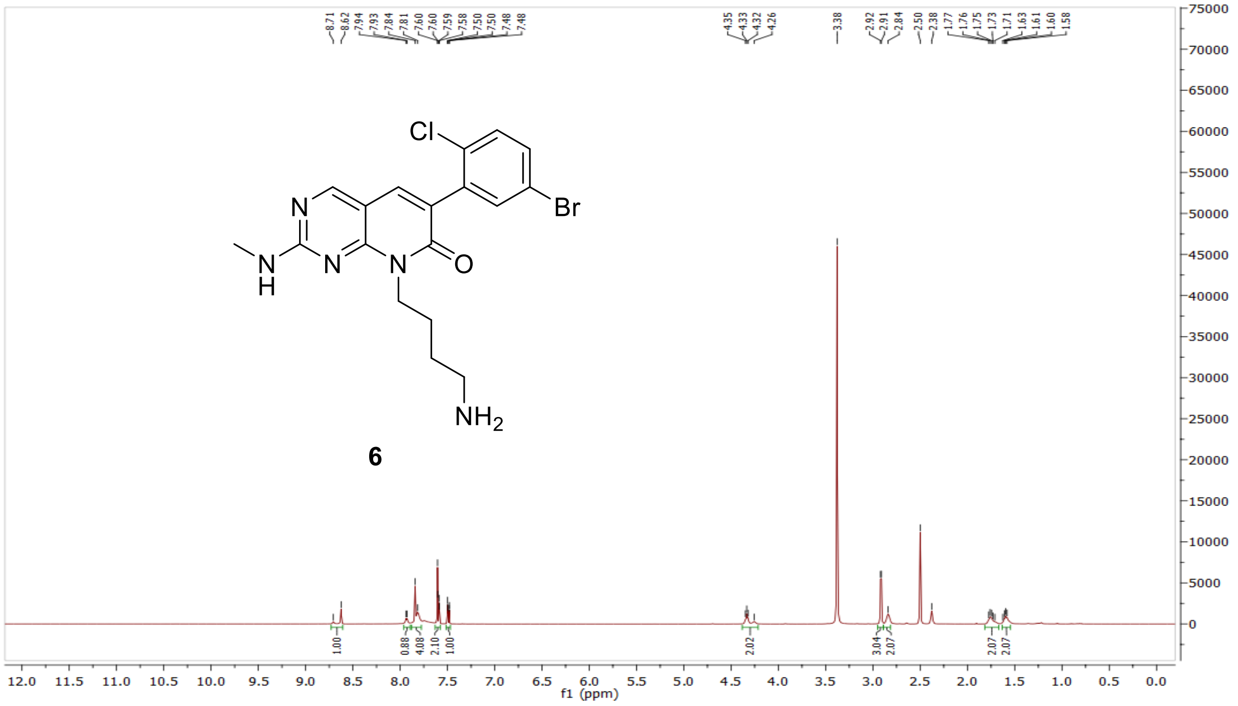

$^{13}C$ NMR compound **6:**

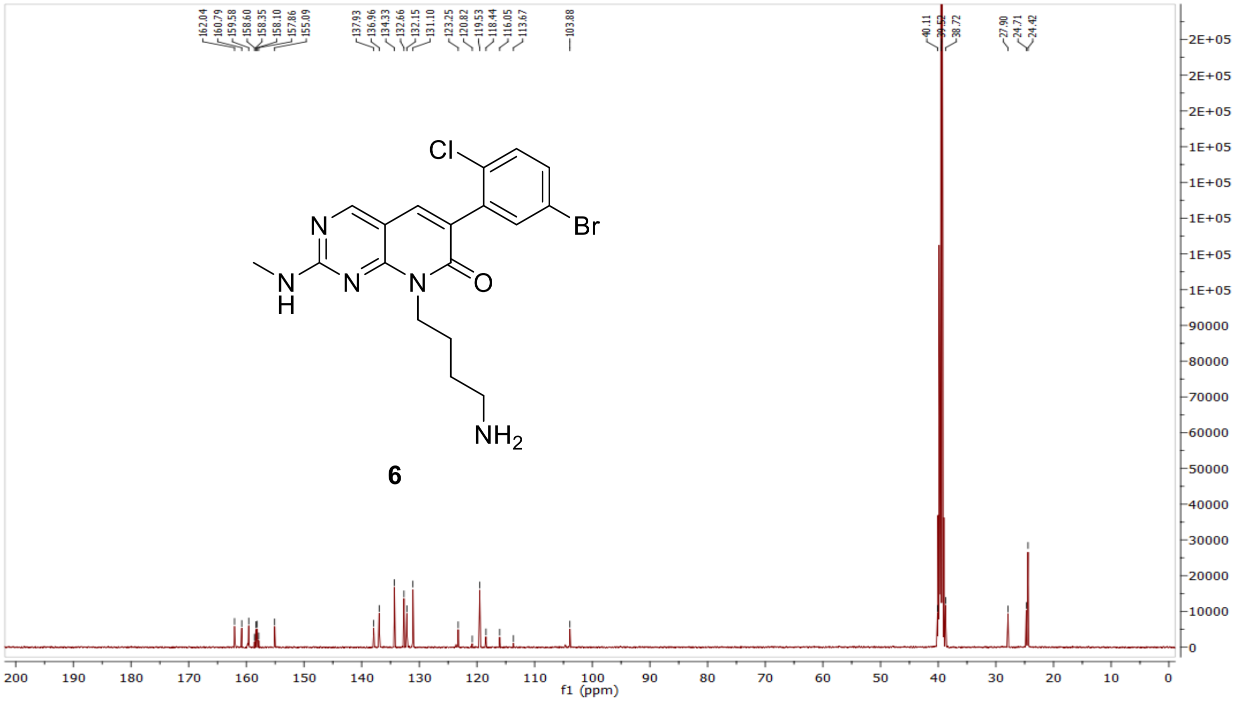

HRMS (FTMS +p MALDI) compound **6**:

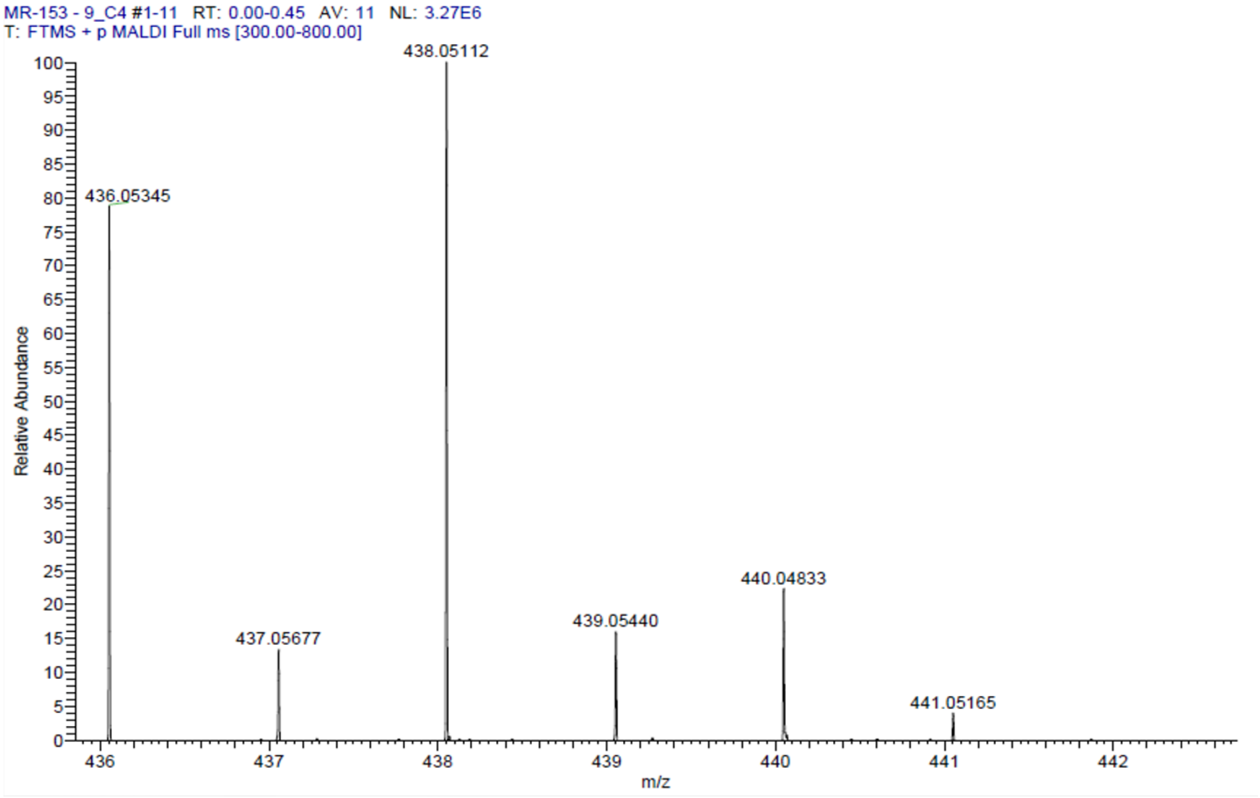

HPLC-chromatograms compound **6**:

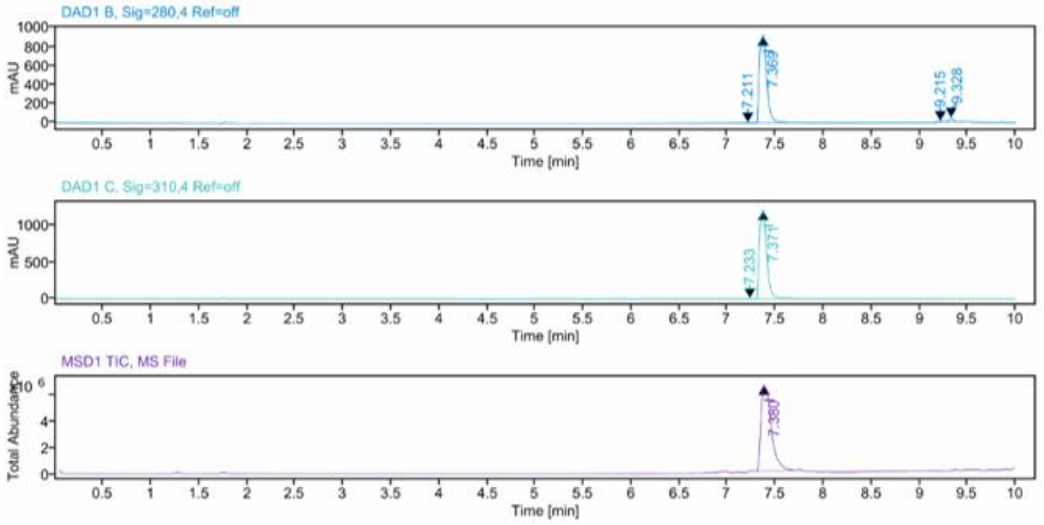


**Sample Purity**

Signal Description DAD1 B, Sig=280,4 Ref=off

| Sample Name | Name | RT | Width | Area | Area% | Height |
|---|---|---|---|---|---|---|
| MR153.9 | | 7.211 | 0.060 | 8.1553 | 0.16 | 2.4581 |
| MR153.9 | | 7.369 | 0.083 | 4866.3120 | 96.60 | 922.8562 |
| MR153.9 | | 9.215 | 0.042 | 25.3915 | 0.50 | 8.6552 |
| MR153.9 | | 9.328 | 0.043 | 137.6212 | 2.73 | 43.4542 |

Max Area% 96.602

UV Signal Purity>95% Pass

Signal Description DAD1 C, Sig=310,4 Ref=off

| Sample Name | Name | RT | Width | Area | Area% | Height |
|---|---|---|---|---|---|---|
| MR153.9 | | 7.233 | 0.060 | 8.9552 | 0.14 | 2.4585 |
| MR153.9 | | 7.371 | 0.082 | 6463.1943 | 99.86 | 1217.7488 |

Max Area% 99.862

UV Signal Purity>95% Pass

**MSD Apex Spectra**

**RT:**

7.380

**Sample Name:**

MR153.9

**Signal Name:**

MSD1TIC

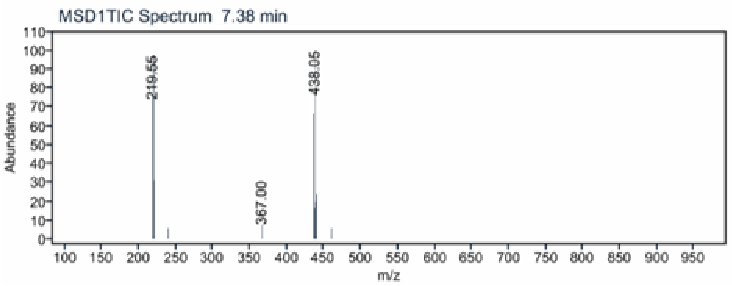

$^{1}H$ NMR compound **7**:

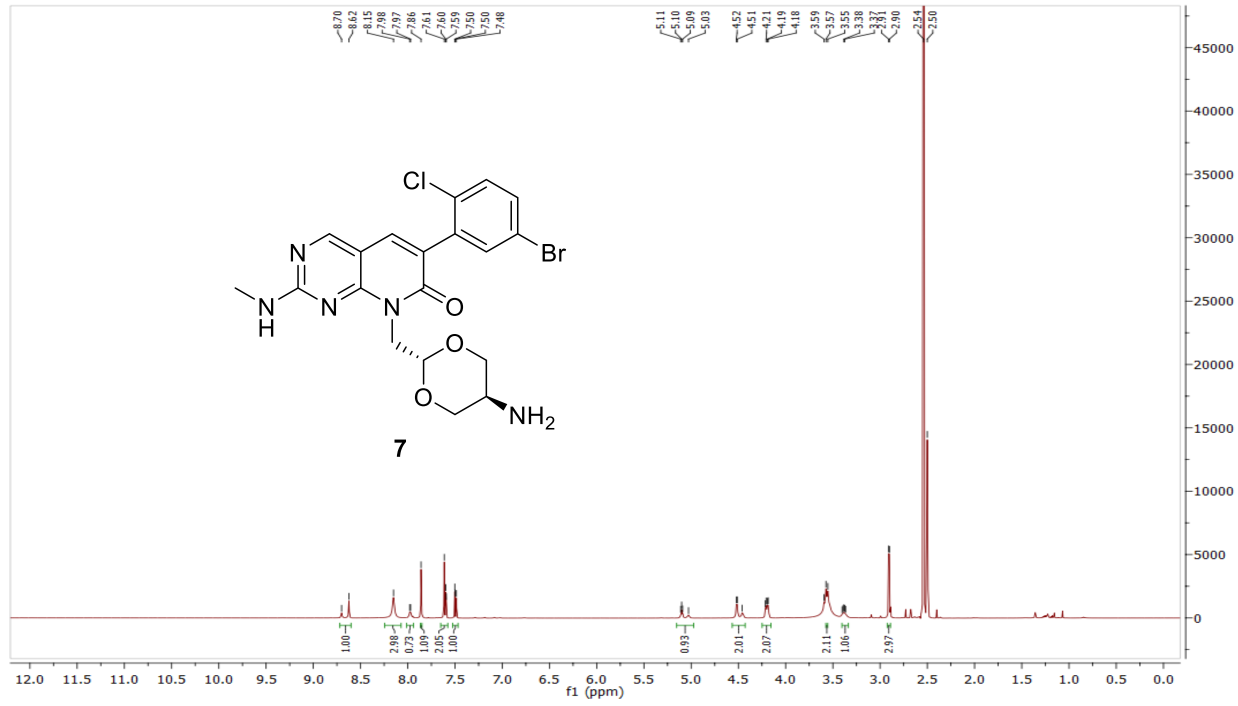


$^{13}C$ NMR compound **7:**

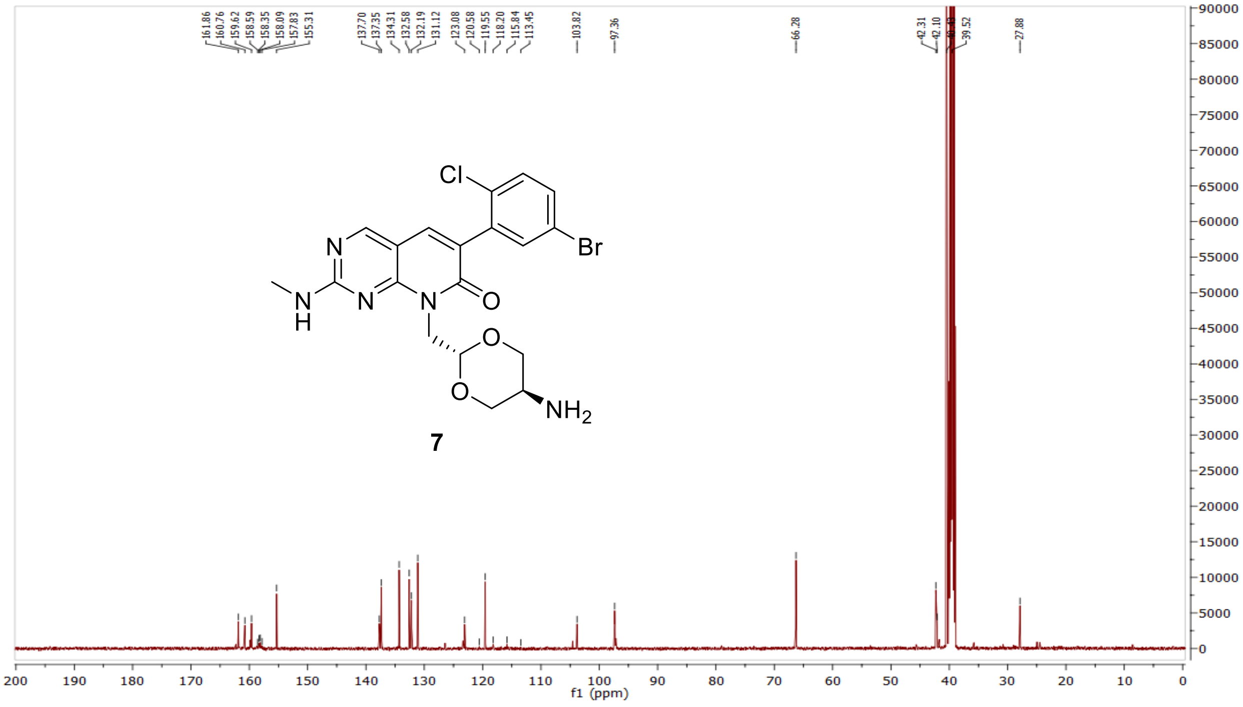

HRMS (FTMS +p MALDI) compound **7**:

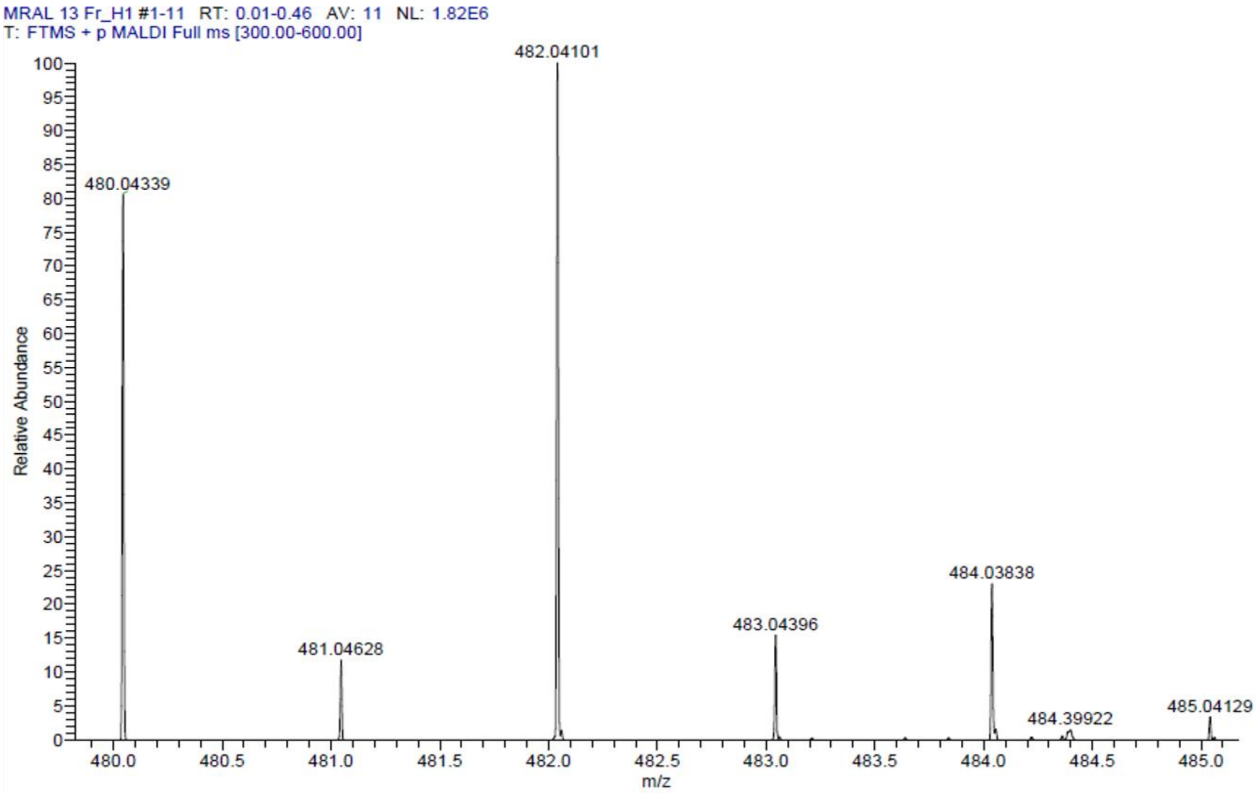

HPLC-chromatograms compound **7**:

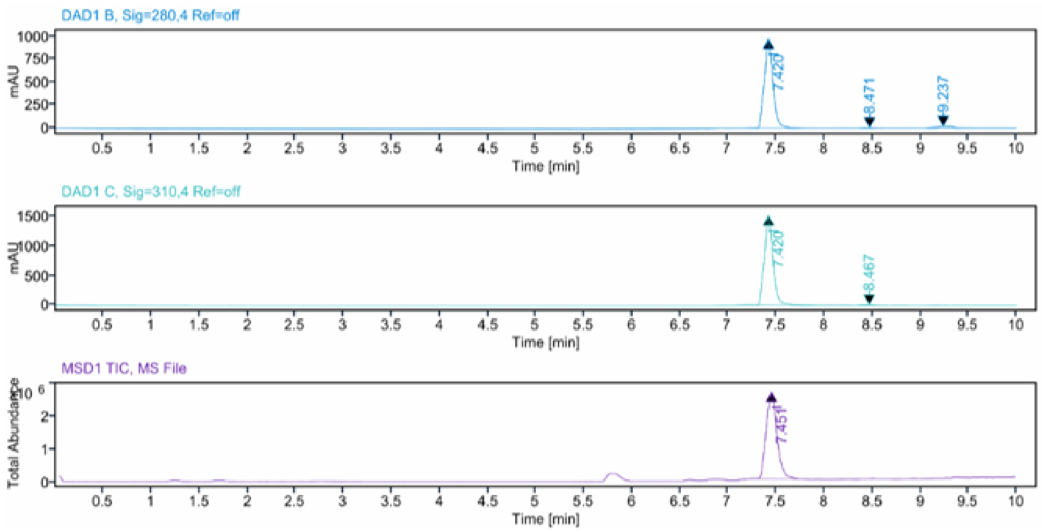


**Sample Purity**

Signal Description DAD1 B, Sig=280,4 Ref=off

| Sample Name | Name | RT | Width | Area | Area% | Height |
|---|---|---|---|---|---|---|
| MRAL13 Fr.1 HPLC | | 7.420 | 0.092 | 6031.2842 | 95.42 | 979.0758 |
| MRAL13 Fr.1 HPLC | | 8.471 | 0.132 | 39.4749 | 0.62 | 4.7332 |
| MRAL13 Fr.1 HPLC | | 9.237 | 0.189 | 249.8396 | 3.95 | 19.5297 |

Max Area% 95.423

UV Signal Purity>95% Pass

Signal Description DAD1 C, Sig=310,4 Ref=off

| Sample Name | Name | RT | Width | Area | Area% | Height |
|---|---|---|---|---|---|---|
| MRAL13 Fr.1 HPLC | | 7.420 | 0.092 | 9379.3438 | 99.43 | 1523.5389 |
| MRAL13 Fr.1 HPLC | | 8.467 | 0.140 | 53.8405 | 0.57 | 6.6170 |

Max Area% 99.429

UV Signal Purity>95% Pass

**MSD Apex Spectra**

**RT:**

7.451

**Sample Name:**

MRAL13 Fr.1 HPLC

**Signal Name:**

MSD1TIC

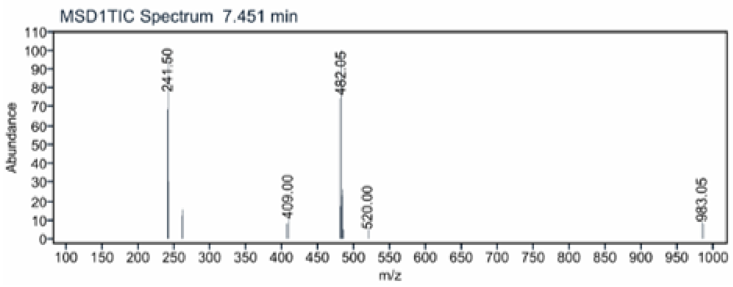

[1]H NMR compound **8**:

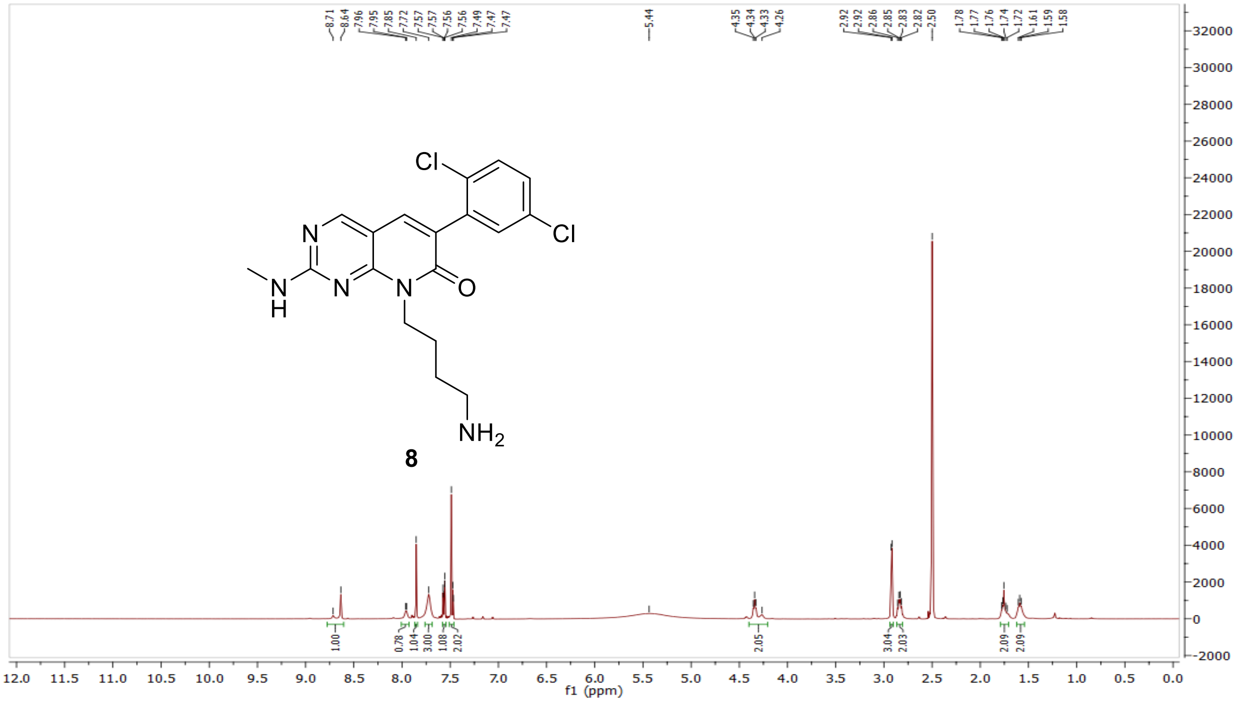


[13]C NMR compound **8:**

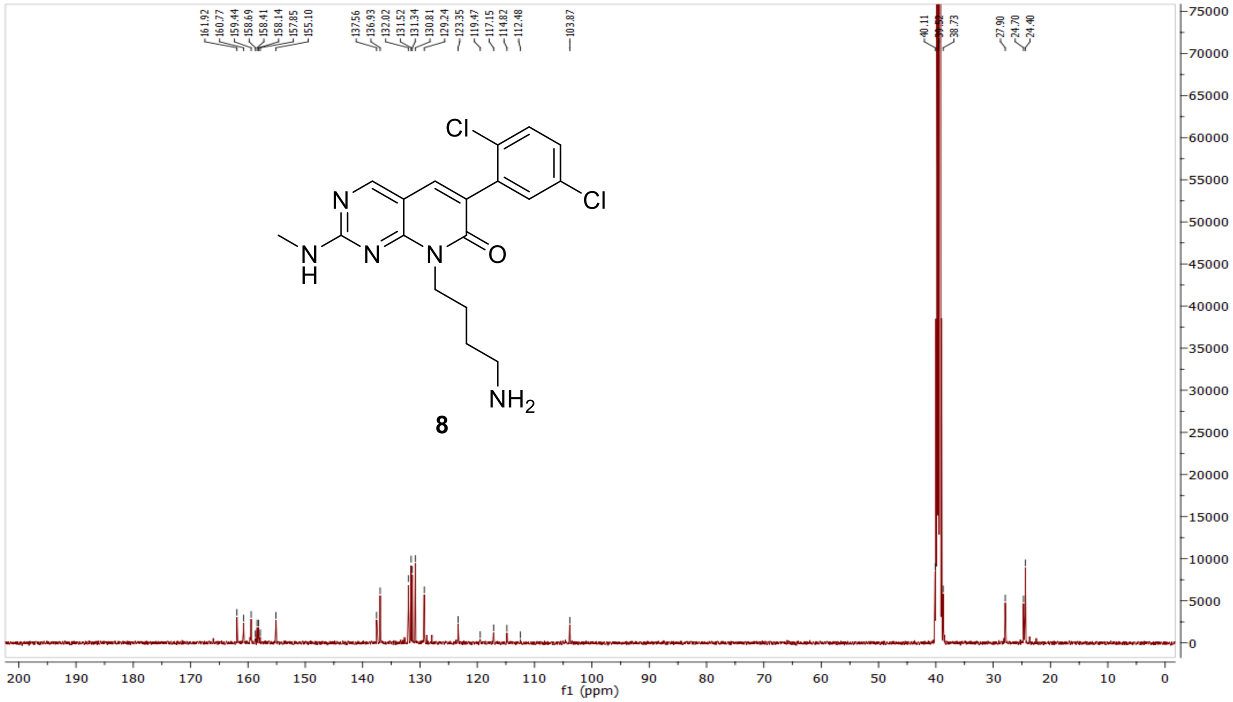

HRMS (FTMS +p MALDI) compound **8**:

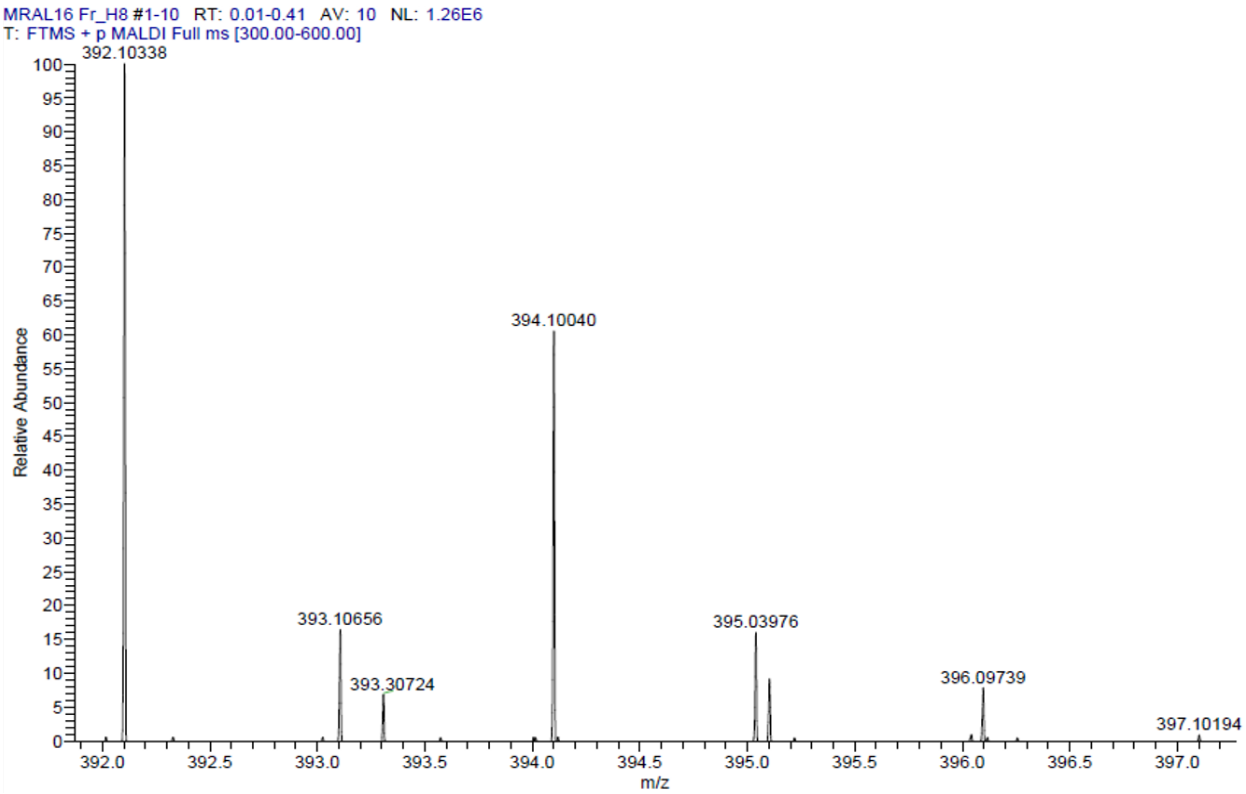

HPLC-chromatograms compound **8**:

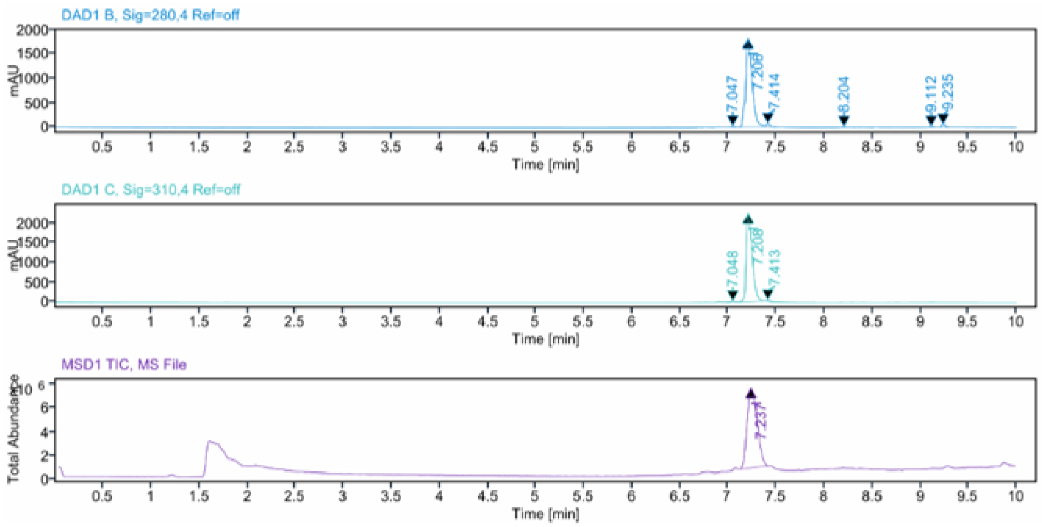


**Sample Purity**

Signal Description DAD1 B, Sig=280,4 Ref=off

| Sample Name | Name | RT | Width | Area | Area% | Height |
|---|---|---|---|---|---|---|
| MRAL16 Fr.2 HPLC | | 7.047 | 0.025 | 42.0888 | 0.47 | 24.5978 |
| MRAL16 Fr.2 HPLC | | 7.206 | 0.069 | 8487.9473 | 95.02 | 1824.8907 |
| MRAL16 Fr.2 HPLC | | 7.414 | 0.031 | 124.0515 | 1.39 | 59.6723 |
| MRAL16 Fr.2 HPLC | | 8.204 | 0.034 | 46.2747 | 0.52 | 22.0561 |
| MRAL16 Fr.2 HPLC | | 9.112 | 0.042 | 59.9236 | 0.67 | 21.8613 |
| MRAL16 Fr.2 HPLC | | 9.235 | 0.038 | 172.3655 | 1.93 | 63.7658 |

Max Area% 95.022

UV Signal Purity>95% Pass

Signal Description DAD1 C, Sig=310,4 Ref=off

| Sample Name | Name | RT | Width | Area | Area% | Height |
|---|---|---|---|---|---|---|
| MRAL16 Fr.2 HPLC | | 7.048 | 0.027 | 60.5523 | 0.61 | 29.1264 |
| MRAL16 Fr.2 HPLC | | 7.208 | 0.068 | 9816.9141 | 98.69 | 2246.9172 |
| MRAL16 Fr.2 HPLC | | 7.413 | 0.030 | 69.9643 | 0.70 | 34.7599 |

Max Area% 98.688

UV Signal Purity>95% Pass

**MSD Apex Spectra**

**RT:**

7.237

**Sample Name:**

MRAL16 Fr.2 HPLC

**Signal Name:**

MSD1TIC

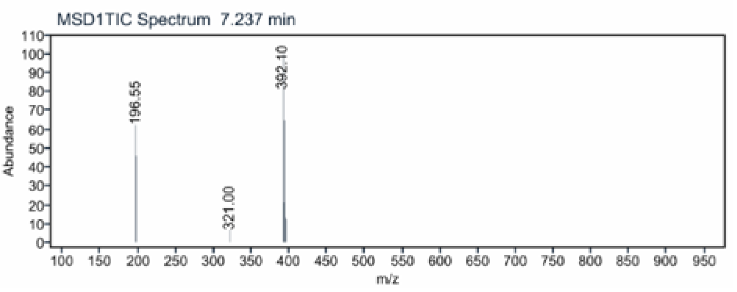


$^1$H NMR compound **9**:

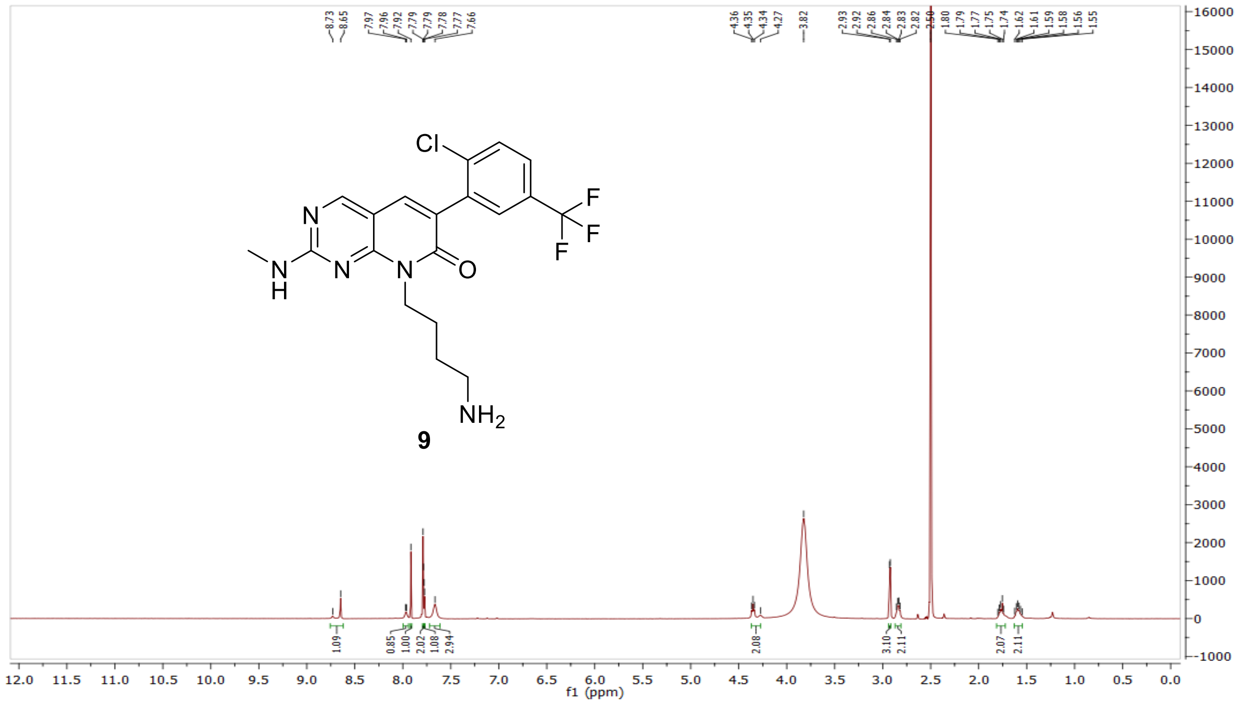

$^{13}$C NMR compound **9:**

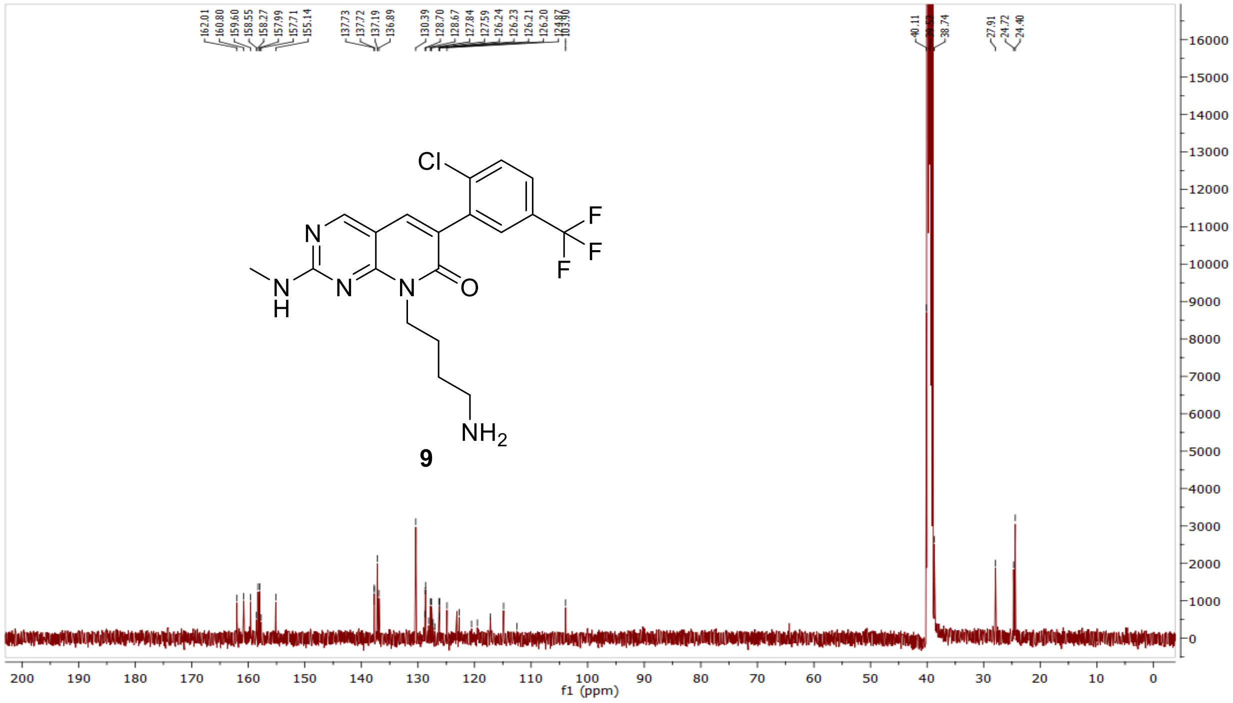


HRMS (FTMS +p MALDI) compound **9**:

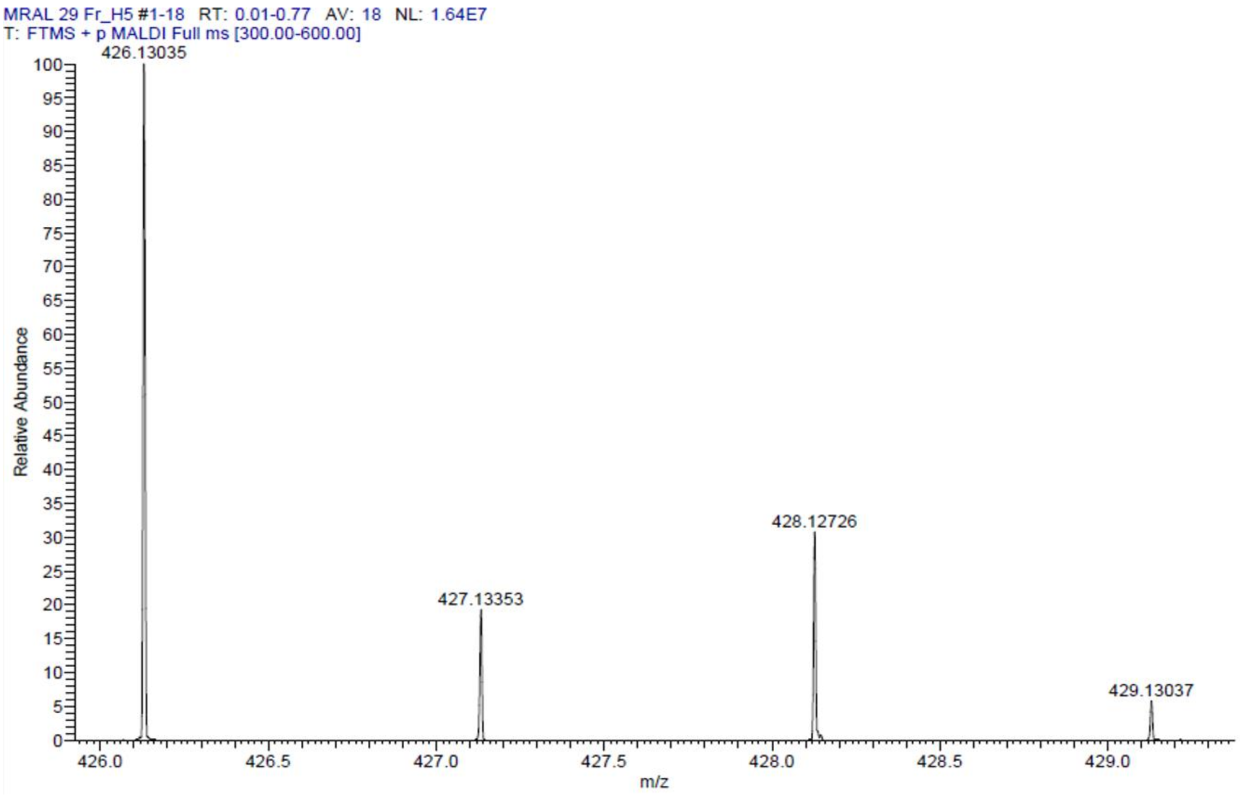

HPLC-chromatograms compound **9**:

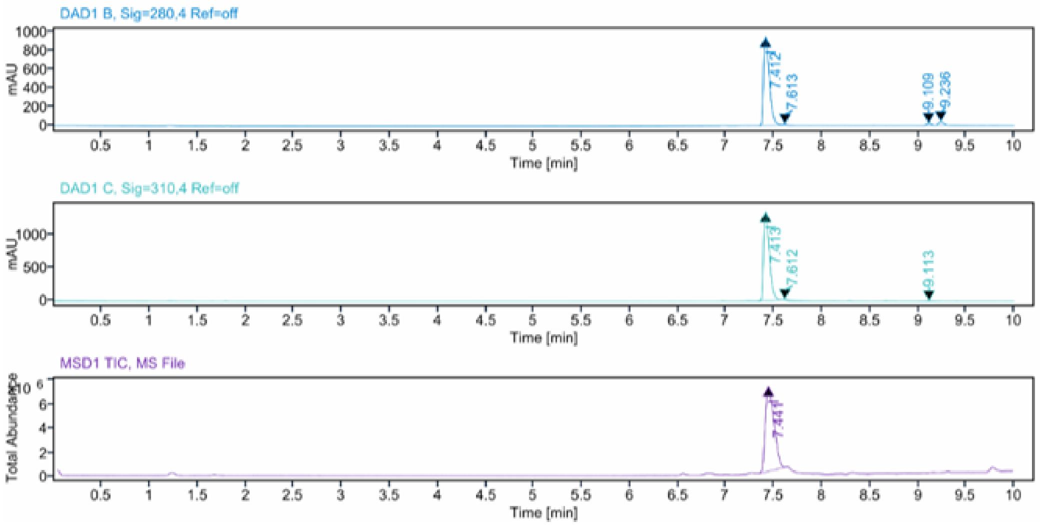


**Sample Purity**

Signal Description DAD1 B, Sig=280,4 Ref=off

| Sample Name | Name | RT | Width | Area | Area% | Height |
|---|---|---|---|---|---|---|
| MRAL29 Fr.1 HPLC | | 7.412 | 0.066 | 3995.0374 | 95.07 | 946.4332 |
| MRAL29 Fr.1 HPLC | | 7.613 | 0.039 | 17.6804 | 0.42 | 7.0205 |
| MRAL29 Fr.1 HPLC | | 9.109 | 0.056 | 60.5463 | 1.44 | 18.0339 |
| MRAL29 Fr.1 HPLC | | 9.236 | 0.059 | 128.9623 | 3.07 | 35.7230 |

Max Area% 95.070

UV Signal Purity>95% Pass

Signal Description DAD1 C, Sig=310,4 Ref=off

| Sample Name | Name | RT | Width | Area | Area% | Height |
|---|---|---|---|---|---|---|
| MRAL29 Fr.1 HPLC | | 7.413 | 0.066 | 5652.7402 | 99.31 | 1339.7534 |
| MRAL29 Fr.1 HPLC | | 7.612 | 0.040 | 25.7766 | 0.45 | 9.6920 |
| MRAL29 Fr.1 HPLC | | 9.113 | 0.072 | 13.6023 | 0.24 | 2.8035 |

Max Area% 99.308

UV Signal Purity>95% Pass

**MSD Apex Spectra**

**RT:**

7.441

**Sample Name:**

MRAL29 Fr.1 HPLC

**Signal Name:**

MSD1TIC

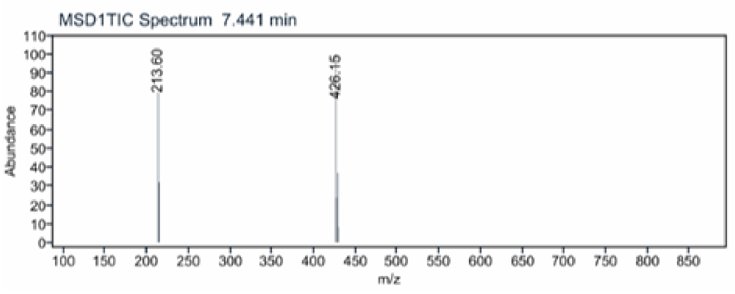

<sup></sup>[1]H NMR compound **10**:

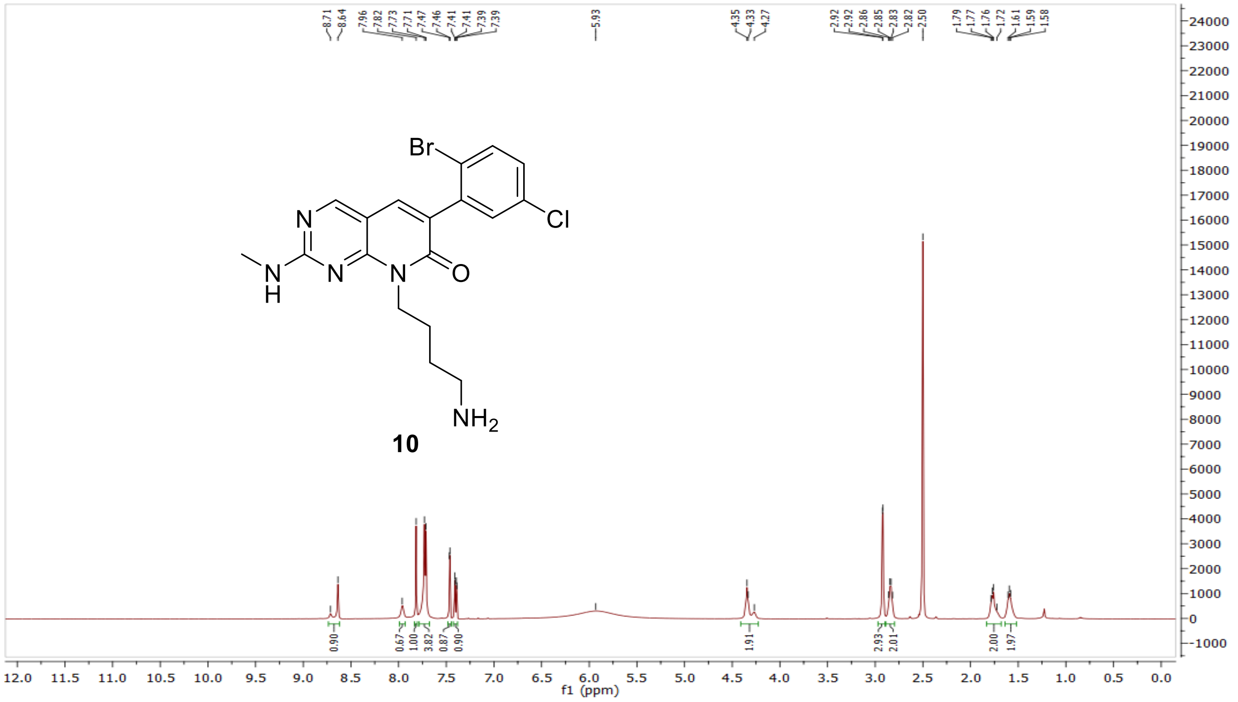

[13]C NMR compound **10:**

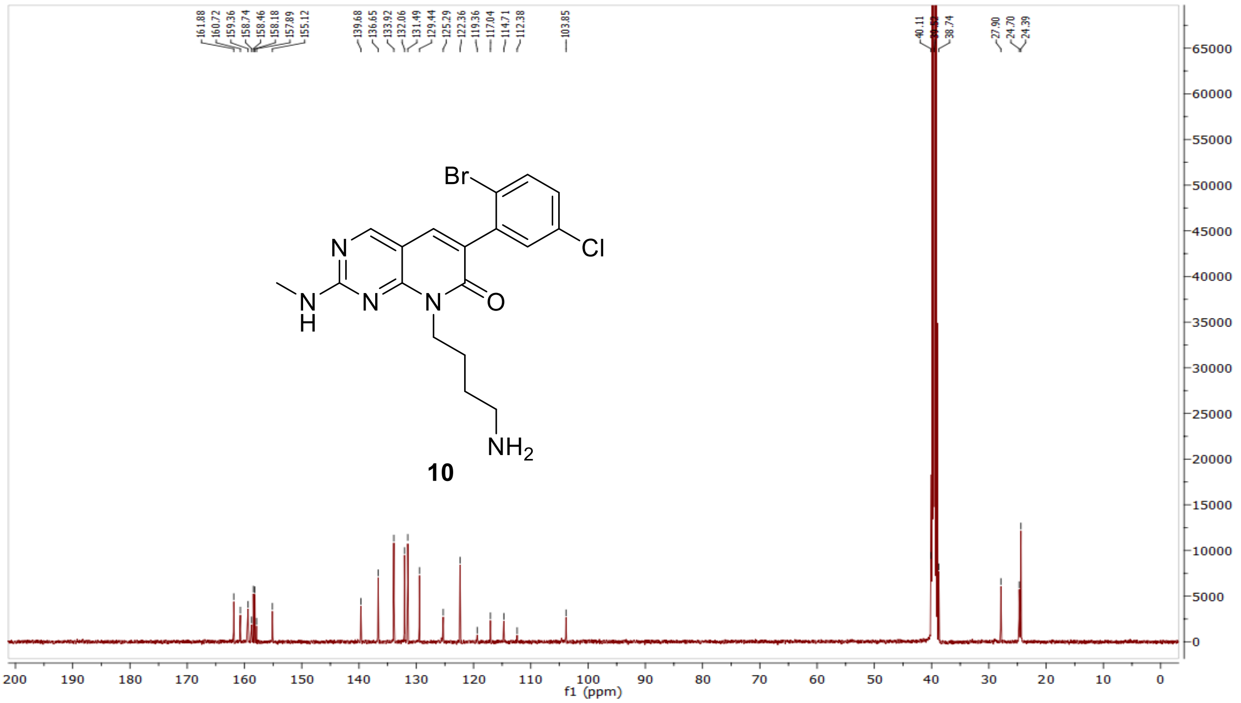

HRMS (FTMS +p MALDI) compound **10**:

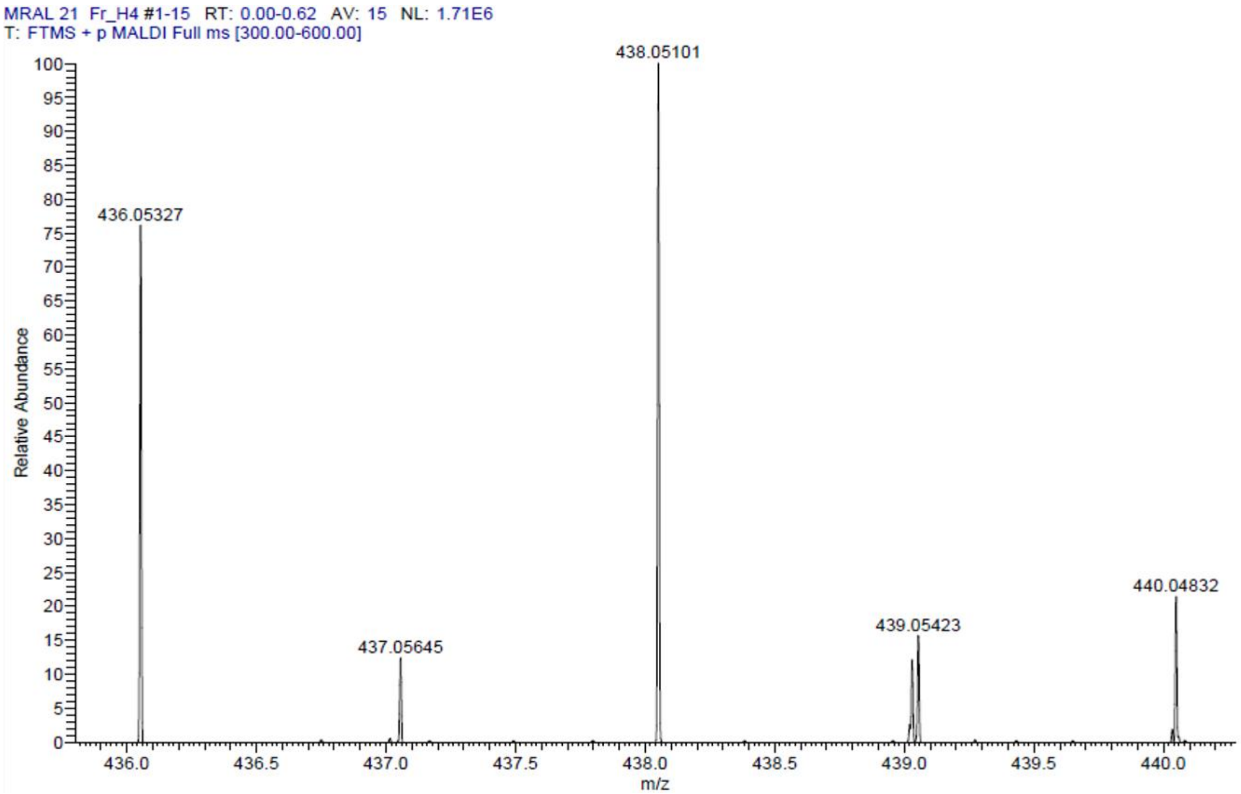

HPLC-chromatograms compound **10**:

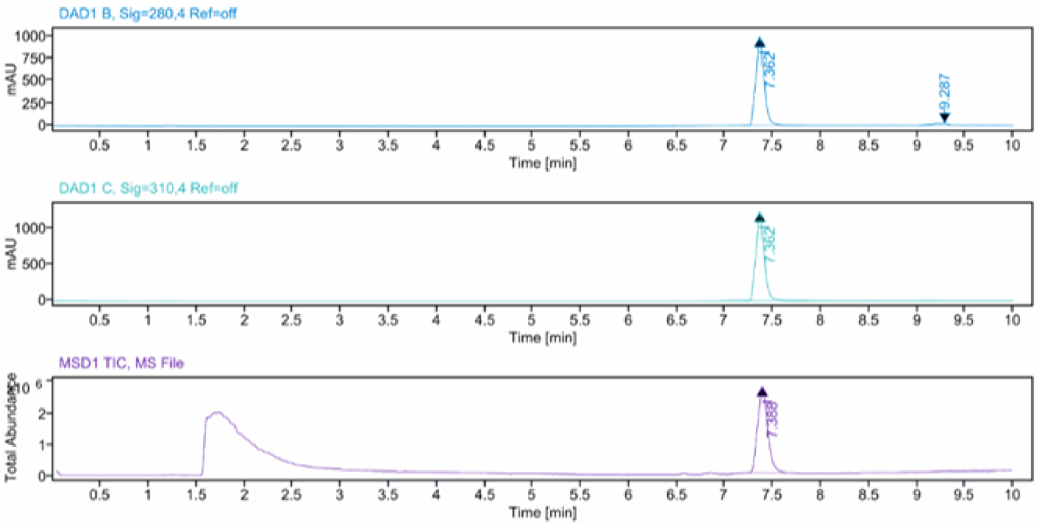


**Sample Purity**

Signal Description DAD1 B, Sig=280,4 Ref=off

| Sample Name | Name | RT | Width | Area | Area% | Height |
|---|---|---|---|---|---|---|
| MRAL21 Fr.1 HPLC | | 7.362 | 0.087 | 5833.6270 | 95.51 | 986.1207 |
| MRAL21 Fr.1 HPLC | | 9.287 | 0.191 | 274.5393 | 4.49 | 20.0200 |

Max Area% 95.505

UV Signal Purity>95% Pass

Signal Description DAD1 C, Sig=310,4 Ref=off

| Sample Name | Name | RT | Width | Area | Area% | Height |
|---|---|---|---|---|---|---|
| MRAL21 Fr.1 HPLC | | 7.362 | 0.087 | 7316.1392 | 100.00 | 1238.5265 |

Max Area% 100.000

UV Signal Purity>95% Pass

**MSD Apex Spectra**

**RT:**

7.388

**Sample Name:**

MRAL21 Fr.1 HPLC

**Signal Name:**

MSD1TIC

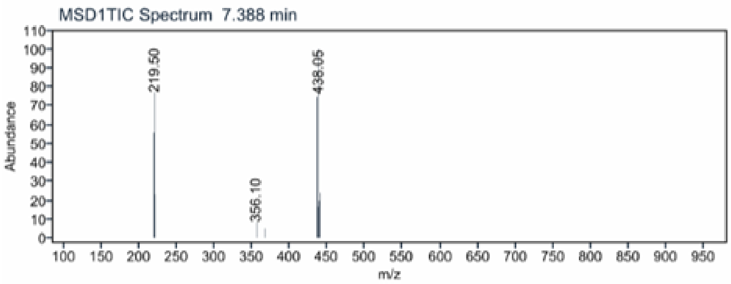

[1]H NMR compound **11**:

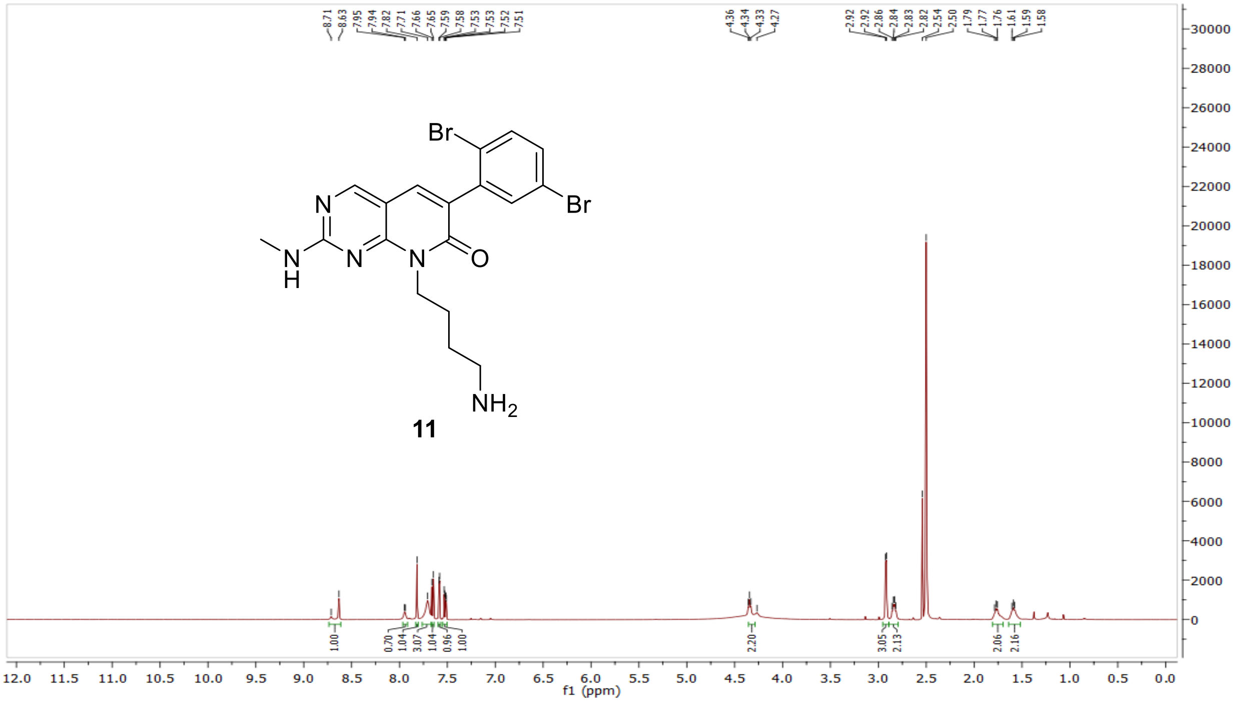

[13]C NMR compound **11:**

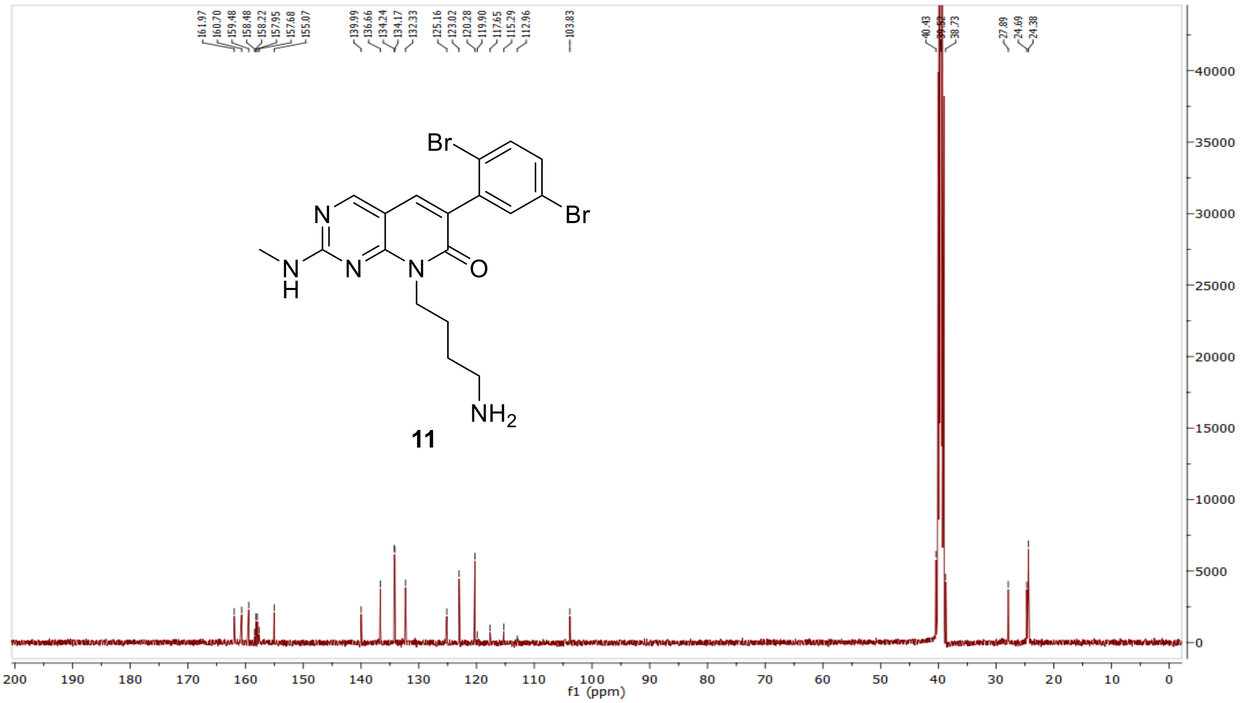

HRMS (FTMS +p MALDI) compound **11**:

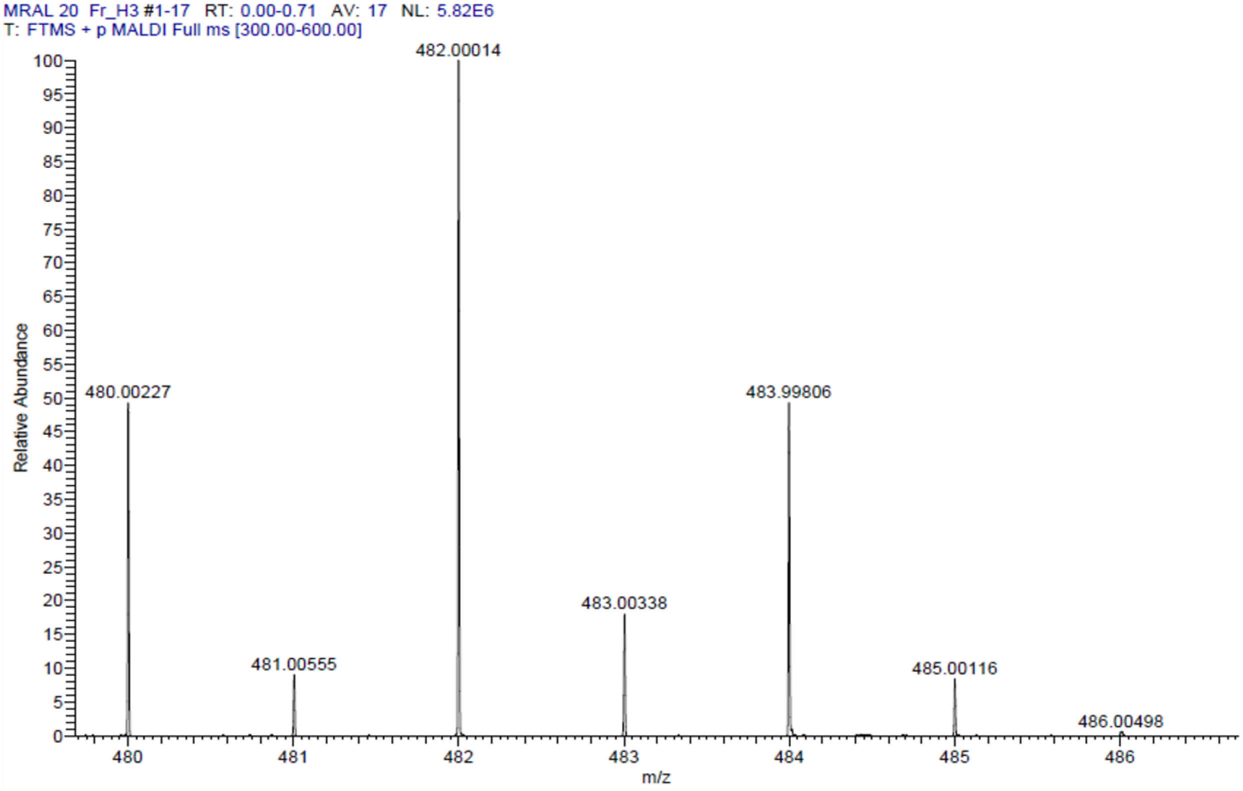

HPLC-chromatograms compound **11:**

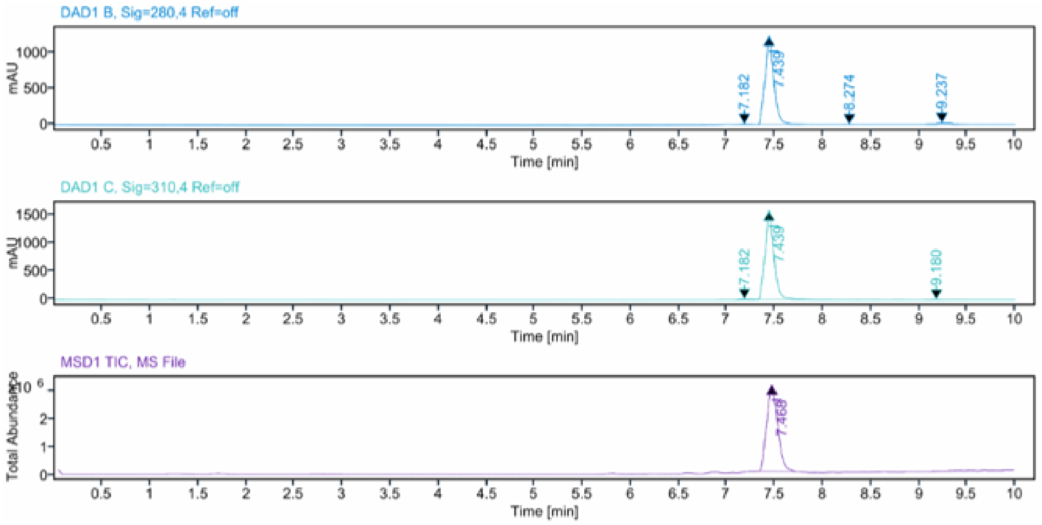


**Sample Purity**

Signal Description DAD1 B, Sig=280,4 Ref=off

| Sample Name | Name | RT | Width | Area | Area% | Height |
|---|---|---|---|---|---|---|
| MRAL20 Fr.1 HPLC | | 7.182 | 0.140 | 54.2987 | 0.67 | 7.6183 |
| MRAL20 Fr.1 HPLC | | 7.439 | 0.094 | 7767.8643 | 95.78 | 1231.5923 |
| MRAL20 Fr.1 HPLC | | 8.274 | 0.143 | 27.8501 | 0.34 | 4.1107 |
| MRAL20 Fr.1 HPLC | | 9.237 | 0.192 | 260.2716 | 3.21 | 19.9874 |

Max Area% 95.778

UV Signal Purity>95% Pass

Signal Description DAD1 C, Sig=310,4 Ref=off

| Sample Name | Name | RT | Width | Area | Area% | Height |
|---|---|---|---|---|---|---|
| MRAL20 Fr.1 HPLC | | 7.182 | 0.130 | 56.6384 | 0.57 | 8.2008 |
| MRAL20 Fr.1 HPLC | | 7.439 | 0.094 | 9933.9531 | 99.32 | 1569.8225 |
| MRAL20 Fr.1 HPLC | | 9.180 | 0.133 | 11.3975 | 0.11 | 1.2712 |

Max Area% 99.320

UV Signal Purity>95% Pass

**MSD Apex Spectra**

**RT:**

7.468

**Sample Name:**

MRAL20 Fr.1 HPLC

**Signal Name:**

MSD1TIC

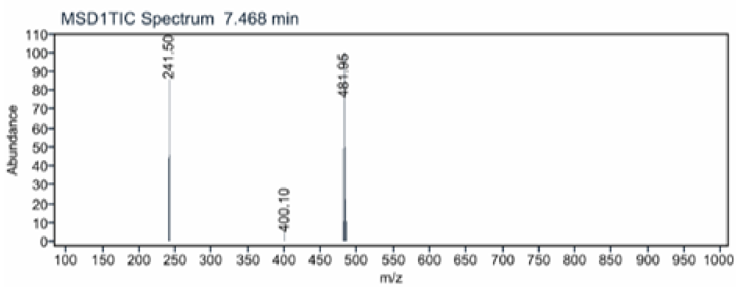

[1]H NMR compound **41**:

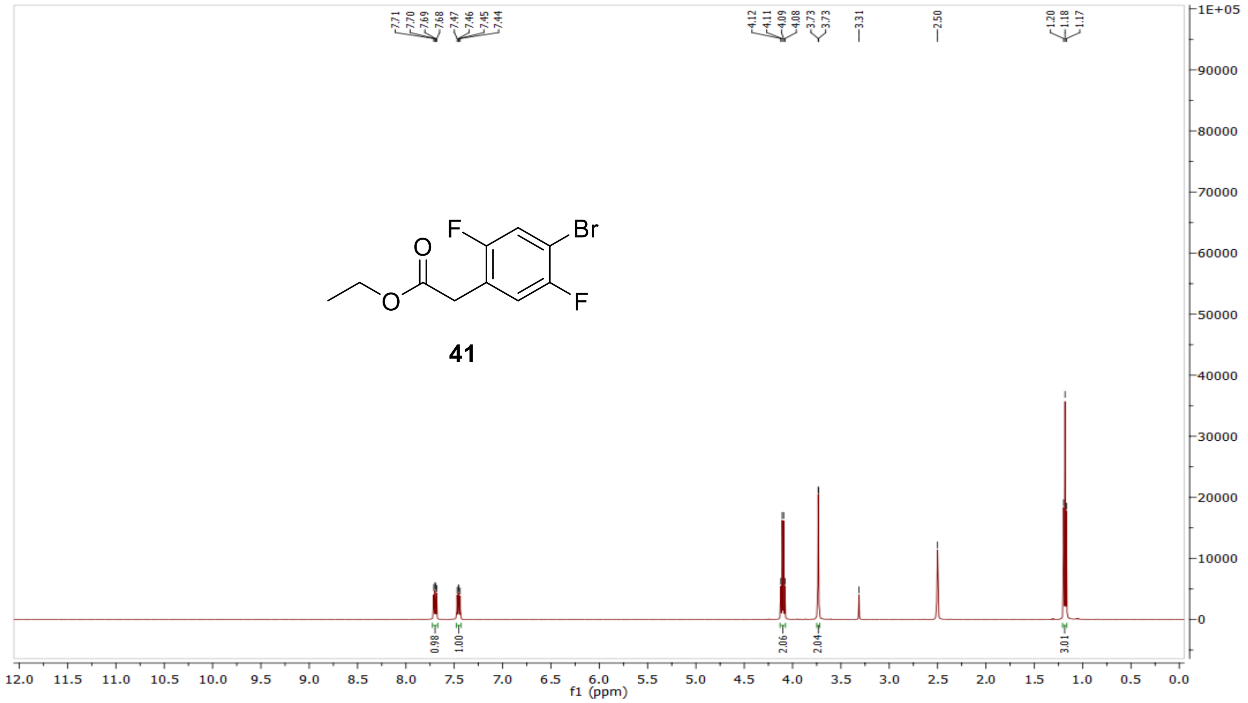


[13]C NMR compound **41:**

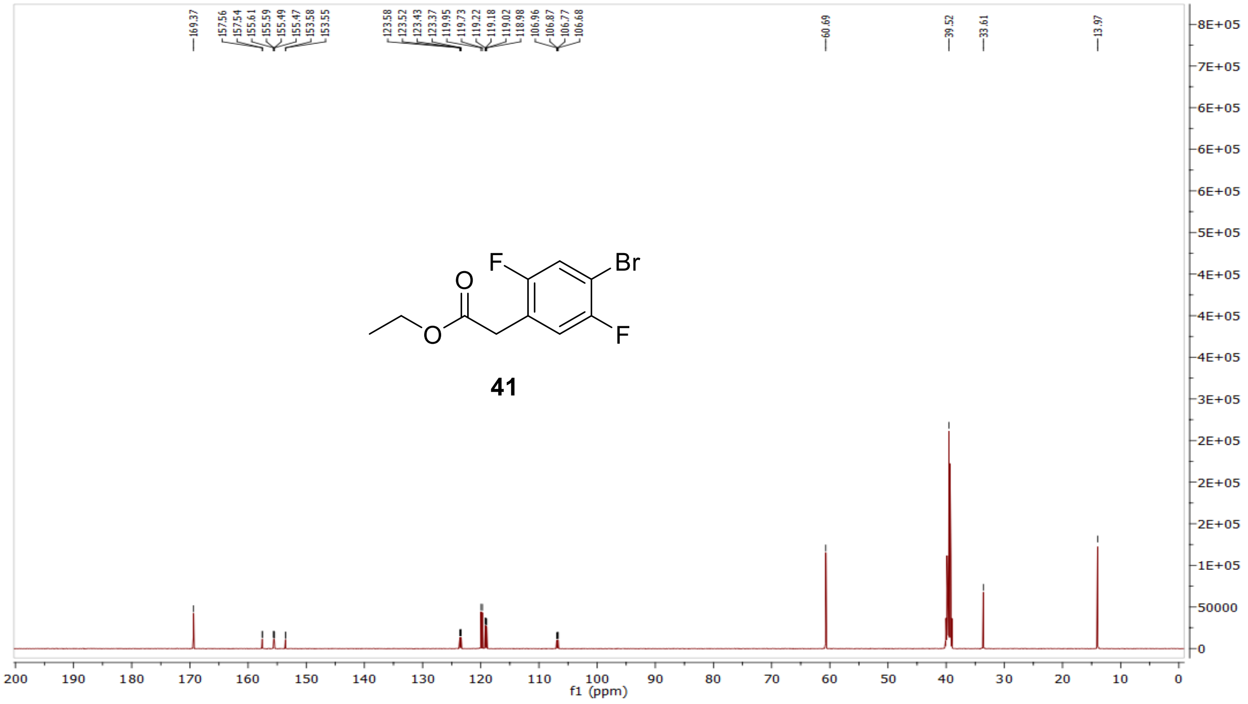

MS (ESI+) compound **41**:

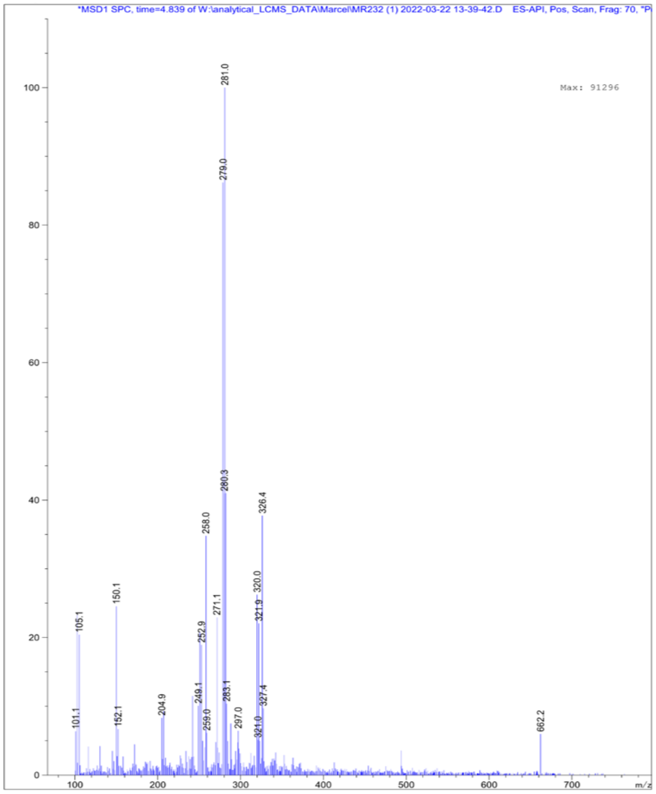

[1]H NMR compound **42**:

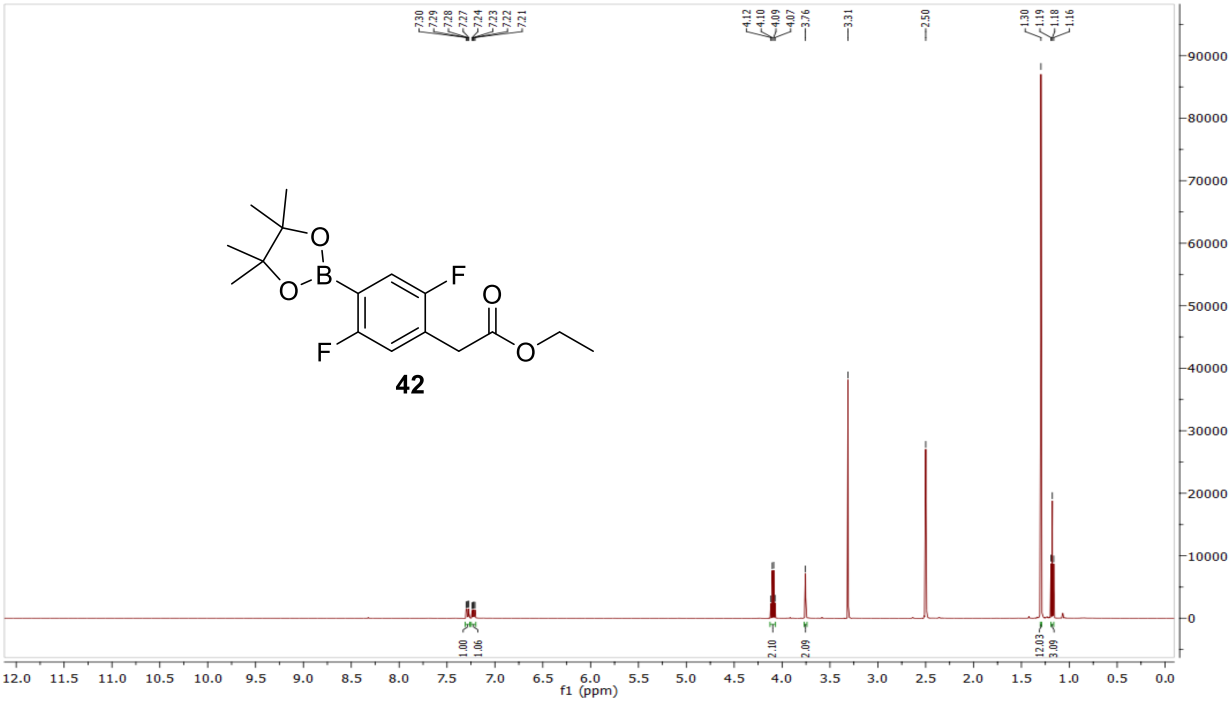

$^{13}C$ NMR compound **42**:

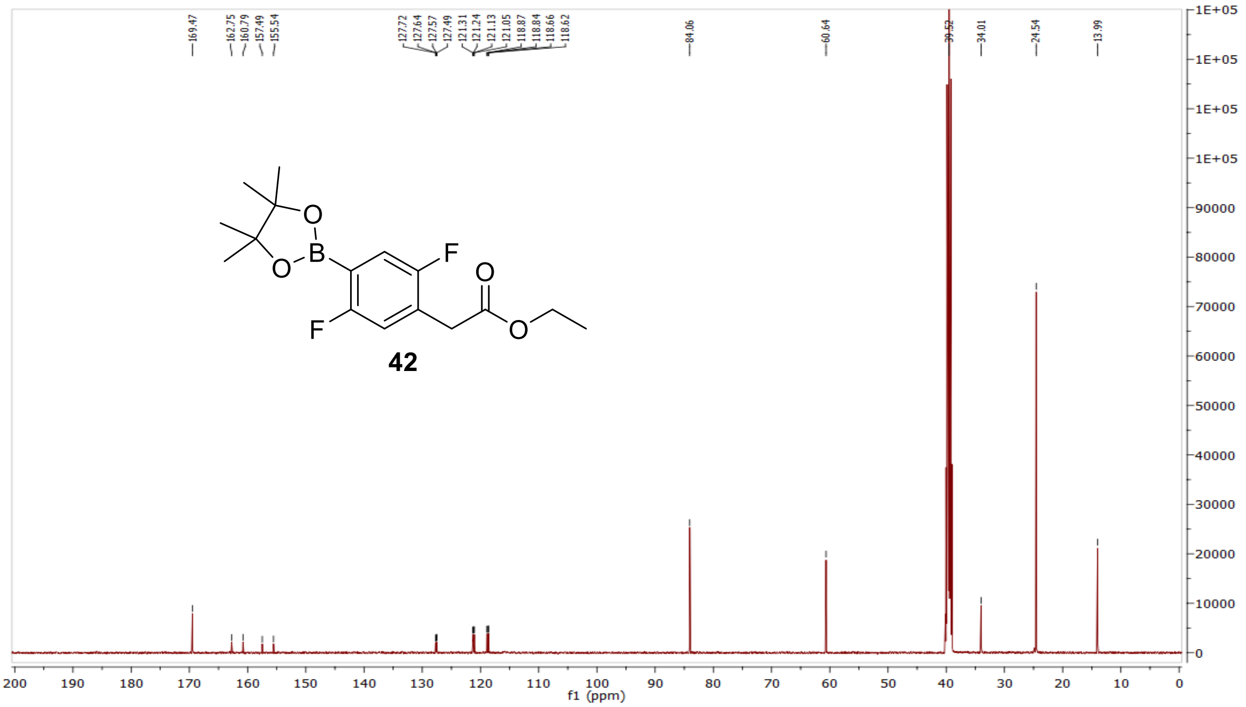

MS (ESI+) compound **42**:

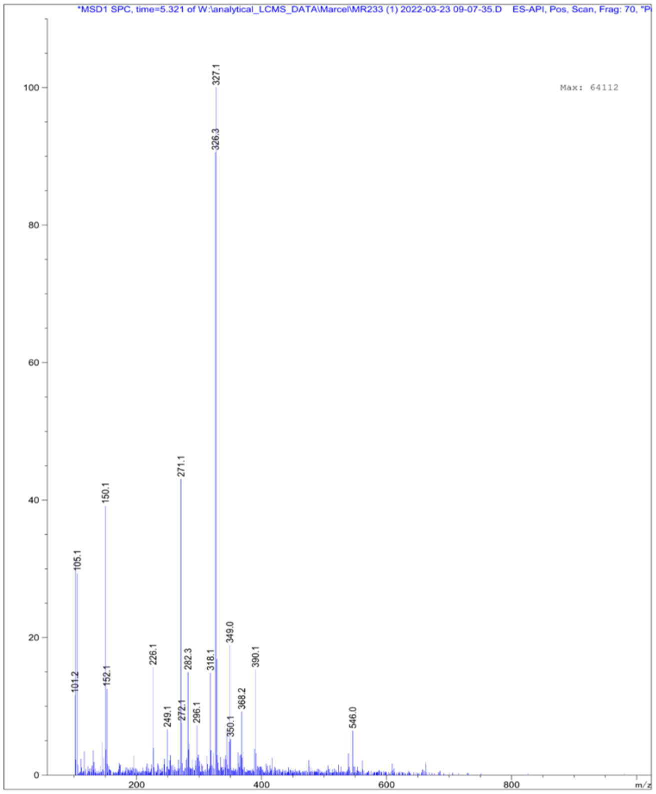

$^{1}H$ NMR compound **43**:

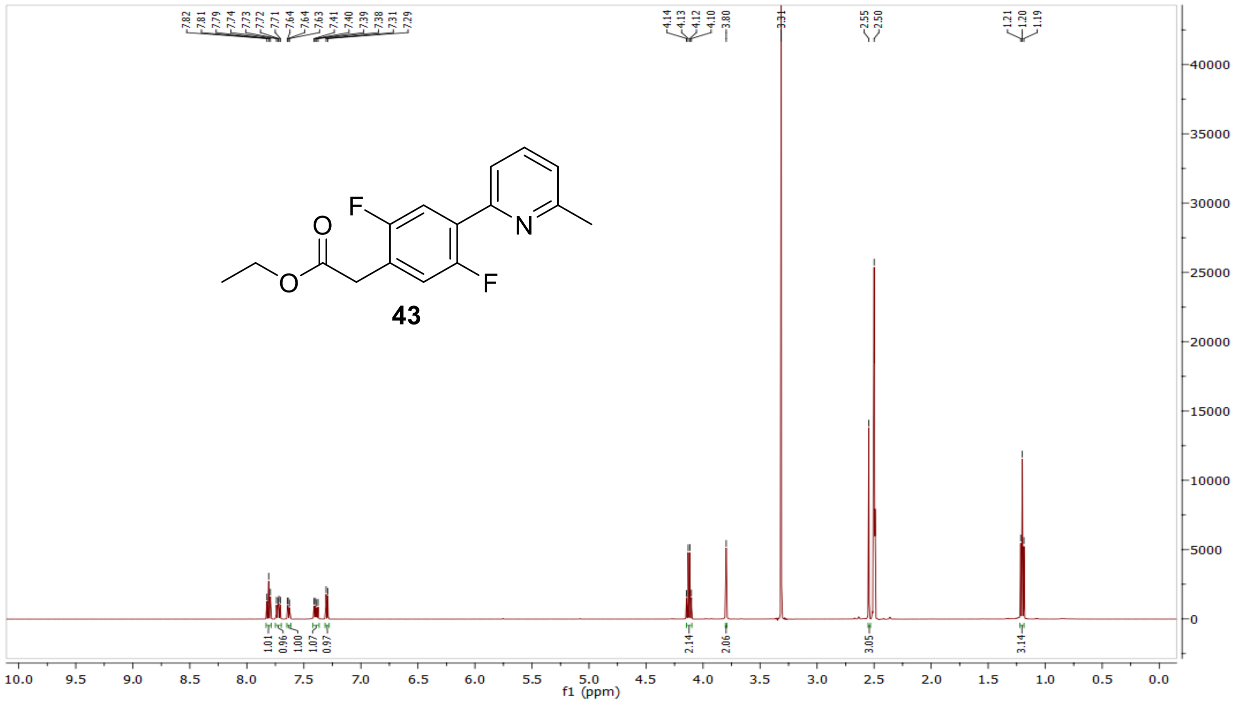

$^{13}C$ NMR compound **43**:

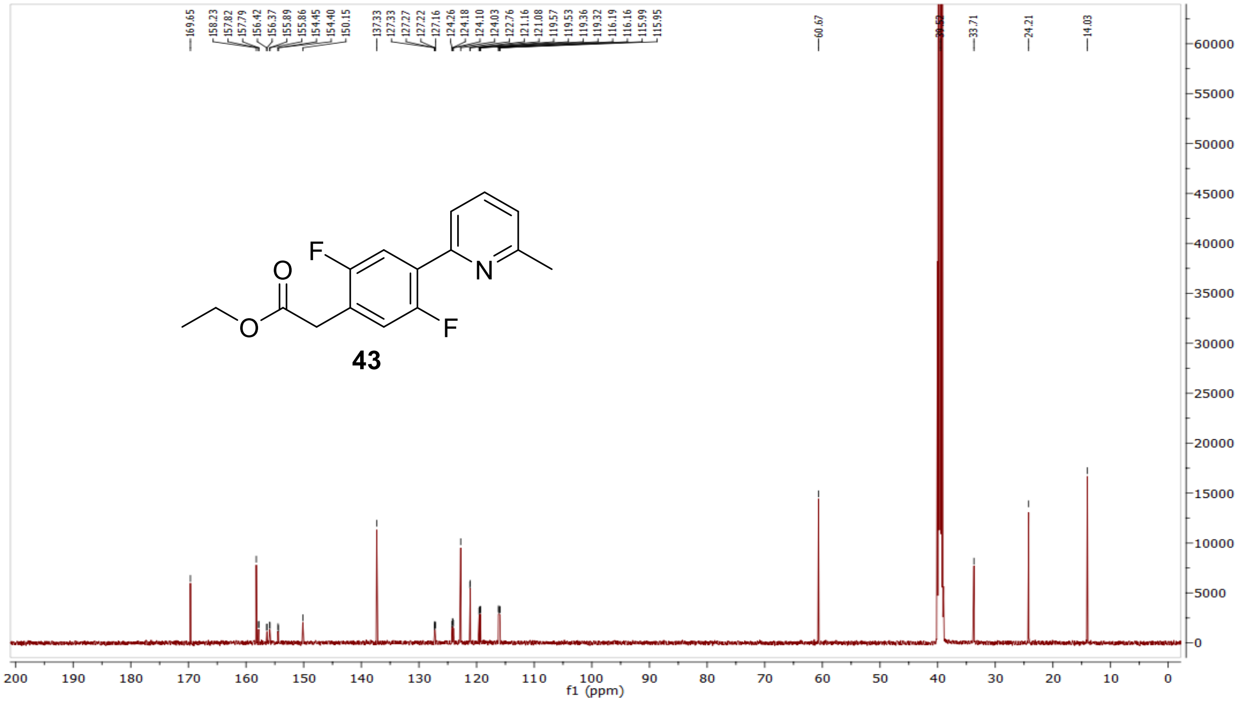

MS (ESI+) compound **43**:

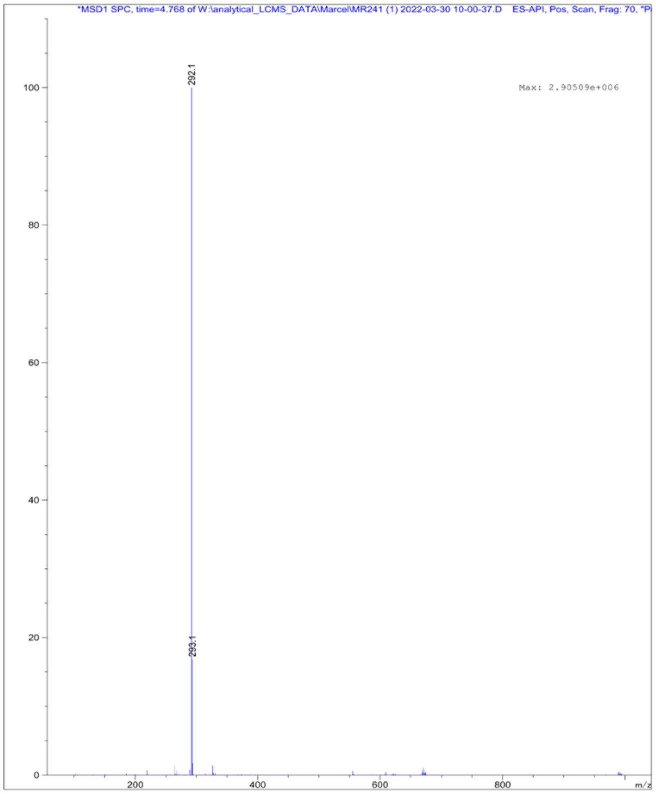

$^1$H NMR compound **44**:

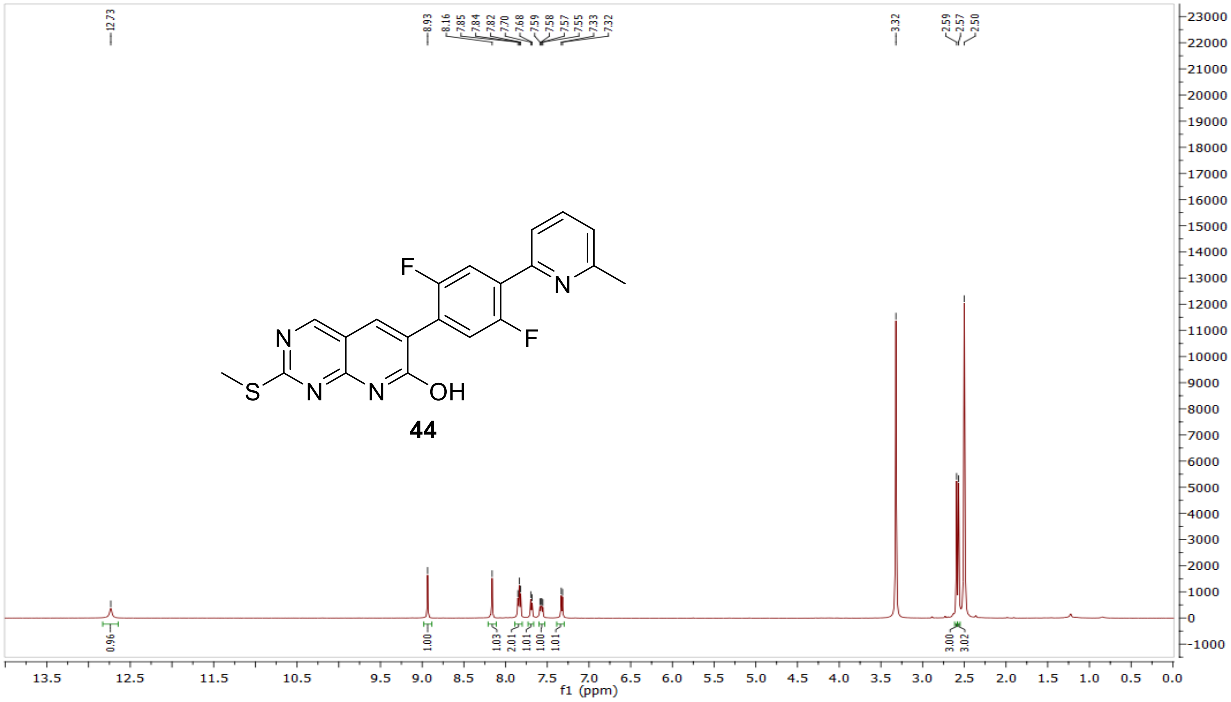

$^{13}$C NMR compound **44:**

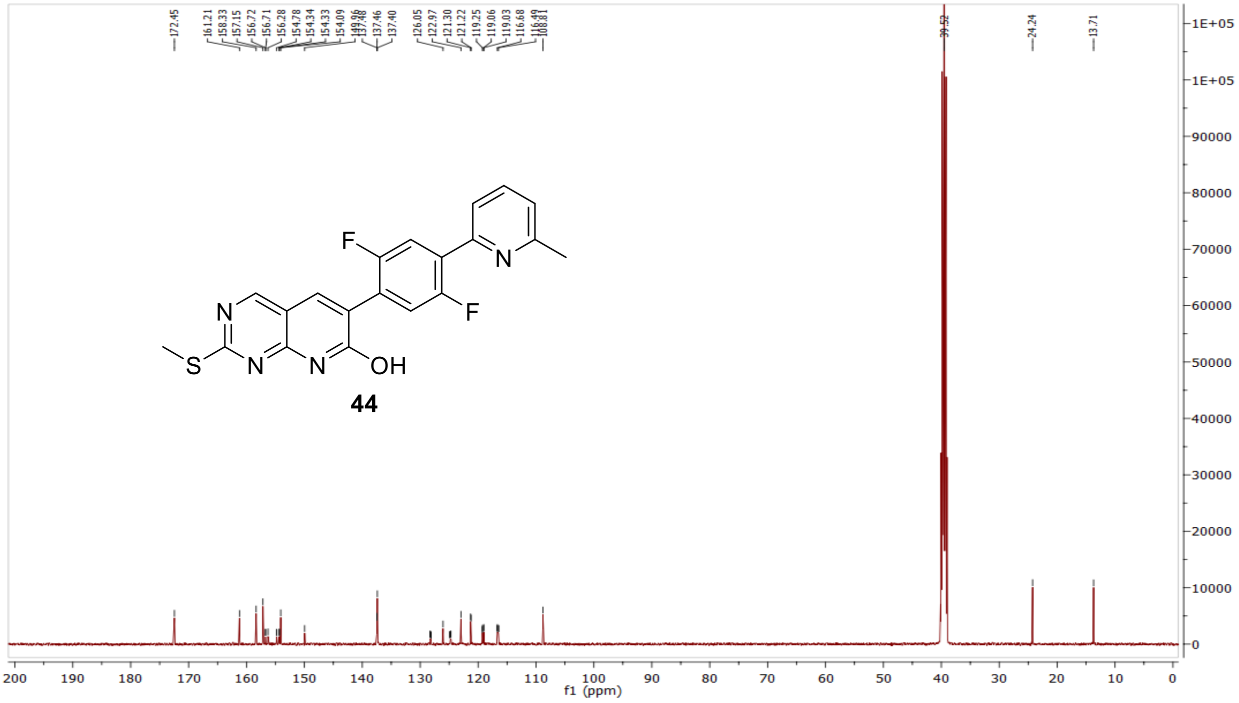


MS (ESI+) compound **44**:

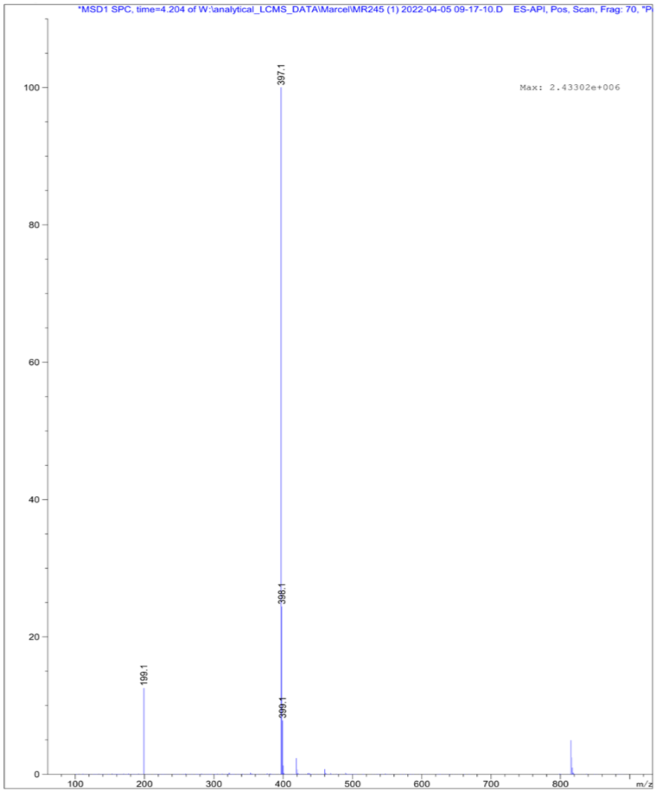

$^{1}$H NMR compound **45**:

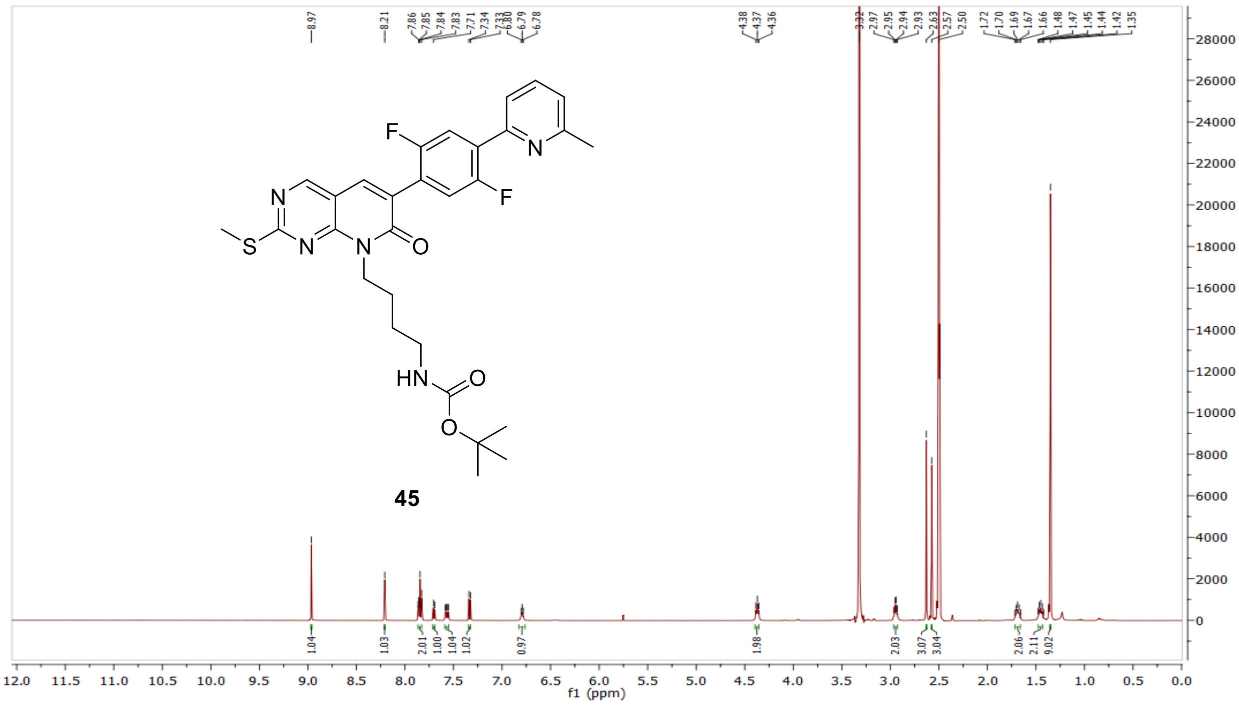

$^{13}$C NMR compound **45:**

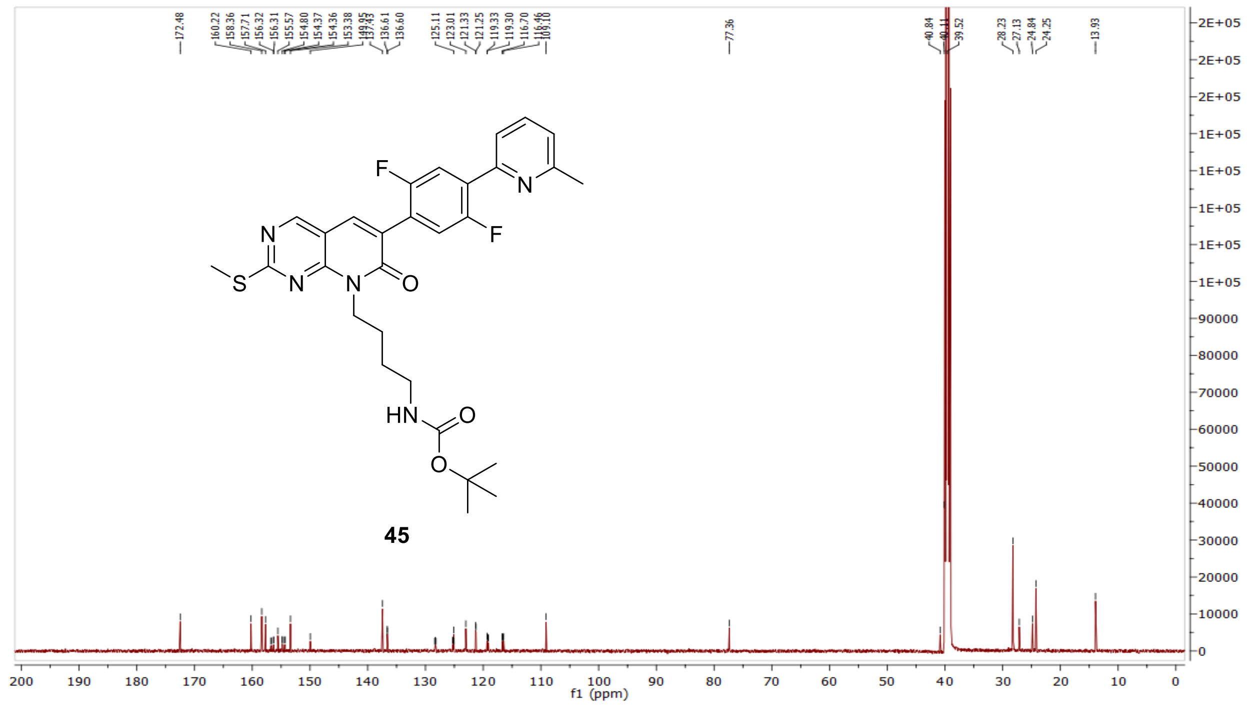

MS (ESI+) compound **45**:

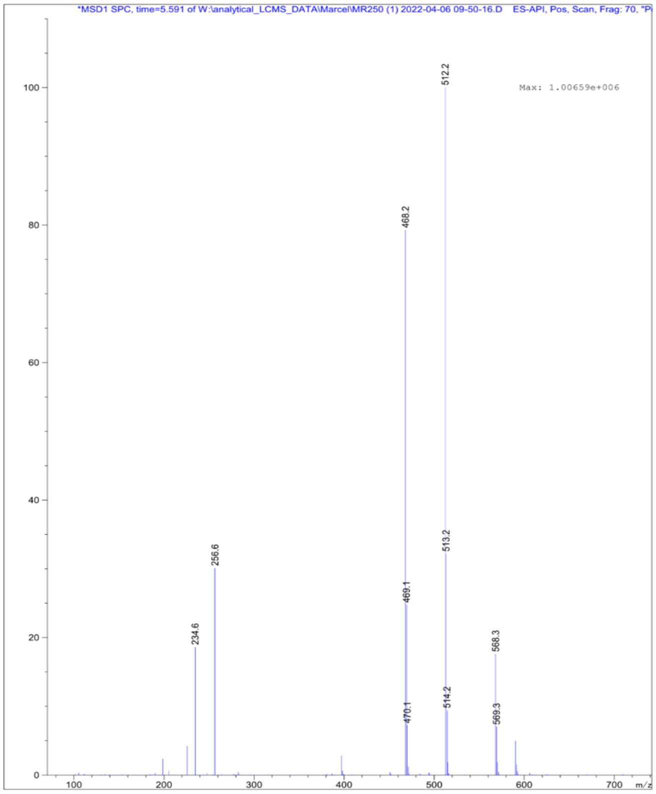

[1]H NMR compound **46**:

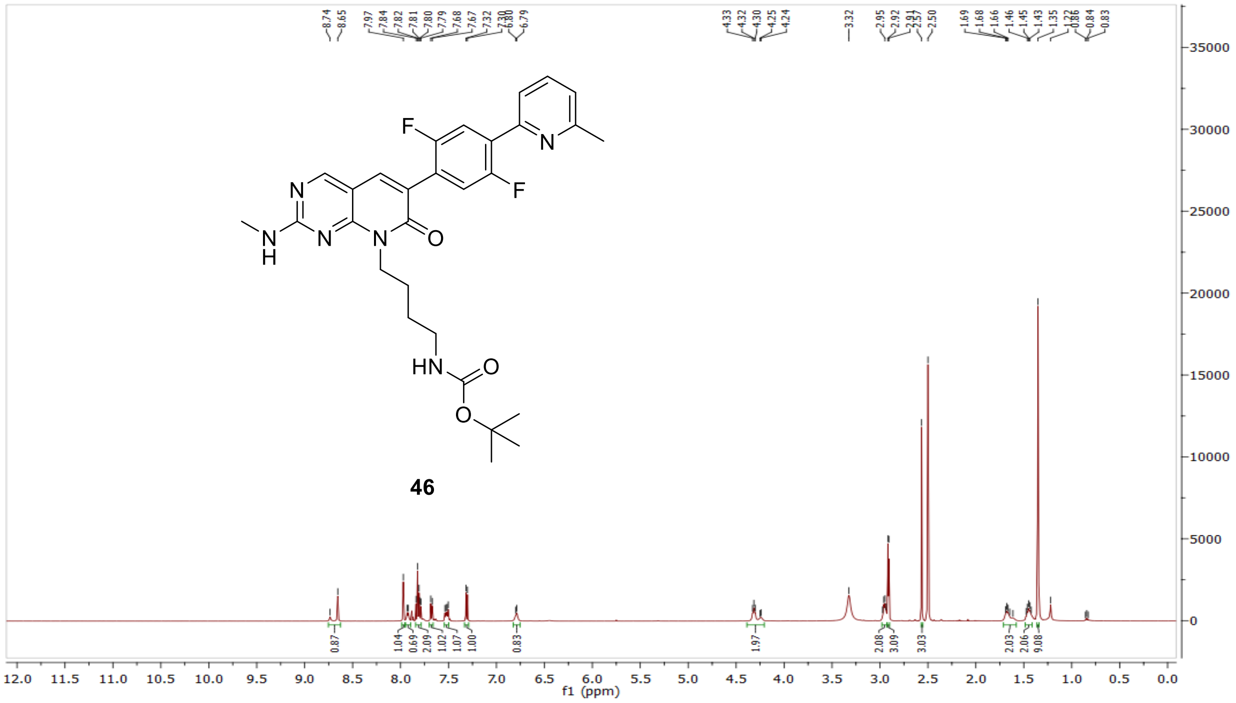

$^{13}$C NMR compound **46:**

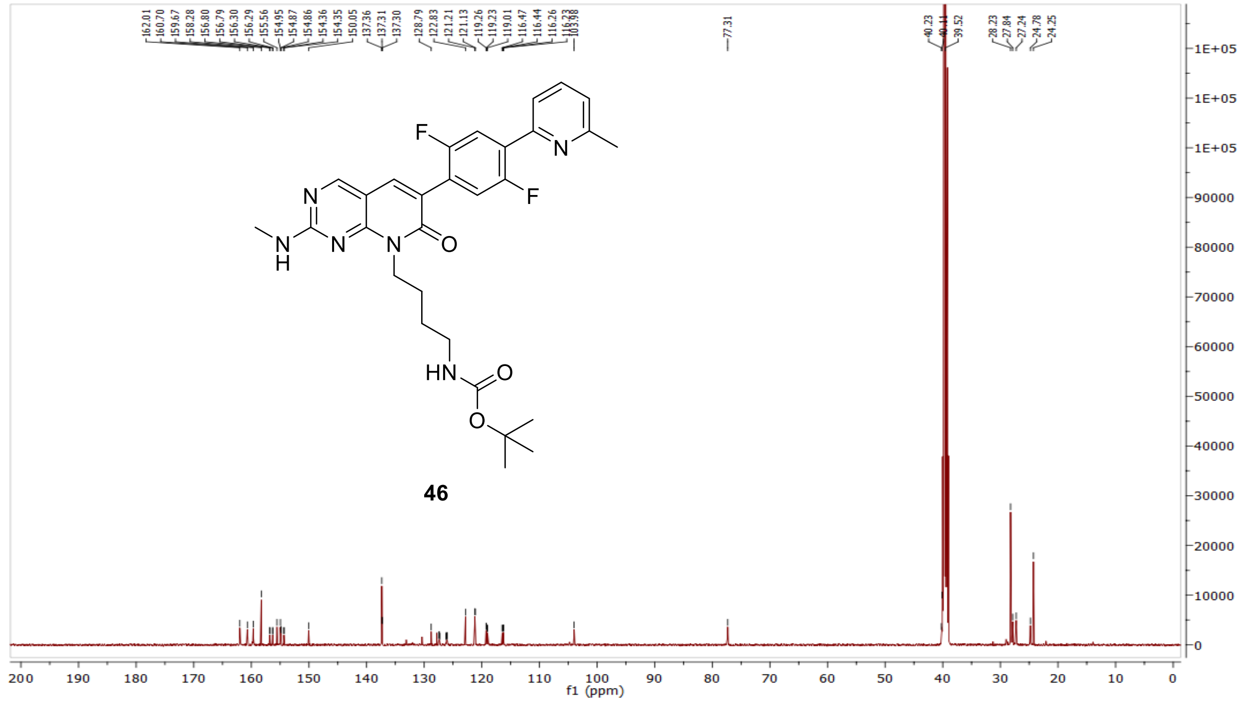


MS (ESI+) compound **46**:

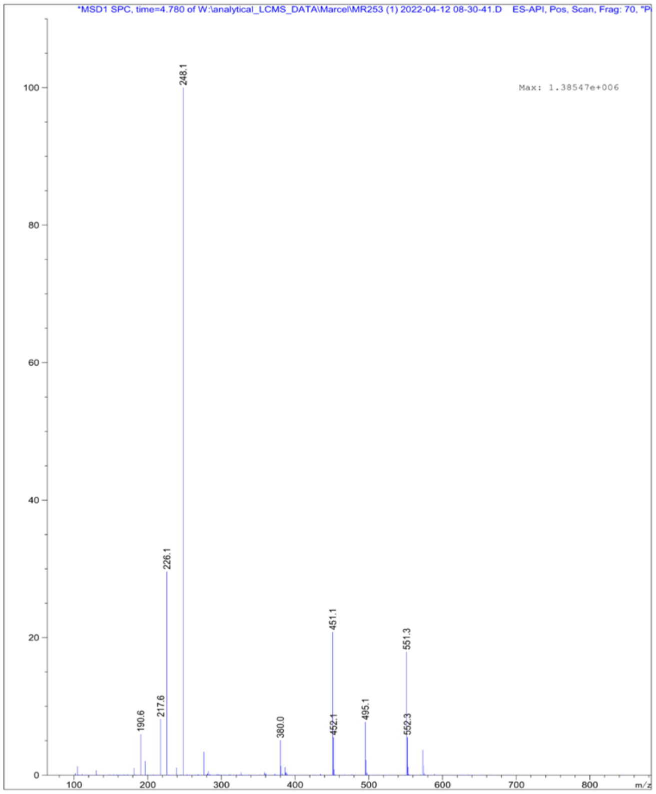

$^{1}H$ NMR compound **12**:

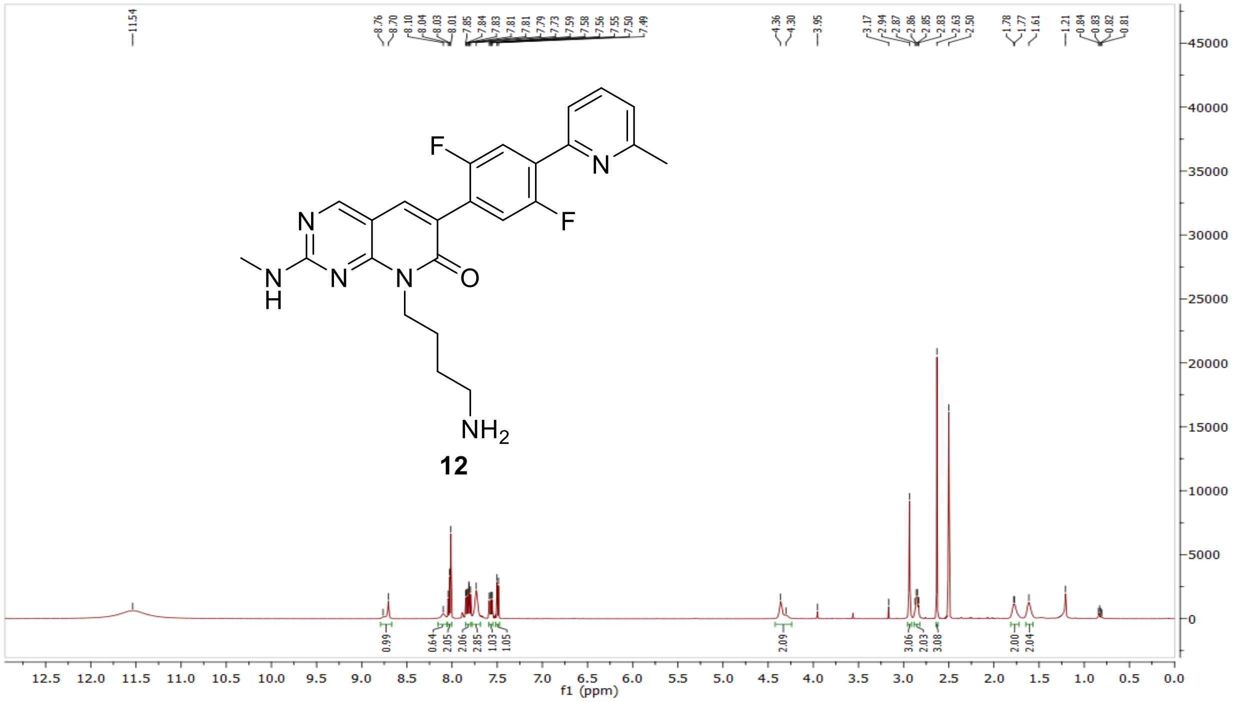

$^{13}C$ NMR compound **12:**

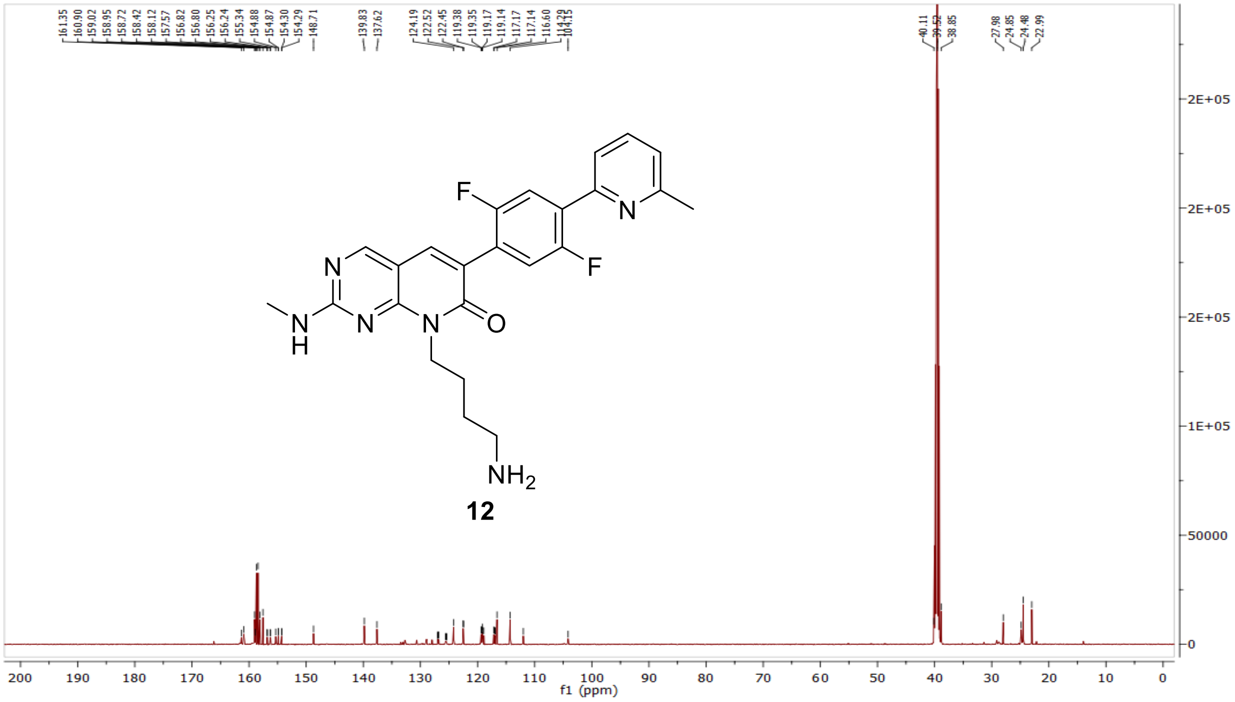

HRMS (FTMS +p MALDI) compound **12**:

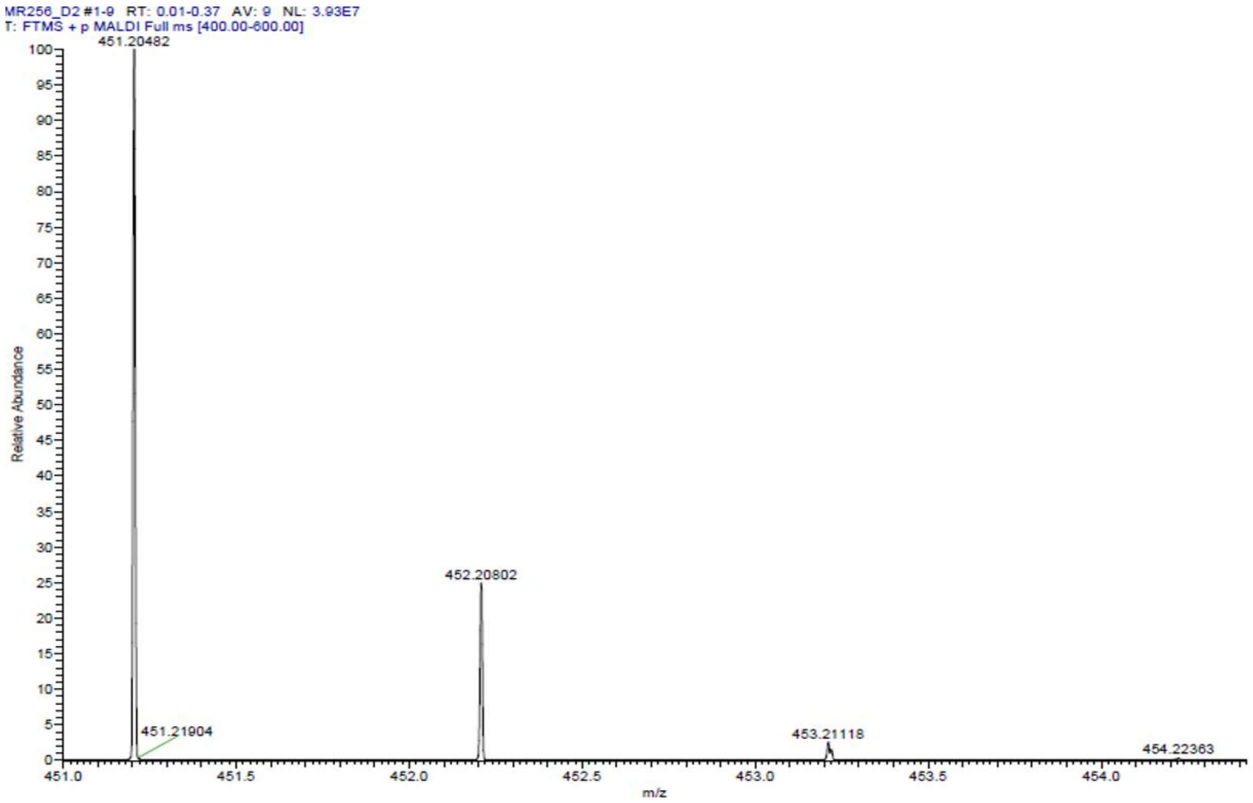

HPLC-chromatograms compound **12**:

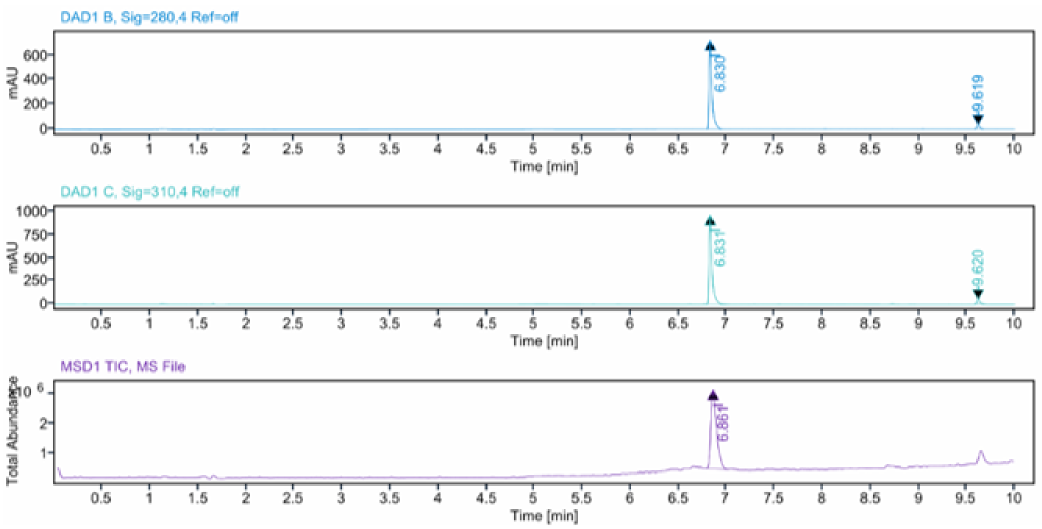


**Sample Purity**

Signal Description DAD1 B, Sig=280,4 Ref=off

| Sample Name | Name | RT | Width | Area | Area% | Height |
|---|---|---|---|---|---|---|
| MR256 | | 6.830 | 0.028 | 1474.1193 | 95.64 | 719.2051 |
| MR256 | | 9.619 | 0.034 | 67.2047 | 4.36 | 30.2529 |

Max Area% 95.640

UV Signal Purity>95% Pass

Signal Description DAD1 C, Sig=310,4 Ref=off

| Sample Name | Name | RT | Width | Area | Area% | Height |
|---|---|---|---|---|---|---|
| MR256 | | 6.831 | 0.028 | 1976.5155 | 95.35 | 961.3381 |
| MR256 | | 9.620 | 0.033 | 96.4102 | 4.65 | 44.9142 |

Max Area% 95.349

UV Signal Purity>95% Pass

**MSD Apex Spectra**

**RT:**

6.861

**Sample Name:**

MR256

**Signal Name:**

MSD1TIC

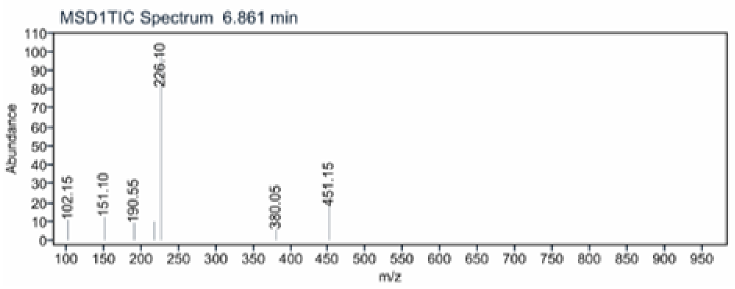
